# Breast Milk Oligosaccharides Contain Immunomodulatory Glucuronic Acid and LacdiNAc

**DOI:** 10.1101/2023.01.16.524336

**Authors:** Chunsheng Jin, Jon Lundstrøm, Emma Korhonen, Ana S. Luis, Daniel Bojar

## Abstract

Breast milk is abundant with functionalized milk oligosaccharides (MOs), to nourish and protect the neonate. Yet we lack a comprehensive understanding of the repertoire and evolution of MOs across Mammalia. We report ∼400 MO-species associations (>100 novel structures) from milk glycomics of nine mostly understudied species: alpaca, beluga whale, black rhinoceros, bottlenose dolphin, impala, L’Hoest’s monkey, pygmy hippopotamus, domestic sheep, and striped dolphin. This revealed the hitherto unknown existence of the LacdiNAc motif (GalNAcβ1-4GlcNAc) in MOs of all species except alpaca, sheep, and striped dolphin, indicating widespread occurrence of this potentially antimicrobial motif in MOs. We also characterize glucuronic acid-containing MOs in the milk of impala, dolphins, sheep, and rhinoceros, previously only reported in cows. We demonstrate that these GlcA-MOs exhibit potent immunomodulatory effects. Our study extends the number of known MOs by >15%. Combined with >1,900 curated MO-species associations, we characterize MO motif distributions, presenting an exhaustive overview of MO biodiversity.

---

While every mammalian species produces breast milk to nurture its young, there is a pronounced diversity in breast milk composition^1^. This ranges from the evolutionarily ancient monotreme milk over marsupial milk, designed to nourish the prematurely born neonates in the pouch^2^, up to placental milk, including humans and their domestic animals. This broad range of needs and niches has to be recapitulated on the molecular level, to endow the respective milk with its required functionality, adjusted during the time after parturition.

Regardless of milk origin, complex carbohydrates are usually among the most abundant components of breast milk^3,4^. Composed of monosaccharides arranged in branching chains, these milk oligosaccharides (MOs) exhibit striking diversity (Fig. 1). Many aspects of infant nutrition^5^, immune system development^6^, microbiome development^7^, and pathogen defense^8^ have all been directly tied to MOs and their biochemical properties. The initial function of MOs seemed to be of defensive nature, with a subsequent expansion into nutrition^9^. Due to this plethora of functions, the relative absence of complex MOs from infant formula harbors consequences for infant health^10,11^.

**Figure 1.**
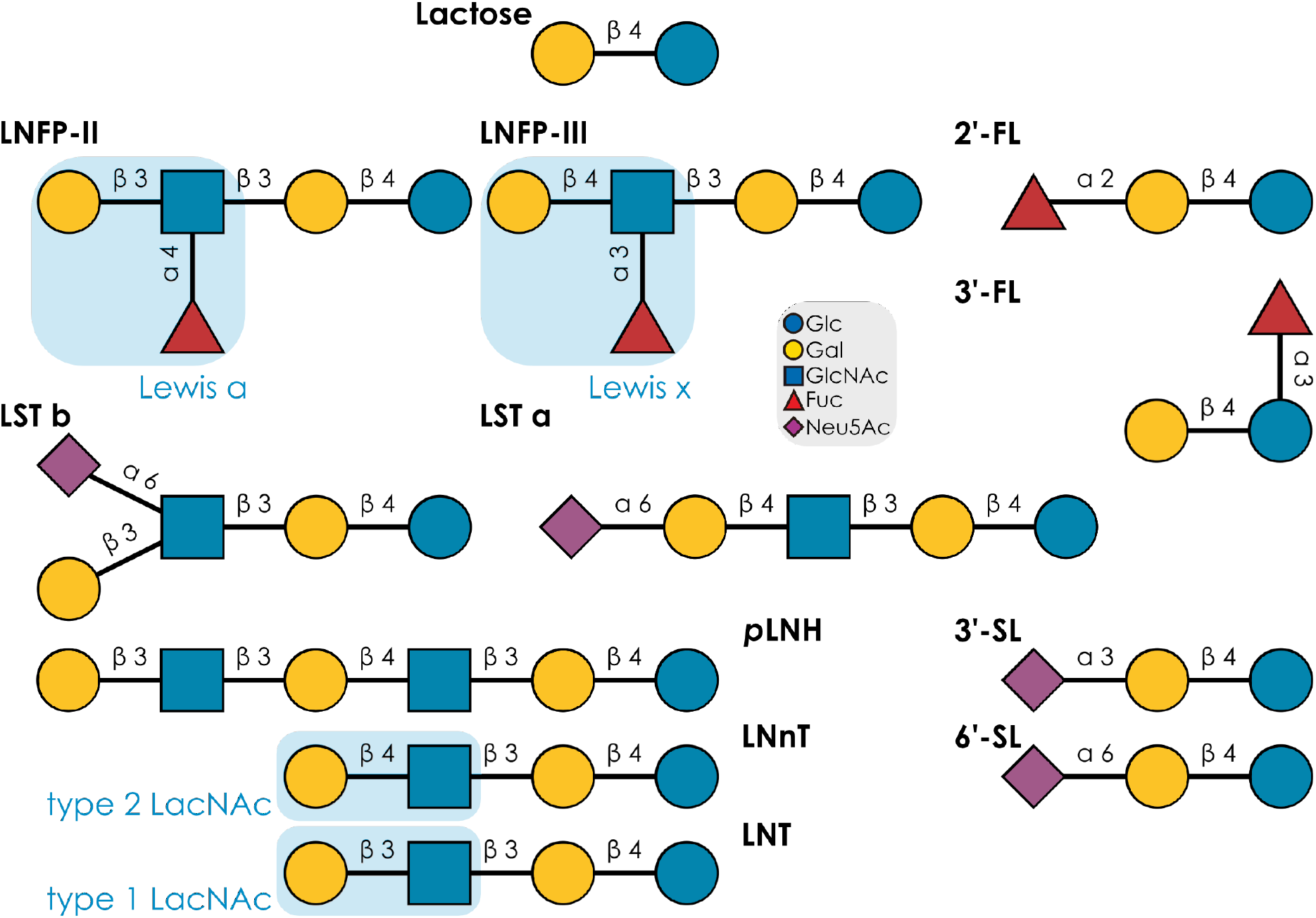
Typical structures and motifs in free milk oligosaccharides. Examples of common milk oligosaccharide structures, with their commonly used name abbreviations and highlighted motifs, are shown via the Symbol Nomenclature For Glycans (SNFG).

In humans, hundreds of unique MOs have been described^12^. While this is often used to imply human exceptionalism, milk from well-investigated domestic animals, such as cows or pigs, exhibit similar orders of magnitude in terms of unique MO structures^13,14^. More importantly, the milk glycomes of different species are characteristic^15–17^, supplying a molecular repertoire to fulfill the functions of breast milk in that species. Usually, motifs within whole glycan structures are functional determinants^18^. Therefore, (species-specific) sequence-to-function relationships exist in MOs and are strictly limited by the available motif repertoire.

These MO repertoires are produced in the mammary gland by an array of glycosyltransferases, enzymes starting from lactose or lactosamine in the Golgi apparatus and consecutively adding monosaccharides to the non-reducing end^19^. While MO biosynthesis, especially in non-human species, is still incompletely resolved, an emerging consensus describes evolutionarily conserved MO repertoire patterns in various taxonomic groups^20,21^. A classic example can be found in marsupials, with characteristic oligo-galactose motifs in their MOs that are not found in that form and abundance in other mammalian groups.

However, only a small fraction of mammalian species has been characterized regarding their MOs, with a particular emphasis on humans and domestic animals / model organisms. Further, due to the structural complexity of MOs, studies investigating MO repertoires that rely on comparison to standards or regular mass spectrometry might be restricted to measuring only simpler / already characterized MOs, as pointed out before^12,22^. Particularly for characterizing new MO structures, tandem mass spectrometry, including probing with specific exoglycosidases, is often necessary^23^, yet infrequently applied when characterizing non-model species. This underestimates the true biodiversity of MOs throughout Mammalia.

Here, we use structural glycomics via liquid chromatography – tandem mass spectrometry (LC-MS/MS) to characterize the milk glycomes of nine species that have been barely, or not at all, investigated regarding their MOs. We identify 393 MO-species associations in this endeavor, including 108 glycan sequences that have not been described before (Supplementary Table 1), out of 172 characterized sequences. We also present the discovery of new motifs for MOs, such as the LacdiNAc motif (GalNAcβ1-4GlcNAc), expanding the known motif repertoire for MOs. We further discover several glucuronic acid (GlcA)-containing MOs in multiple species, previously only identified in cows in recent work^24^. Next to expanding the motif repertoire, we therefore also broaden the monosaccharide repertoire in MOs. On human immune cells, these GlcA-MOs demonstrate immunomodulatory effects that seem more potent than established immunomodulators such as 2-fucosyllactose (2’-FL). We contextualize our findings within a newly curated dataset of 1,902 MO observations in >100 species, constituting an exhaustive overview of our current knowledge about MO biodiversity. Engaging in various sequence and motif analyses, we reveal clusters of motif usage in mammalian MOs. We envision our findings to substantially extend the known biodiversity of MOs and improve our understanding of biochemical properties and functions of these important constituents of breast milk.

## Results

### Exploring the free milk glycome of uncharacterized mammals

Our extensive literature curation yielded MO data (beyond merely lactose) for 106 species. However, estimates^25^ place the total number of mammalian species at about 6,500, suggesting that we only have information for ∼1% of all milk glycomes, and, even where we do have some information, we lack the exhaustiveness applied to human or cow MOs. We thus decided to analyze the free milk oligosaccharides of uncharacterized or undercharacterized mammalian species, using an in-depth structural glycomics approach, to improve overall biodiversity comparisons across Mammalia and expand our knowledge of this important nutrient.

The availability of breast milk from exotic mammalian species is, however, rather limited, which determined our species choice in collaborations with European zoological institutions. This resulted in milk from nine species (seven Artiodactyla, one Perissodactyla, one Primates), with all samples constituting mature milk (n = 11), and one additional sample of colostrum and pre-milk each. Most of these species had zero previously reported glycans in general and thus add to our knowledge of biodiversity in glycobiology.

### LacdiNAc is widespread in MOs of Artiodactyla species

In alpaca milk (*Lama pacos*, Fig. 2a; not studied previously), we detected MOs that were sulfated on galactose or *N*-acetylglucosamine (GlcNAc), including extended structures such as Neu5Acα2-6Galβ1-4GlcNAc6Sβ1-3Galβ1-4Glc (Supplementary Dataset 1, slide 89). Unexpectedly, we only measured two fucosylated glycans, present in low abundance, while sialylated glycans (predominantly Neu5Acα2-6) were more common. This included unusual structures such as Neu5Acα2-3GlcNAcβ1-6Galβ1-4Glc (Supplementary Dataset 1, slide 50), with the rarely observed sialic acid-GlcNAc linkage. Particularly high-abundance MOs were the neutral LNnT (lacto-*N*-neotetraose) and LNnH (lacto-*N*-neohexaose), and the sialylated S-LNnH. Other investigated camelids, *Camelus bactrianus* and *Camelus dromedarius*, differ from this by exhibiting more fucosylated structures, including blood group epitopes^16^. Overlaps, such as S-LNnH in *C. bactrianus* and *C. dromedarius*^26,27^, might imply that further investigation of alpaca milk also uncovers these fucosylated MO motifs. We note that *L. pacos* seems to constitute the first example of sulfated MOs in the family of camelids.

**Figure 2.**
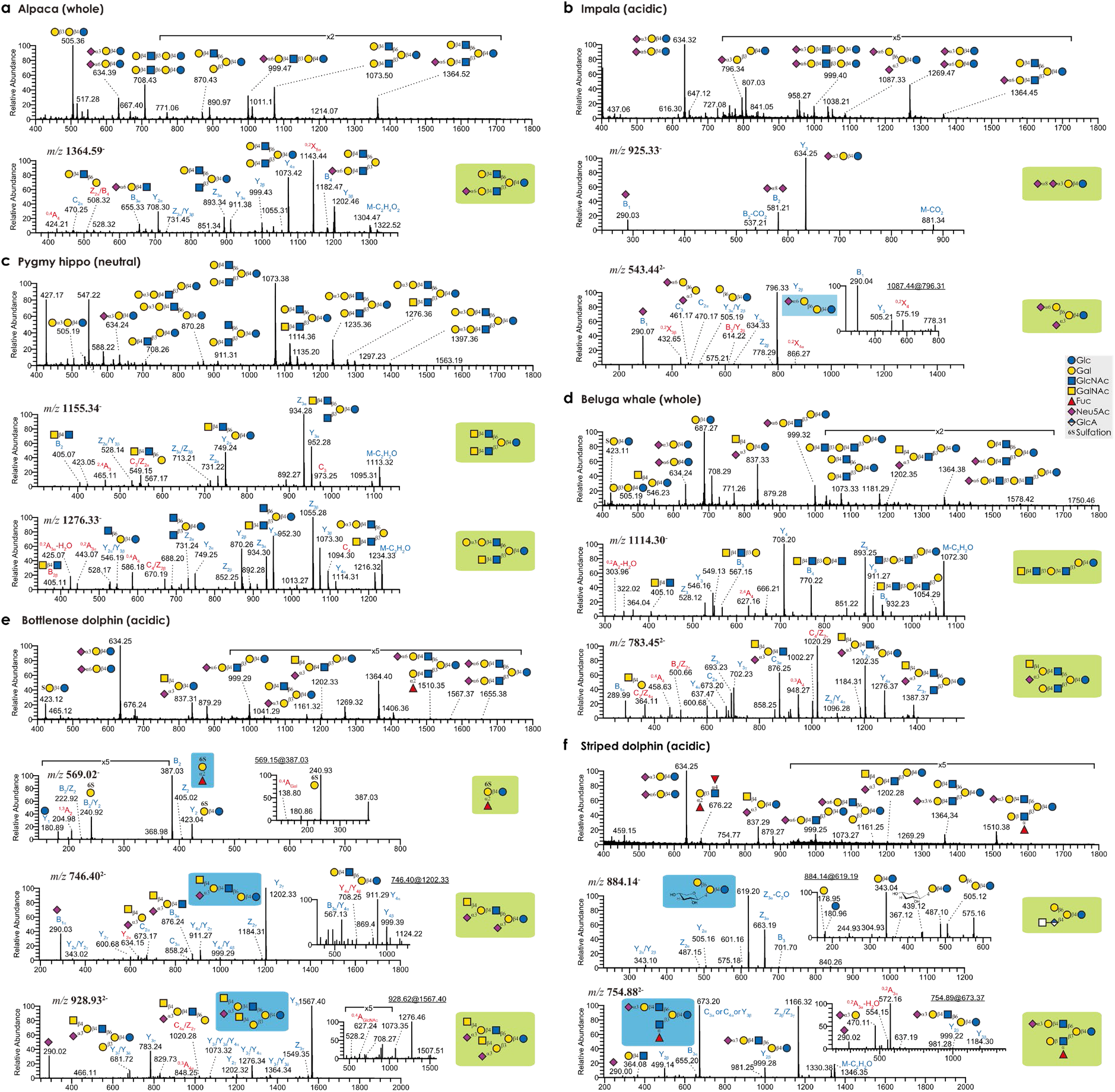
Free milk oligosaccharides from understudied Artiodactyla species. **a-f**, Annotated mass spectra of free MOs for newly investigated mammalian species. Shown are the milk glycans, identified via negative-ion mode LC-ESI-MS/MS, from *Lama pacos* (a), *Aepyceros melampus* (b), *Choeropsis liberiensis* (c), *Delphinapterus leucas* (d), *Tursiops truncatus* (e), *Stenella coeruleoalba* (f). The MS/MS and MS^3^ (insert) spectra of selected free MOs (structures shadowed in green or blue, respectively) from each species are annotated with selected fragmentation ions, which are degenerated by glycosidic or cross-ring cleavage. The C/Z cleavage of typical fragments is used to determine the branching unit (e.g., m/z 506 in A and m/z 549, 670 in c) according to the Domon and Costello nomenclature^28^. The ^0,2^X cleavage of sialic acid (e.g., m/z 432 in b) is diagnostic of α2,6-linked sialic acid. The cross-ring cleavage (^0,2^A_GlcNAc_-H_2_O) of distal GlcNAc residues is indicative of terminal β1,4-linked Gal or GalNAc (e.g., m/z 425 in c and m/z 304 in d). Structures are depicted using the Symbol Nomenclature for Glycans (SNFG).

Our results from impalas (*Aepyceros melampus*, Fig. 2b; not studied previously) initially resembled *L. pacos*: lack of fucosylated glycans and presence of sulfated MOs. The most abundant MO, by far, was 3-sialyllactose (3’-SL), followed by 6-sialyllactose (6’-SL). Notably, we identified one lactosamine-based structure, Neu5Acα2-6Galβ1-4GlcNAc, and several structures containing the alpha-Gal motif, immunogenic in humans^29^. Additional findings included extended sulfated (Gal?1-3Gal6Sβ1-4Glc; Supplementary Dataset 1, slide 22) and phosphorylated (Gal?1-3Galβ1-4GlcP; Supplementary Dataset 1, slide 21) MOs, and highly unusual structures, such as Neu5Acα2-3GlcNAcβ1-6(Galβ1-4)Glc (Supplementary Dataset 1, slide 51). We also detected Neu5Gc as a sialic acid, and a Neu5Acα2-8Neu5Ac motif on one glycan, uncommon on MOs. Several structures, such as Neu5Acα2-3Galα1-3Galβ1-4Glc (Supplementary Dataset 1, slide 47), have, to our knowledge, never been described as a MO and represents a new glycan sequence, sialyl-a3GL. Other bovines, such as *Addax nasomaculatus*^30^, *Bos taurus*^16,31^, or *Bubalus bubalis*^12^, share characteristics such as extended sialic acids, usage of Neu5Gc, and lactosamine-based MOs, supporting their conservation in this family.

Most importantly, six of our novel glycans in *A. melampus*, such as GalNAcβ1-4GlcNAcβ1-3Galβ1-4Glc and GalNAcβ1-4GlcNAcβ1-3(Galβ1-4GlcNAcβ1-6)Galβ1-4Glc (Supplementary Dataset 1, slide 44 and 92), exhibited a terminal LacdiNAc motif (GalNAcβ1-4GlcNAc), an important motif in protein-linked glycans that has not yet been described in MOs. The two examples constitute GalNAc-substituted versions of LNnT and LNnH, and we propose to refer to them as LdiNnT (lacto-*N,N*-neotetraose) and LdiNnH. GalNAc in general constitutes a relatively rare monosaccharide in MOs and its usage within a LacdiNAc motif exhibits important functions in protein- and lipid-linked glycans^32–34^, likely also indicating a functional role in MOs. We additionally note the unusual appearance of sialylated LacdiNAc structures in impala milk, with the example of Neu5Acα2-3(GalNAcβ1-4)GlcNAcβ1-3(Galβ1-4GlcNAcβ1-6)Galβ1-4Glc (Supplementary Dataset 1, slide 120).

In the pygmy hippo (*Choeropsis liberiensis*, Fig. 2c), the first hippopotamid to be investigated with regard to glycans, we report an abundance of LNnH and some of the LacdiNAc-substituted LdiNnH, described above for impala. In total, we identify 10 LacdiNAc-containing MOs, out of 60 characterized structures. *C. liberiensis* expresses both Neu5Ac and Neu5Gc in its MOs and we found several sulfated MOs. Next to an abundance of alpha-Gal motifs, other structures of interest included Fucα1-2(GalNAcα1-3)Galβ1-4Glc, exhibiting an A-type blood group antigen, GalNAcα1-3GalNAcβ1-3Galα1-4Galβ1-4Glc, identified in pre-milk, and Galα1-3Galβ1-4GlcNAcβ1-3[GalNAcβ1-4(Neu5Acα2-6)GlcNAcβ1-6]Galβ1-4Glc, an extended structure representing an LNnH backbone with an alpha-Gal motif and a sialylated LacdiNAc motif (Supplementary Dataset 1, slide 38, 63, 146). Sialylation of GlcNAc residues is still insufficiently understood. Even in humans, the responsible enzyme is not yet fully known, with indications for ST6GALNAC6^35^.

### Aquatic Artiodactyla species exhibit GlcA in their MOs

Another major finding from our impala milk was the presence of glucuronic acid (GlcA) in one glycan, GlcAβ1-4Glc (Supplementary Dataset 1, slide 2). Usually, GlcA is found in proteoglycans. Only recently, the first GlcA-containing MOs were reported, albeit not fully structurally characterized, in cow milk, another bovine species, with speculations about their role in immunomodulation^24^. We thus set out to further investigate this understudied area of MO biochemical diversity.

To analyze this phenomenon outside of Bovidae, our samples included three aquatic mammals: beluga whales and two dolphin species, an understudied group. Previous research only reported the presence of trace amounts of 3’-SL in beluga whale milk, with even the presence of lactose being questionable^36^. This contributed to the impression of a general MO paucity in aquatic animals, with a reliance on milk fats for energy and evolutionary loss of complex MOs^21^. However, we characterized 60 unique MOs in beluga whale (Fig. 2d), including highly elaborate dodecasaccharides (e.g., Supplementary Dataset 1, slide 172).

Next to the already described 3SL, we report several MOs carrying the Sd^a^ epitope (e.g., **Neu5Acα2-3(GalNAcβ1-4)Gal**β1-4Glc; Sd^a^ epitope bold; Supplementary Dataset 1, slide 52), its Neu5Gc-substituted version (Neu5Gcα2-3(GalNAcβ1-4)Galβ1-4Glc; Supplementary Dataset 1, slide 54), and many other extended sialylated structures. Neutral MOs in beluga whales were dominated by LNnT, LNH, and nLc6. Similar to pygmy hippos, relatively closely related to whales, we identified blood group A, alpha-Gal, and Lewis x epitopes. Lactosamine-based MOs, sulfated MOs, and Neu5Gc-containing MOs were also detected.

Similar to impalas and pygmy hippos, we found nine LacdiNAc-containing structures in the milk of *D. leucas*. Next to the ones already described above, this included a GalNAc-substituted version of para-LNnH (GalNAcβ1-4GlcNAcβ1-3Galβ1-4GlcNAcβ1-3Galβ1-4Glc; Supplementary Dataset 1, slide 94), para-LdiNnH in our proposed nomenclature, and sialylated LacdiNAc structures.

Bottlenose dolphin milk has been tentatively analyzed, with nine reported MO structures^17,37^. Our results, comprising 63 unique structures from two individuals, extend these findings (Fig. 2e). Next to the already reported Sd^a^-lactose tetrasaccharide^37^, we, for instance, report MOs with two Sd^a^ motifs (Neu5Acα2-3(GalNAcβ1-4)Galβ1-4GlcNAcβ1-6(Neu5Acα2-3(GalNAcβ1-4)Galβ1-3)Galβ1-4Glc; Supplementary Dataset 1, slide 163) or extended a3GL (Fucα1-2Galα1-3Galβ1-4Glc; Supplementary Dataset 1, slide 31). The dominant neutral structure was LNnT but we again identified several LacdiNAc-containing MOs, such as LdiNnT. Other noteworthy motifs include a high degree of fucosylation, Neu5Gc-Sd^a^, blood group A, Lewis x, sulfation, and alpha-Gal.

Two exceptional findings in the milk of bottlenose dolphins were sulfated 2’-FL and GlcA-containing MOs. We present Fucα1-2Gal6Sβ1-4Glc (6-sulfo-2’-FL; Supplementary Dataset 1, slide 20) as the first fully structurally characterized sulfo-fucosyllactose, a class of MOs that was recently suggested to be present in human milk and may modulate immune activity^38^. We also discovered an extended GlcA-containing MO in bottlenose dolphin milk, with the sequence HexNAc?1-?GlcAβ1-4(Galβ1-6)Galβ1-4Glc (Supplementary Dataset 1, slide 62), in both studied individuals, which we confirmed via digestion with a β-glucuronidase. We note that, outside of proteoglycans, GlcA in an internal position is highly unusual.

We then analyzed the milk of striped dolphins (Fig. 2f; not studied previously), which exhibited the same GlcA-containing MO and its biosynthetic predecessor, HexNAc?1-?GlcAβ1-4Galβ1-4Glc (Supplementary Dataset 1, slide 43). In general, the 45 characterized MOs were reminiscent of bottlenose dolphins, with many Sd^a^ motifs, an abundance of LNnT, and a high degree of fucosylation, including Lewis x structures, DFL, and α2,6-sialylation. Of note, we detected no LacdiNAc structures in this sample.

Overall, these findings clearly question the presumed MO paucity of aquatic mammals. They also strengthen the conservation of LacdiNAc and, at least in Delphinidae, indicate a potential conservation of GlcA usage in MOs, expanding previous conceptions of the chemical diversity of MOs.

### Conservation of LacdiNAc and GlcA in MOs of Perissodactyla

From Perissodactyla, we studied the black rhinoceros, *Diceros bicornis*, with colostrum and mature milk samples from the same individual. In total, we report 43 unique MO structures (Fig. 3a), exceeding the previously reported seven MOs for this species^17^, which only contained neutral MOs. Here, we note an abundance of 3’-SL (and an absence of 6’-SL), combined with many MOs containing the Sd^a^ motif. We also note the relative dearth of Neu5Gc-containing MOs in this species. For neutral MOs, LNnP-I and LNH were dominant and we report several lactosamine-based MOs. In contrast to previous work^17^, we did not detect any fucosylated MOs in either sample.

**Figure 3.**
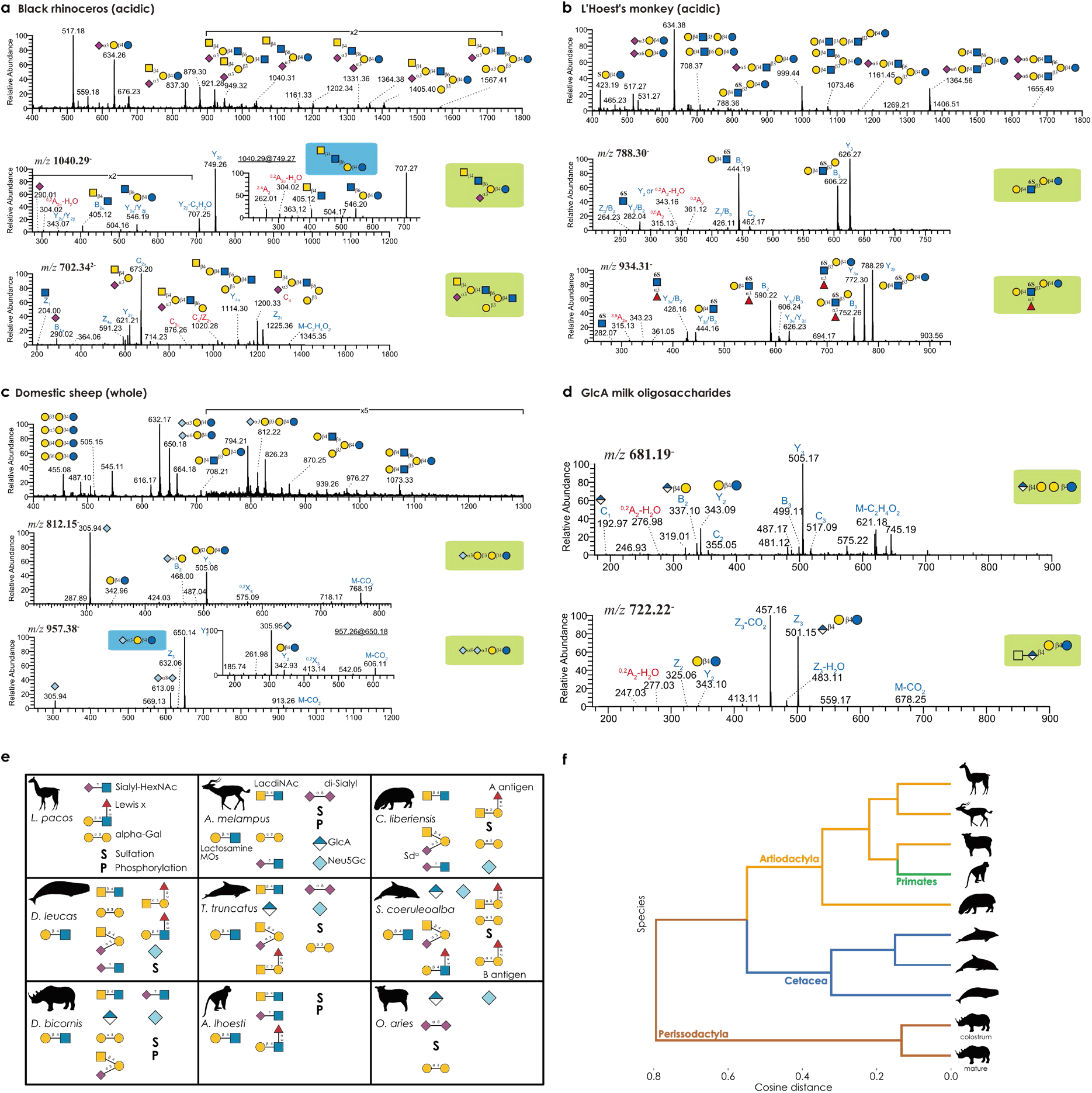
Characterizing understudied species from other taxonomic orders. **a-c**, Annotated mass spectra of free MOs for newly investigated mammalian species. Shown are the milk glycans, identified via LC-ESI-MS/MS, from *Diceros bicornis* (a), *Allochrocebus lhoesti* (b), and *Ovis aries* (c). **d**, Annotated mass spectra of additional GlcA-containing MOs. **e**, Characteristic milk oligosaccharide motifs. For each newly investigated species, a schematic view of common structural motifs, as identified here, is shown. Structures are depicted using the Symbol Nomenclature for Glycans (SNFG). **f**, Clustering of species by their milk glycome. Using the relative abundances from Supplementary Table 1, we used glycowork to calculate relative abundances of motifs and converted this into a cosine-based distance matrix of species. This was used for hierarchical clustering, with additional annotation of higher-order taxonomic groups that these species belonged to.

We also discovered the new sialyl-a3GL, first described in impalas, in this sample. Focusing on our two main discoveries in this work, LacdiNAc and GlcA-MOs, we wanted to know whether a different taxonomic order, Perissodactyla, would also exhibit these features. Indeed, we structurally characterized LacdiNAc-based MOs in the milk of black rhinoceros, such as GalNAcβ1-4GlcNAcβ1-6(Neu5Acα2-3)Galβ1-4Glc (Supplementary Dataset 1, slide 81), also present in beluga whales. We also report sialylated LacdiNAc structures, GalNAcβ1-4(Neu5Acα2-6)GlcNAcβ1-3Galβ1-4Glc (Supplementary Dataset 1, slide 82), shared with impalas and beluga whales. Additionally, we even identified four GlcA-containing MOs in the colostrum of *D. bicornis*, the abovementioned structures and an extended trisaccharide GlcAβ1-4Galβ1-3Galβ1-4Glc (Supplementary Dataset 1, slide 36). The existence of LacdiNAc- and GlcA-containing MOs in multiple glycans, individuals, species, and taxonomic orders strongly suggests some degree of conservation of these features in MOs.

### LacdiNAc is also found in MOs of Primates

We next investigated the milk from a third taxonomic order, Primates. In our analyzed primate, L’Hoest’s monkey (*Allochrocebus lhoesti*, Fig. 3b; not studied previously), we report extended sulfated MOs and several fucosylated and sialylated structures, predominantly Neu5Acα2-6, and one lactosamine-based structure, Neu5Acα2-6Galβ1-4GlcNAc. After lactose, the most-abundant MO was LSTb (sialyl-lacto-*N*-tetraose b), and we additionally detected a sulfated variant of LSTb with lower abundance. Goto and colleagues^39^ also reported sulfated MOs in another Old World monkey, *Papio hamadryas*, and the production of LSTc in *P. hamadryas* and *Macaca mulatta*. However, the finding of LSTb expression and lactosamine-based MOs in *A. lhoesti* seem to be unique among Old World monkeys, for now.

While no GlcA-containing MOs were identified in this sample, we did detect a LacdiNAc glycan in this species, GalNAcβ1-4GlcNAcβ1-3Galβ1-4Glc or LdiNnT. Another Old World monkey, *M. mulatta*, was also reported to exhibit GalNAc in its MOs^39^, yet not in the characteristic LacdiNAc motif. We speculate that the LacdiNAc motif might be found in other Old World monkeys as well. The existence of LacdiNAc-exhibiting MOs in the three orders Artiodactyla, Perissodactyla, and Primates, as shown here, suggests that this motif could be conserved in the whole of Mammalia.

Lastly, we turned to domestic sheep, *Ovis aries*, as a relatively common model system, to assess whether the breadth of structures we uncovered in our more exotic species was also recapitulated in more established species. We characterized 21 unique MO structures in mature sheep milk (Fig. 3c), recapitulating all major MOs identified in previous work^15^ and extending it by structures such as disialyl-lactose (Neu5Acα2-8Neu5Acα2-3Galβ1-4Glc). While we did not detect LacdiNAc-containing structures in our sample, we did detect GlcAβ1-4Glc, similar to the impala and rhinoceros, demonstrating another case of GlcA-containing MOs (Fig. 3d).

Taken together, we clearly show that the complexity of non-human MOs has been underestimated. Of particular importance, we present 108 entirely new glycan sequences (Fig. 4) out of 172 characterized sequences, extending the total number of unique MOs (estimated at 550-600 based on our extensive literature dataset) by >15% with our single study. New sequences, motifs, and monosaccharides, all conserved to at least some degree within Mammalia (Fig. 3e), demonstrate that this realm likely still is full of discoveries and surprises, including many family- and order-specific structures (Fig. 3f), such as the Sd^a^ motif that is highly enriched in Perissodactyla and Cetacea, or others that are likely to be antimicrobial/antiviral or otherwise functional.

**Figure 4.**
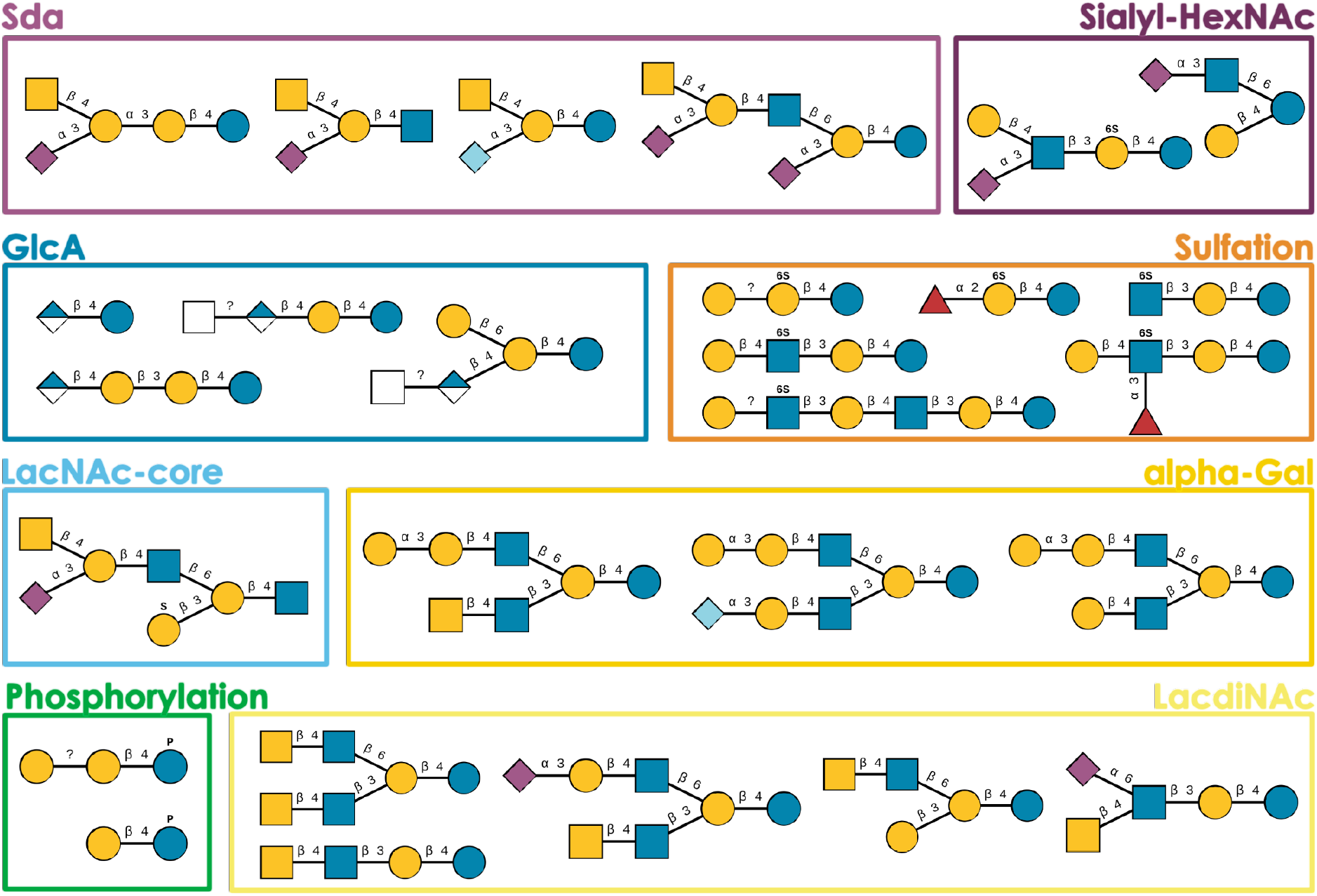
Newly discovered milk oligosaccharide structures. Some of the newly characterized oligosaccharide structures that are reported here are shown as examples of particularly salient glycan motifs in our dataset, shown in the SNFG style.

### Surveying the milk glycomes of Mammalia

To gain a sense of the evolution of MOs across mammals, we required a comprehensive dataset, allowing conclusive statements about, e.g., motif distributions, according to our current knowledge. Surveying all 1,902 (non-unique) published MOs, and combining them with our 393 observations, we find that approximately 80% of known MOs from more than 100 species stem from the taxonomic orders Primates, Artiodactyla, and Carnivora (Fig. 5a). These orders contain humans (Primates) and livestock such as cows or goats (Artiodactyla), highlighting the focus on model organisms and commercially / biomedically relevant species, which may not be representative of the number of species per taxonomic order. Even within these orders, coverage per species varies (Fig. 5b), from species with lactose as the sole described MO up to the extensively investigated humans. The number of unique MOs per species strongly correlated with the number of publications on the MOs of that species (Spearman’s rank correlation coefficient; ρ = 0.655, p < 0.001). This clearly shows that, from a repertoire size perspective, there is little reason to believe that non-human species do not exhibit order-of-magnitude similar MO diversity, making biodiversity mapping crucial to understand the biosynthetic capabilities of Mammalia.

**Figure 5.**
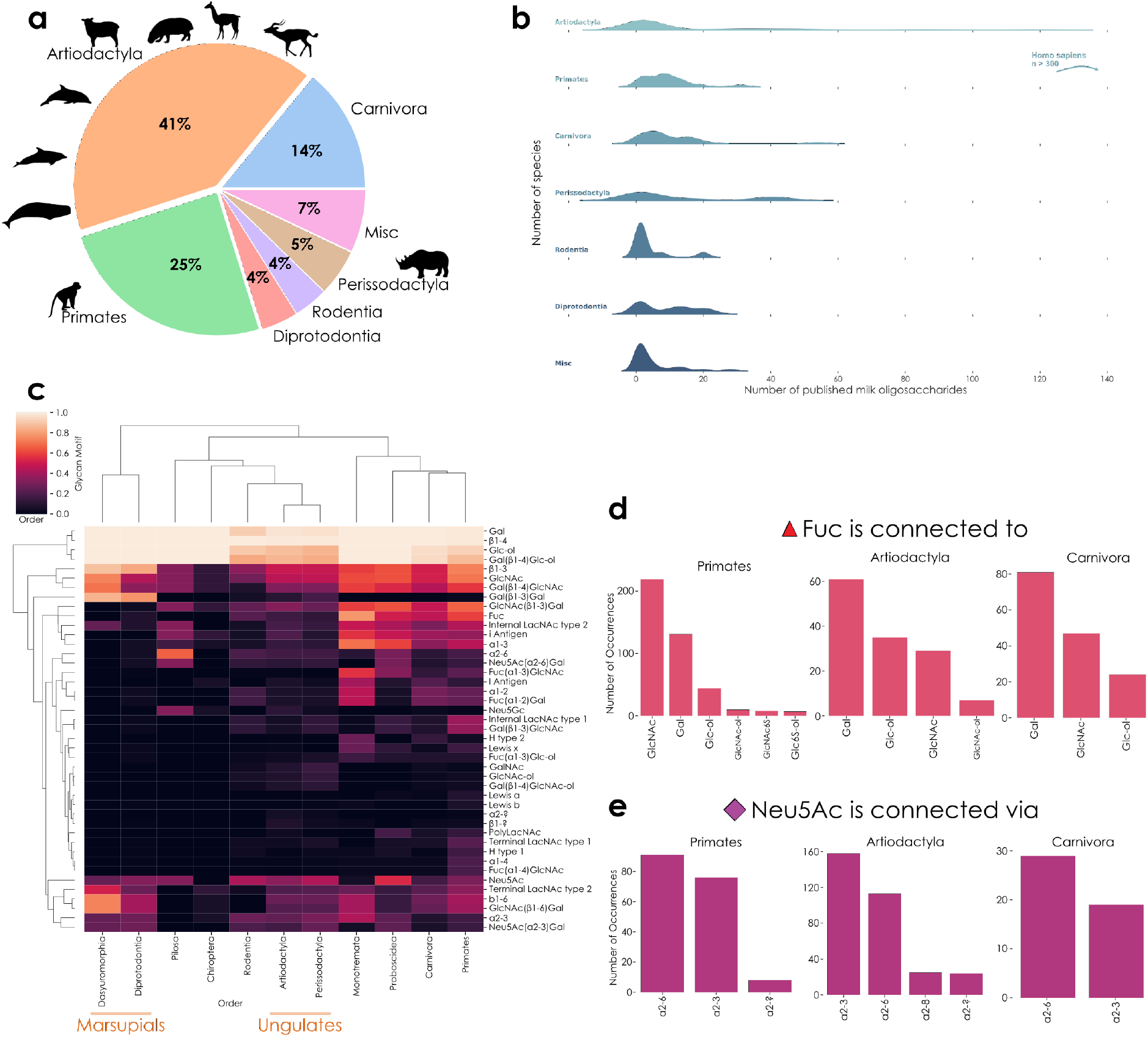
Properties of all known MOs. **a**, Milk oligosaccharides by taxonomic order. A pie chart with the proportion of milk glycans from various mammalian orders is shown, with rare orders being grouped under “Misc”. **b**, Glycan distribution by taxonomic order. For each order, the number of milk glycans per species is visualized as a distribution via a ridge plot, with the noted exception of the “outlier” *Homo sapiens*. **c**, Heatmap of milk glycan motifs. For taxonomic orders with at least five unique known MOs, we analyzed their proportion of motifs using the annotate_dataset function of glycowork version 0.6, depicted via a hierarchically-clustered heatmap. **d-e**, Linkage analysis of the monosaccharides fucose (Fuc) and *N*-acetylneuraminic acid (Neu5Ac) in milk glycans. Using glycowork, we analyzed to which monosaccharides Fuc was connected (d) and via which linkages Neu5Ac (e) was found in milk glycans of the three most exhaustively explored taxonomic orders.

MOs have different sequences, with functional consequences, which also differ by species. Therefore, we sought to characterize the distribution of MO motifs by taxonomic order (Fig. 5c). Marsupial orders (Dasyuromorphia and Diprotodontia) clearly separated from other orders, mainly due to a preponderance of neutral MO motifs in marsupials. Other orders, including Primates or Proboscidea, characteristically exhibited fucose- and sialic acid-containing motifs in their MOs, including Lewis blood group antigens.

Focusing on sequence characteristics of the three best-represented orders (Primates, Artiodactyla, and Carnivora), we analyzed their usage of fucose and sialic acids, monosaccharides crucial for anti-pathogenic effects of MOs^40,41^. While fucose was preferentially linked to GlcNAc in Primates, such as in Lewis motifs, we identified galactose as the most frequent neighbor of fucose in MOs of Artiodactyla and Carnivora (Fig. 5d). For sialic acid, Primates exhibited approximately the same amount of α2-3 and α2-6 linkages, while Artiodactyla species seemed to prefer Neu5Ac in its α2-3 configuration in their MOs (Fig. 5e). We also analyzed post-biosynthetic MO modifications, uncovering widespread sulfation of monosaccharides at C6 in Artiodactyla, Primates, and Rodentia, while sulfation at C3 was largely restricted to marsupials (Diprotodontia/Dasyuromorphia) and, to a lesser extent, Carnivora (Extended Data Fig. 1).

### GlcA milk oligosaccharides are potent immunomodulators

Building on previous speculations regarding immunomodulatory effects of GlcA^24^, we set out to test the effect of various MO formulations on macrophage activation by LPS. For this, we added chemically pure individual MOs to differentiated THP-1 macrophages – unstimulated and stimulated with LPS – and assessed the response via a multiplex cytokine assay. For 11 out of 13 measured cytokines, LPS treatment significantly upregulated cytokine production compared to baseline levels (Fig. 6a-c, Extended Data Fig. 2b), indicating robust immunostimulation of the macrophages without significant impact on the cell viability (Extended Data Fig. 2a).

**Figure 6.**
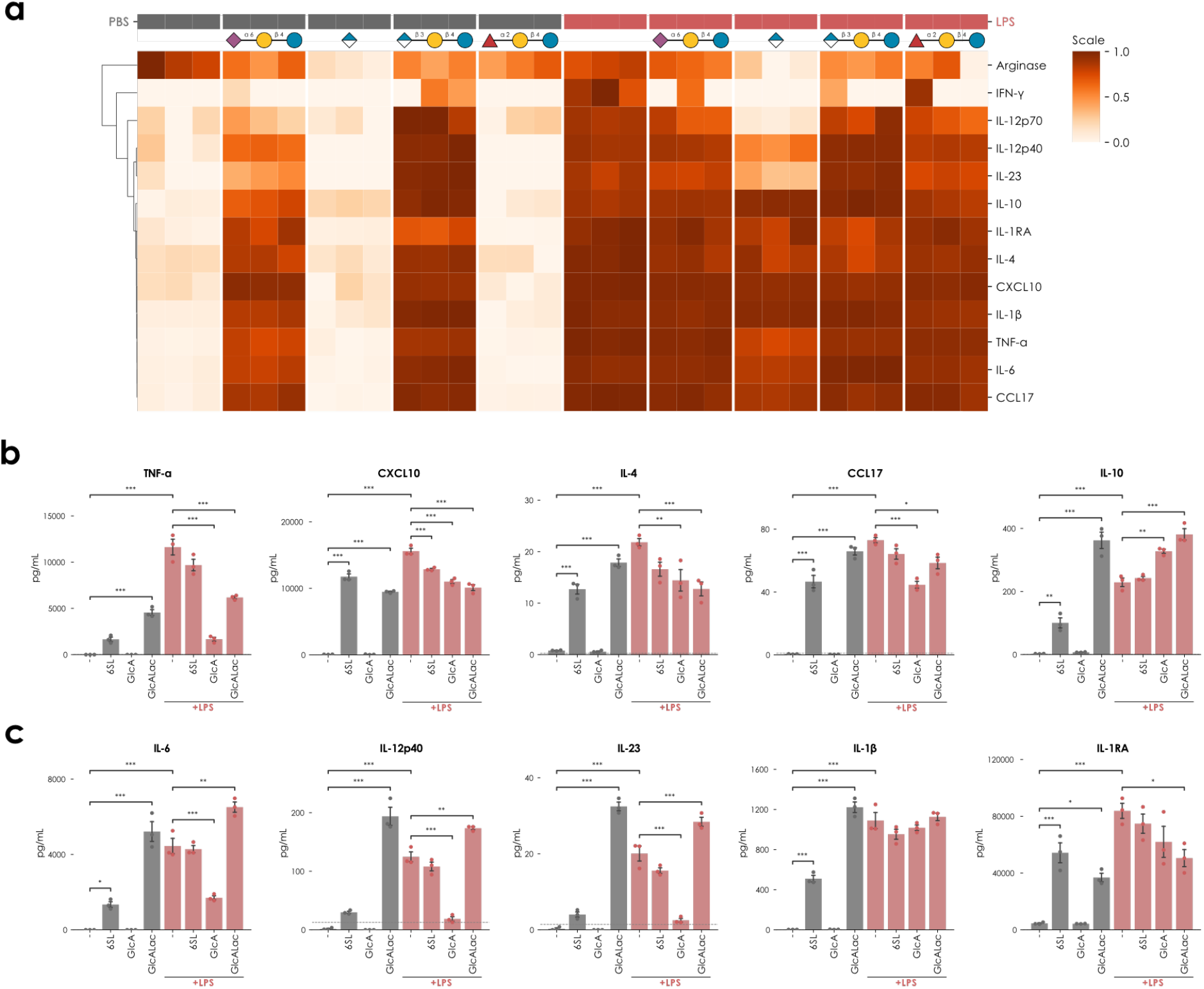
Glucuronylated milk glycans are immunomodulatory. **a**, Heatmap of cytokine concentrations of Arginase, IFN-γ, IL-12p70, IL-12p40, IL-23, IL-10, IL-1RA, IL-4, CXCL10, IL-1β, TNF-α, IL-6, and CCL17 from the culture supernatant of THP-1 cells unstimulated (grey) or stimulated with LPS (red) in the absence or presence of various MO-derived glycan structures. The row dimension was normalized and clustered by correlation distance. **b-c**, Quantification of cytokine concentration of TNF-α, CXCL10, IL-4, CCL17, IL-10, IL-6, IL-12p40, IL-23, IL-1β, and IL-1RA from the culture supernatant of THP-1 cells unstimulated (grey) or stimulated with LPS (red) in the absence or presence of various MO-derived glycan structures. The dashed line indicates the limit of detection as determined by the standard curve of each analyte. Significant differences were established via a one-way ANOVA with Tukey’s multiple comparison test. ***, p < 0.001; **, p < 0.01; *, p < 0.05.

In the absence of LPS stimulation, treatment with 2’-FL had no impact on the production of macrophage-derived cytokines. In contrast, treatment with acidic MOs – 6’-SL and GlcALac – robustly induced cytokine production, with the GlcALac-induced levels exceeding those achieved by LPS treatment alone for IL-10, IL-6, IL-12p40, IL-23, and IL-1β. Interestingly, this induction was not observed for the treatment with glucuronic acid as a monosaccharide, indicating the importance of the lactose core structure for its effect (Fig. 6b-c).

Supporting previous reports on immunomodulatory functions of MOs^42^, we observed significant inhibition of LPS-induced production of pro-inflammatory cytokines. Compared to 6’-SL, GlcALac had a more potent effect in dampening the LPS-induced production of Th1-directing molecules, TNF-α and CXCL10, and Th2-directing molecules, IL-4 and CCL17, in addition to a more potent induction of anti-inflammatory IL-10 (Fig. 6b). Moreover, and in contrast to 6’-SL, GlcALac treatment further enhanced the LPS-induced production of Th17-inducing cytokines, IL-6, IL-12p40, and IL-23, while also promoting IL-1β signaling through downregulating its inhibitor, IL-1RA (Fig. 6b). Taken together, these results demonstrate a potent immunomodulatory effect of GlcA-MOs and could indicate a physiological function, such as regulating the infant response to gut-colonizing bacteria through promoting the polarization of IL-10-dependent non-pathogenic Th17 cells^43,44^.

## Discussion

To obtain a comprehensive overview of the biodiversity and evolution of breast milk oligosaccharides, a representative set of Mammalia needs to be studied. However, even though we present and analyze data from a set of over 100 mammalian species, large swaths of Mammalia still constitute frontiers in MO research. Taxonomic orders such as Chiroptera, Cingulata, Didelphimorphia, Eulipotyphla, Lagomorpha, Peramelemorphia, Pilosa, Rodentia, Sirenia, or Tubulidentata are poorly characterized or not at all, regarding their MOs. Including diverse environments, from the sea to the air, we expect many more unexpected biodiversity discoveries in their MOs, akin to the MO LacdiNAc motif uncovered here. Even already characterized species may deserve a revisit, as evidenced by our discovery of GlcA-MOs in sheep milk. Additionally, MO descriptions of most exotic mammals rely on a single study and a single population, sometimes even a single individual and/or a single time-point during lactation. We are therefore convinced that the diversity and complexity of the various non-human milk glycomes is currently substantially underestimated.

One property of LacdiNAc-decorated glycans in other glycan classes is to facilitate the adhesion of bacteria such as *Helicobacter pylori*^32^. Aligning with functions of other MOs, preventing the adhesion of toxins, bacteria, and viruses by mimicking adhesion epitopes^45^, we propose that the herein discovered LacdiNAc motif in MOs serves a similar purpose for yet to be identified neonate pathogens in LacdiNAc-expressing species. Exhibited by cetaceans, primates, and both ungulates, we envision that the LacdiNAc motif is far from rare in MOs, despite its obscurity to this point. Its biosynthesis remains unclear yet, though it is likely that the isoenzymes B4GALNT3/4, responsible for LacdiNAc synthesis on protein-linked glycans^46^, are involved.

Together with recent findings by Gray et al., describing GlcA-containing MOs in cow milk^24^, we contribute towards the emerging conclusion that GlcA seems to be used more broadly in MOs, at least in Artiodactyla and Perissodactyla. This provides a third modality for negative charge in MOs, next to sialic acids and sulfation. In cows, GlcA also seems to be post-biosynthetically sulfated^24^, potentially implying that the human natural killer-1 (HNK-1, CD57) epitope, important in immune regulation^47^, could be found in MOs. Combined with our macrophage immunomodulation results and the paucity of known pathogens that could be inhibited by GlcA, we speculate that physiological functions of GlcA-containing MOs might relate to immune modulation, an important area for MOs^6^, such as facilitating the establishment of the infant microbiome.

While for human MOs specific functions have been extensively investigated^42,48,49^, functional roles of MO motifs or structures in most animals still remain elusive. So far, MO functions seem to fall into the broad categories of nutrition, anti-pathogen, and immunomodulation^4,8,50,51^. Future work could investigate potential anti-pathogen functionality of the described structures, in cases where species-specific pathogens are known. Computational methods, predicting binding to pathogen glycan-binding proteins^52^, and screening methods, such as glycan arrays^53^, could shed light on functions of motifs such as the herein discovered MO LacdiNAc, considering its abovementioned connection to *Helicobacter pylori*^32^.

Our work demonstrates that MOs, especially non-human MOs, are substantially more diverse than currently appreciated. In multiple species, and multiple oligosaccharides within those species, we show that the MO repertoire contains more monosaccharides (GlcA) and more motifs (e.g., LacdiNAc) than canonically assumed. We anticipate that further exploration of MOs will uncover further biochemical diversity. Our curated and measured dataset represents a currently exhaustive collection of what is known about MO biodiversity, and we anticipate that it will be used to produce new insights in glycobiology, MO biosynthesis, and breast milk function.

## Online Methods

### Literature dataset curation

To generate a comprehensive resource of all currently known milk oligosaccharides, we manually inspected all 19,902 published articles on PubMed (term=milk+carbohydrate) between 1898 and December 2021. Milk oligosaccharides were extracted from papers with sequence information on species-specific milk oligosaccharides. This was supplemented by various Google searches for publications that are not indexed on PubMed (e.g., “milk oligosaccharides”, “milk glycans”, “mammalian breast milk”). We also compared and completed our dataset with milk oligosaccharides from GlyCosmos^54^, GlyConnect^55^, GlyGen^56^, and GlycoStore^57^. All milk oligosaccharides were formatted into IUPAC-condensed nomenclature and paired with their full taxonomic information. This resulted in 1,902 species-specific MO records from 168 species (with 62 species having only been reported to contain lactose in their breast milk). All curated milk oligosaccharides can be found in Supplementary Table 2 and stored within our Python package, glycowork^58^.

### Breast milk samples from uncharacterized mammals

Milk samples from alpaca (*Lama pacos*) and L’Hoest’s monkey (*Allochrocebus lhoesti*) were donated by Benoît Quintard from the Zoo Mulhouse (France). Samples from impala (*Aepyceros melampus*) were provided by Thierry Petit from the Zoo La Palmyre (France). Breast milk from pygmy hippo (*Choreopsis liberiensis*) and black rhinoceros (*Diceros bicornis*) was donated by Florine Wedlarski from Zoo de Doué la Fontaine (France). Milk from beluga whale (*Delphinapterus leucas*), bottlenose dolphin (*Tursiops truncatus*), and striped dolphin (*Stenella coeruleoalba*) were provided by Jose Luis Crespo from Fundación Oceanogràfic de la Comunitat Valenciana (Spain). Sheep milk (*Ovis aries*) was gifted by Rykets Gård (Sweden). All milk samples constituted mature breast milk, sampled according to local regulations, with an additional colostrum sample from black rhinoceros and a pre-milk sample from pygmy hippo.

### Milk oligosaccharide extraction

All breast milk samples followed the same protocol. Per sample, 500 μL of breast milk were diluted with the same amount of distilled water and centrifuged at 4,000 g for 30 min at 4°C. The skimmed milk was separated from the milk fat layer and the cell pellet and transferred to a fresh tube. Then, two volumes of cold 96% ethanol were added to one volume of milk and incubated at 4°C overnight to precipitate protein. Subsequently, the sample was again centrifuged at 4,000 g for 30 min at 4°C. The milk oligosaccharide-containing supernatant was then transferred into a fresh tube, frozen, and lyophilized.

Freeze-dried milk oligosaccharides were suspended in water (250 µL/mL original milk). Trace proteins were removed by spin-filter (10 kDa cutoff, 11,000 rpm for 10 min, Sigma-Aldrich, St. Louis, MO, USA). Reduction was carried out with 0.5 M NaBH_4_ and 20 mM NaOH at 50°C overnight. The samples were desalted using cation exchange resin (AG^®^50WX8, Bio-Rad, Hercules, CA, USA) packed onto a ZipTip C18 tip (Sigma-Aldrich). After drying via SpeedVac, additional methanol was added to remove residual borate by evaporation. The resulting glycans were then analyzed via liquid chromatograph-electrospray ionization tandem mass spectrometry (LC-MS/MS) with or without further fractionation. Relative abundances of experimentally determined MOs can be found in Supplementary Table 1.

The released MOs were further fractionated into neutral and acidic fraction using DEAE Sephadex A-25 (GH Healthcare, Uppsala, Sweden). As lactose is the dominant MO in the neutral fraction, which will suppress the signal of other minor neutral MOs during LC-MS/MS, it was removed using carbon solid phase cartridge (HyperSep™ Hypercarb™ SPE cartridges 25 mL, Thermo Scientific, Sweden). In brief, the cartridge was conditioned with 3 × 500 µL of 90% MeCN with 0.1% TFA, 3 × 500 µL of 0.1% TFA. After applying the MOs, lactose was eluted with 3 × 500 µL 8% MeCN with 0.1% TFA. The neutral MOs were further eluted with 3 × 500 µL 65% MeCN with 0.1% TFA, dried by centrifugation evaporation, and stored at - 20°C until analysis. To characterize the type of Lewis structure, MOs without reduction were also analyzed.

For 70 mL of *C. liberiensis* pre-milk, we performed affinity chromatography to enrich for GalNAc-terminated MOs, via 1 mL of VVL (*Vicia villosa* lectin) linked to agarose beads. Elution was performed by heating the bound material at 70°C for one hour (denaturing VVL). Then, we collected the first four fractions for analysis (each fraction equals one column volume, or 1.2 mL).

### Analysis of milk oligosaccharides via LC-MS/MS

Milk oligosaccharides were analyzed by liquid chromatograph-electrospray ionization tandem mass spectrometry. The oligosaccharides, dissolved in water, were separated on a column (10 cm × 250 µm) packed in-house with 5 µm porous graphite particles (Hypercarb, Thermo-Hypersil, Runcorn, UK). The oligosaccharides were injected onto the column and eluted with an acetonitrile gradient (Buffer A, 10 mM ammonium bicarbonate; Buffer B, 10 mM ammonium bicarbonate in 80% acetonitrile). The gradient (0-45% Buffer B) was eluted for 46 min, followed by a wash step with 100% Buffer B, and equilibrated with Buffer A in the next 24 min.

The samples were analyzed in negative ion mode on a LTQ linear ion trap mass spectrometer (Thermo Electron, San José, CA), with an IonMax standard ESI source equipped with a stainless-steel needle kept at –3.5 kV. Compressed air was used as nebulizer gas. The heated capillary was kept at 270°C, and the capillary voltage was –50 kV. A full scan (m/z 340 or 380-2000, two microscan, maximum 100 ms, target value of 30,000) was performed, followed by data-dependent MS^2^ scans (two microscans, maximum 100 ms, target value of 10,000) with normalized collision energy of 35%, isolation window of 2.5 units, activation q=0.25, and activation time 30 ms). The threshold for MS^2^ was set to 300 counts. Data acquisition and processing were conducted with Xcalibur software (Version 2.0.7). For comparing MO abundances between samples, individual glycan structures were quantified relative to the total content by integrating the extracted ion chromatogram peak area. The area under the curve (AUC) of each structure was normalized to the total AUC and expressed as a percentage. The peak area was processed by Progenesis QI (Nonlinear Dynamics Ltd., Newcastle upon Tyne, UK).

### Linkage determination of milk oligosaccharides

MOs were identified from their MS/MS spectra by manual annotation together with exoglycosidase verification. Diagnostic fragmentation ions for assignment were investigated as described previously^59–61^. For instance, the Lewis types were determined by diagnostic ions resulting from double glycosidic cleavages (C/Z ions) encompassing 4-linked branches obtained from MS/MS of non-reducing MOs. In case of Lewis a/b, the presence was supported by the fragmentation ions at *m/z* 348, while Lewis x/y was indicated by the fragmentation ions at *m/z* 364 and 510, respectively^61^.

MOs were treated, prior to re-analysis by LC-MS/MS, with different exoglycosidases, alone or in combination, including α2-3 neuraminidase S (NEB), α2-3,6,8,9 neuraminidase A (NEB), α1-3,6-galactosidase (NEB), β1-4 galactosidase S (NEB), α-N-acetylgalactosaminidase (NEB), and *E. coli* β-glucuronidase (Megazyme, desalted and concentrated 10 times with a 10 kDa cutoff spinfilter before use). The recombinant *Bacteroides thetaiotaomicron* 3SGal sulfatase (BT1626) was described previously^62^. As appropriate, the treated glycans were desalted as described above. Changes in re-analysis were used for structural elucidation according to the known enzyme specificity. All proposed structures were informed by MS/MS information as well as enzymatic degradation, with example spectra located in Supplementary Dataset 1.

### Immunomodulation analysis of milk oligosaccharides

Human THP-1 cells (ATCC, TIB-202) were maintained in Roswell Park Memorial Institute 1640 medium (RPMI 1640; Gibco, A1049101) supplemented with 10% (v/v) fetal bovine serum (FBS; Nordic Biolabs, FBS-HI-12A) and 1% (v/v) penicillin-streptomycin solution (Sigma-Aldrich, P4333-100ML) in a humidified atmosphere containing 5% CO_2_ at 37 °C. 4 × 10^4^ cells/well were seeded in a 96-well plate and differentiated into macrophage-like cells by treatment with 25 nM phorbol-12-myristate-13-acetate (PMA; Sigma-Aldrich, P8139-1MG) for 48 hours. After exchanging the medium and an additional 24-hour rest period, the cells were challenged with 100 ng/mL lipopolysaccharide (LPS; Sigma-Aldrich, L4391-1MG) in the absence or presence of 1 mg/mL chemically pure MOs (2-fucosyllactose: 2’-FL, Sigma-Aldrich, SMB00933-10MG; 6-sialyllactose: 6’-SL, Sigma-Aldrich, 40817-1MG; Glucuronic acid: GlcA, Sigma-Aldrich, G5269-10MG, Glucuronyl-lactose: GlcALac, Elicityl Oligotech, GLY180-50%-5MG). As a control, we also included MO treatment of the cells in the absence of LPS. After 24 hours, the culture supernatants were collected and analyzed for the levels of key macrophage cytokines using a multiplex immunoassay based on fluorescence-encoded beads (LEGENDplex; Biolegend, 740503) according to the manufacturer’s instructions. The samples were acquired on an Accuri C6 Plus Flow Cytometer (BD) and the results were analyzed using LEGENDplex Data Analysis Software Suite (Biolegend, Qognit).

### Motif analysis

All motif analyses were performed using the Python package glycowork (version 0.6.0)^58^. The motif distribution was visualized via the motif.analysis.make_heatmap function while the linkage patterns were analyzed with the motif.analysis.characterize_monosaccharide function. In all analyses, only species with at least five recorded MOs were considered. Clustering of our species was performed by annotating motifs in their MOs via the motif.annotate.annotate_dataset function, followed by averaging relative abundances across motifs, calculating pairwise cosine distances of species, and a hierarchical clustering.

### Statistical analysis

All statistical tests in this work were done via a one-way ANOVA with Tukey’s multiple comparison test. In all cases, significance was defined as p < 0.05, after correcting for multiple testing.

## Data and Code Availability

All code used here is available in the Python package glycowork version 0.6. Data curated or generated here can be found in the supplementary tables as well as stored as internal datasets within glycowork. The glycomics MS raw files have been deposited in the GlycoPOST database under the ID GPST000317 (https://glycopost.glycosmos.org/preview/197848368063b54edf7c2f6; code:7373).

## Acknowledgments

The authors would like to thank the Zoo Mulhouse, Zoo La Palmyre, Zoo de Doué la Fontaine, Fundación Oceanogràfic de la Comunitat Valenciana, and Rykets Gård for generous sample donations. This work was funded by a Branco Weiss Fellowship – Society in Science awarded to D.B., by the Knut and Alice Wallenberg Foundation, and the University of Gothenburg, Sweden. Mass spectrometry measurements in this work were supported by the BioMS National Infrastructure in Sweden.

## Author Contributions

C.J., E.K., and J.L. prepared and measured milk samples. J.L. conducted the macrophage experiments. A.S.L. prepared the sulfatase. D.B. prepared the literature dataset. C.J., D.B., and J.L. generated the figures. All authors wrote and edited the manuscript.

