## Supplemental Figures for "Breast Milk Oligosaccharides Contain Immunomodulatory Glucuronic Acid and LacdiNAc"

**Supplementary Figures**

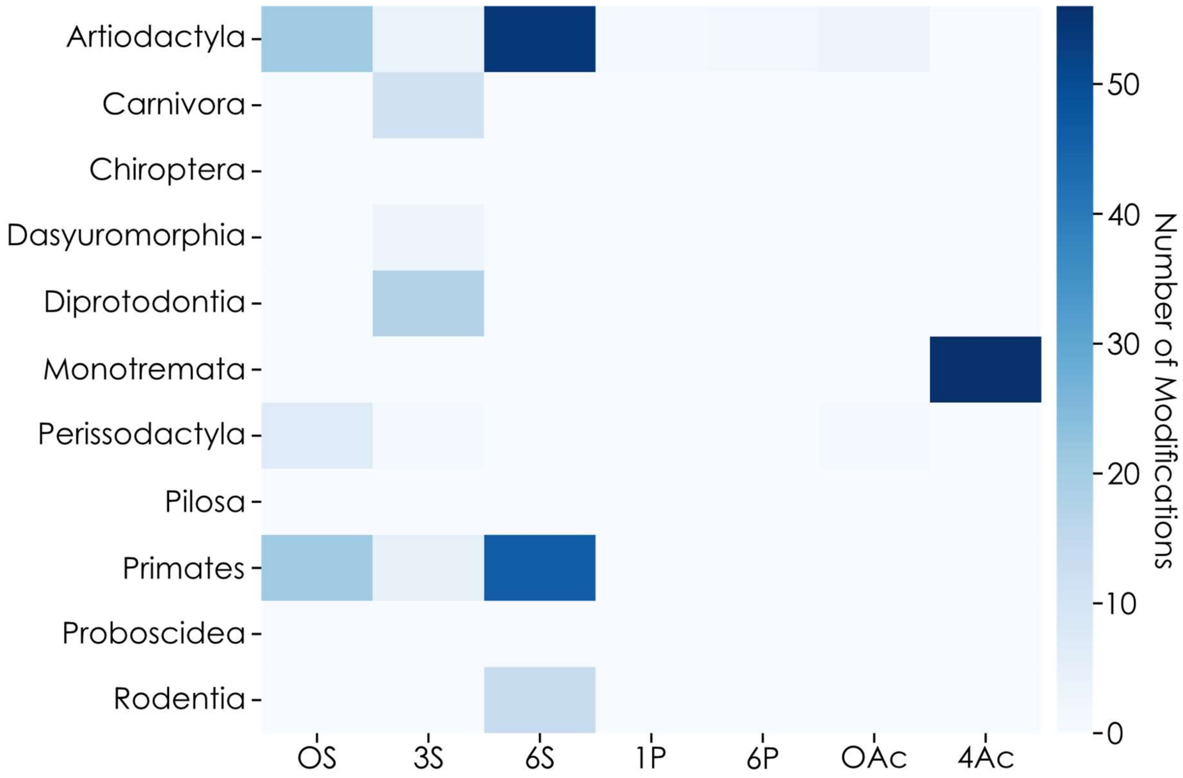


**Extended Data Figure 1. Distribution of post-biosynthetic modifications across milk oligosaccharides.** For all species with more than five milk oligosaccharides, we counted the occurrence of sulfation (OS, 3S, 6S), phosphorylation (1P, 6P), and acetylation (OAc, 4Ac) and summed them by taxonomic order, depicted as a heatmap.

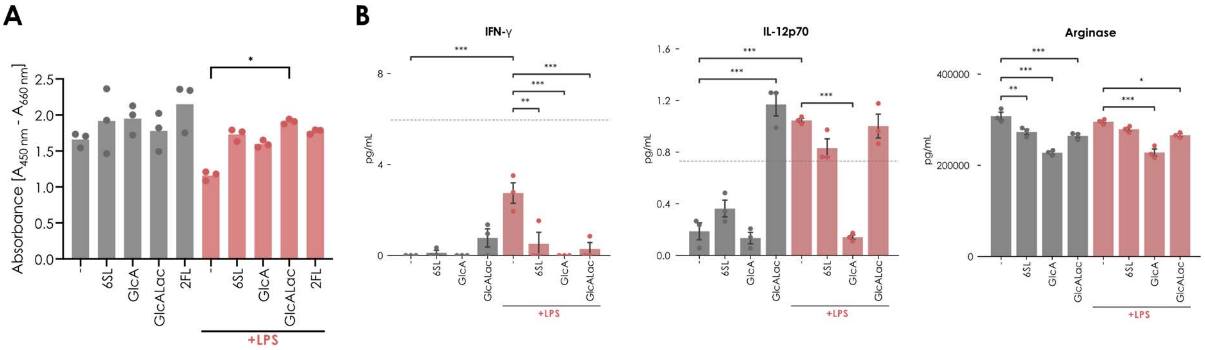


**Extended Data Figure 2. Effect of glucuronylated milk glycans on viability and other cytokines.** **a,** XTT viability assay of THP-1 cells unstimulated (grey) or stimulated with LPS (red) in the absence or presence of various MO-derived glycan structures. **b,** Measurement of cytokine concentration of IFN-γ, IL-12p70, and Arginase from the culture supernatant of THP-1 cells unstimulated (grey) or stimulated with LPS (red) in the absence or presence of various MO-derived glycan structures. The dashed line indicates the limit of detection as determined by the standard curve of each analyte. Significant differences were established via a one-way ANOVA with Tukey's multiple comparison test. ***, $p < 0.001$; **, $p < 0.01$; *, $p < 0.05$.
