## Supplemental Dataset 1 for "Breast Milk Oligosaccharides Contain Immunomodulatory Glucuronic Acid and LacdiNAc"

### Slide 1
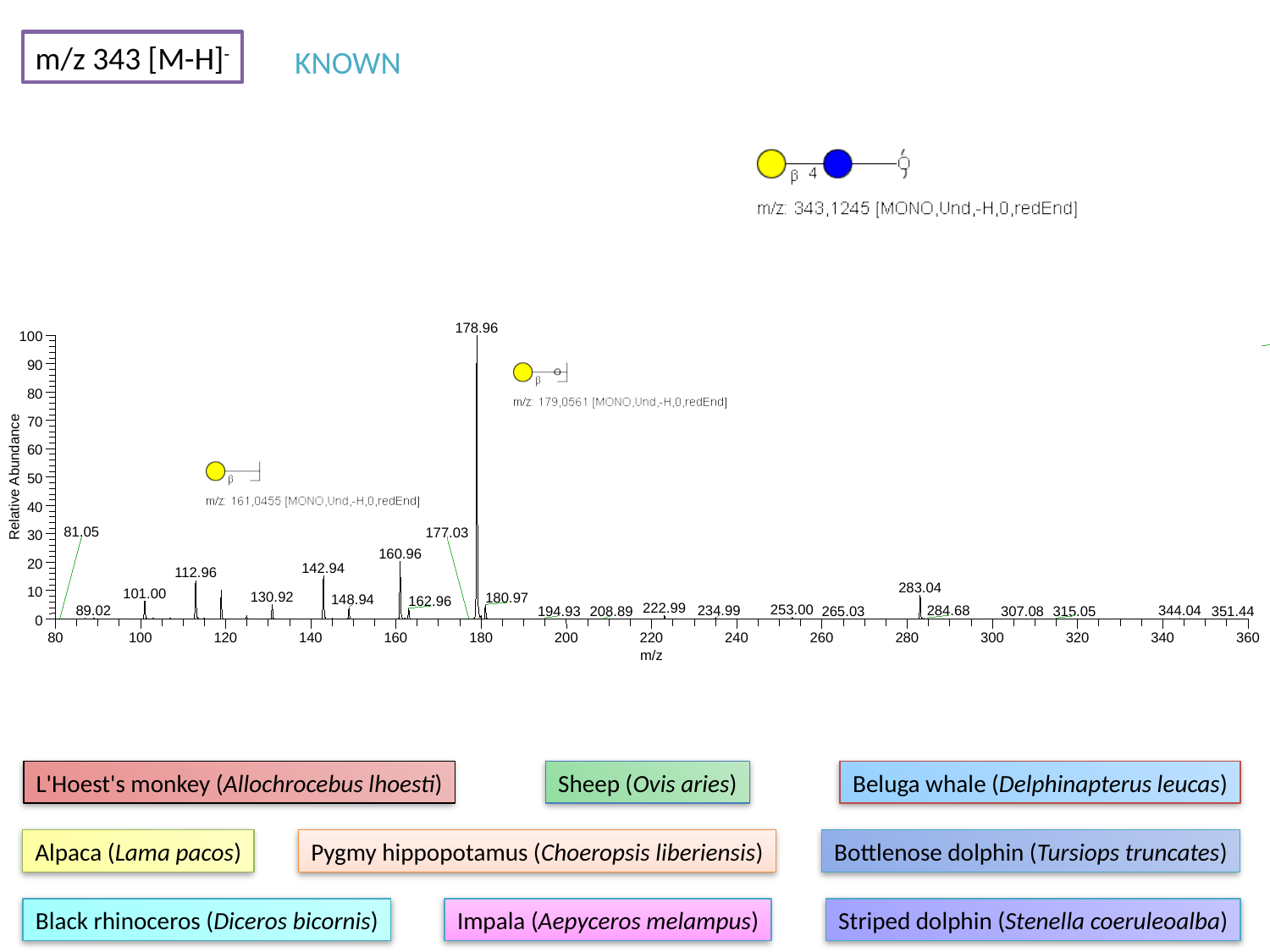

m/z 343 [M-H]^-^
KNOWN
178.96
100
90
80
70
60
Relative Abundance
50
40
81.05
177.03
30
160.96
20
142.94
112.96
283.04
10
101.00
130.92
180.97
148.94
162.96
222.99
253.00
89.02
234.99
284.68
344.04
265.03
194.93
208.89
307.08
315.05
351.44
0
80
100
120
140
160
180
200
220
240
260
280
300
320
340
360
m/z
L'Hoest's monkey (Allochrocebus lhoesti)
Sheep (Ovis aries)
Beluga whale (Delphinapterus leucas)
Alpaca (Lama pacos)
Pygmy hippopotamus (Choeropsis liberiensis)
Bottlenose dolphin (Tursiops truncates)
Black rhinoceros (Diceros bicornis)
Impala (Aepyceros melampus)
Striped dolphin (Stenella coeruleoalba)

### Slide 2
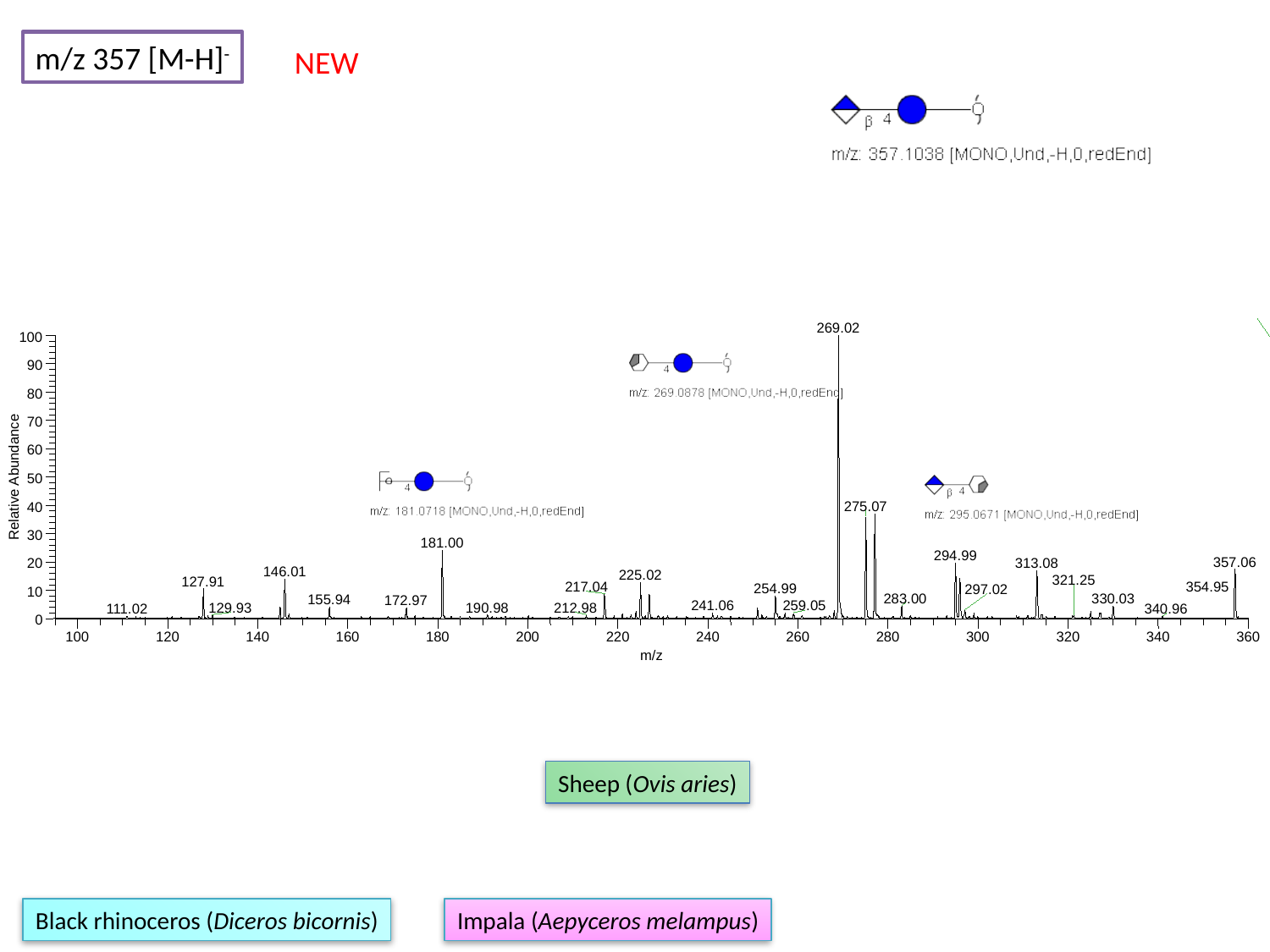

m/z 357 [M-H]^-^
NEW
269.02
100
90
80
70
60
Relative Abundance
50
275.07
40
30
181.00
294.99
357.06
313.08
20
146.01
225.02
321.25
127.91
217.04
354.95
254.99
297.02
10
283.00
330.03
155.94
172.97
241.06
259.05
190.98
212.98
129.93
111.02
340.96
0
100
120
140
160
180
200
220
240
260
280
300
320
340
360
m/z
Sheep (Ovis aries)
Black rhinoceros (Diceros bicornis)
Impala (Aepyceros melampus)

### Slide 3
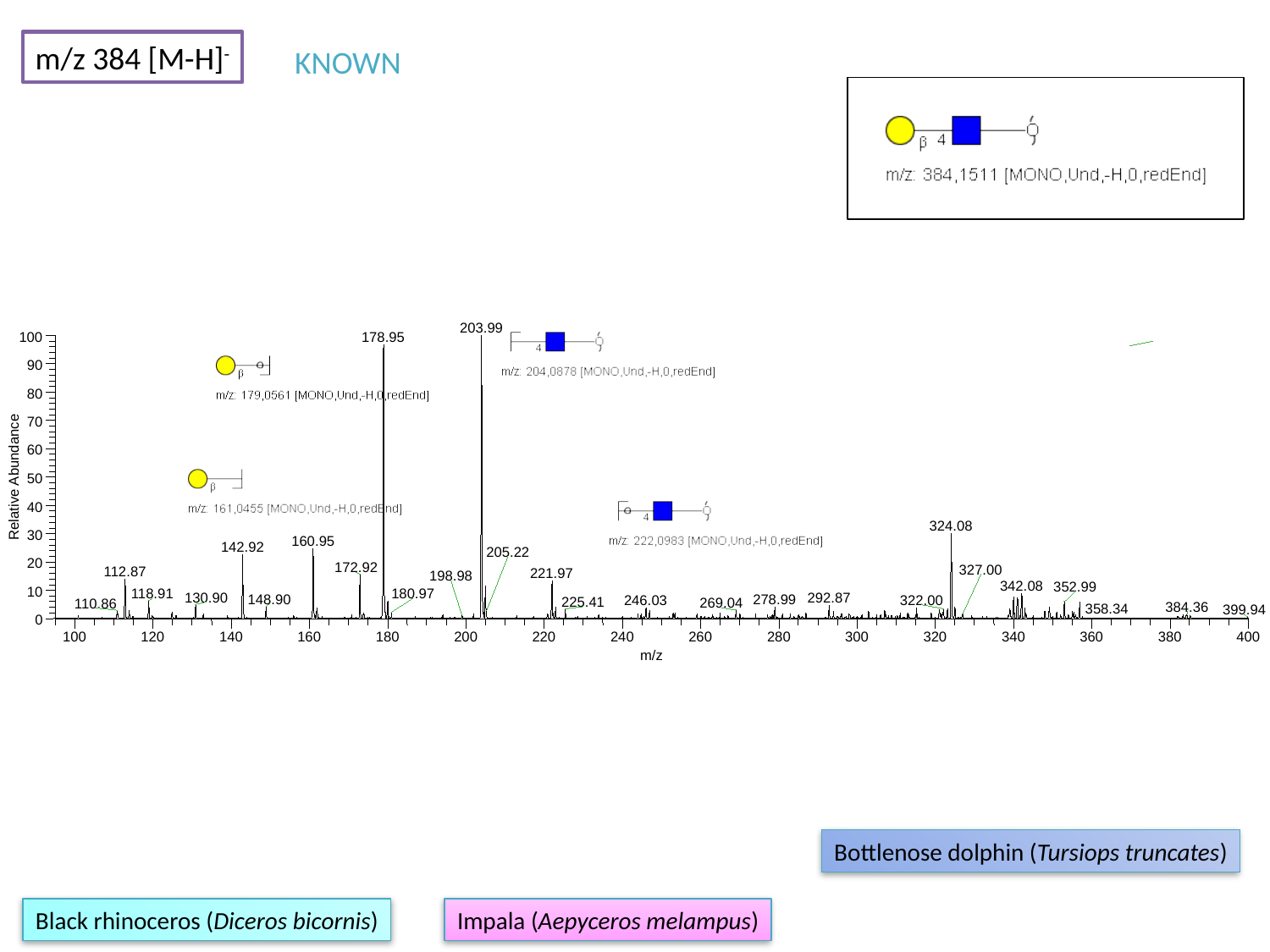

m/z 384 [M-H]^-^
KNOWN
203.99
100
178.95
90
80
70
60
Relative Abundance
50
40
324.08
30
160.95
142.92
205.22
20
172.92
327.00
112.87
221.97
198.98
342.08
352.99
10
180.97
118.91
130.90
292.87
148.90
278.99
246.03
322.00
225.41
269.04
110.86
384.36
0
100
120
140
160
180
200
220
240
260
280
300
320
340
360
380
400
m/z
358.34
399.94
Bottlenose dolphin (Tursiops truncates)
Black rhinoceros (Diceros bicornis)
Impala (Aepyceros melampus)

### Slide 4
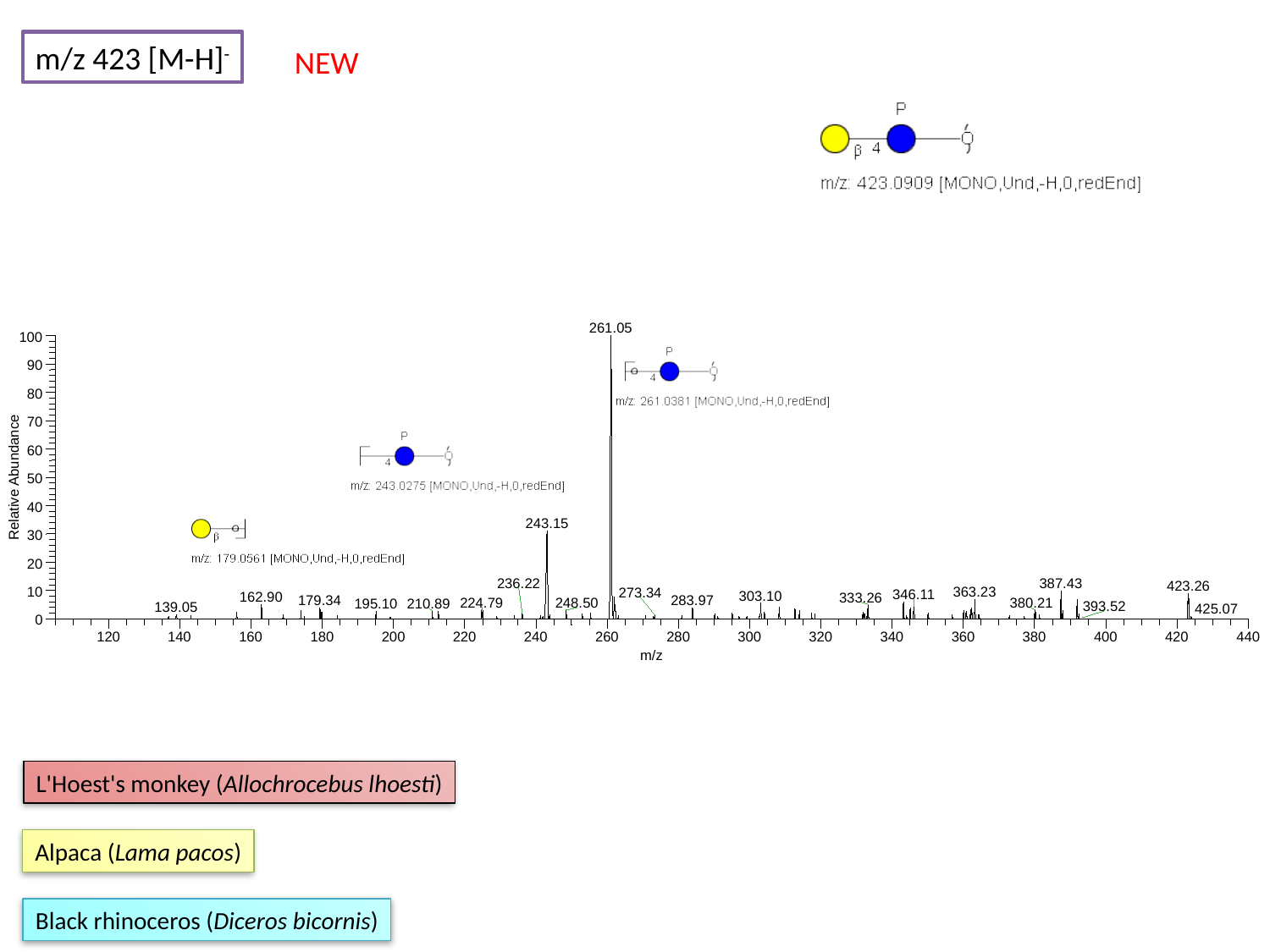

m/z 423 [M-H]^-^
NEW
261.05
100
90
80
70
60
Relative Abundance
50
40
243.15
30
20
236.22
387.43
423.26
363.23
10
273.34
346.11
303.10
162.90
333.26
179.34
283.97
224.79
248.50
380.21
210.89
195.10
393.52
139.05
425.07
0
120
140
160
180
200
220
240
260
280
300
320
340
360
380
400
420
440
m/z
L'Hoest's monkey (Allochrocebus lhoesti)
Alpaca (Lama pacos)
Black rhinoceros (Diceros bicornis)

### Slide 5
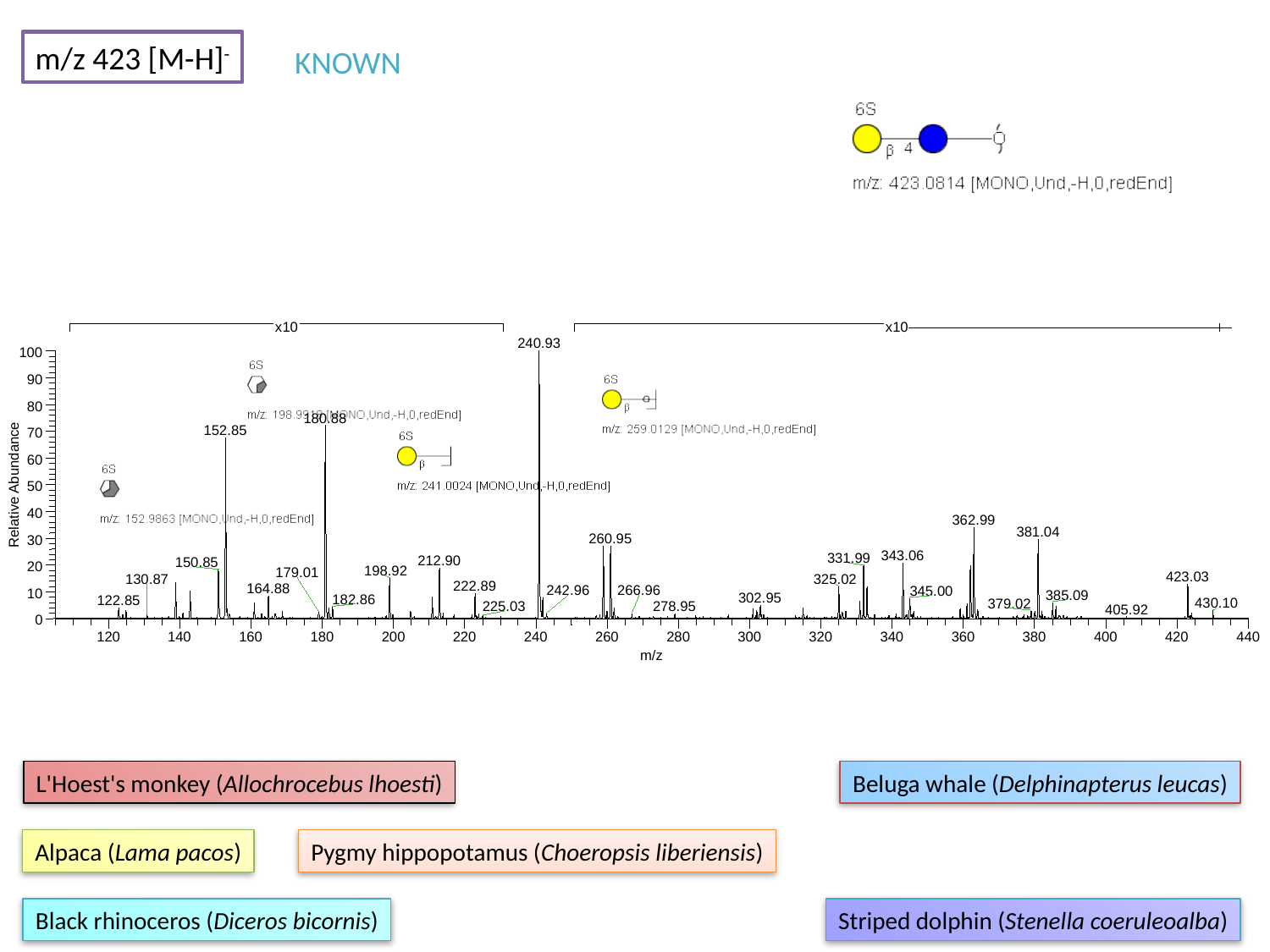

m/z 423 [M-H]^-^
KNOWN
240.93
100
90
80
180.88
152.85
70
60
Relative Abundance
50
40
362.99
381.04
260.95
30
343.06
331.99
212.90
150.85
20
198.92
179.01
423.03
325.02
222.89
164.88
242.96
345.00
10
385.09
302.95
182.86
122.85
379.02
0
120
140
160
180
200
220
240
260
280
300
320
340
360
380
400
420
440
m/z
x10
x10
130.87
266.96
430.10
225.03
278.95
405.92
L'Hoest's monkey (Allochrocebus lhoesti)
Beluga whale (Delphinapterus leucas)
Alpaca (Lama pacos)
Pygmy hippopotamus (Choeropsis liberiensis)
Black rhinoceros (Diceros bicornis)
Striped dolphin (Stenella coeruleoalba)

### Slide 6
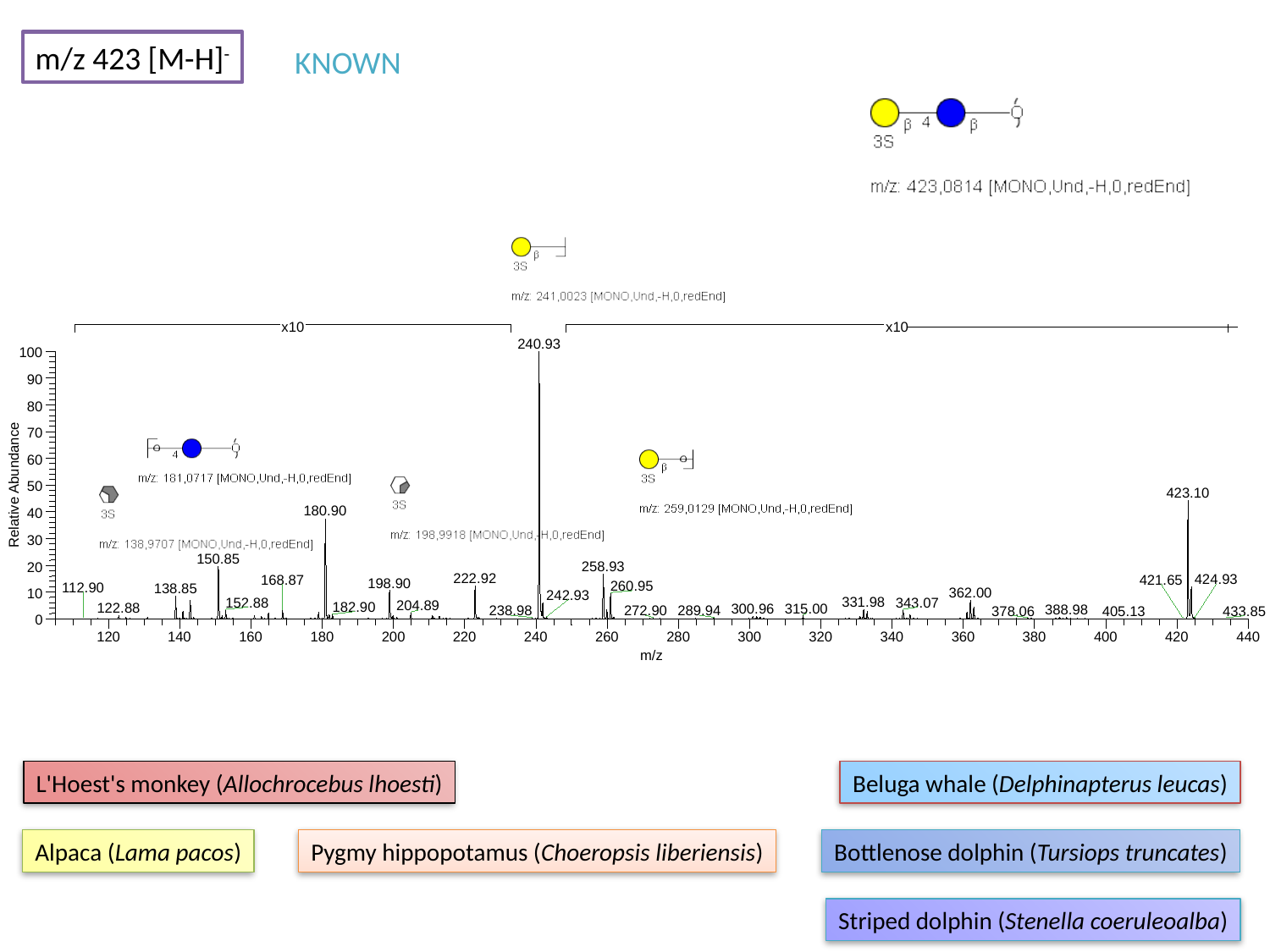

m/z 423 [M-H]^-^
KNOWN
240.93
100
90
80
70
60
Relative Abundance
50
423.10
180.90
40
30
150.85
258.93
20
222.92
168.87
198.90
260.95
138.85
362.00
10
242.93
331.98
152.88
343.07
204.89
182.90
122.88
315.00
300.96
388.98
238.98
289.94
0
120
140
160
180
200
220
240
260
280
300
320
340
360
380
400
420
440
m/z
x10
x10
424.93
421.65
112.90
272.90
378.06
405.13
433.85
L'Hoest's monkey (Allochrocebus lhoesti)
Beluga whale (Delphinapterus leucas)
Alpaca (Lama pacos)
Pygmy hippopotamus (Choeropsis liberiensis)
Bottlenose dolphin (Tursiops truncates)
Striped dolphin (Stenella coeruleoalba)

### Slide 7
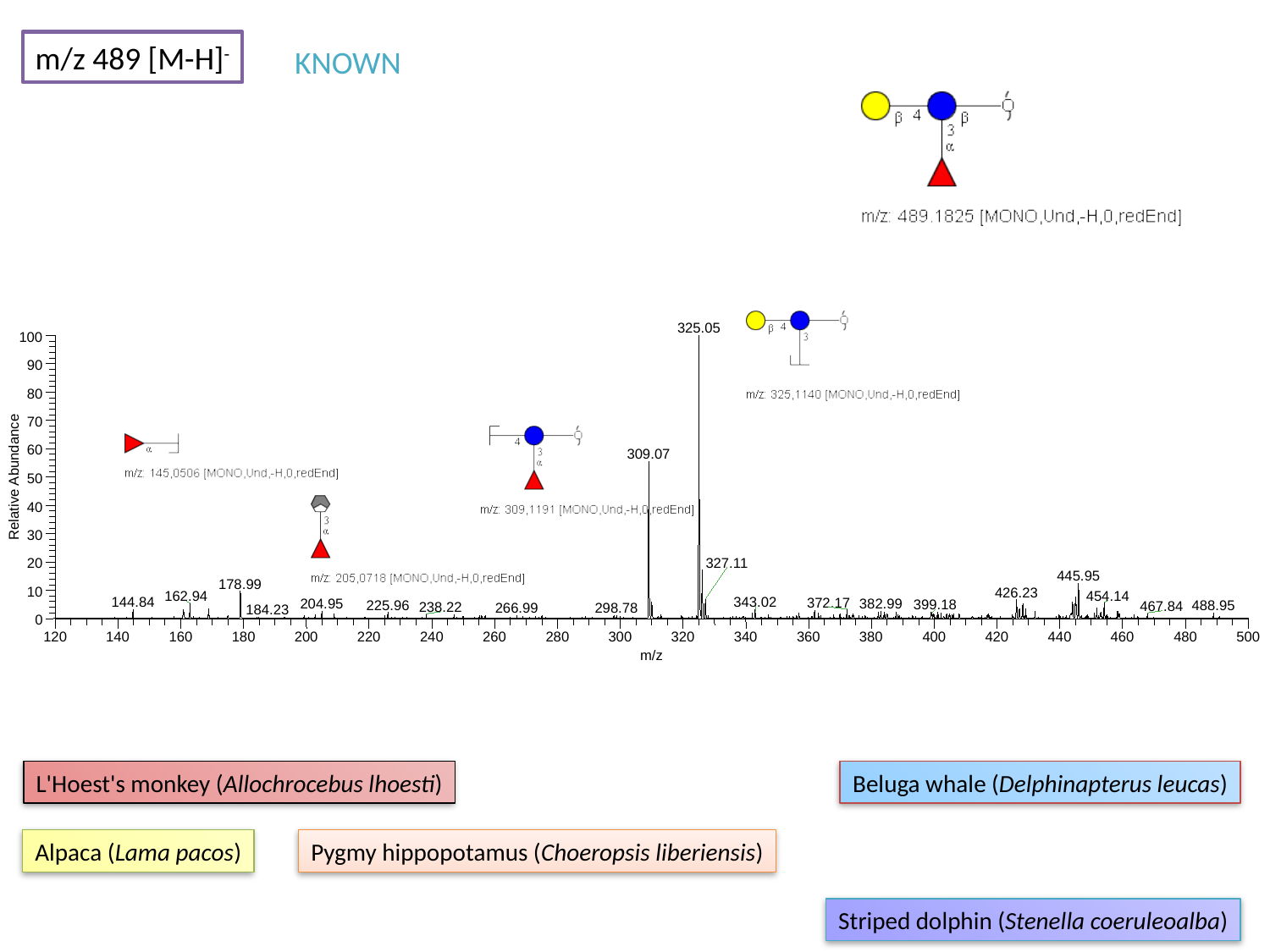

m/z 489 [M-H]^-^
KNOWN
325.05
100
90
80
70
60
309.07
Relative Abundance
50
40
30
327.11
20
445.95
178.99
10
426.23
454.14
162.94
343.02
144.84
372.17
382.99
204.95
399.18
0
120
140
160
180
200
220
240
260
280
300
320
340
360
380
400
420
440
460
480
500
m/z
225.96
488.95
467.84
238.22
266.99
298.78
184.23
L'Hoest's monkey (Allochrocebus lhoesti)
Beluga whale (Delphinapterus leucas)
Alpaca (Lama pacos)
Pygmy hippopotamus (Choeropsis liberiensis)
Striped dolphin (Stenella coeruleoalba)

### Slide 8
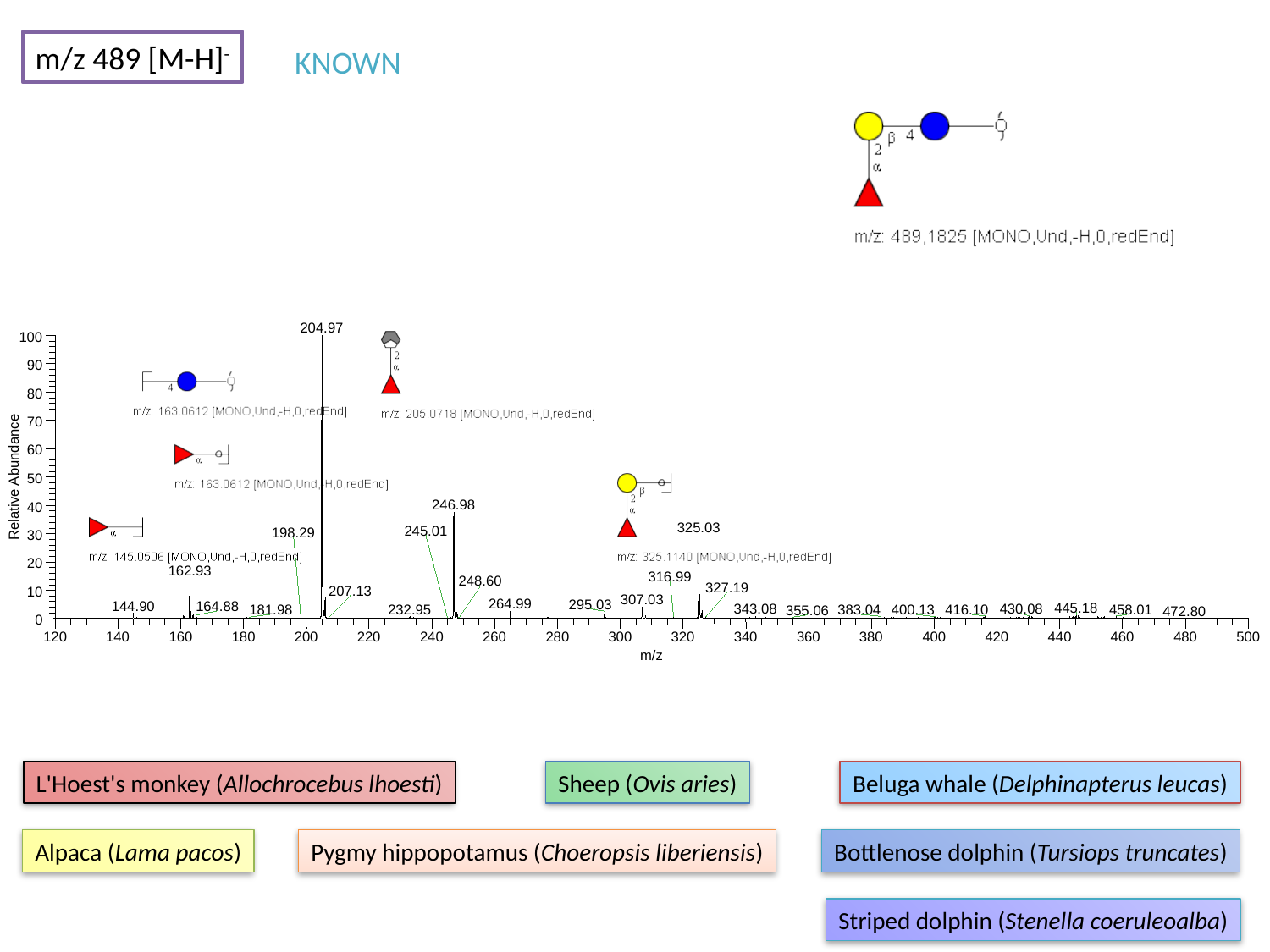

m/z 489 [M-H]^-^
KNOWN
204.97
100
90
80
70
60
Relative Abundance
50
246.98
40
325.03
30
20
162.93
10
307.03
264.99
295.03
144.90
164.88
445.18
343.08
430.08
232.95
416.10
400.13
0
120
140
160
180
200
220
240
260
280
300
320
340
360
380
400
420
440
460
480
500
m/z
245.01
198.29
316.99
248.60
327.19
207.13
458.01
181.98
383.04
355.06
472.80
L'Hoest's monkey (Allochrocebus lhoesti)
Sheep (Ovis aries)
Beluga whale (Delphinapterus leucas)
Alpaca (Lama pacos)
Pygmy hippopotamus (Choeropsis liberiensis)
Bottlenose dolphin (Tursiops truncates)
Striped dolphin (Stenella coeruleoalba)

### Slide 9
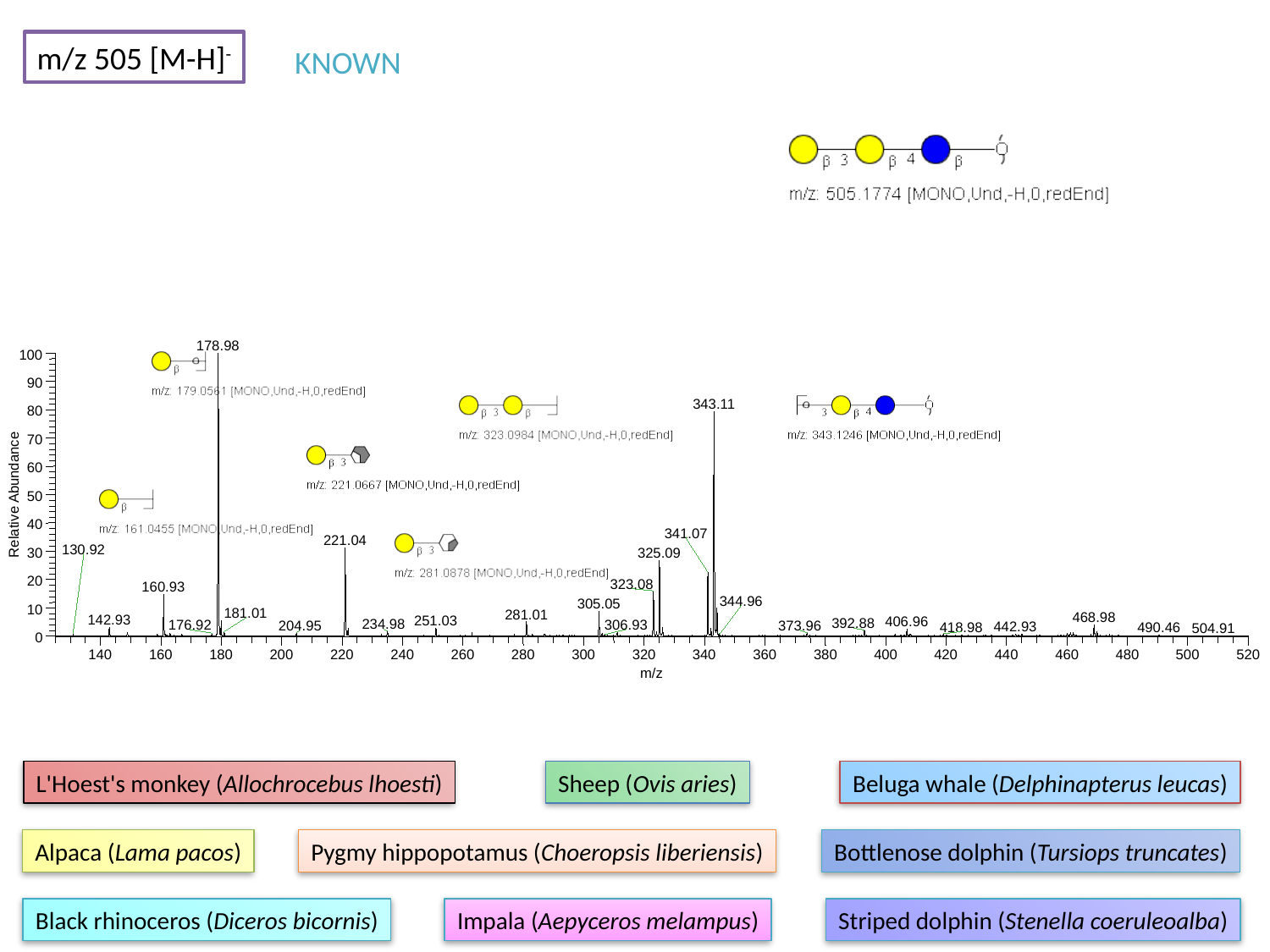

m/z 505 [M-H]^-^
KNOWN
178.98
100
90
343.11
80
70
60
Relative Abundance
50
40
341.07
221.04
30
325.09
20
323.08
160.93
305.05
10
281.01
468.98
142.93
251.03
406.96
392.88
0
140
160
180
200
220
240
260
280
300
320
340
360
380
400
420
440
460
480
500
520
m/z
130.92
344.96
181.01
234.98
306.93
176.92
204.95
373.96
442.93
418.98
490.46
504.91
L'Hoest's monkey (Allochrocebus lhoesti)
Sheep (Ovis aries)
Beluga whale (Delphinapterus leucas)
Alpaca (Lama pacos)
Pygmy hippopotamus (Choeropsis liberiensis)
Bottlenose dolphin (Tursiops truncates)
Black rhinoceros (Diceros bicornis)
Impala (Aepyceros melampus)
Striped dolphin (Stenella coeruleoalba)

### Slide 10
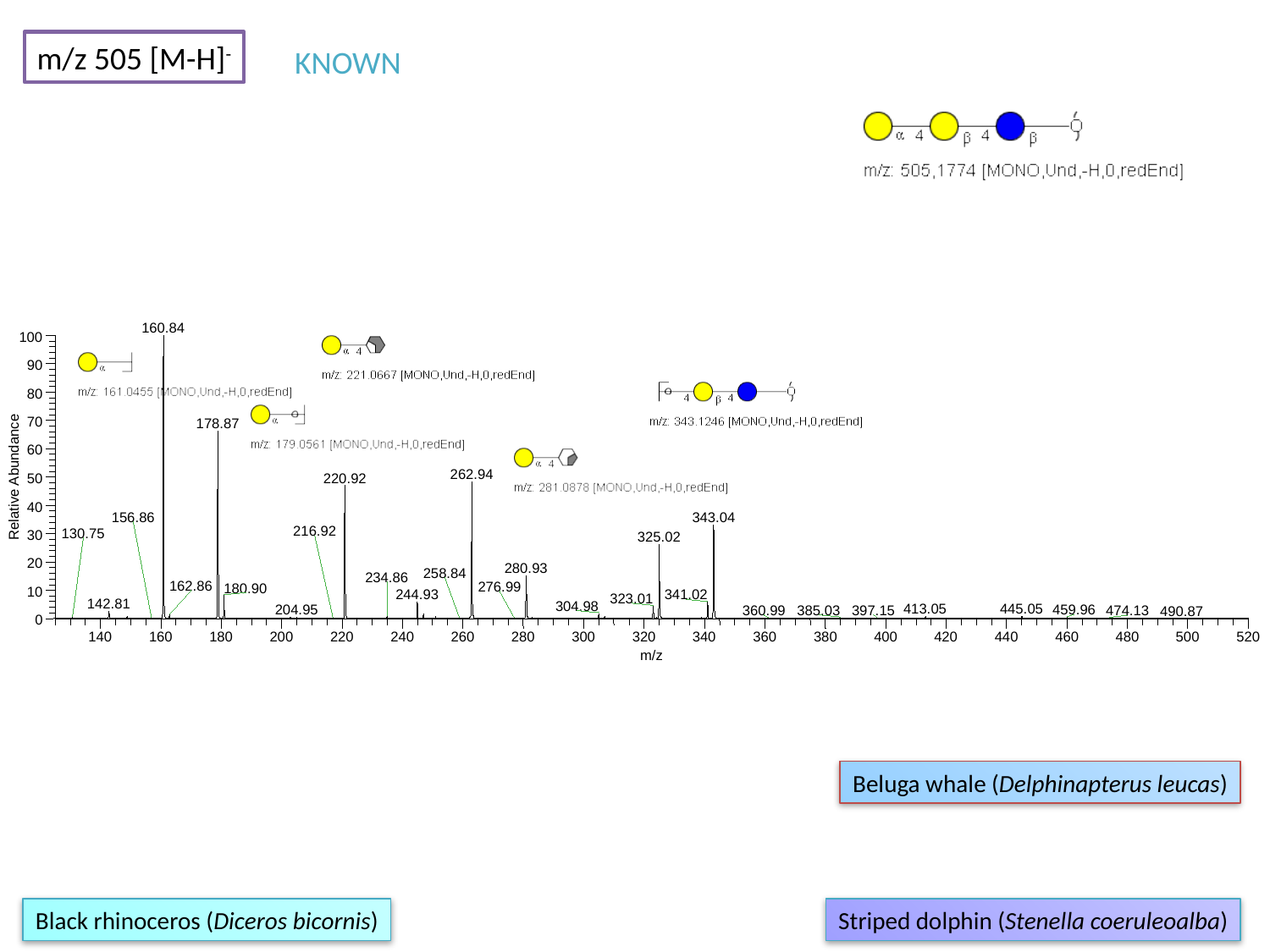

m/z 505 [M-H]^-^
KNOWN
160.84
100
90
80
70
178.87
60
262.94
Relative Abundance
220.92
50
40
343.04
30
325.02
20
280.93
180.90
10
244.93
341.02
323.01
142.81
304.98
0
140
160
180
200
220
240
260
280
300
320
340
360
380
400
420
440
460
480
500
520
m/z
156.86
216.92
130.75
258.84
234.86
162.86
276.99
413.05
445.05
204.95
459.96
385.03
397.15
360.99
474.13
490.87
Beluga whale (Delphinapterus leucas)
Black rhinoceros (Diceros bicornis)
Striped dolphin (Stenella coeruleoalba)

### Slide 11
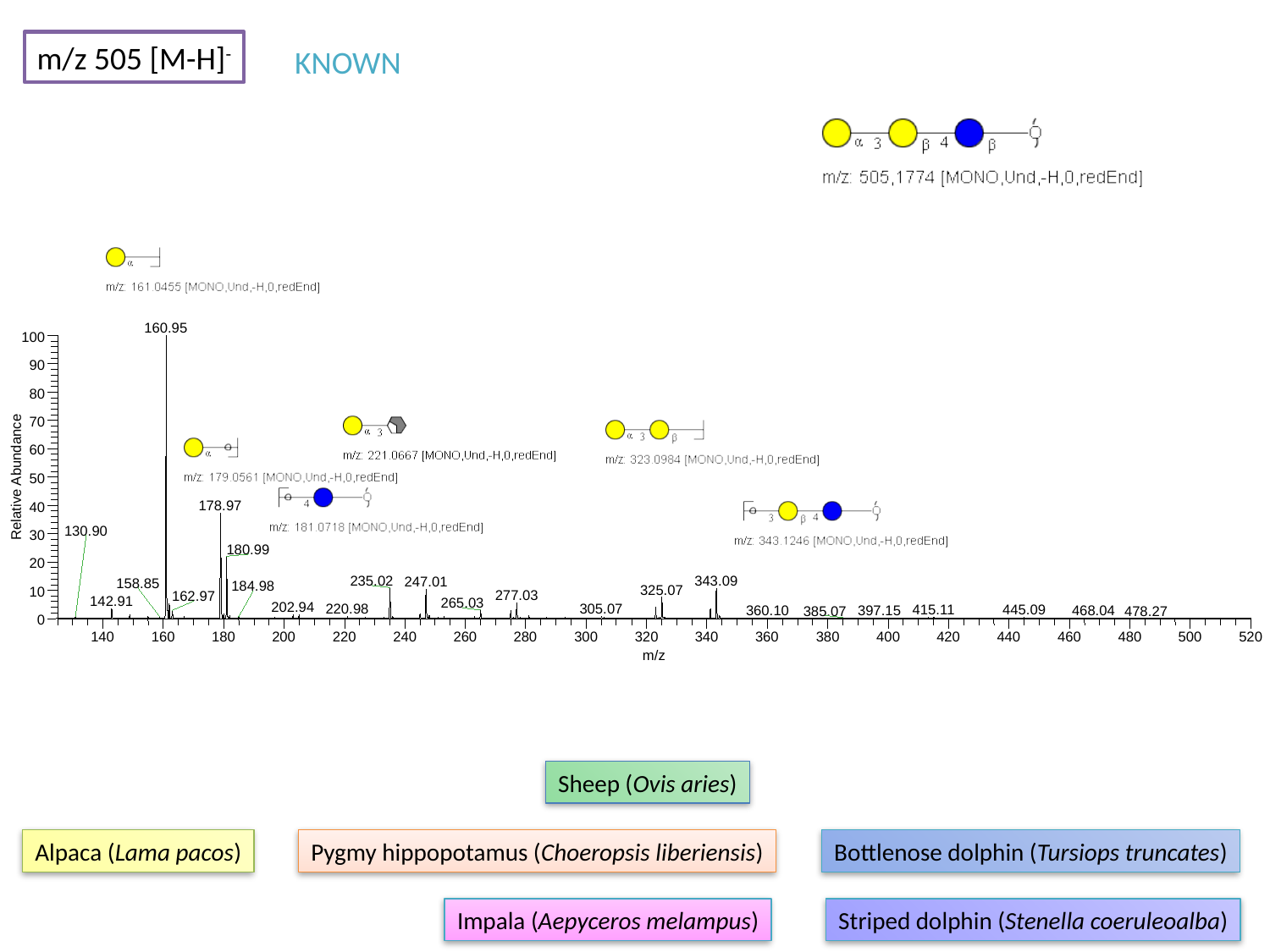

m/z 505 [M-H]^-^
KNOWN
160.95
100
90
80
70
60
Relative Abundance
50
178.97
40
30
180.99
20
235.02
343.09
247.01
325.07
10
277.03
162.97
142.91
265.03
202.94
305.07
0
140
160
180
200
220
240
260
280
300
320
340
360
380
400
420
440
460
480
500
520
m/z
130.90
158.85
184.98
220.98
415.11
445.09
360.10
397.15
468.04
385.07
478.27
Sheep (Ovis aries)
Alpaca (Lama pacos)
Pygmy hippopotamus (Choeropsis liberiensis)
Bottlenose dolphin (Tursiops truncates)
Impala (Aepyceros melampus)
Striped dolphin (Stenella coeruleoalba)

### Slide 12
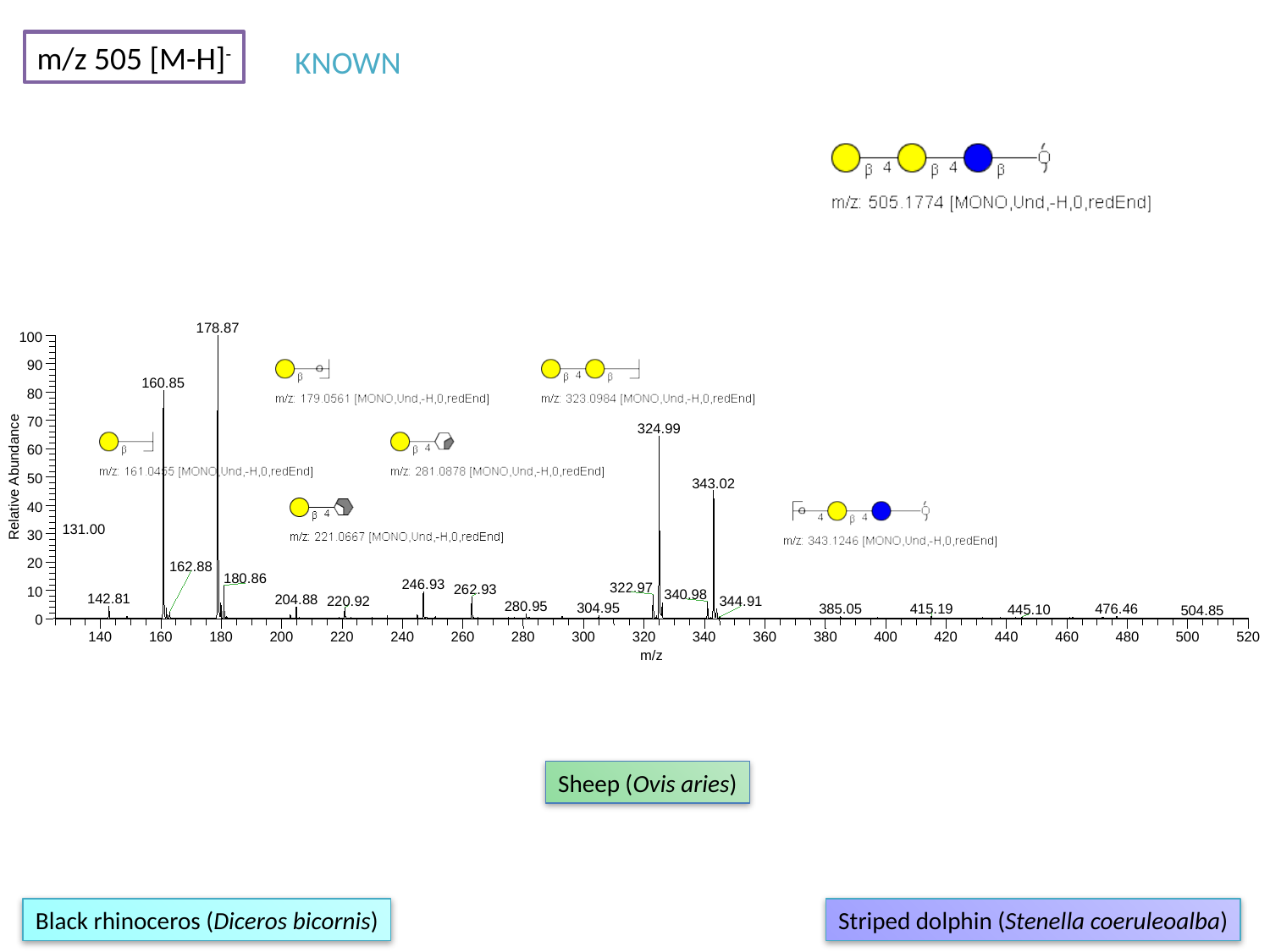

m/z 505 [M-H]^-^
KNOWN
178.87
100
90
160.85
80
70
324.99
60
Relative Abundance
50
343.02
40
30
20
180.86
246.93
322.97
262.93
10
340.98
142.81
204.88
220.92
0
140
160
180
200
220
240
260
280
300
320
340
360
380
400
420
440
460
480
500
520
m/z
131.00
162.88
344.91
280.95
304.95
385.05
415.19
476.46
445.10
504.85
Sheep (Ovis aries)
Black rhinoceros (Diceros bicornis)
Striped dolphin (Stenella coeruleoalba)

### Slide 13
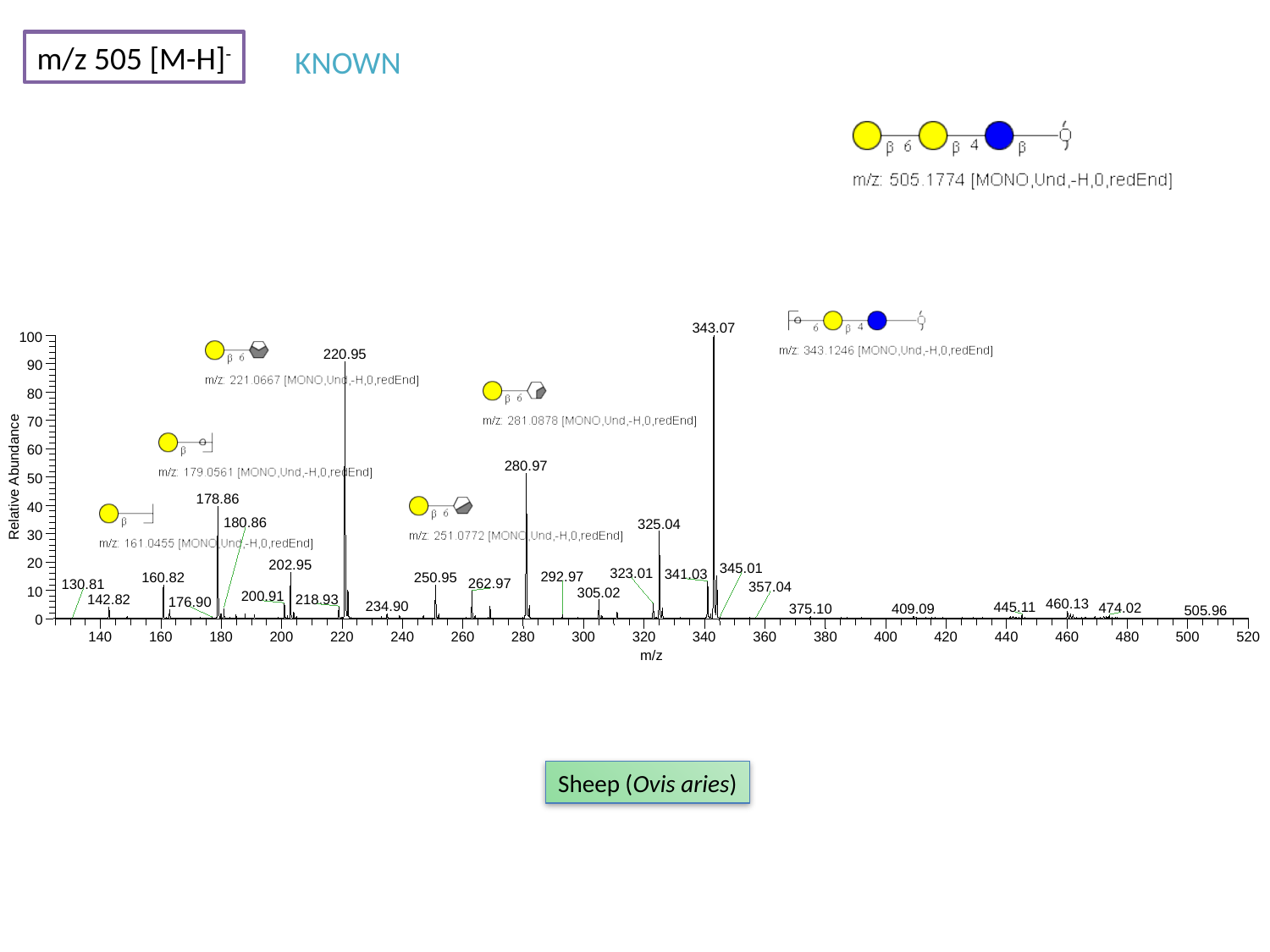

m/z 505 [M-H]^-^
KNOWN
343.07
100
220.95
90
80
70
60
280.97
Relative Abundance
50
178.86
40
325.04
30
20
202.95
323.01
341.03
160.82
250.95
262.97
10
305.02
200.91
0
140
160
180
200
220
240
260
280
300
320
340
360
380
400
420
440
460
480
500
520
m/z
180.86
345.01
292.97
130.81
357.04
218.93
142.82
176.90
460.13
234.90
445.11
474.02
409.09
375.10
505.96
Sheep (Ovis aries)

### Slide 14
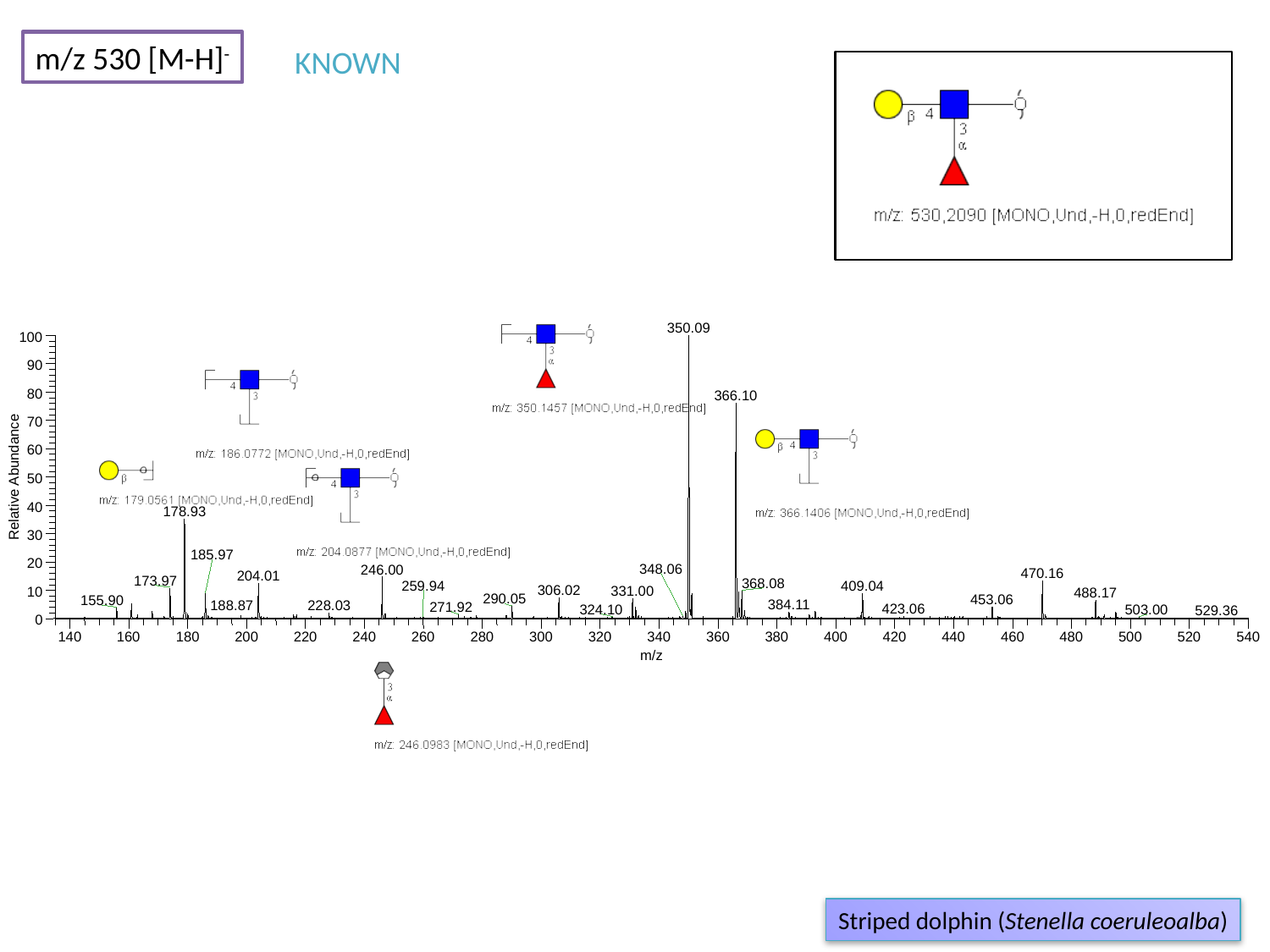

m/z 530 [M-H]^-^
KNOWN
350.09
100
90
80
366.10
70
60
Relative Abundance
50
40
178.93
30
185.97
20
246.00
470.16
204.01
173.97
368.08
409.04
10
488.17
0
140
160
180
200
220
240
260
280
300
320
340
360
380
400
420
440
460
480
500
520
540
m/z
348.06
259.94
306.02
331.00
290.05
453.06
155.90
384.11
188.87
228.03
271.92
423.06
324.10
503.00
529.36
Striped dolphin (Stenella coeruleoalba)

### Slide 15
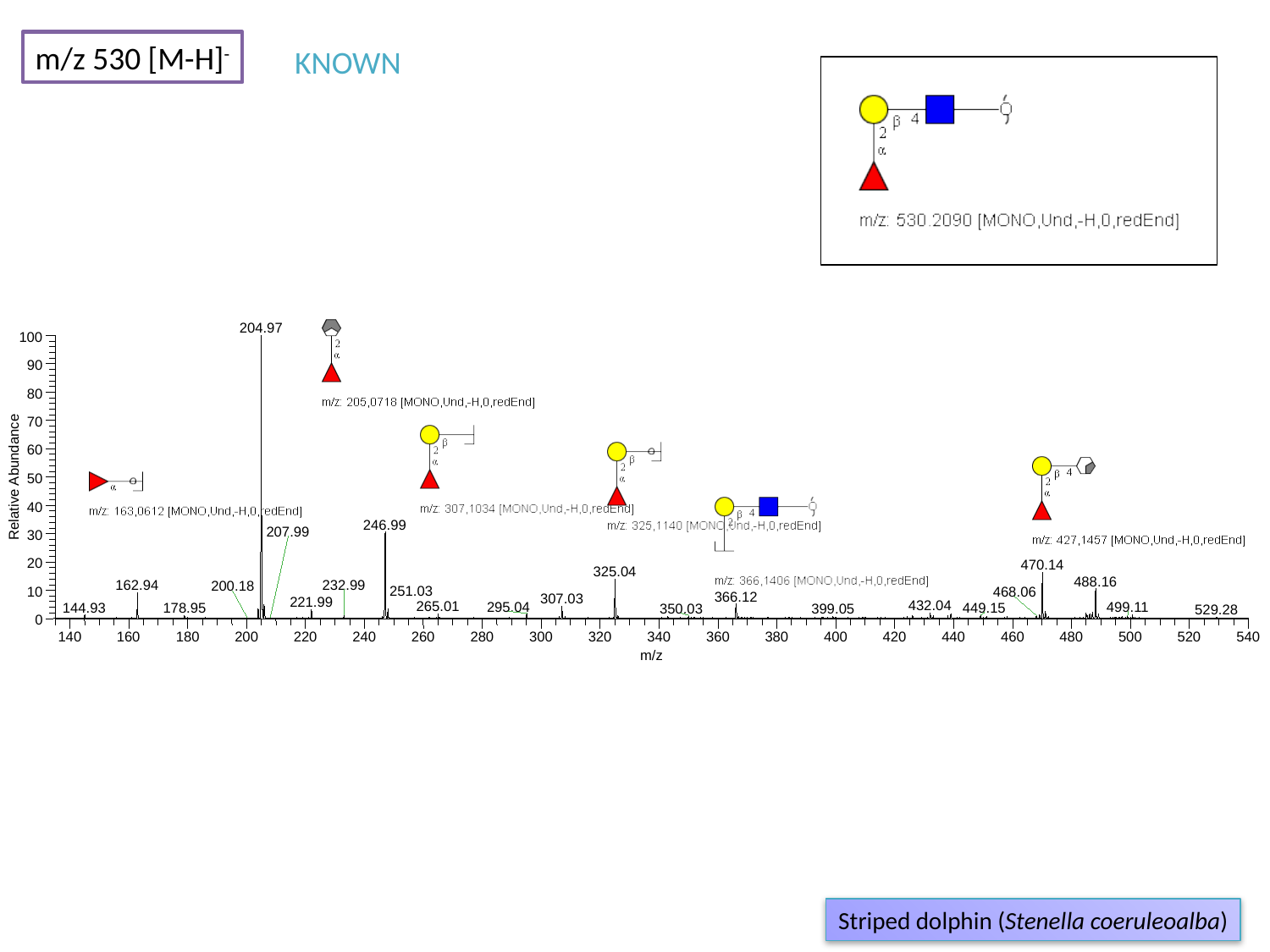

m/z 530 [M-H]^-^
KNOWN
204.97
100
90
80
70
60
Relative Abundance
50
40
246.99
30
20
470.14
325.04
488.16
162.94
10
366.12
307.03
221.99
432.04
265.01
295.04
144.93
0
140
160
180
200
220
240
260
280
300
320
340
360
380
400
420
440
460
480
500
520
540
m/z
207.99
232.99
200.18
251.03
468.06
499.11
178.95
449.15
350.03
399.05
529.28
Striped dolphin (Stenella coeruleoalba)

### Slide 16
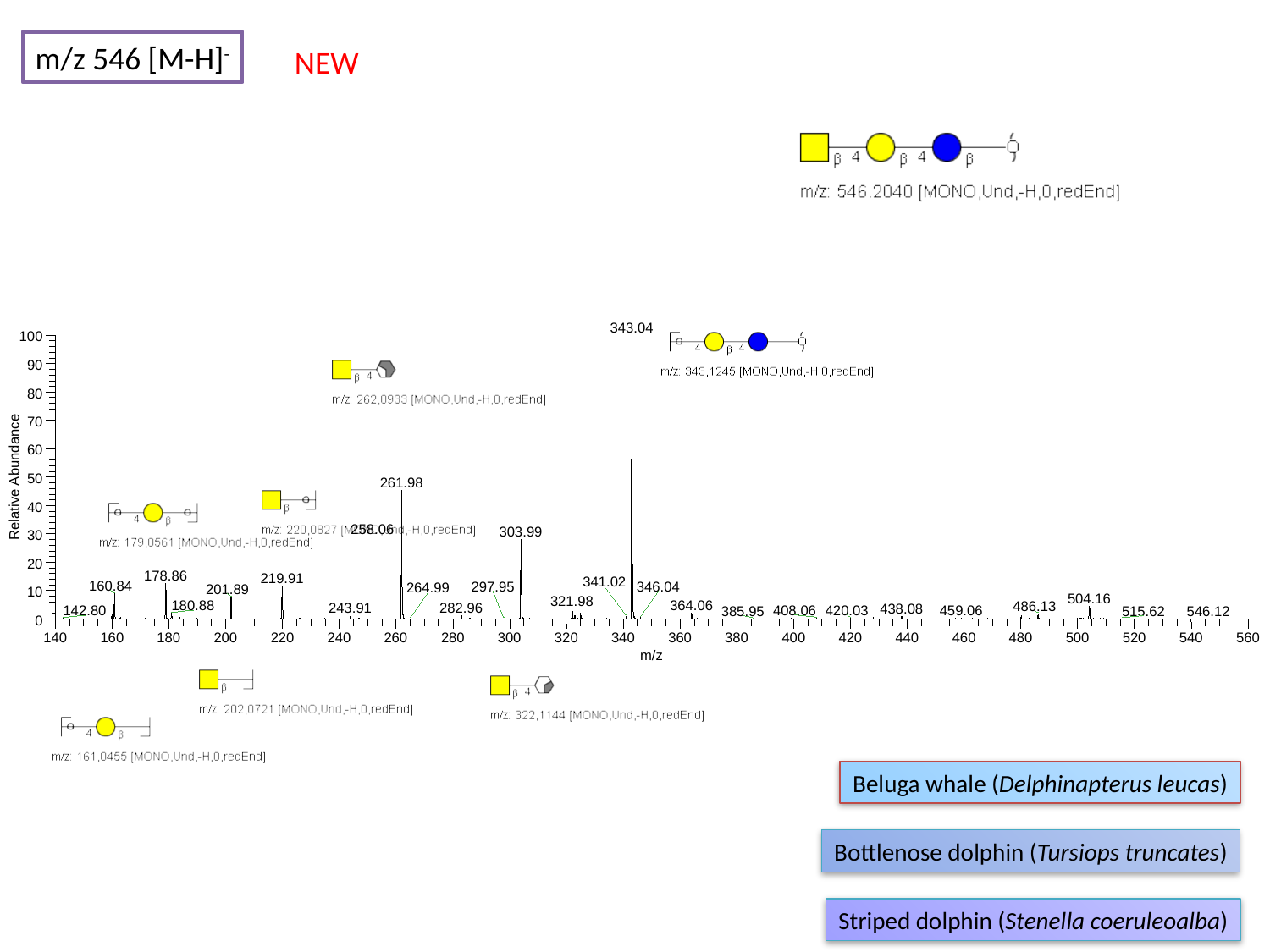

m/z 546 [M-H]^-^
NEW
343.04
100
90
80
70
60
Relative Abundance
50
261.98
40
303.99
30
20
178.86
219.91
160.84
201.89
10
504.16
0
140
160
180
200
220
240
260
280
300
320
340
360
380
400
420
440
460
480
500
520
540
560
m/z
258.06
341.02
346.04
297.95
264.99
321.98
364.06
180.88
486.13
243.91
282.96
438.08
459.06
408.06
420.03
142.80
385.95
515.62
546.12
Beluga whale (Delphinapterus leucas)
Bottlenose dolphin (Tursiops truncates)
Striped dolphin (Stenella coeruleoalba)

### Slide 17
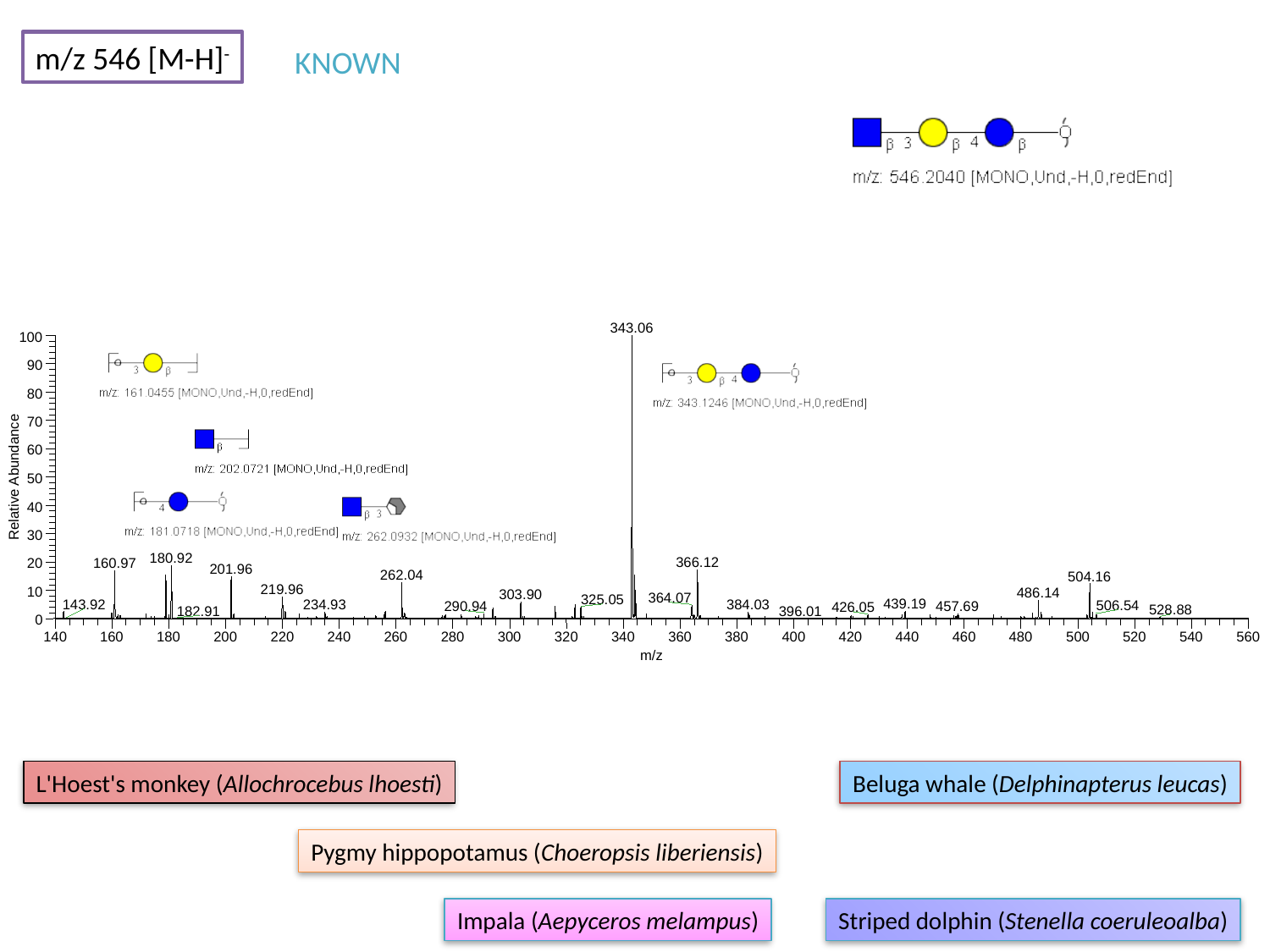

m/z 546 [M-H]^-^
KNOWN
343.06
100
90
80
70
60
Relative Abundance
50
40
30
180.92
366.12
20
160.97
201.96
262.04
504.16
219.96
10
486.14
303.90
0
140
160
180
200
220
240
260
280
300
320
340
360
380
400
420
440
460
480
500
520
540
560
m/z
364.07
325.05
439.19
143.92
384.03
234.93
506.54
290.94
457.69
426.05
528.88
182.91
396.01
L'Hoest's monkey (Allochrocebus lhoesti)
Beluga whale (Delphinapterus leucas)
Pygmy hippopotamus (Choeropsis liberiensis)
Impala (Aepyceros melampus)
Striped dolphin (Stenella coeruleoalba)

### Slide 18
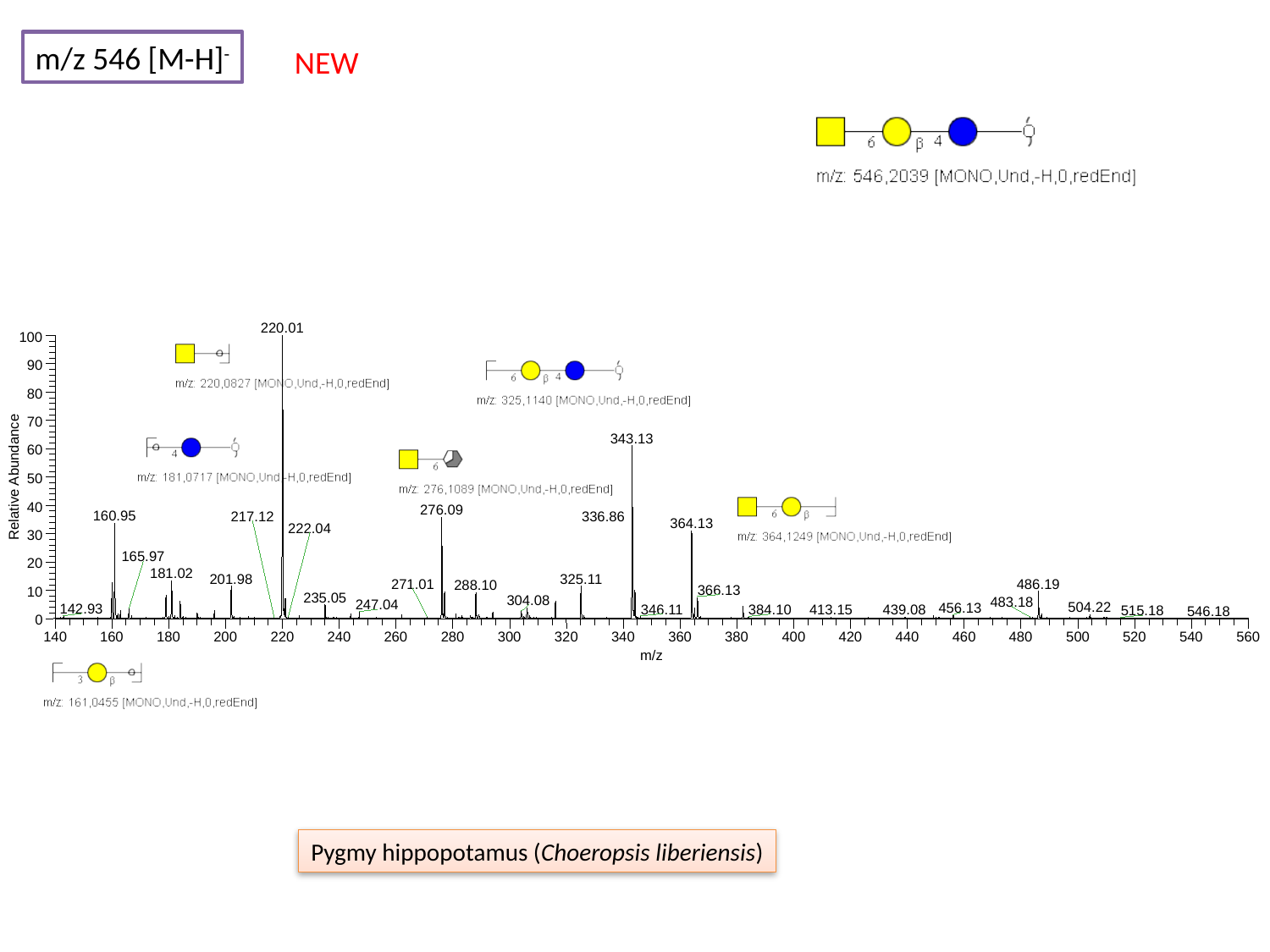

m/z 546 [M-H]^-^
NEW
220.01
100
90
80
70
343.13
60
Relative Abundance
50
40
276.09
160.95
364.13
30
20
181.02
201.98
325.11
486.19
288.10
10
0
140
160
180
200
220
240
260
280
300
320
340
360
380
400
420
440
460
480
500
520
540
560
m/z
217.12
336.86
222.04
165.97
271.01
366.13
235.05
304.08
483.18
247.04
504.22
456.13
142.93
346.11
384.10
413.15
439.08
515.18
546.18
Pygmy hippopotamus (Choeropsis liberiensis)

### Slide 19
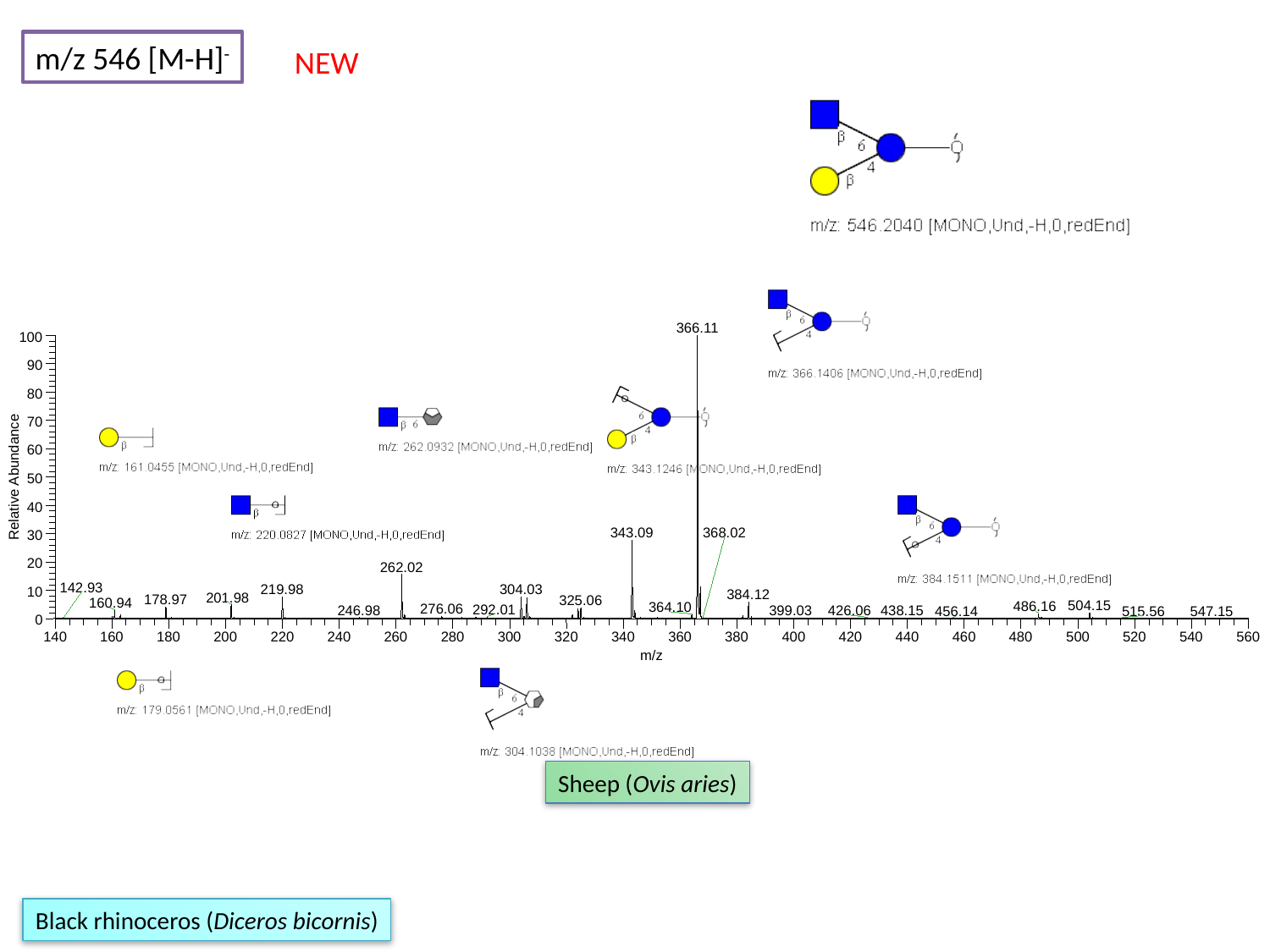

m/z 546 [M-H]^-^
NEW
366.11
100
90
80
70
60
Relative Abundance
50
40
343.09
30
20
262.02
304.03
219.98
10
384.12
201.98
178.97
325.06
0
140
160
180
200
220
240
260
280
300
320
340
360
380
400
420
440
460
480
500
520
540
560
m/z
368.02
142.93
160.94
504.15
486.16
364.10
276.06
292.01
246.98
438.15
426.06
399.03
456.14
515.56
547.15
Sheep (Ovis aries)
Black rhinoceros (Diceros bicornis)

### Slide 20
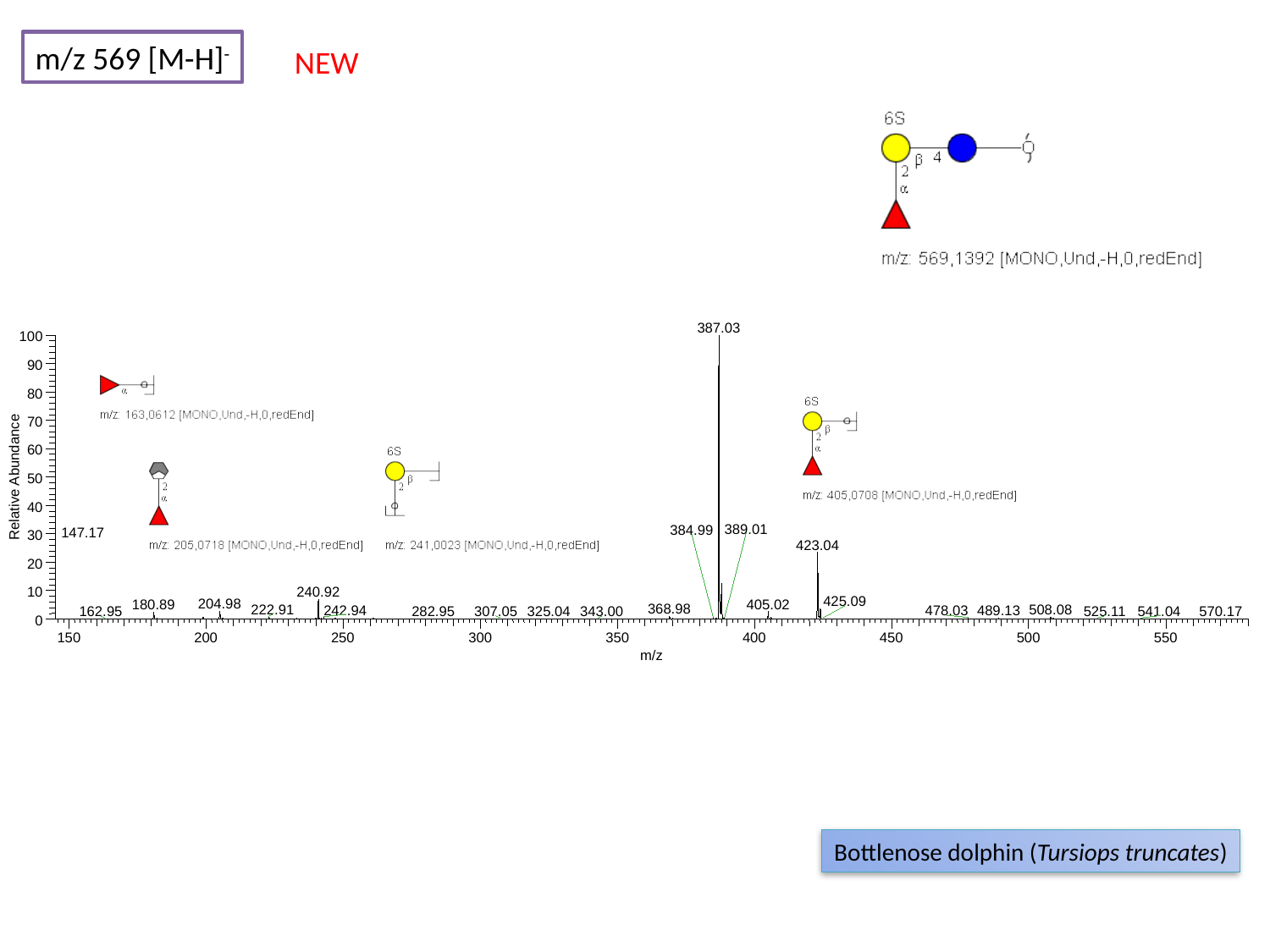

m/z 569 [M-H]^-^
NEW
387.03
100
90
80
70
60
Relative Abundance
50
40
389.01
384.99
147.17
30
423.04
20
240.92
10
425.09
204.98
405.02
180.89
368.98
508.08
222.91
242.94
489.13
478.03
307.05
325.04
162.95
282.95
343.00
525.11
541.04
570.17
0
150
200
250
300
350
400
450
500
550
m/z
Bottlenose dolphin (Tursiops truncates)

### Slide 21
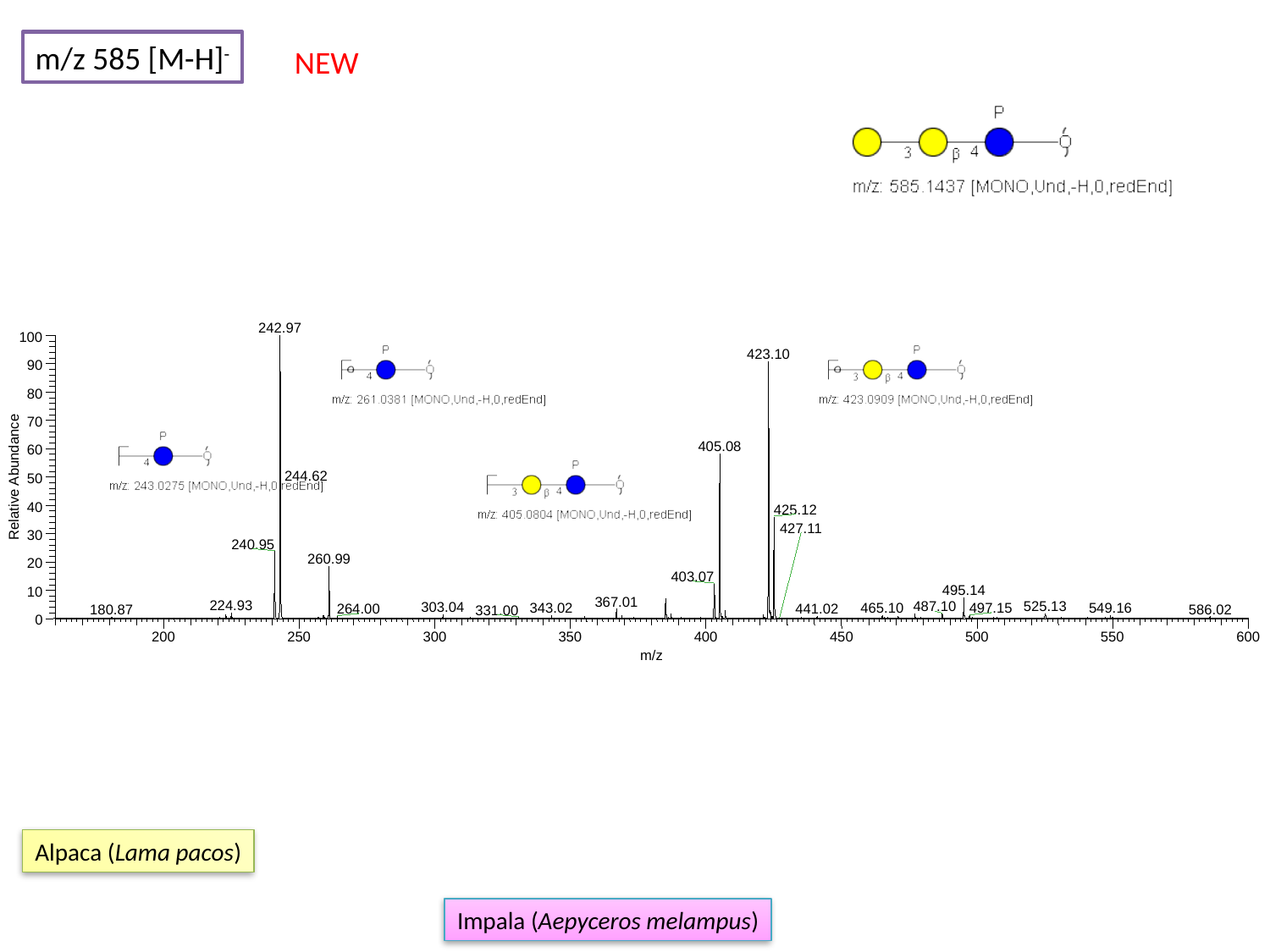

m/z 585 [M-H]^-^
NEW
242.97
100
423.10
90
80
70
405.08
60
244.62
Relative Abundance
50
40
425.12
427.11
30
240.95
260.99
20
403.07
495.14
10
367.01
224.93
525.13
487.10
303.04
465.10
549.16
343.02
497.15
441.02
264.00
586.02
180.87
331.00
0
200
250
300
350
400
450
500
550
600
m/z
Alpaca (Lama pacos)
Impala (Aepyceros melampus)

### Slide 22
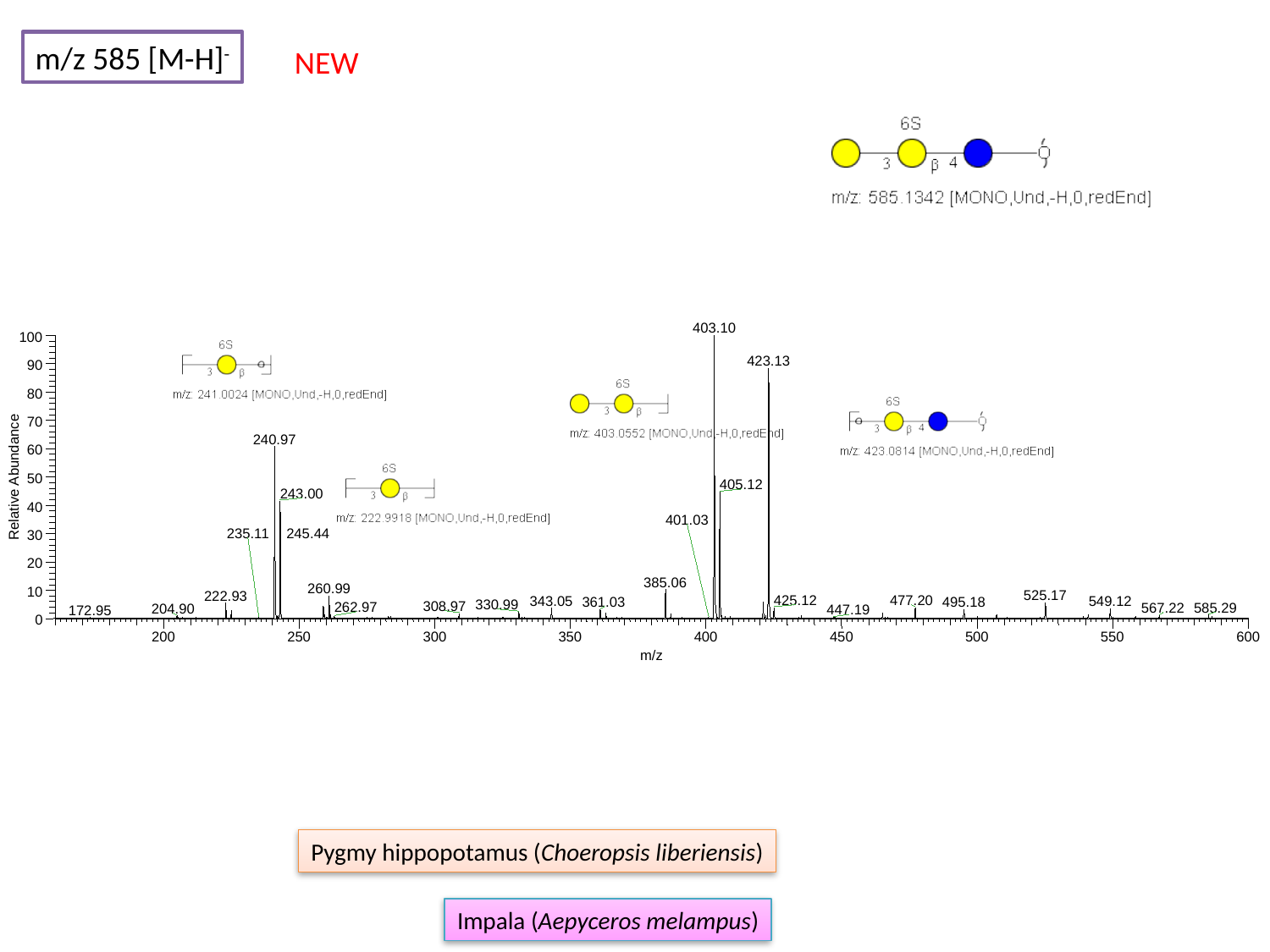

m/z 585 [M-H]^-^
NEW
403.10
100
423.13
90
80
70
240.97
60
Relative Abundance
50
405.12
243.00
40
401.03
235.11
245.44
30
20
385.06
260.99
10
525.17
222.93
425.12
477.20
343.05
549.12
361.03
495.18
330.99
308.97
262.97
585.29
567.22
204.90
447.19
172.95
0
200
250
300
350
400
450
500
550
600
m/z
Pygmy hippopotamus (Choeropsis liberiensis)
Impala (Aepyceros melampus)

### Slide 23
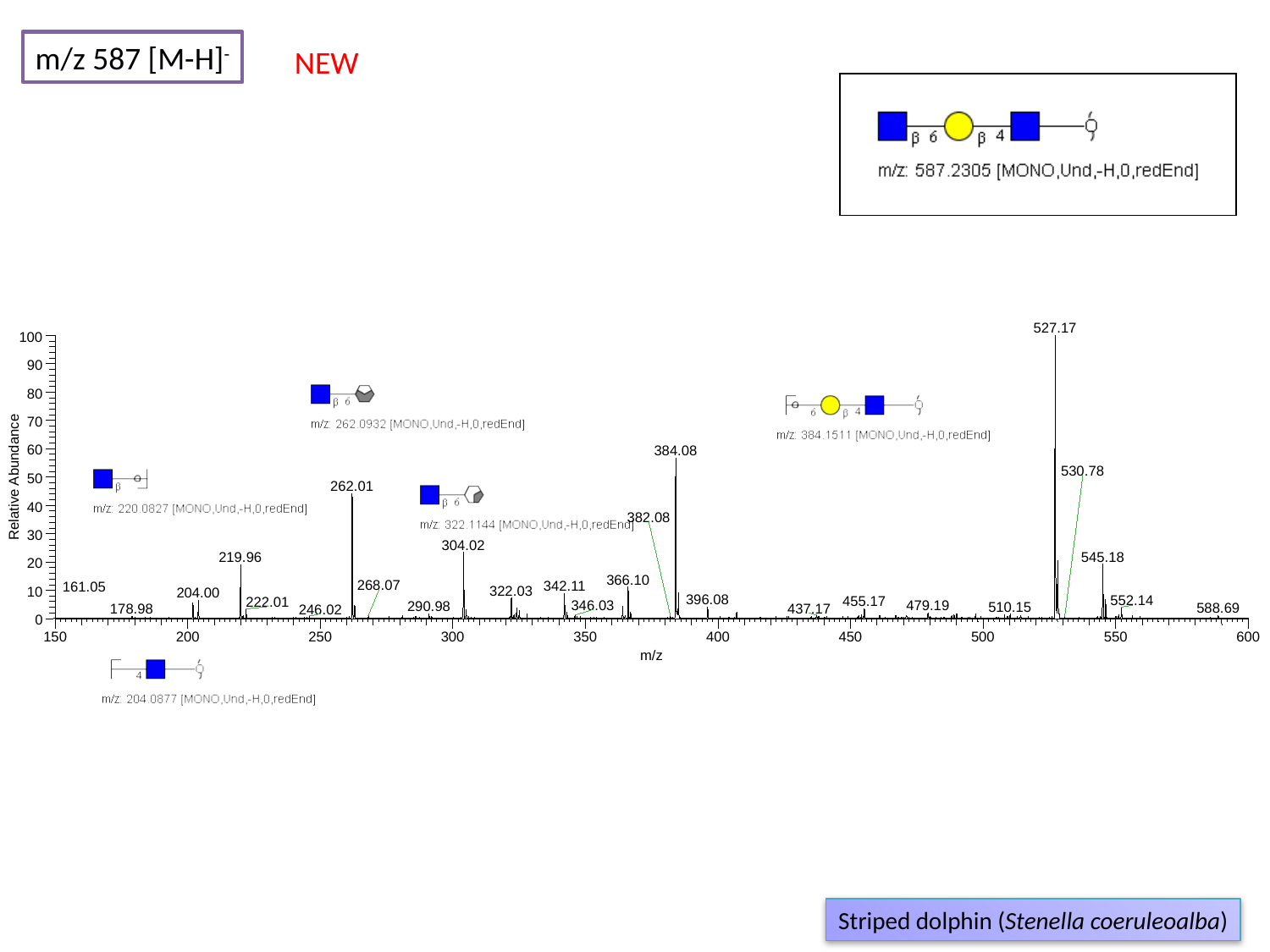

m/z 587 [M-H]^-^
NEW
527.17
100
90
80
70
60
384.08
530.78
Relative Abundance
50
262.01
40
382.08
30
304.02
545.18
219.96
20
366.10
268.07
342.11
161.05
322.03
10
204.00
396.08
552.14
455.17
222.01
346.03
479.19
290.98
510.15
588.69
437.17
178.98
246.02
0
150
200
250
300
350
400
450
500
550
600
m/z
Striped dolphin (Stenella coeruleoalba)

### Slide 24
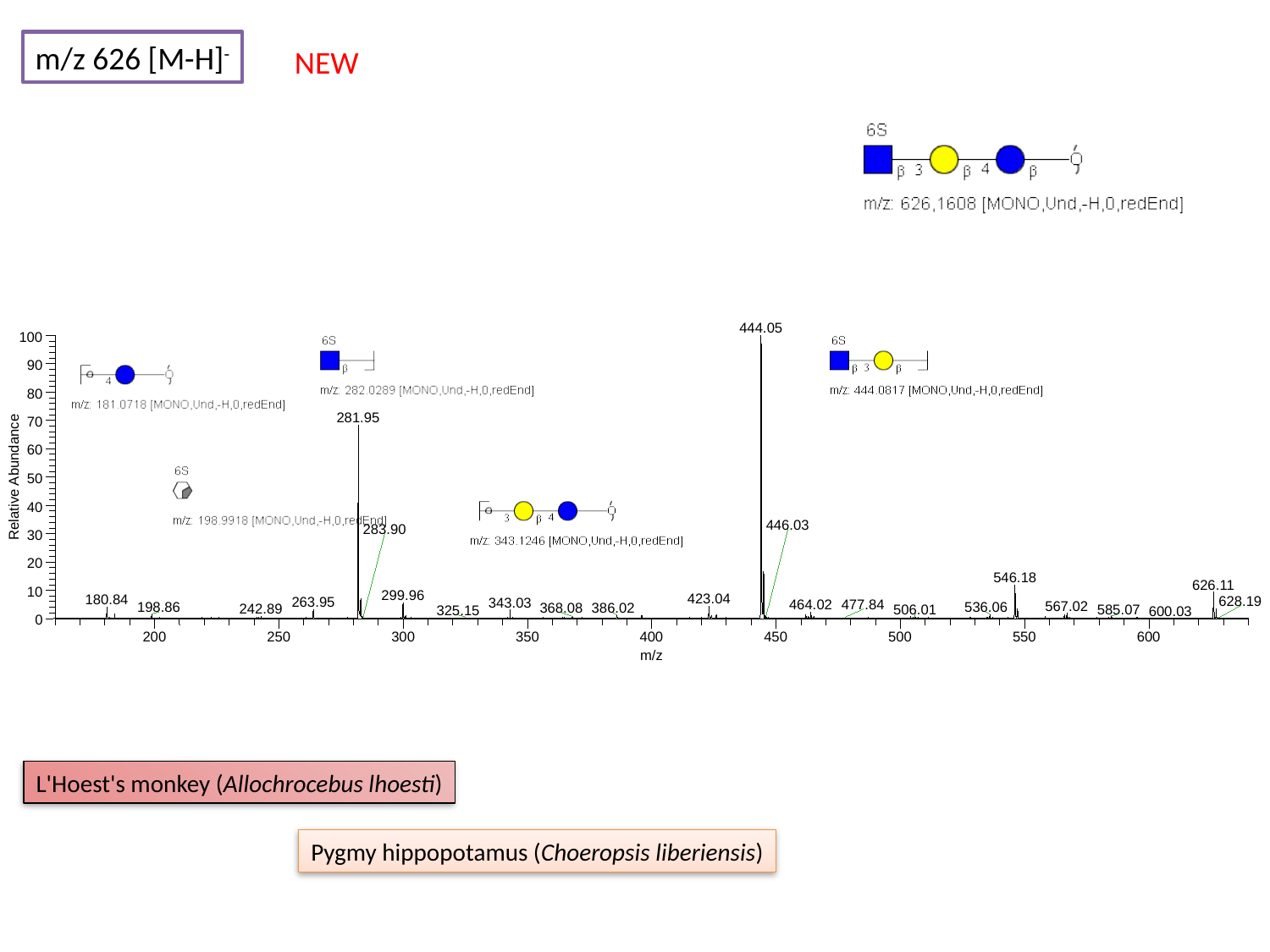

m/z 626 [M-H]^-^
NEW
444.05
100
90
80
281.95
70
60
Relative Abundance
50
40
446.03
283.90
30
20
546.18
626.11
10
299.96
423.04
180.84
628.19
263.95
343.03
464.02
477.84
567.02
198.86
536.06
368.08
386.02
242.89
506.01
585.07
325.15
600.03
0
200
250
300
350
400
450
500
550
600
m/z
L'Hoest's monkey (Allochrocebus lhoesti)
Pygmy hippopotamus (Choeropsis liberiensis)

### Slide 25
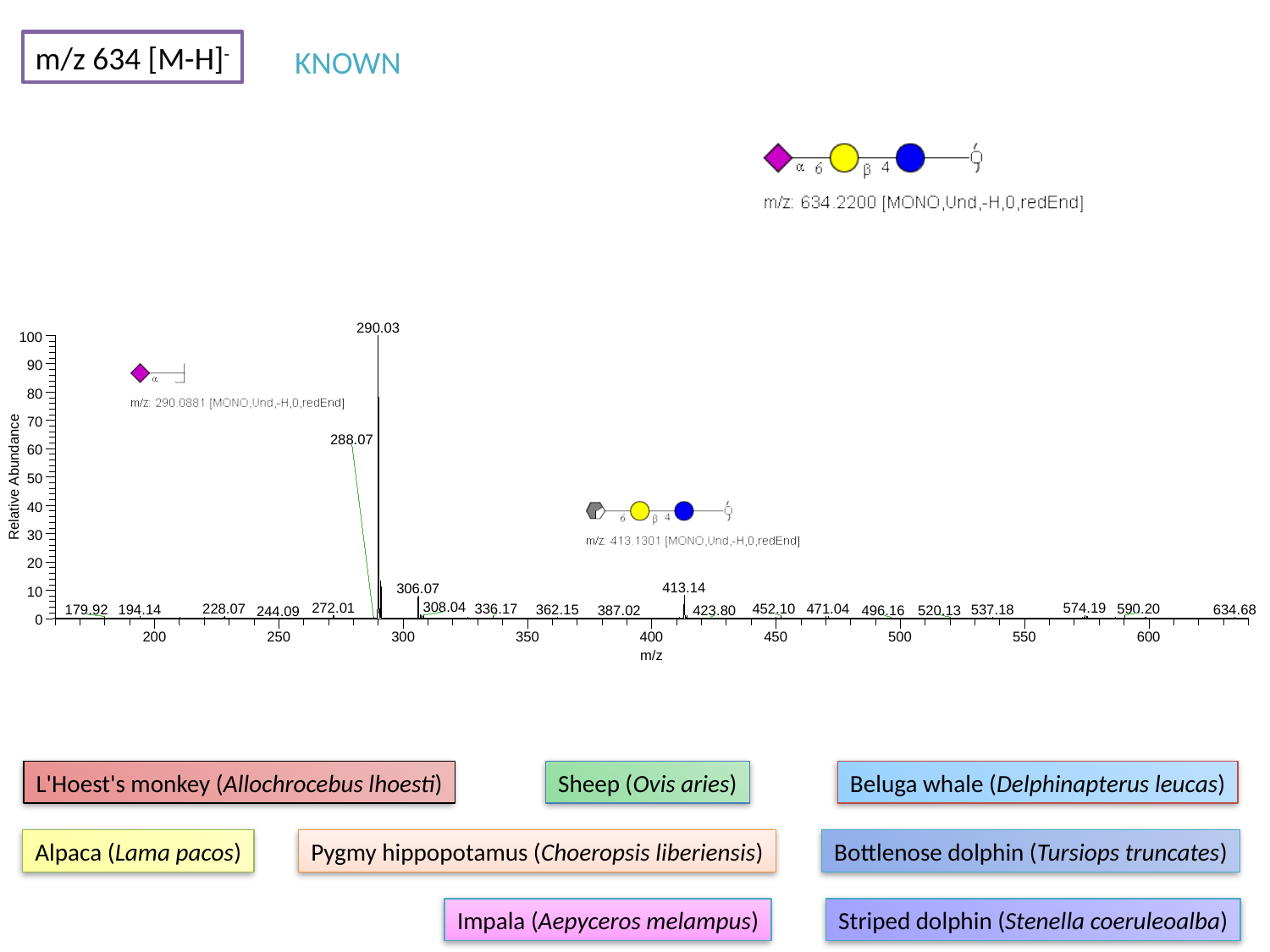

m/z 634 [M-H]^-^
KNOWN
290.03
100
90
80
70
288.07
60
Relative Abundance
50
40
30
20
413.14
306.07
10
308.04
272.01
574.19
590.20
452.10
471.04
228.07
336.17
194.14
179.92
362.15
537.18
634.68
387.02
520.13
423.80
496.16
244.09
0
200
250
300
350
400
450
500
550
600
m/z
L'Hoest's monkey (Allochrocebus lhoesti)
Sheep (Ovis aries)
Beluga whale (Delphinapterus leucas)
Alpaca (Lama pacos)
Pygmy hippopotamus (Choeropsis liberiensis)
Bottlenose dolphin (Tursiops truncates)
Impala (Aepyceros melampus)
Striped dolphin (Stenella coeruleoalba)

### Slide 26
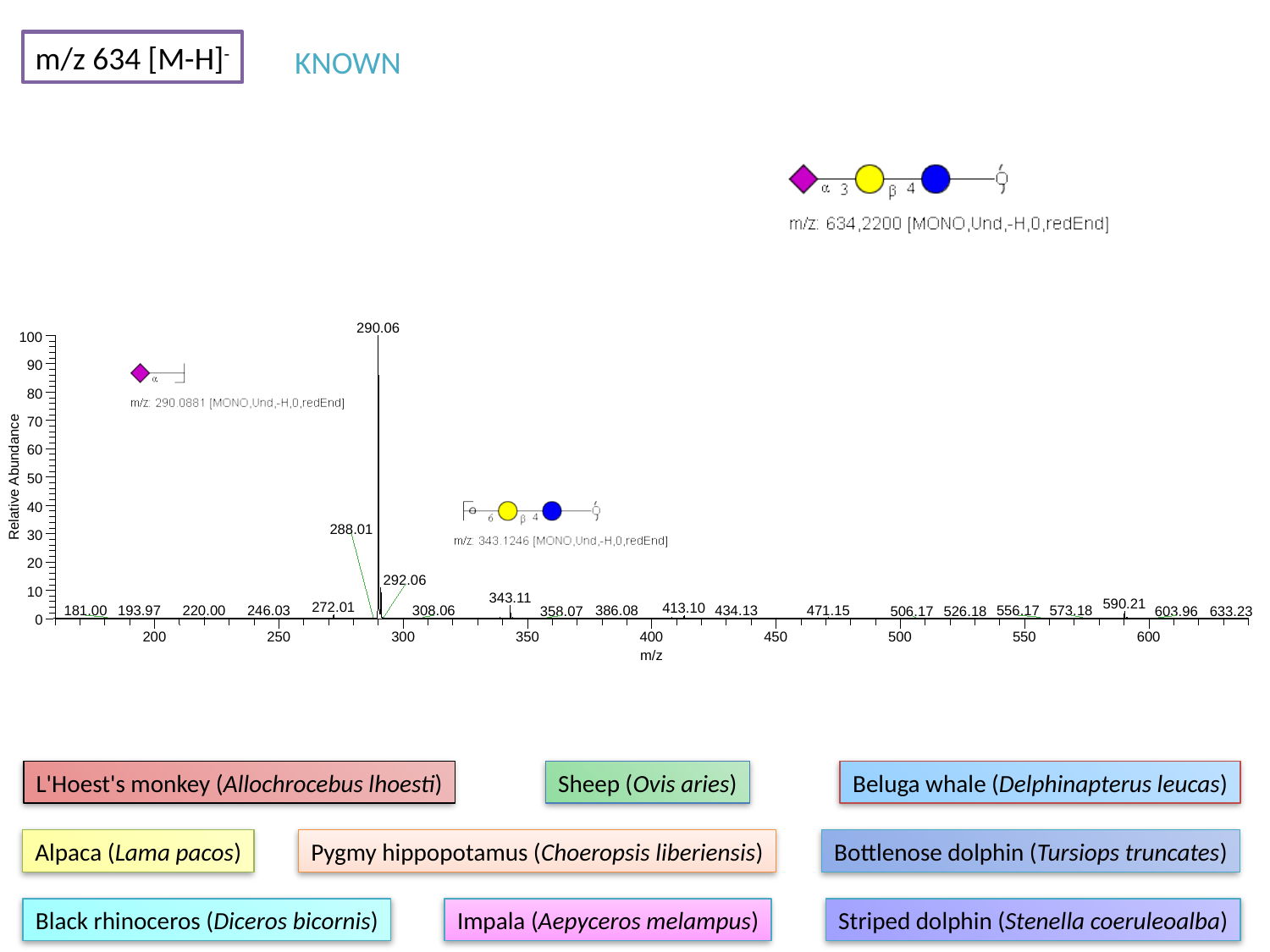

m/z 634 [M-H]^-^
KNOWN
290.06
100
90
80
70
60
Relative Abundance
50
40
288.01
30
20
292.06
10
343.11
590.21
272.01
413.10
220.00
471.15
246.03
434.13
193.97
386.08
181.00
308.06
556.17
573.18
358.07
506.17
526.18
603.96
633.23
0
200
250
300
350
400
450
500
550
600
m/z
L'Hoest's monkey (Allochrocebus lhoesti)
Sheep (Ovis aries)
Beluga whale (Delphinapterus leucas)
Alpaca (Lama pacos)
Pygmy hippopotamus (Choeropsis liberiensis)
Bottlenose dolphin (Tursiops truncates)
Black rhinoceros (Diceros bicornis)
Impala (Aepyceros melampus)
Striped dolphin (Stenella coeruleoalba)

### Slide 27
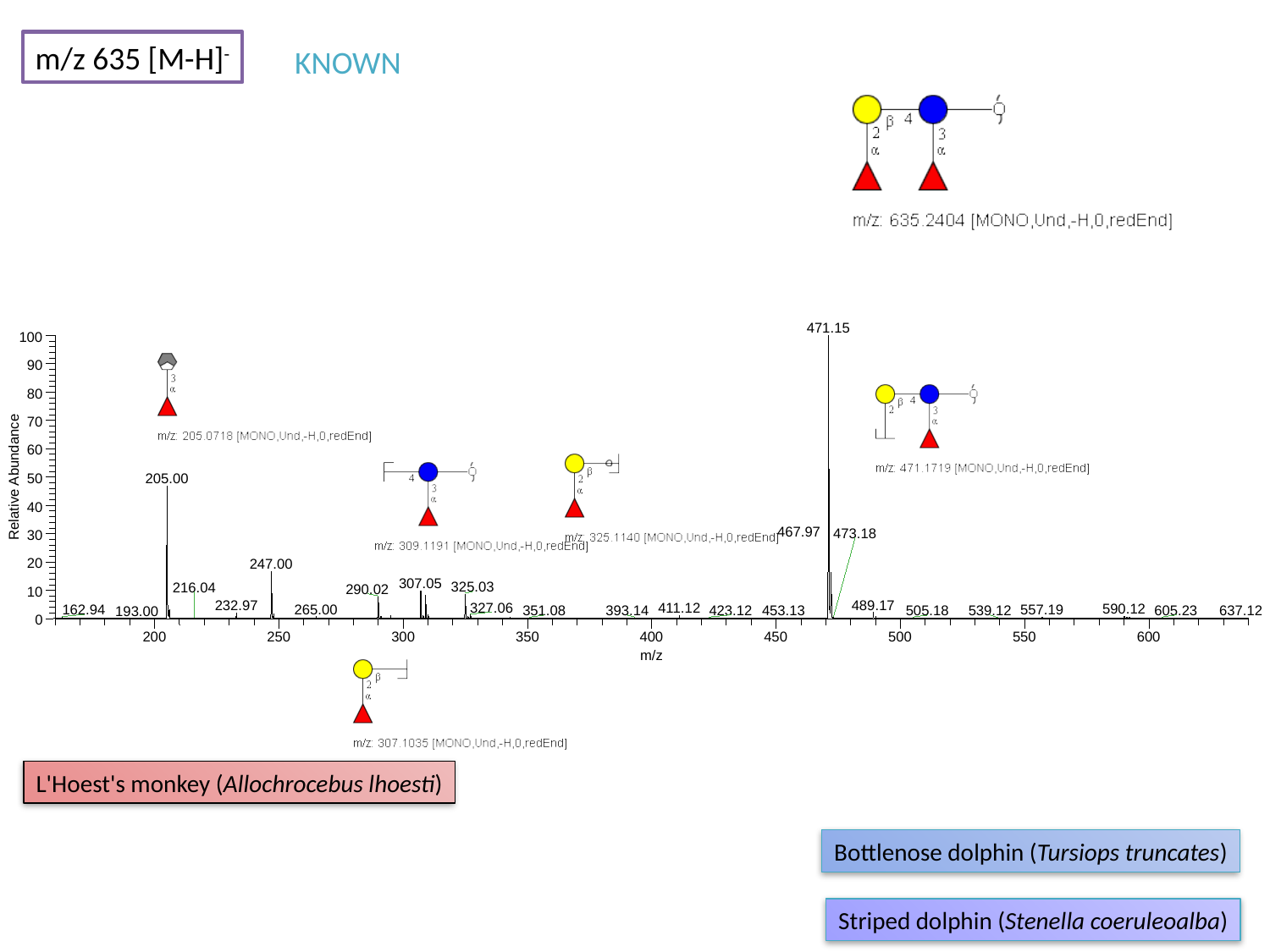

m/z 635 [M-H]^-^
KNOWN
471.15
100
90
80
70
60
Relative Abundance
205.00
50
40
467.97
473.18
30
20
247.00
307.05
325.03
216.04
290.02
10
489.17
232.97
327.06
411.12
590.12
265.00
162.94
557.19
423.12
453.13
605.23
351.08
393.14
505.18
539.12
637.12
193.00
0
200
250
300
350
400
450
500
550
600
m/z
L'Hoest's monkey (Allochrocebus lhoesti)
Bottlenose dolphin (Tursiops truncates)
Striped dolphin (Stenella coeruleoalba)

### Slide 28
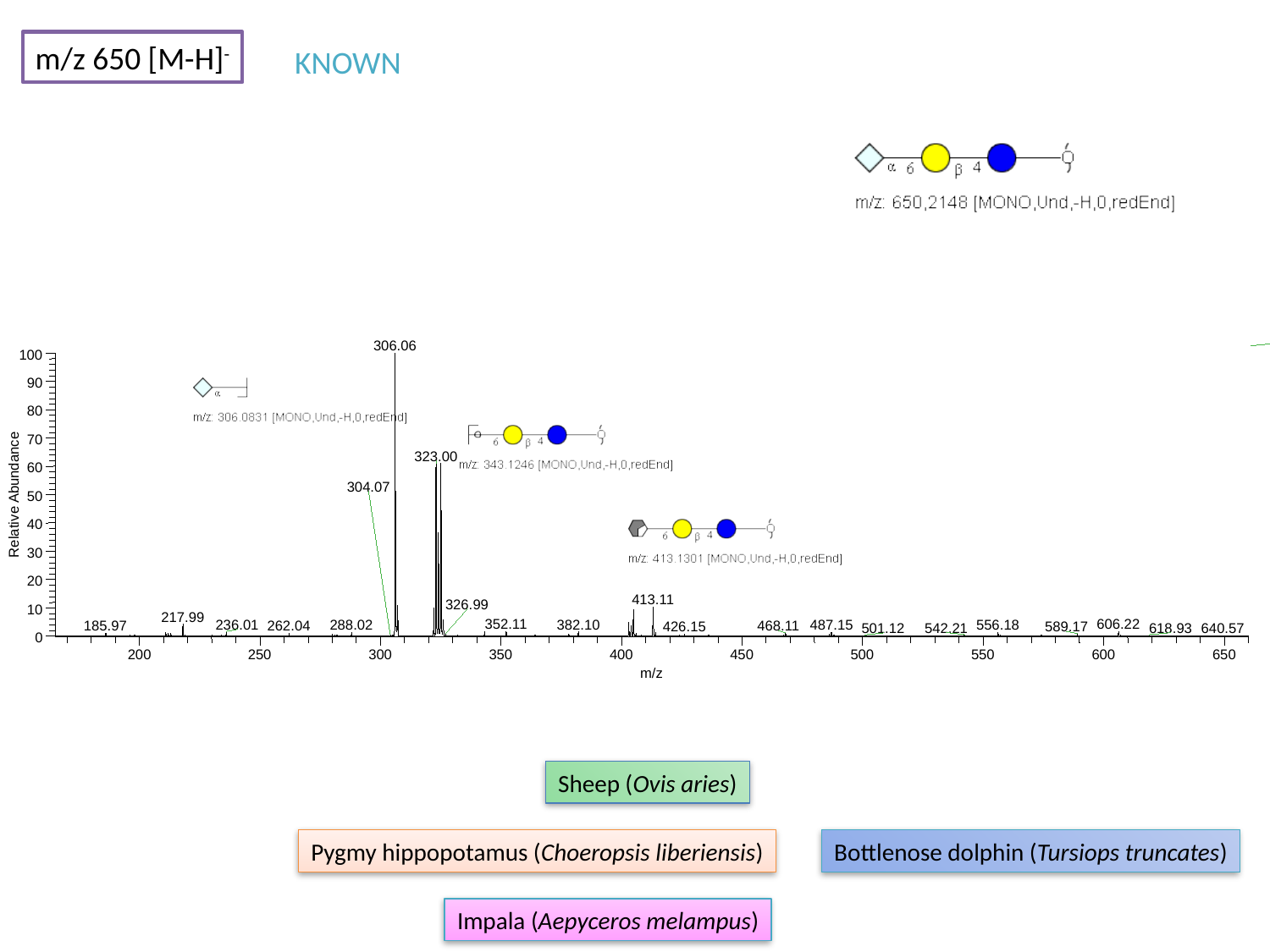

m/z 650 [M-H]^-^
KNOWN
306.06
100
90
80
70
323.00
60
304.07
Relative Abundance
50
40
30
20
413.11
326.99
10
217.99
352.11
606.22
382.10
236.01
288.02
487.15
556.18
185.97
262.04
468.11
426.15
589.17
542.21
618.93
501.12
640.57
0
200
250
300
350
400
450
500
550
600
650
m/z
Sheep (Ovis aries)
Pygmy hippopotamus (Choeropsis liberiensis)
Bottlenose dolphin (Tursiops truncates)
Impala (Aepyceros melampus)

### Slide 29
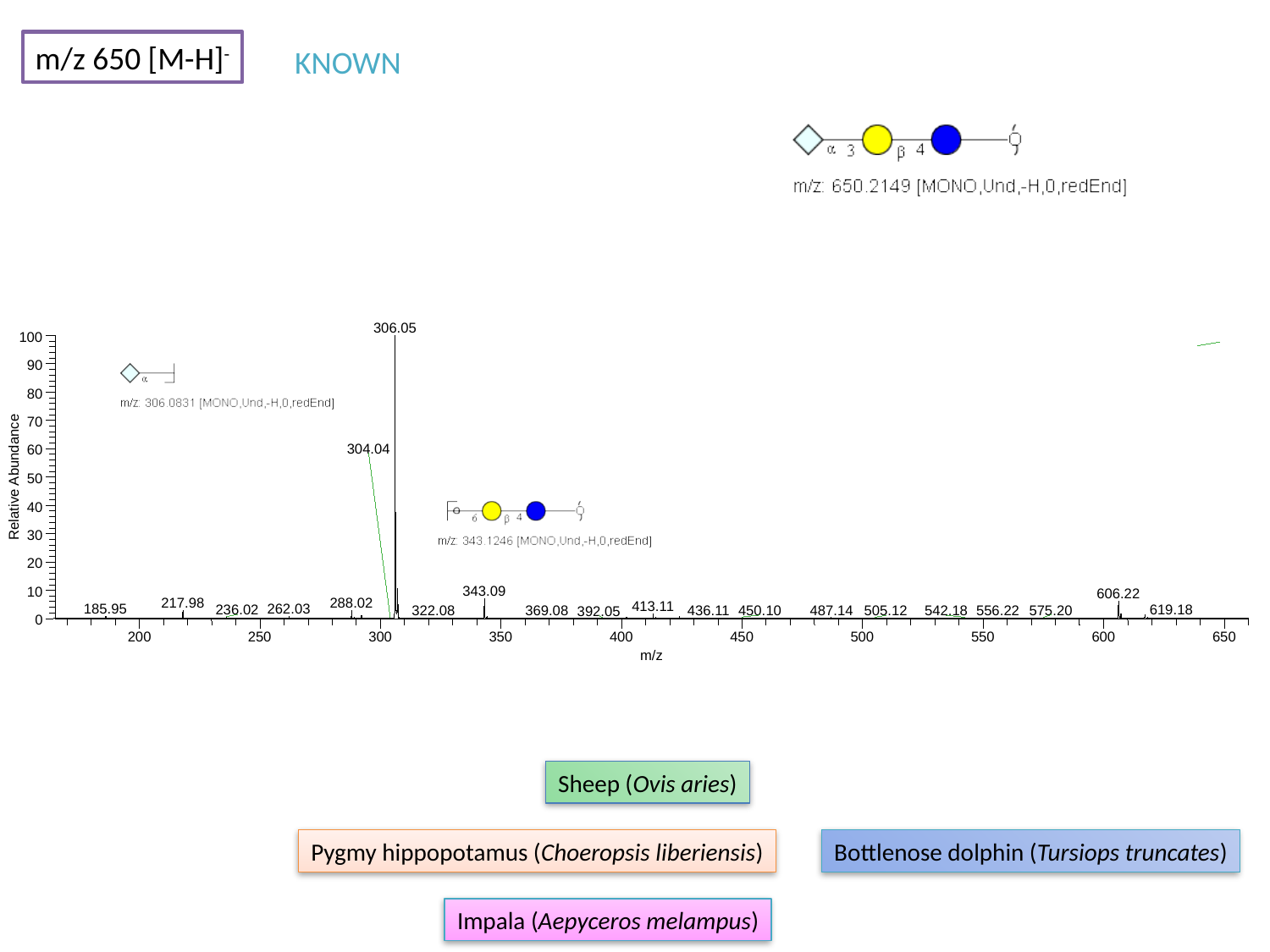

m/z 650 [M-H]^-^
KNOWN
306.05
100
90
80
70
304.04
60
Relative Abundance
50
40
30
20
343.09
10
606.22
288.02
217.98
413.11
185.95
262.03
619.18
236.02
487.14
436.11
556.22
575.20
450.10
322.08
369.08
505.12
542.18
392.05
0
200
250
300
350
400
450
500
550
600
650
m/z
Sheep (Ovis aries)
Pygmy hippopotamus (Choeropsis liberiensis)
Bottlenose dolphin (Tursiops truncates)
Impala (Aepyceros melampus)

### Slide 30
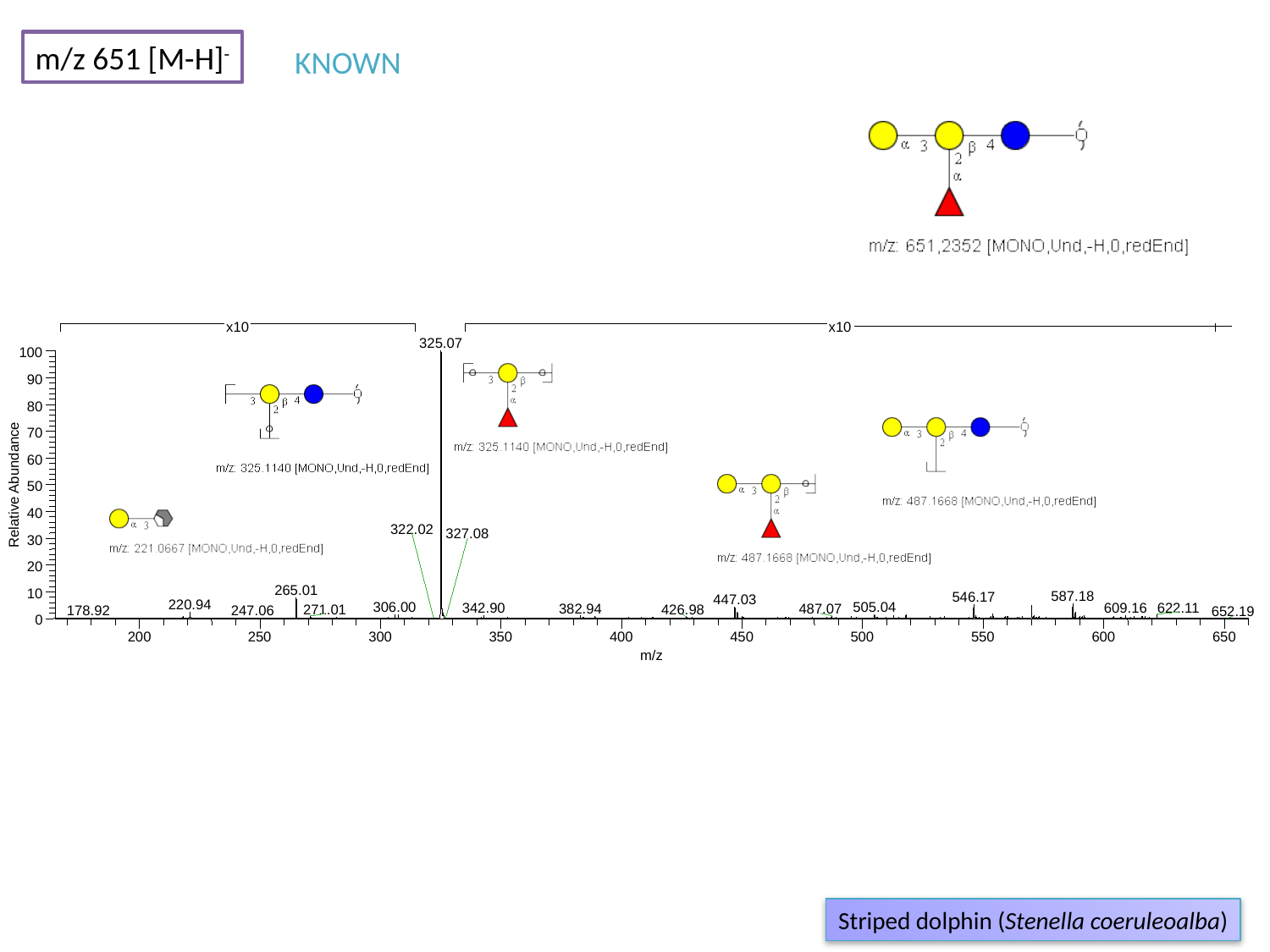

m/z 651 [M-H]^-^
KNOWN
x10
x10
325.07
100
90
80
70
60
Relative Abundance
50
40
322.02
327.08
30
20
265.01
10
587.18
546.17
447.03
220.94
306.00
505.04
622.11
342.90
609.16
382.94
487.07
271.01
426.98
247.06
178.92
652.19
0
200
250
300
350
400
450
500
550
600
650
m/z
Striped dolphin (Stenella coeruleoalba)

### Slide 31
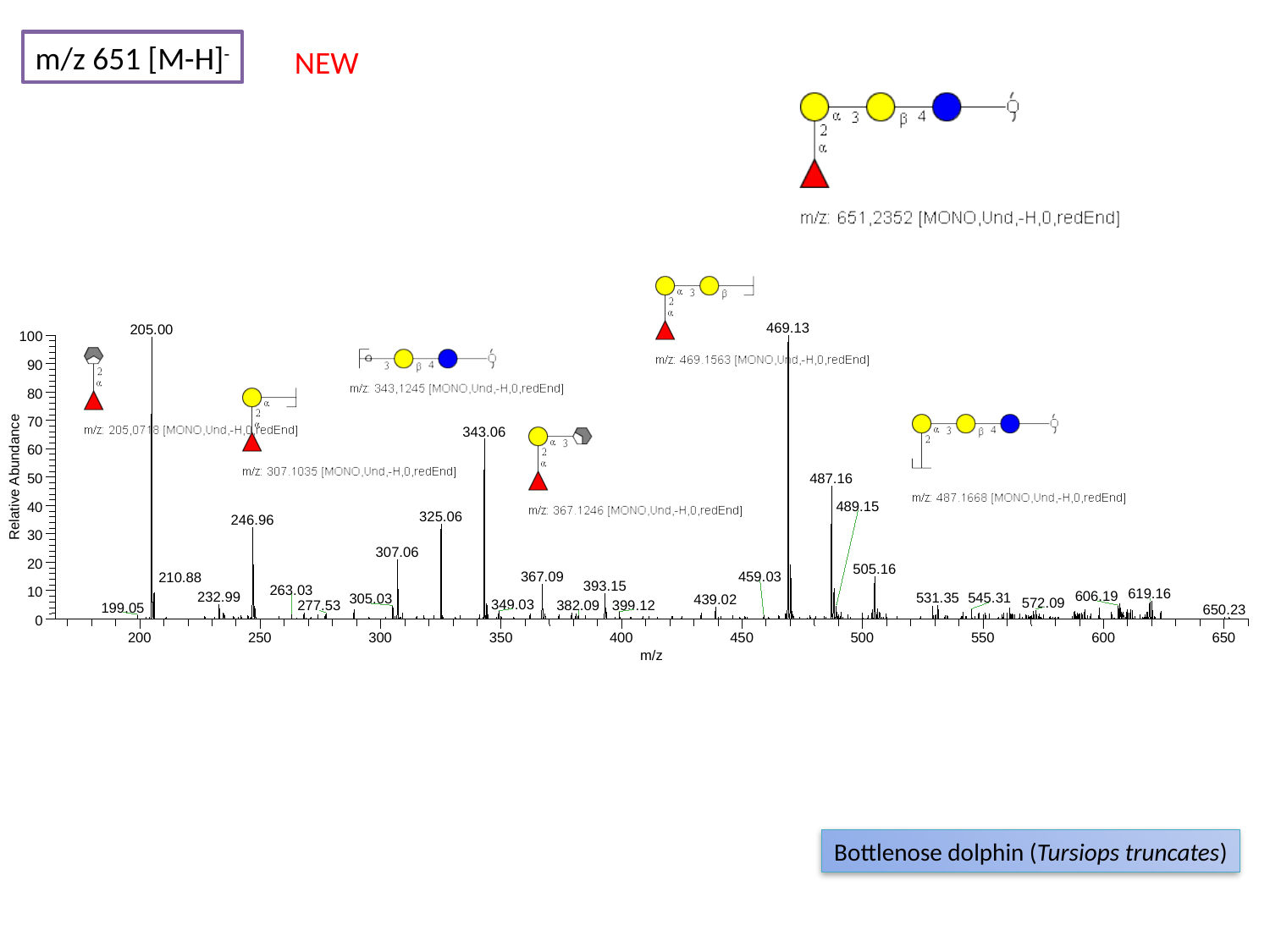

m/z 651 [M-H]^-^
NEW
469.13
205.00
100
90
80
70
343.06
60
Relative Abundance
50
487.16
489.15
40
325.06
246.96
30
307.06
20
505.16
367.09
459.03
210.88
393.15
263.03
10
619.16
606.19
232.99
531.35
545.31
305.03
439.02
572.09
349.03
399.12
277.53
382.09
199.05
650.23
0
200
250
300
350
400
450
500
550
600
650
m/z
Bottlenose dolphin (Tursiops truncates)

### Slide 32
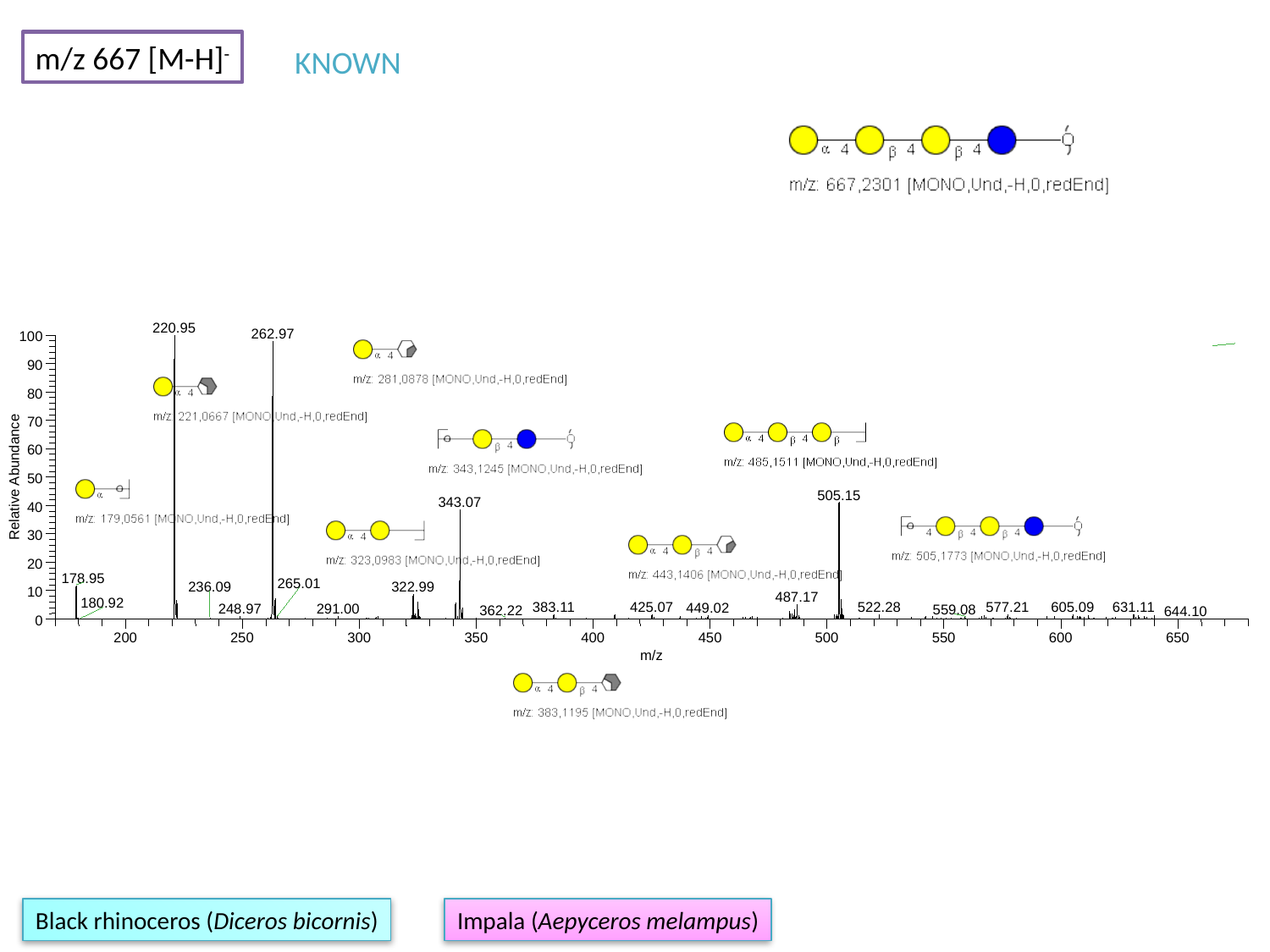

m/z 667 [M-H]^-^
KNOWN
220.95
262.97
100
90
80
70
60
Relative Abundance
50
505.15
343.07
40
30
20
178.95
265.01
322.99
236.09
10
487.17
180.92
425.07
631.11
383.11
605.09
522.28
577.21
449.02
248.97
291.00
559.08
362.22
644.10
0
200
250
300
350
400
450
500
550
600
650
m/z
Black rhinoceros (Diceros bicornis)
Impala (Aepyceros melampus)

### Slide 33
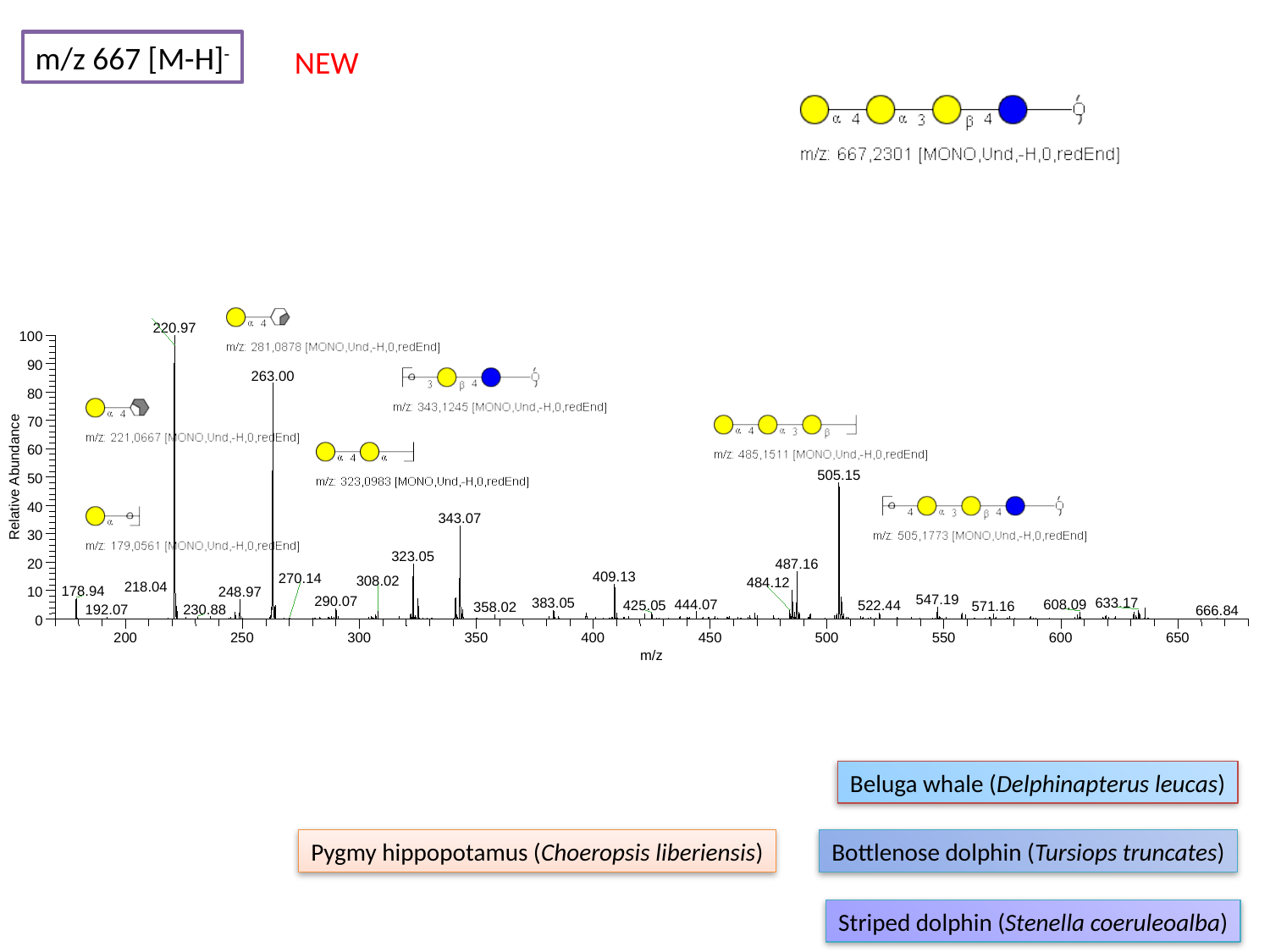

m/z 667 [M-H]^-^
NEW
220.97
100
90
263.00
80
70
60
505.15
Relative Abundance
50
40
343.07
30
323.05
20
487.16
409.13
270.14
308.02
484.12
218.04
178.94
10
248.97
547.19
290.07
633.17
383.05
444.07
608.09
522.44
425.05
571.16
358.02
230.88
192.07
666.84
0
200
250
300
350
400
450
500
550
600
650
m/z
Beluga whale (Delphinapterus leucas)
Pygmy hippopotamus (Choeropsis liberiensis)
Bottlenose dolphin (Tursiops truncates)
Striped dolphin (Stenella coeruleoalba)

### Slide 34
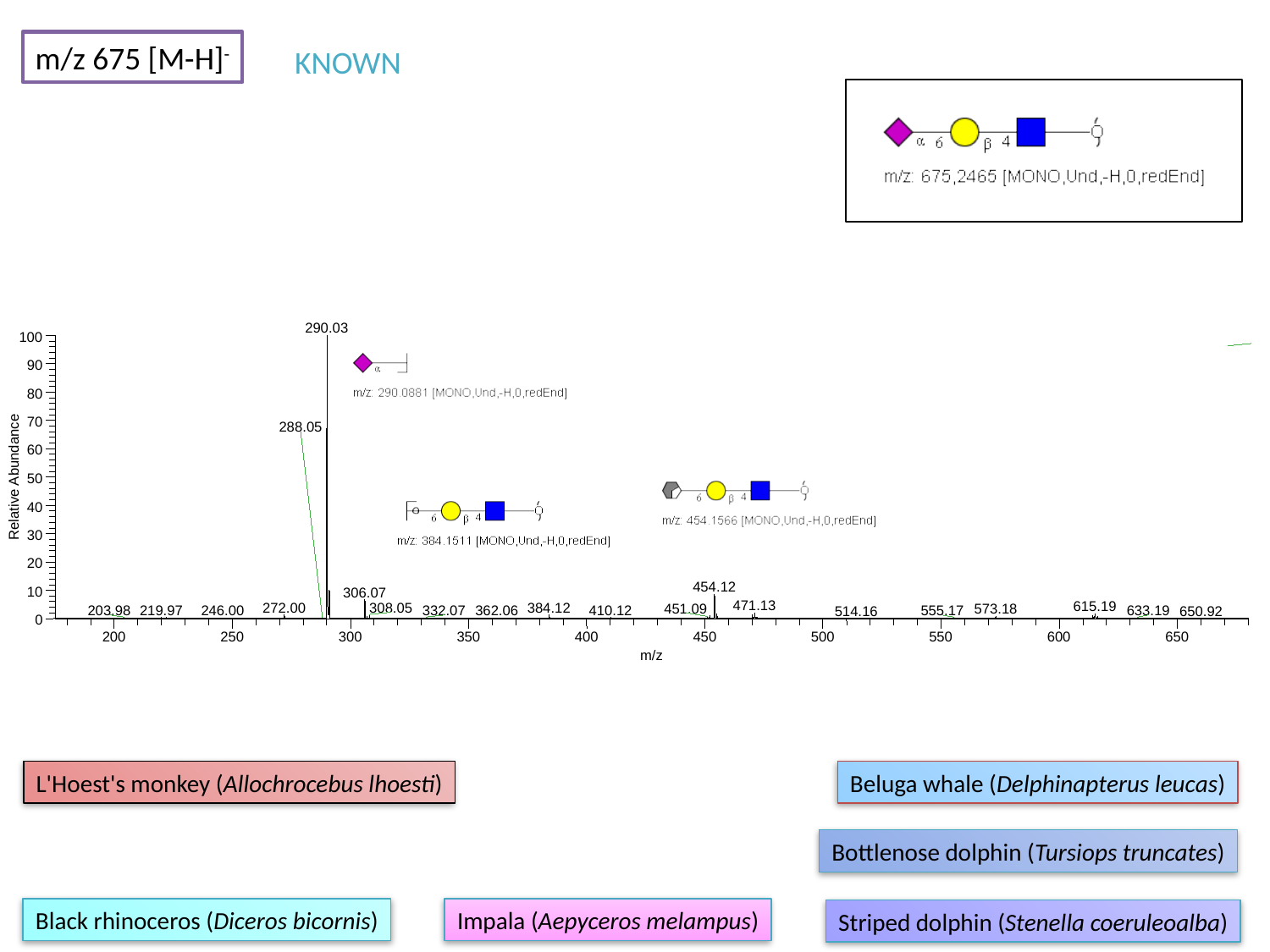

m/z 675 [M-H]^-^
KNOWN
290.03
100
90
80
70
288.05
60
Relative Abundance
50
40
30
20
454.12
10
306.07
471.13
615.19
272.00
384.12
308.05
451.09
573.18
219.97
410.12
203.98
633.19
246.00
332.07
362.06
555.17
514.16
650.92
0
200
250
300
350
400
450
500
550
600
650
m/z
L'Hoest's monkey (Allochrocebus lhoesti)
Beluga whale (Delphinapterus leucas)
Bottlenose dolphin (Tursiops truncates)
Black rhinoceros (Diceros bicornis)
Impala (Aepyceros melampus)
Striped dolphin (Stenella coeruleoalba)

### Slide 35
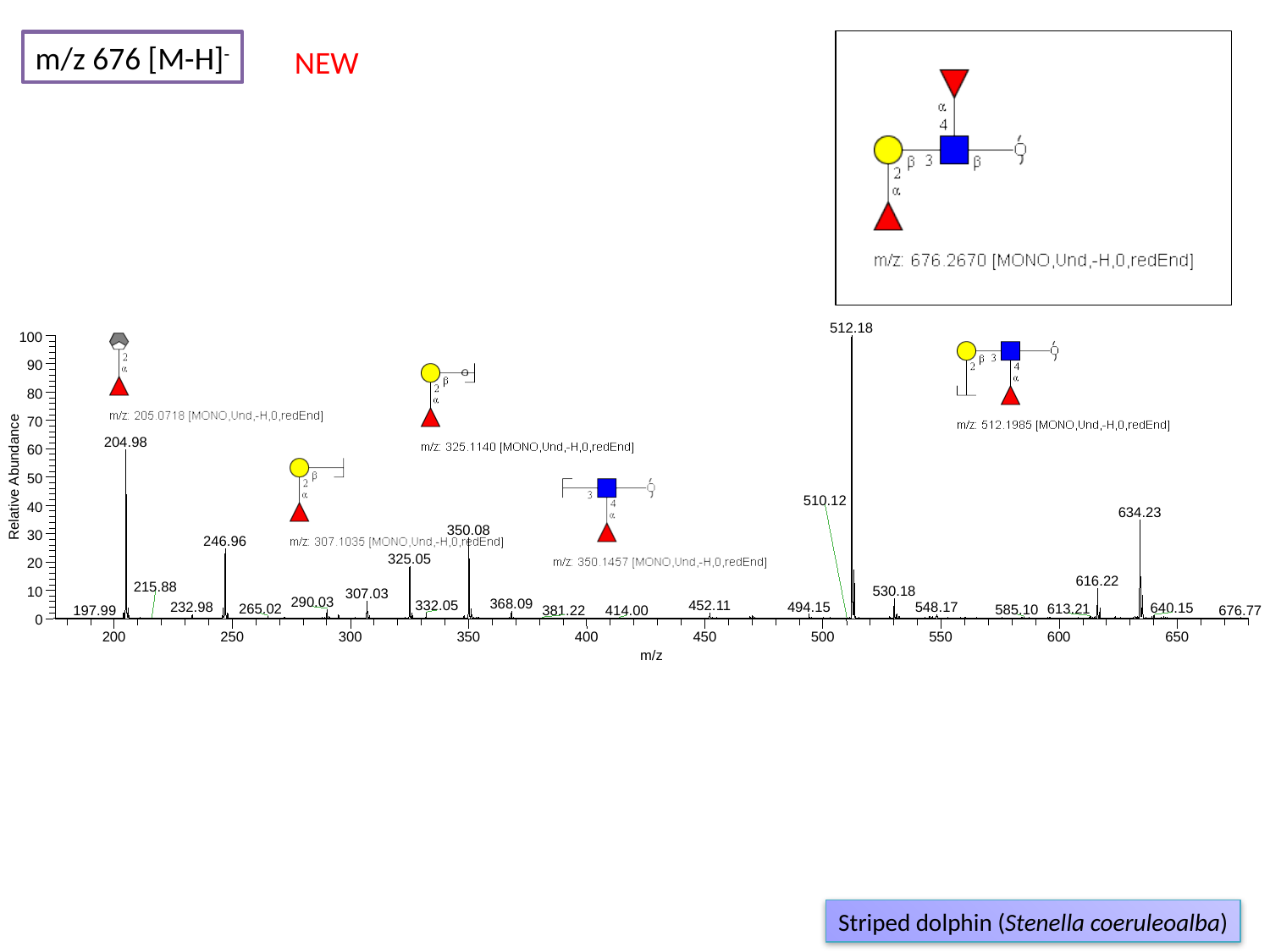

m/z 676 [M-H]^-^
NEW
512.18
100
90
80
70
204.98
60
Relative Abundance
50
510.12
40
634.23
350.08
30
246.96
325.05
20
616.22
215.88
530.18
10
307.03
290.03
368.09
332.05
452.11
494.15
232.98
548.17
640.15
265.02
613.21
585.10
676.77
381.22
197.99
414.00
0
200
250
300
350
400
450
500
550
600
650
m/z
Striped dolphin (Stenella coeruleoalba)

### Slide 36
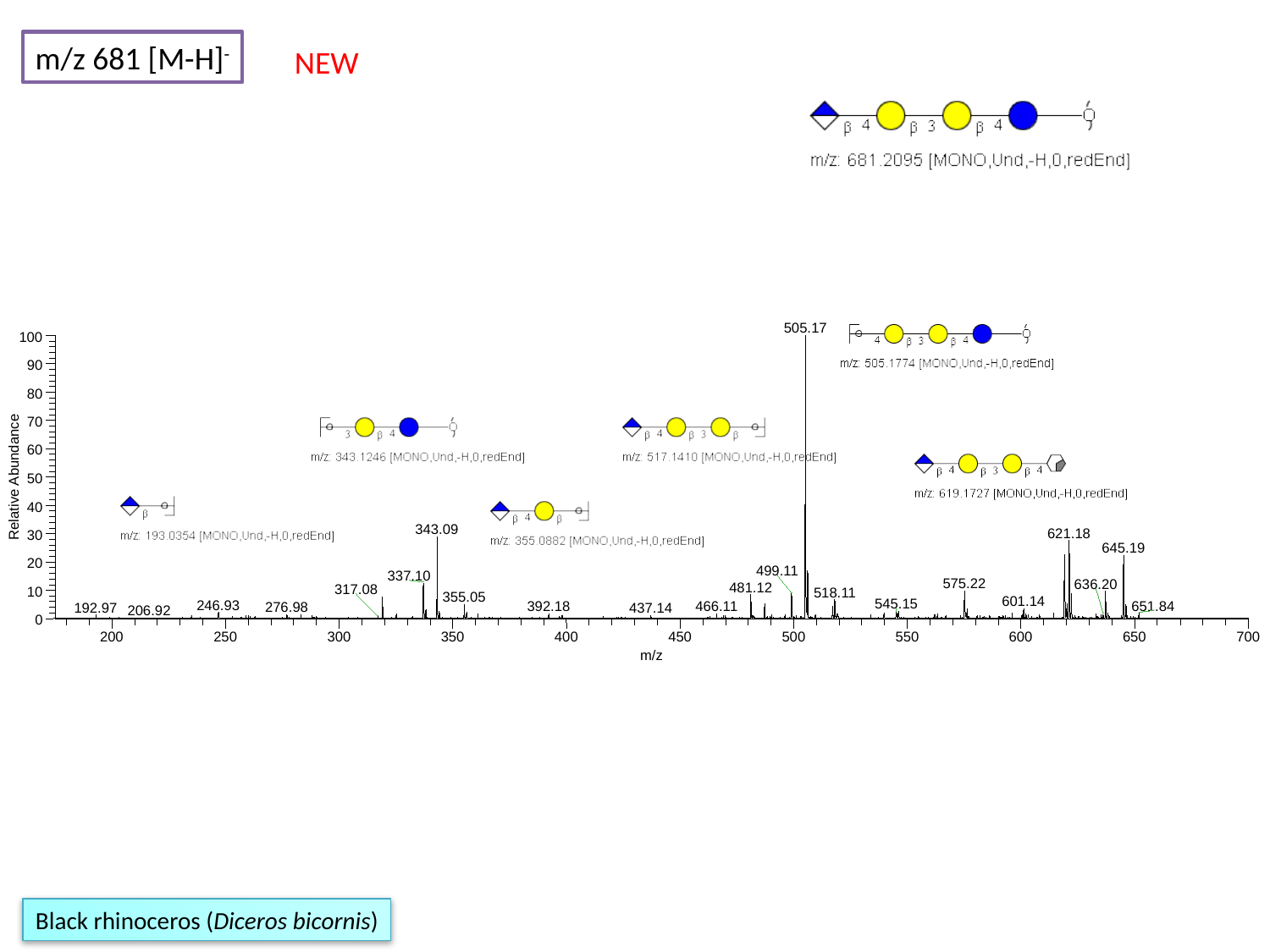

m/z 681 [M-H]^-^
NEW
505.17
100
90
80
70
60
Relative Abundance
50
40
343.09
621.18
30
645.19
20
499.11
337.10
575.22
636.20
481.12
317.08
10
518.11
355.05
601.14
545.15
246.93
651.84
466.11
392.18
276.98
192.97
437.14
206.92
0
200
250
300
350
400
450
500
550
600
650
700
m/z
Black rhinoceros (Diceros bicornis)

### Slide 37
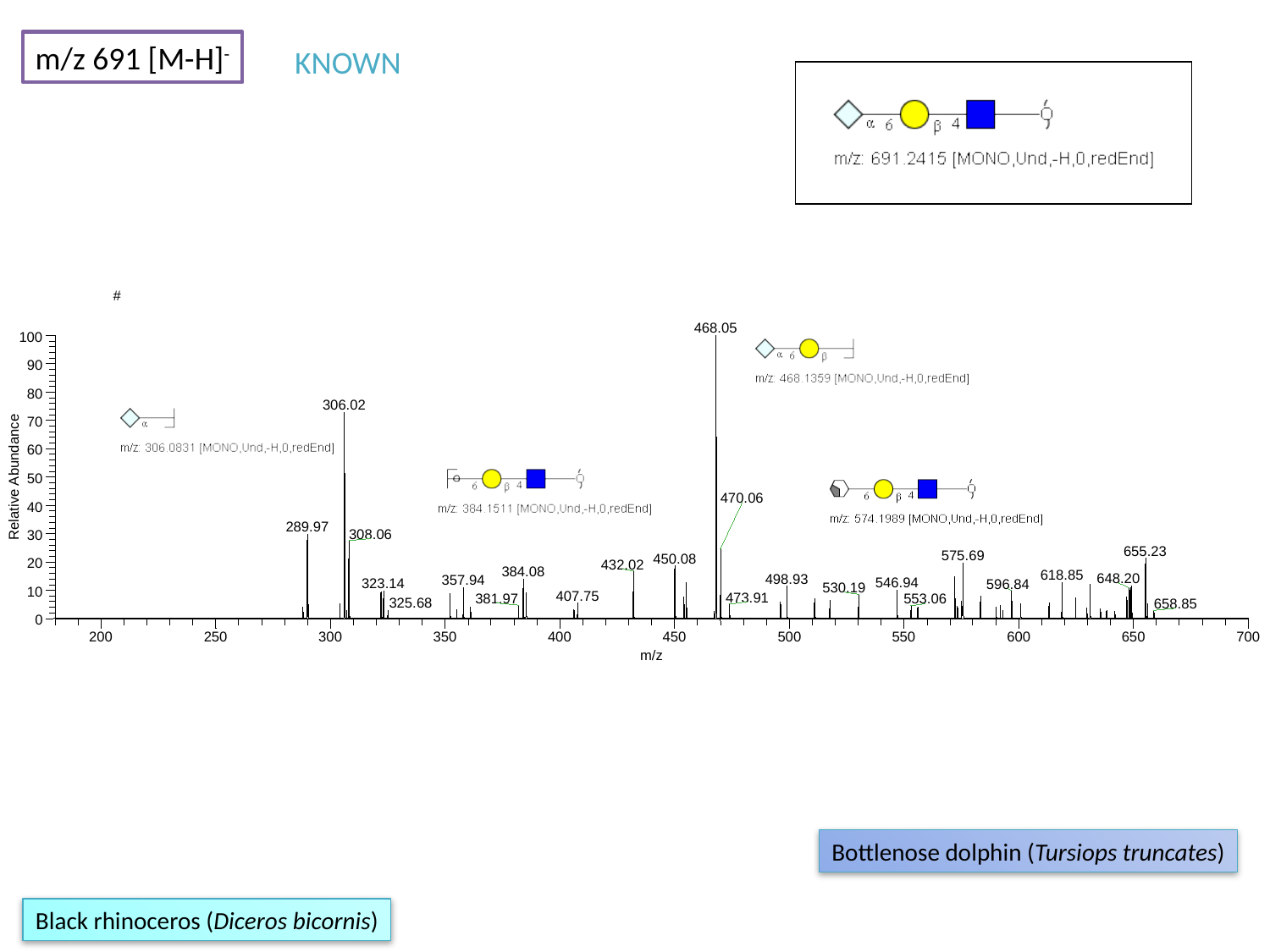

m/z 691 [M-H]^-^
KNOWN
#
468.05
100
90
80
306.02
70
60
Relative Abundance
50
470.06
40
289.97
308.06
30
655.23
575.69
450.08
20
432.02
384.08
618.85
648.20
498.93
357.94
546.94
323.14
596.84
530.19
10
407.75
473.91
381.97
553.06
325.68
658.85
0
200
250
300
350
400
450
500
550
600
650
700
m/z
Bottlenose dolphin (Tursiops truncates)
Black rhinoceros (Diceros bicornis)

### Slide 38
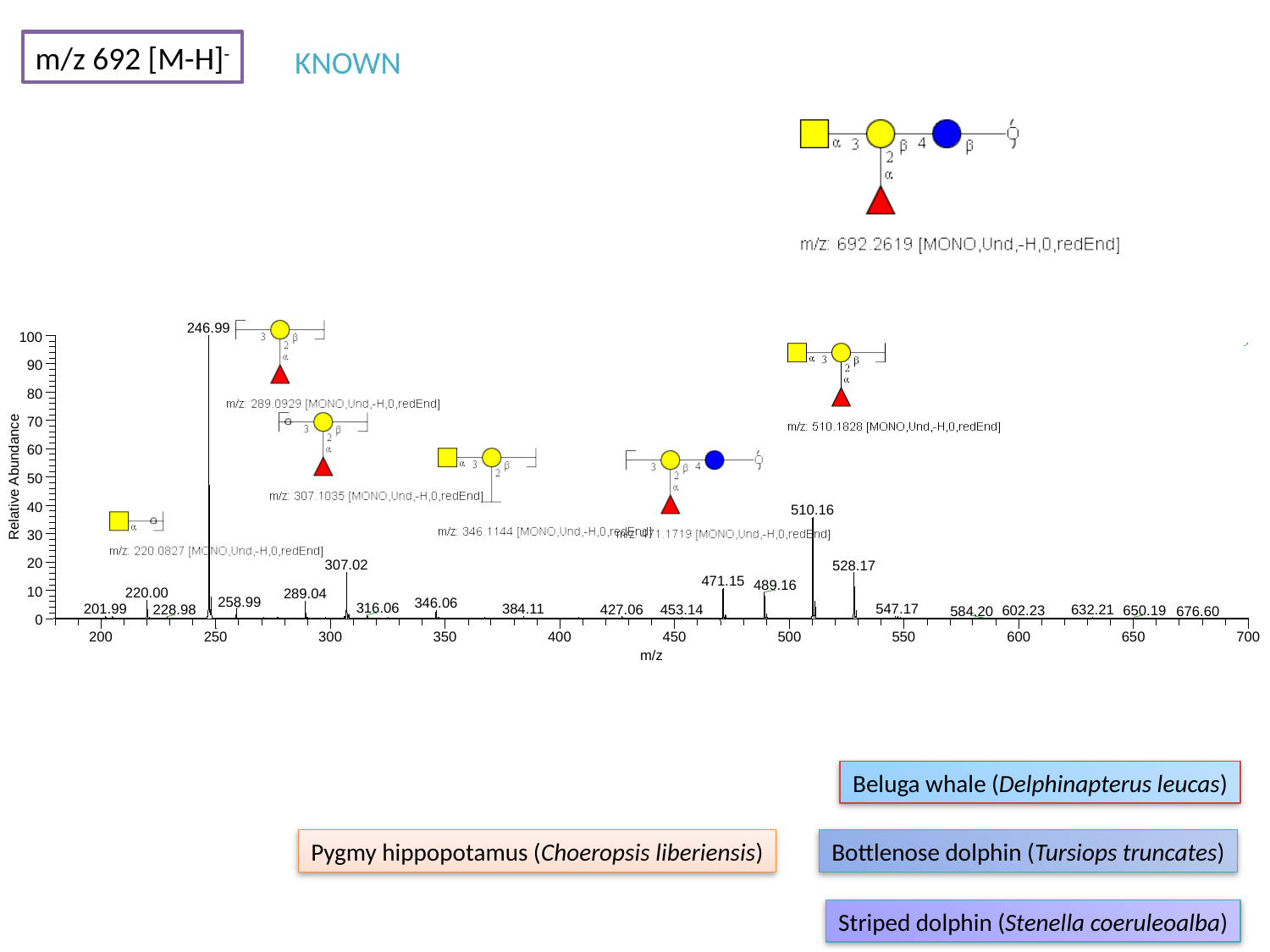

m/z 692 [M-H]^-^
KNOWN
246.99
100
90
80
70
60
Relative Abundance
50
40
510.16
30
20
307.02
528.17
471.15
489.16
10
220.00
289.04
258.99
346.06
316.06
201.99
384.11
547.17
427.06
453.14
632.21
228.98
650.19
602.23
584.20
676.60
0
200
250
300
350
400
450
500
550
600
650
700
m/z
Beluga whale (Delphinapterus leucas)
Pygmy hippopotamus (Choeropsis liberiensis)
Bottlenose dolphin (Tursiops truncates)
Striped dolphin (Stenella coeruleoalba)

### Slide 39
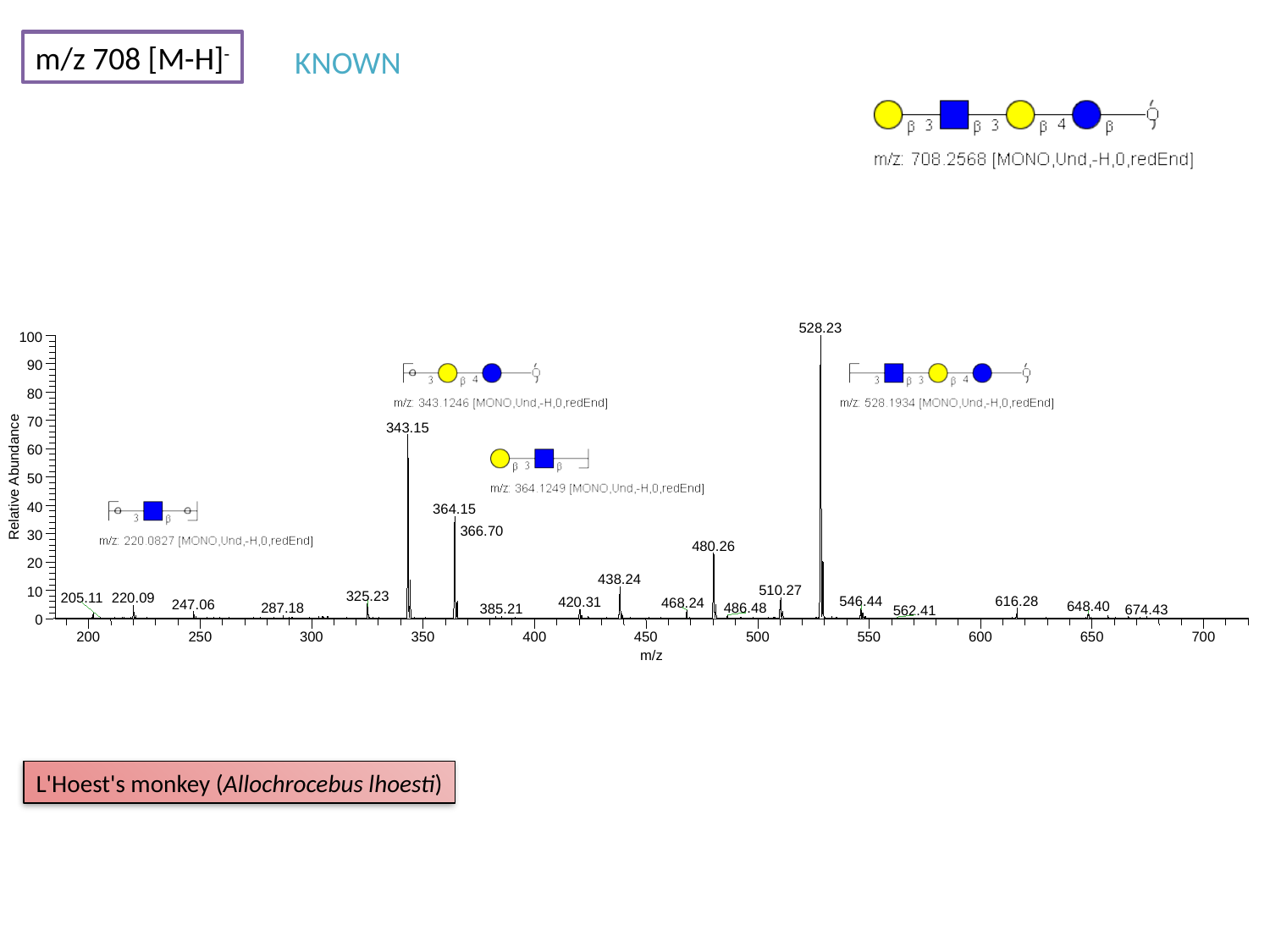

m/z 708 [M-H]^-^
KNOWN
528.23
100
90
80
70
343.15
60
Relative Abundance
50
40
364.15
366.70
30
480.26
20
438.24
510.27
10
325.23
205.11
220.09
616.28
546.44
420.31
468.24
247.06
648.40
287.18
486.48
385.21
674.43
562.41
0
200
250
300
350
400
450
500
550
600
650
700
m/z
L'Hoest's monkey (Allochrocebus lhoesti)

### Slide 40
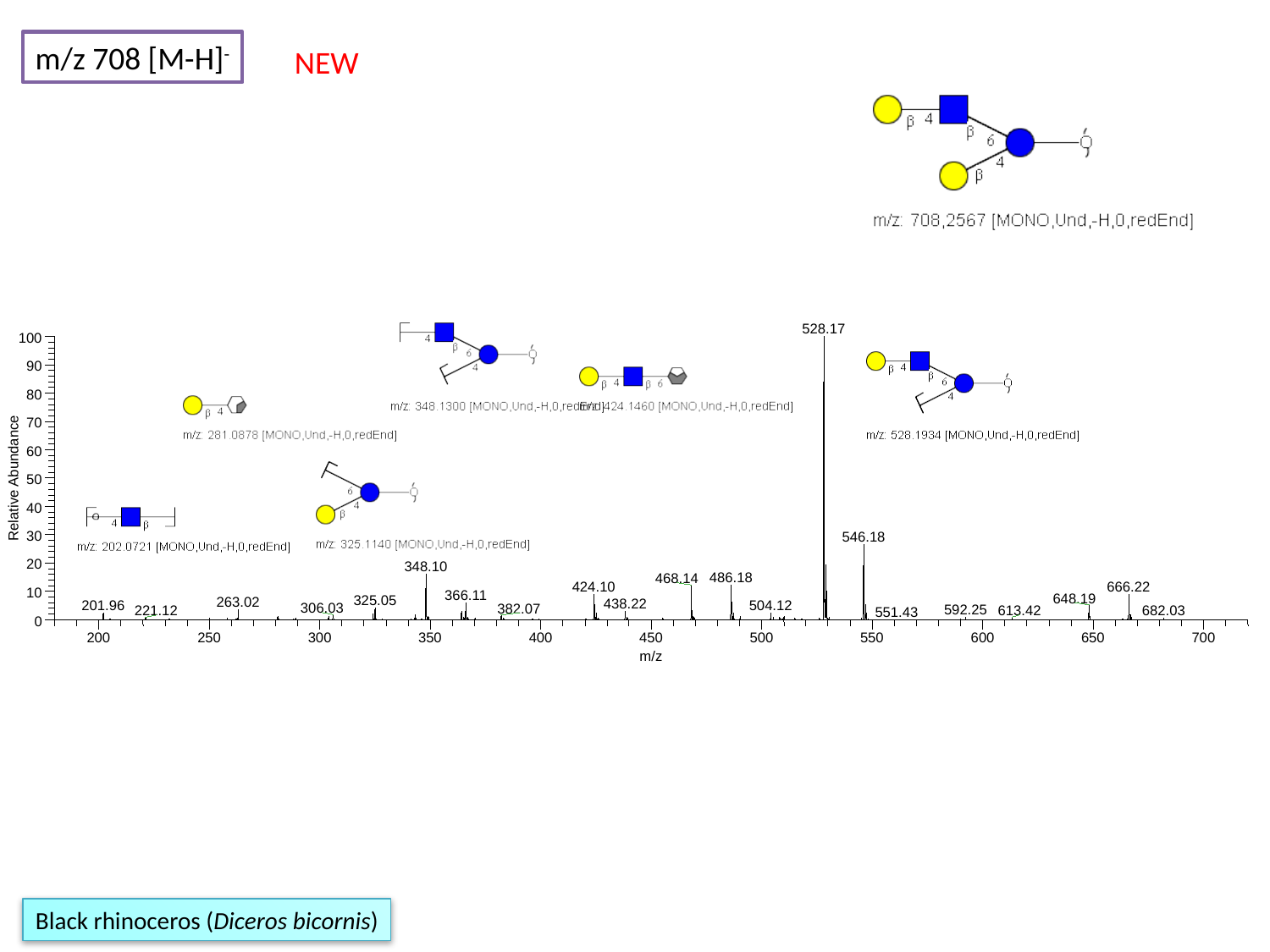

m/z 708 [M-H]^-^
NEW
528.17
100
90
80
70
60
Relative Abundance
50
40
30
546.18
20
348.10
486.18
468.14
424.10
666.22
10
366.11
648.19
325.05
263.02
438.22
201.96
504.12
306.03
382.07
592.25
613.42
221.12
682.03
551.43
0
200
250
300
350
400
450
500
550
600
650
700
m/z
Black rhinoceros (Diceros bicornis)

### Slide 41
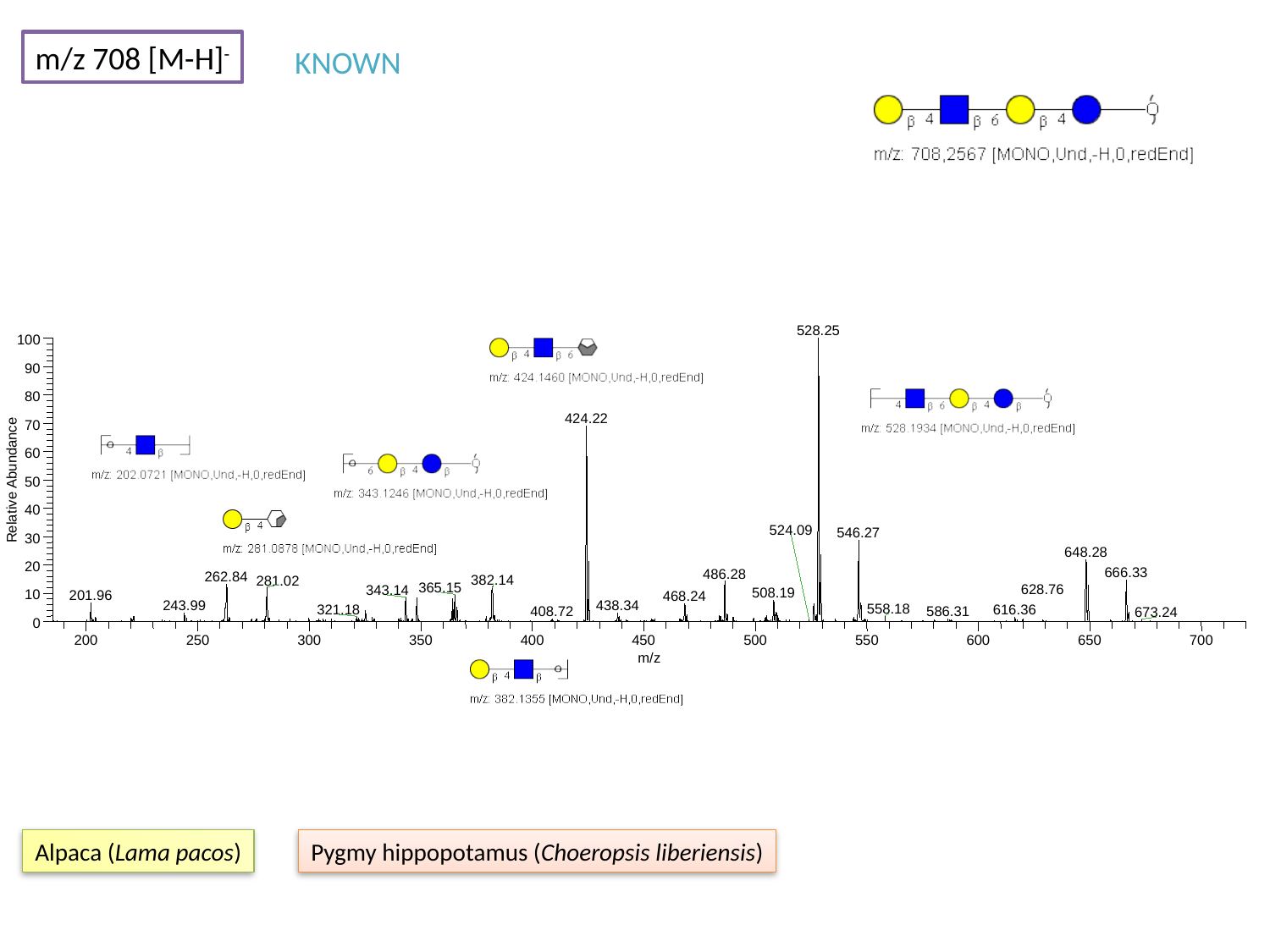

m/z 708 [M-H]^-^
KNOWN
528.25
100
90
80
424.22
70
60
Relative Abundance
50
40
524.09
546.27
30
648.28
20
666.33
486.28
262.84
382.14
281.02
365.15
628.76
343.14
508.19
10
201.96
468.24
243.99
438.34
558.18
321.18
616.36
586.31
408.72
673.24
0
200
250
300
350
400
450
500
550
600
650
700
m/z
Alpaca (Lama pacos)
Pygmy hippopotamus (Choeropsis liberiensis)

### Slide 42
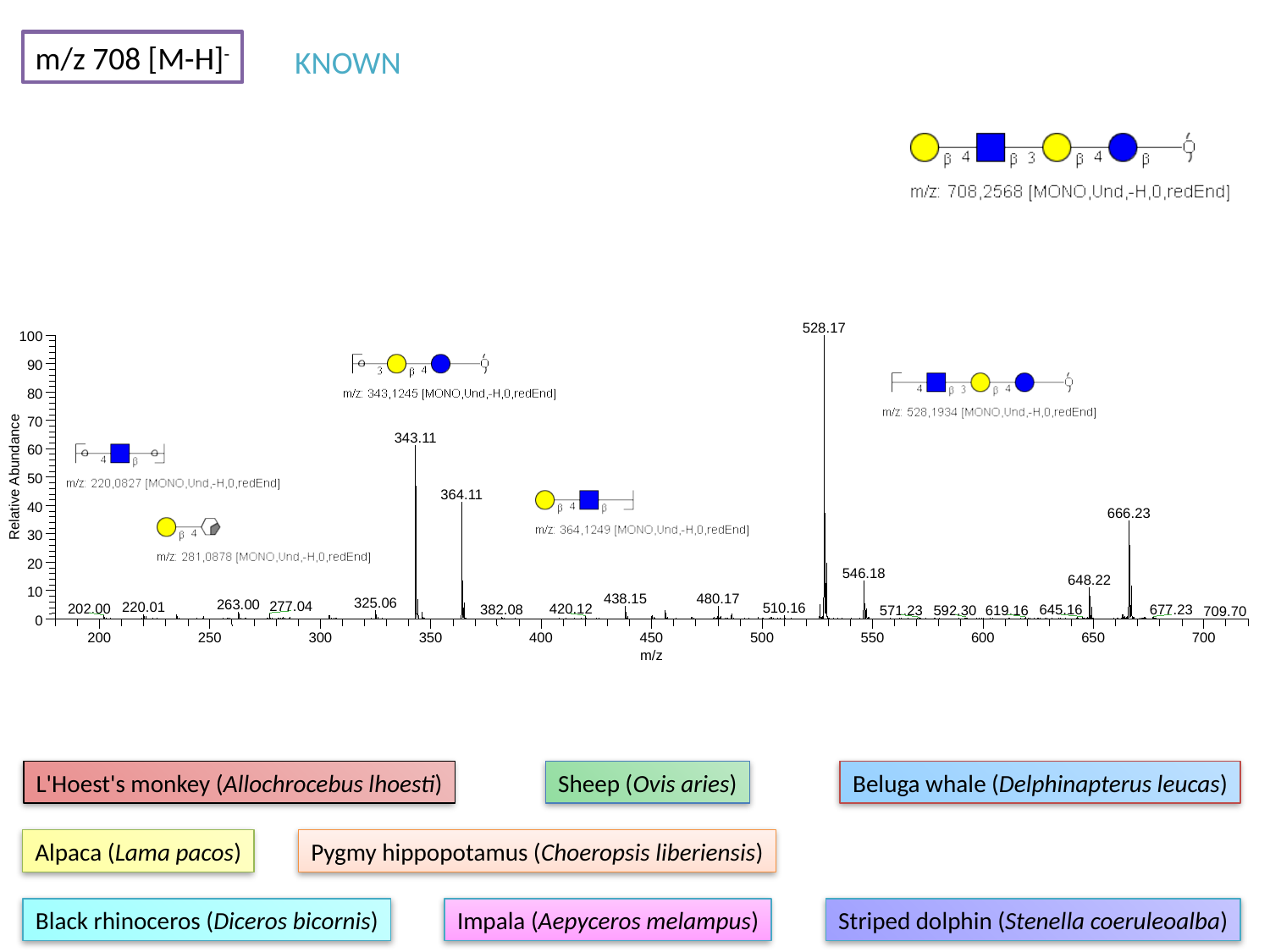

m/z 708 [M-H]^-^
KNOWN
528.17
100
90
80
70
343.11
60
Relative Abundance
50
364.11
40
666.23
30
20
546.18
648.22
10
480.17
438.15
325.06
263.00
277.04
220.01
510.16
202.00
420.12
382.08
645.16
677.23
619.16
571.23
592.30
709.70
0
200
250
300
350
400
450
500
550
600
650
700
m/z
L'Hoest's monkey (Allochrocebus lhoesti)
Sheep (Ovis aries)
Beluga whale (Delphinapterus leucas)
Alpaca (Lama pacos)
Pygmy hippopotamus (Choeropsis liberiensis)
Black rhinoceros (Diceros bicornis)
Impala (Aepyceros melampus)
Striped dolphin (Stenella coeruleoalba)

### Slide 43
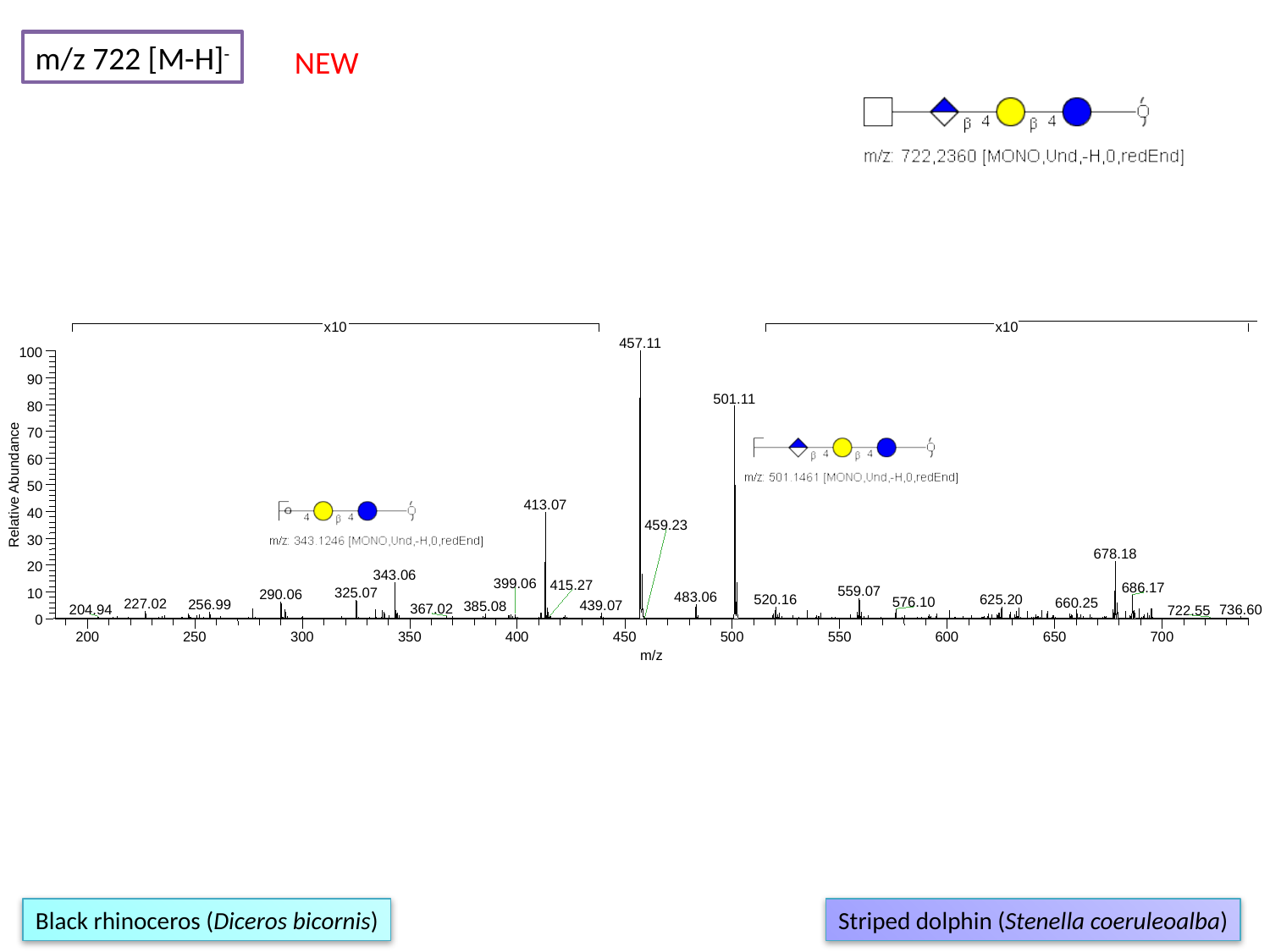

m/z 722 [M-H]^-^
NEW
x10
x10
457.11
100
90
501.11
80
70
60
Relative Abundance
50
413.07
40
459.23
30
678.18
20
343.06
399.06
415.27
686.17
559.07
325.07
10
290.06
483.06
625.20
520.16
576.10
660.25
227.02
256.99
439.07
385.08
367.02
736.60
204.94
722.55
0
200
250
300
350
400
450
500
550
600
650
700
m/z
Black rhinoceros (Diceros bicornis)
Striped dolphin (Stenella coeruleoalba)

### Slide 44
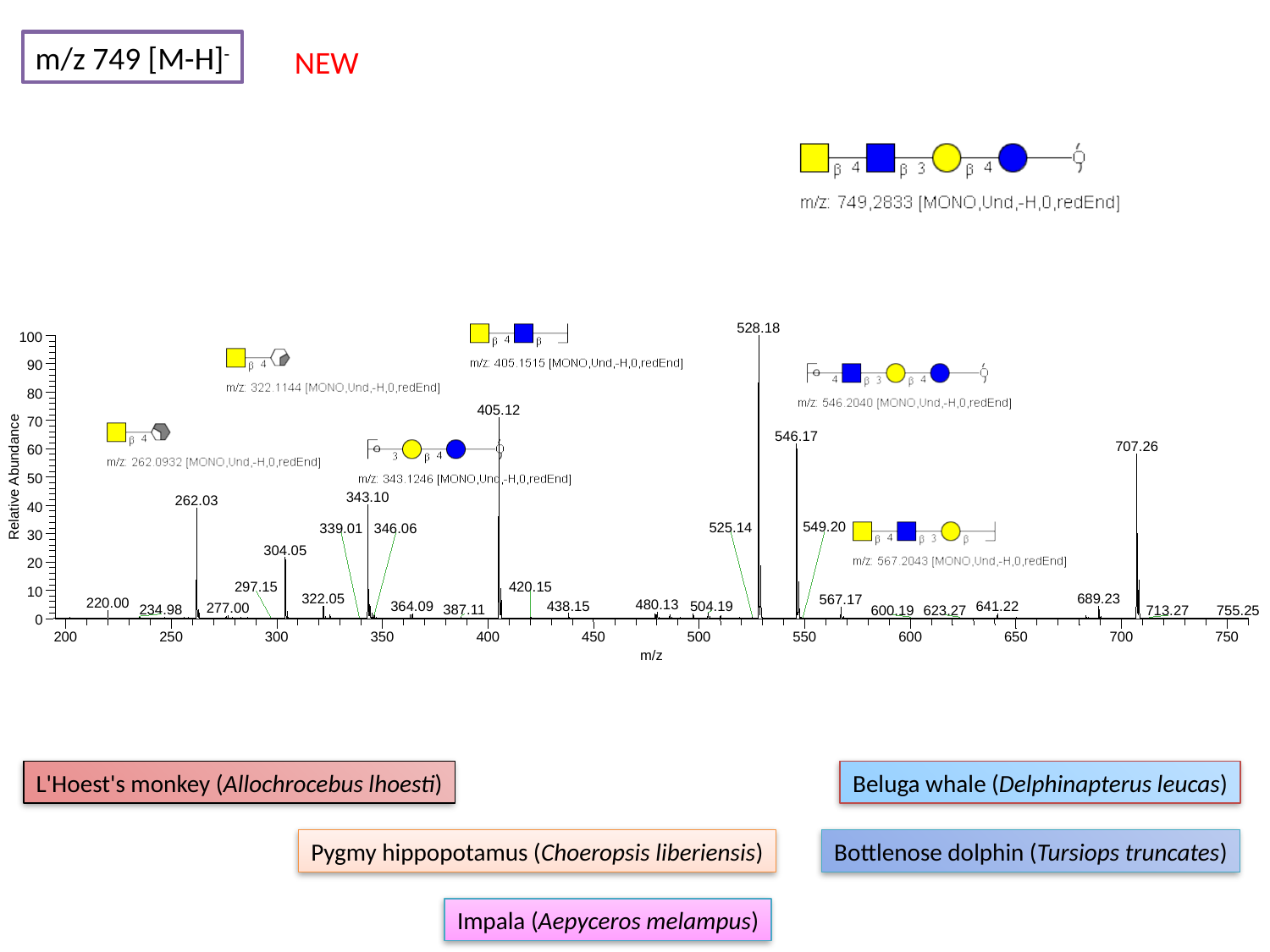

m/z 749 [M-H]^-^
NEW
528.18
100
90
80
405.12
70
546.17
707.26
60
Relative Abundance
50
343.10
262.03
40
549.20
525.14
339.01
346.06
30
304.05
20
297.15
420.15
10
322.05
689.23
567.17
220.00
480.13
438.15
504.19
364.09
641.22
277.00
234.98
387.11
600.19
623.27
713.27
755.25
0
200
250
300
350
400
450
500
550
600
650
700
750
m/z
L'Hoest's monkey (Allochrocebus lhoesti)
Beluga whale (Delphinapterus leucas)
Pygmy hippopotamus (Choeropsis liberiensis)
Bottlenose dolphin (Tursiops truncates)
Impala (Aepyceros melampus)

### Slide 45
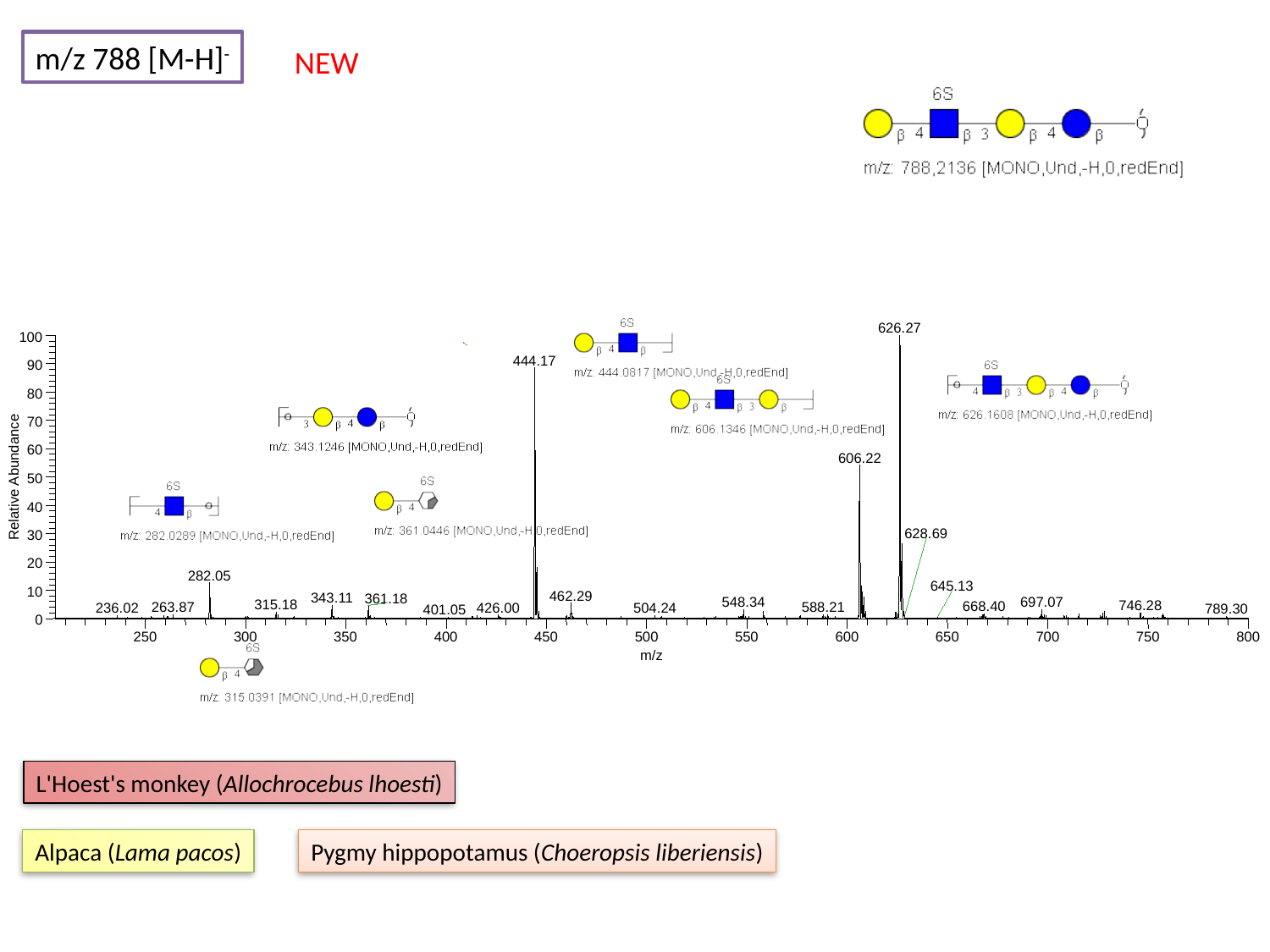

m/z 788 [M-H]^-^
NEW
626.27
100
444.17
90
80
70
60
606.22
Relative Abundance
50
40
628.69
30
20
282.05
645.13
10
462.29
343.11
361.18
548.34
697.07
315.18
746.28
668.40
263.87
588.21
426.00
236.02
504.24
789.30
401.05
0
250
300
350
400
450
500
550
600
650
700
750
800
m/z
L'Hoest's monkey (Allochrocebus lhoesti)
Alpaca (Lama pacos)
Pygmy hippopotamus (Choeropsis liberiensis)

### Slide 46
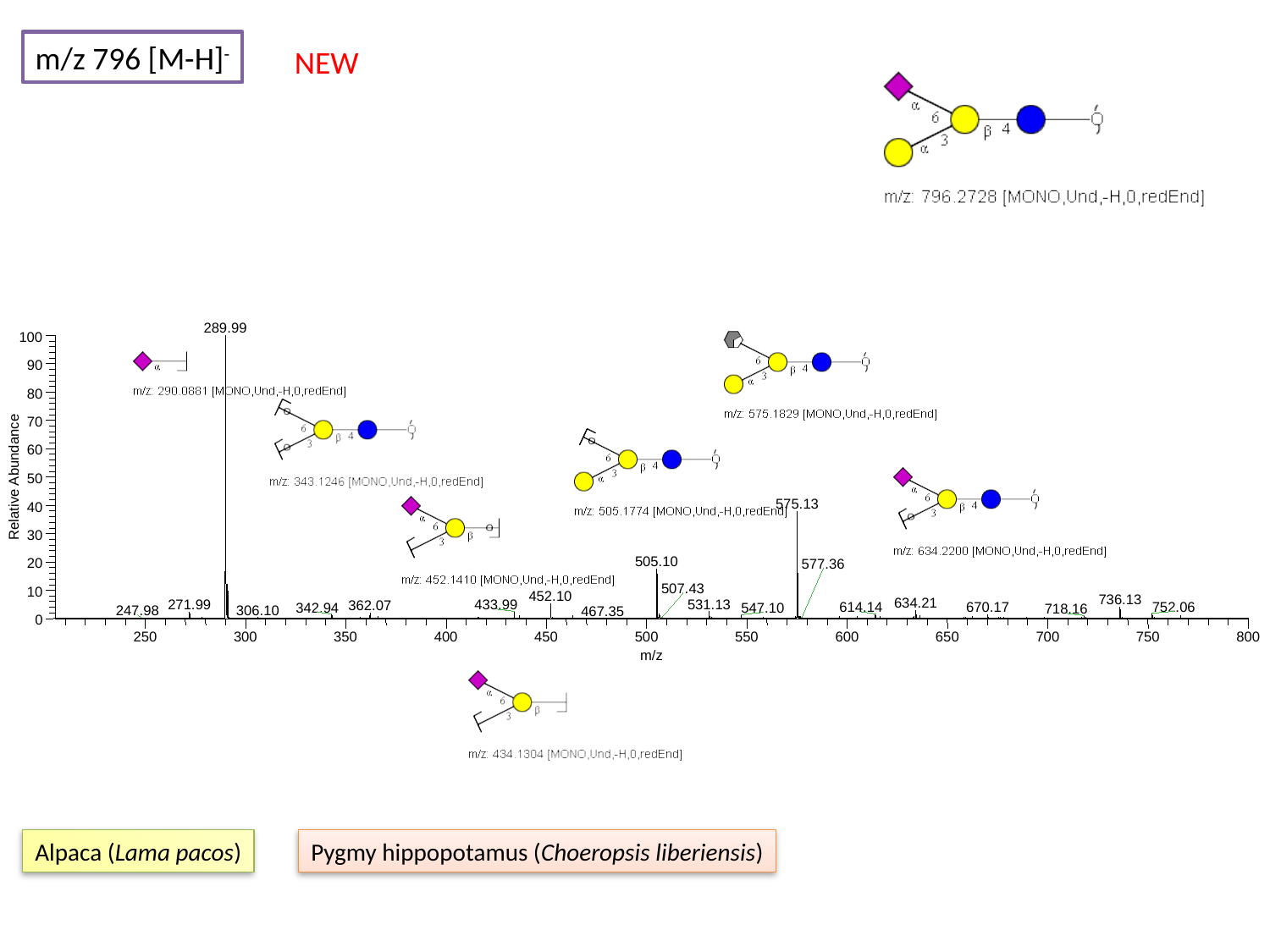

m/z 796 [M-H]^-^
NEW
289.99
100
90
80
70
60
Relative Abundance
50
575.13
40
30
505.10
20
577.36
507.43
10
452.10
736.13
634.21
531.13
271.99
433.99
362.07
752.06
614.14
670.17
342.94
547.10
718.16
306.10
247.98
467.35
0
250
300
350
400
450
500
550
600
650
700
750
800
m/z
Alpaca (Lama pacos)
Pygmy hippopotamus (Choeropsis liberiensis)

### Slide 47
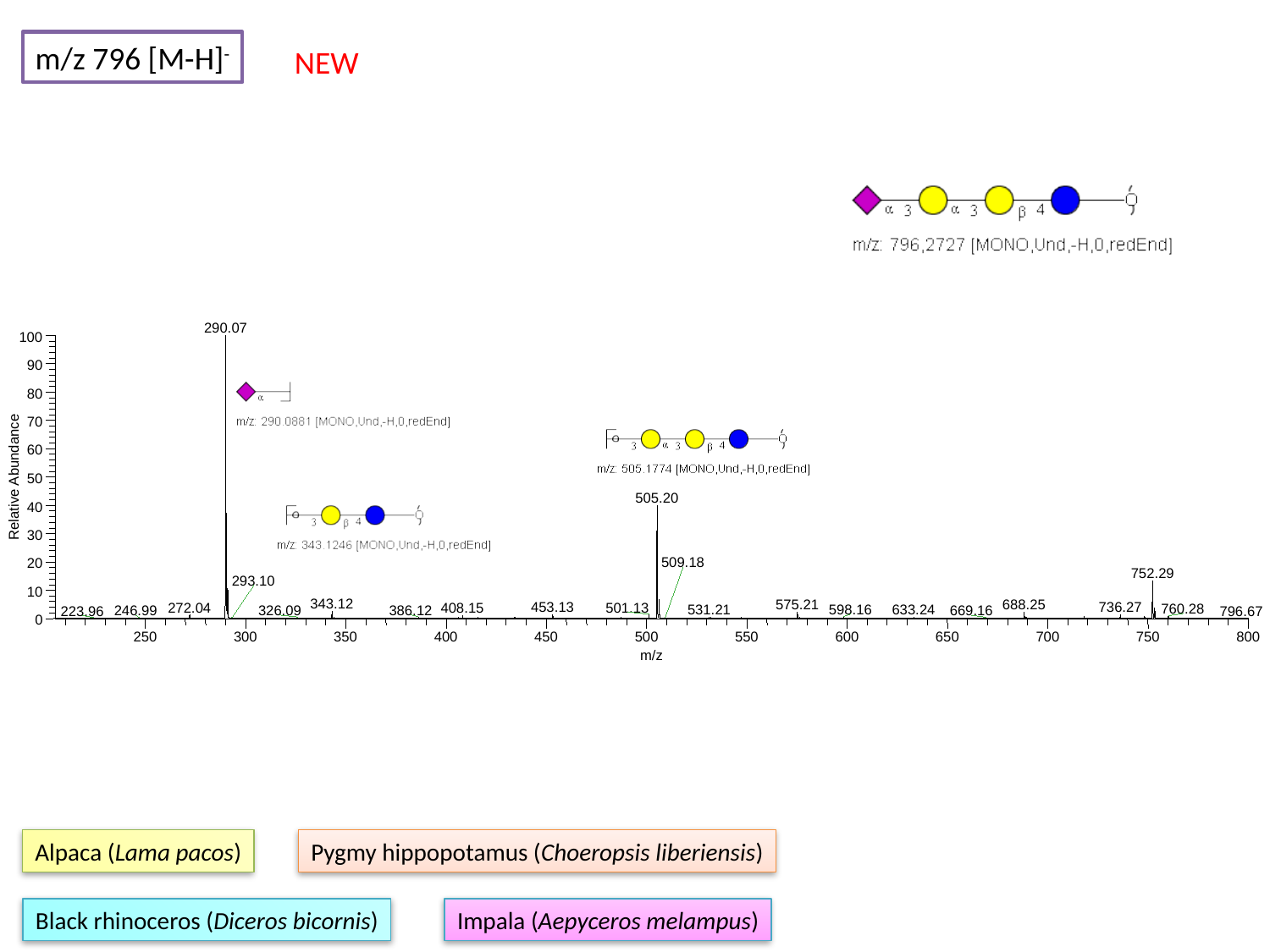

m/z 796 [M-H]^-^
NEW
290.07
100
90
80
70
60
Relative Abundance
50
505.20
40
30
509.18
20
752.29
293.10
10
343.12
575.21
688.25
453.13
736.27
272.04
501.13
408.15
760.28
531.21
598.16
633.24
386.12
326.09
246.99
669.16
223.96
796.67
0
250
300
350
400
450
500
550
600
650
700
750
800
m/z
Alpaca (Lama pacos)
Pygmy hippopotamus (Choeropsis liberiensis)
Black rhinoceros (Diceros bicornis)
Impala (Aepyceros melampus)

### Slide 48
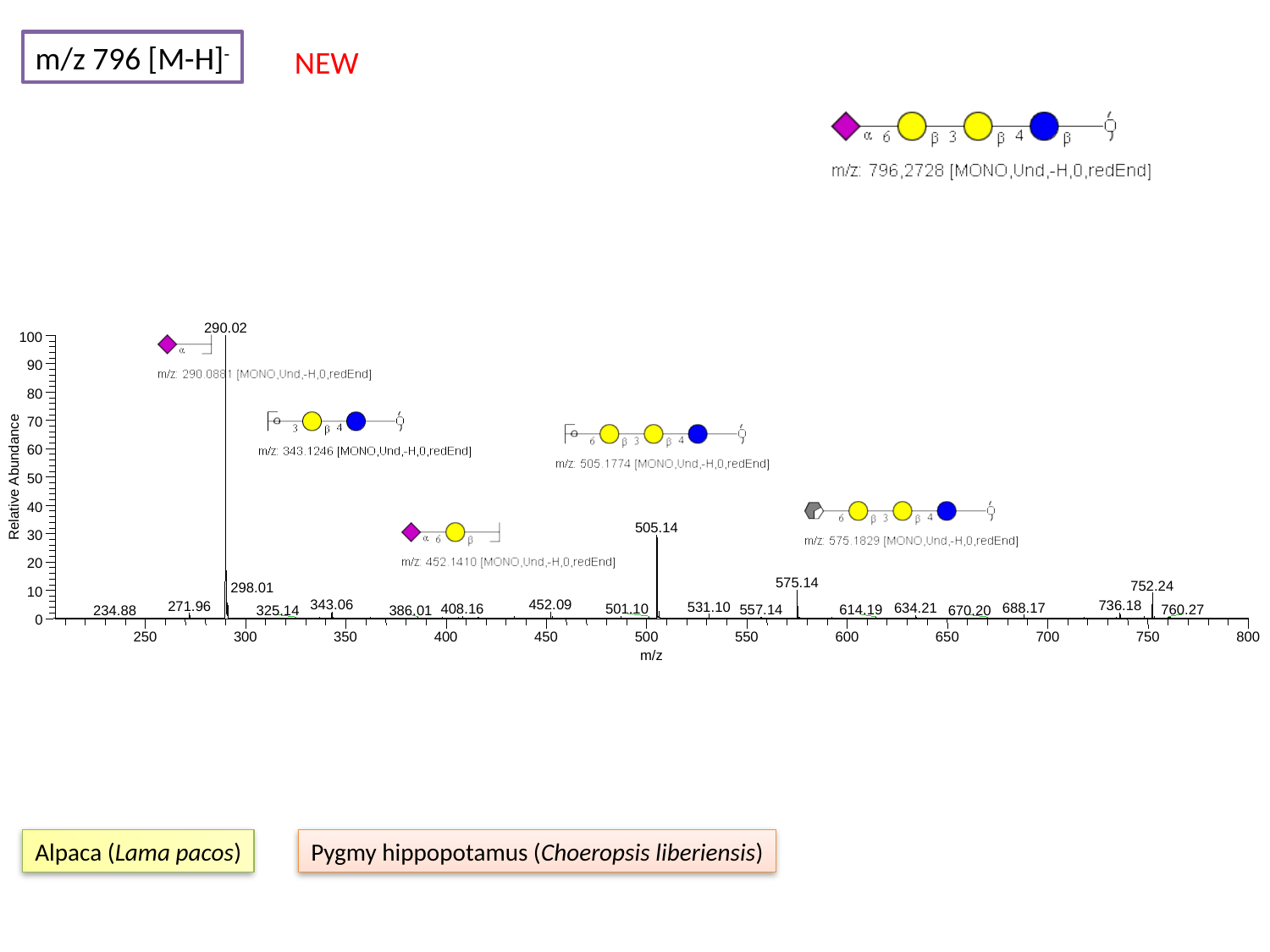

m/z 796 [M-H]^-^
NEW
290.02
100
90
80
70
60
Relative Abundance
50
40
505.14
30
20
575.14
752.24
298.01
10
343.06
452.09
736.18
271.96
531.10
688.17
634.21
408.16
501.10
760.27
557.14
614.19
386.01
325.14
670.20
234.88
0
250
300
350
400
450
500
550
600
650
700
750
800
m/z
Alpaca (Lama pacos)
Pygmy hippopotamus (Choeropsis liberiensis)

### Slide 49
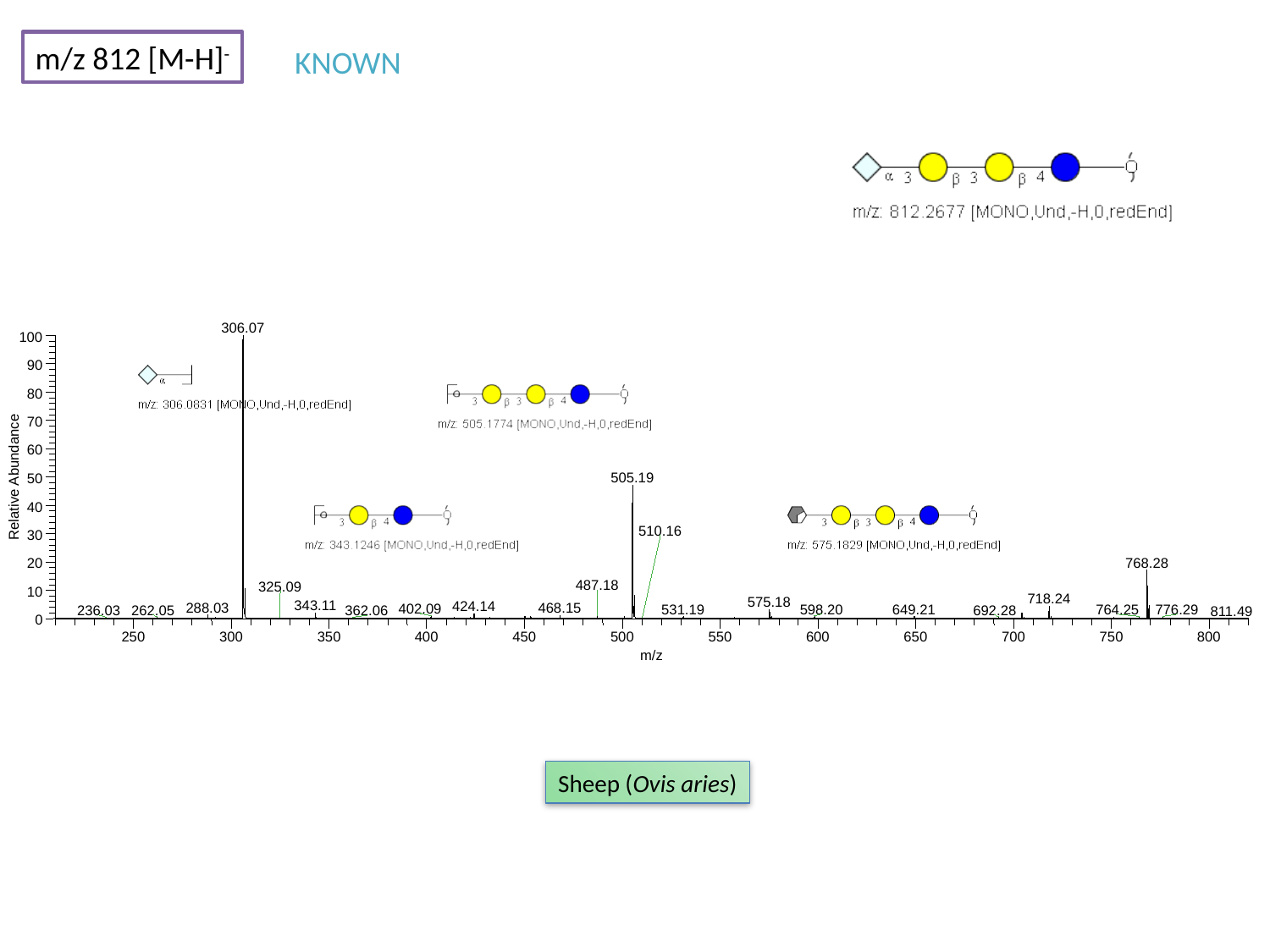

m/z 812 [M-H]^-^
KNOWN
306.07
100
90
80
70
60
Relative Abundance
505.19
50
40
510.16
30
768.28
20
487.18
325.09
10
718.24
575.18
343.11
424.14
288.03
468.15
402.09
531.19
598.20
649.21
764.25
776.29
262.05
692.28
236.03
362.06
811.49
0
250
300
350
400
450
500
550
600
650
700
750
800
m/z
Sheep (Ovis aries)

### Slide 50
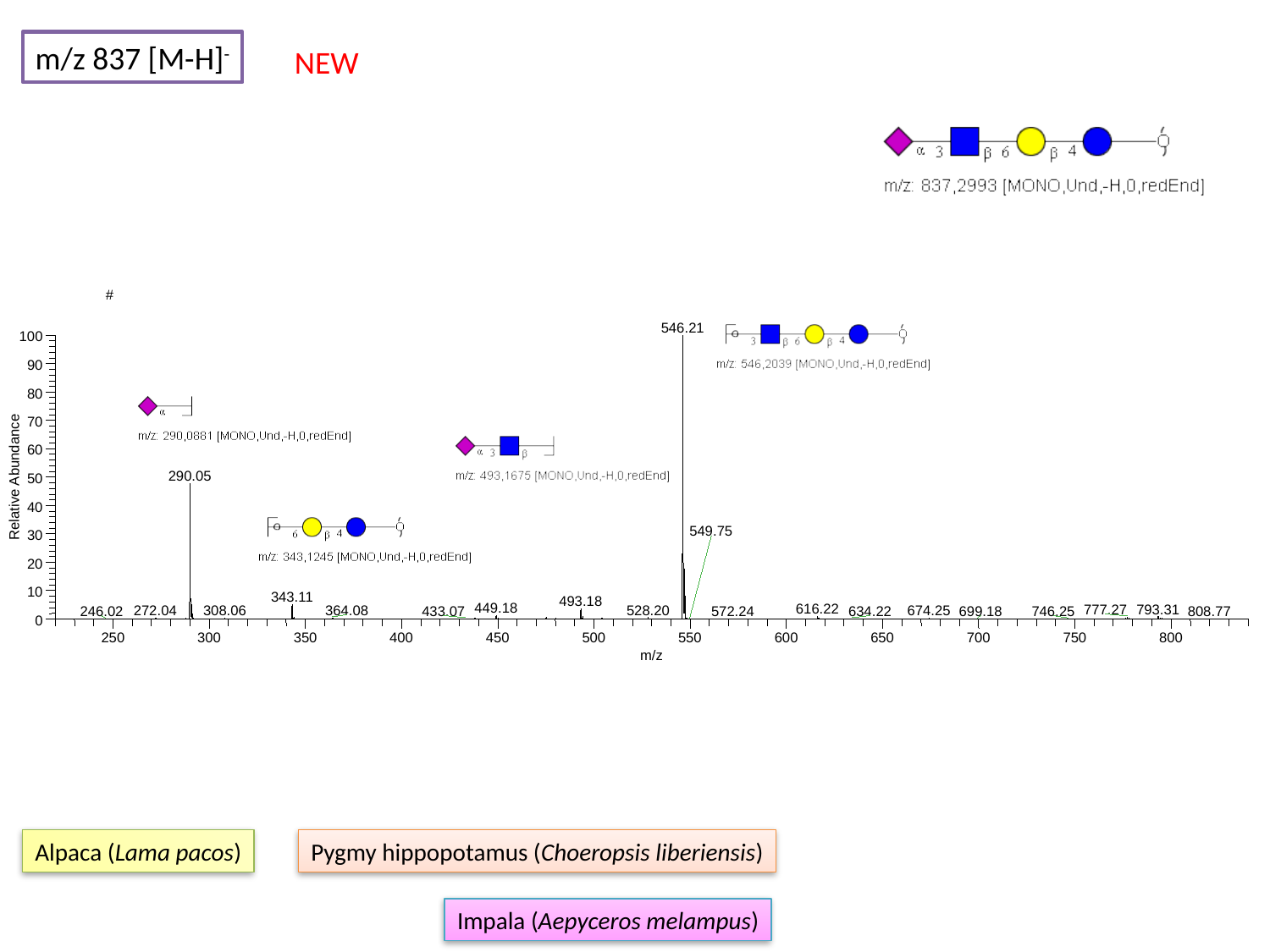

m/z 837 [M-H]^-^
NEW
#
546.21
100
90
80
70
60
290.05
Relative Abundance
50
40
549.75
30
20
10
343.11
493.18
449.18
616.22
793.31
777.27
528.20
272.04
364.08
308.06
674.25
433.07
634.22
246.02
572.24
699.18
746.25
808.77
0
250
300
350
400
450
500
550
600
650
700
750
800
m/z
Alpaca (Lama pacos)
Pygmy hippopotamus (Choeropsis liberiensis)
Impala (Aepyceros melampus)

### Slide 51
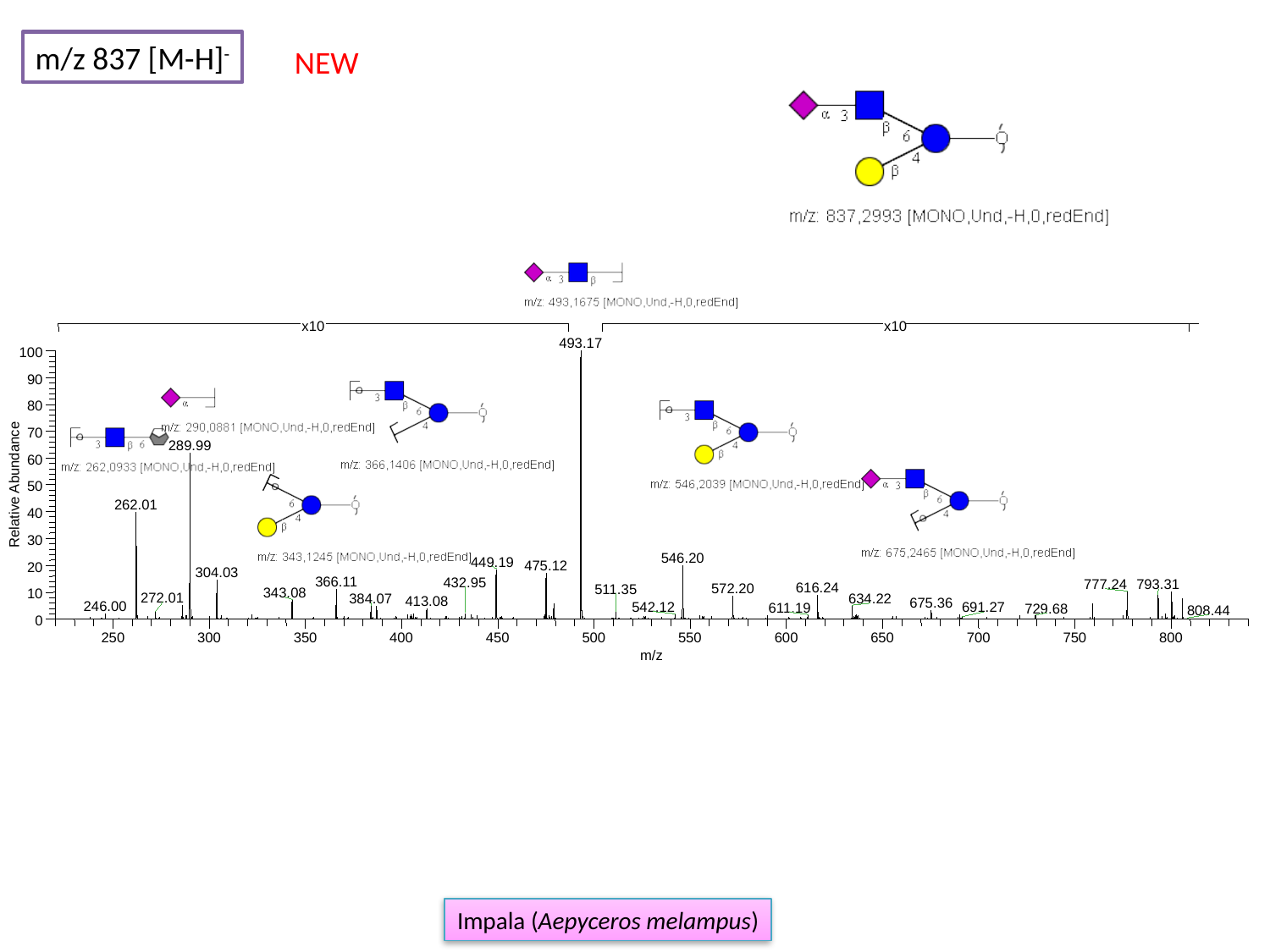

m/z 837 [M-H]^-^
NEW
493.17
100
90
80
70
289.99
60
Relative Abundance
50
262.01
40
30
546.20
449.19
475.12
20
304.03
366.11
432.95
777.24
793.31
616.24
572.20
511.35
343.08
10
272.01
384.07
634.22
413.08
675.36
246.00
542.12
691.27
611.19
729.68
808.44
0
250
300
350
400
450
500
550
600
650
700
750
800
m/z
x10
x10
Impala (Aepyceros melampus)

### Slide 52
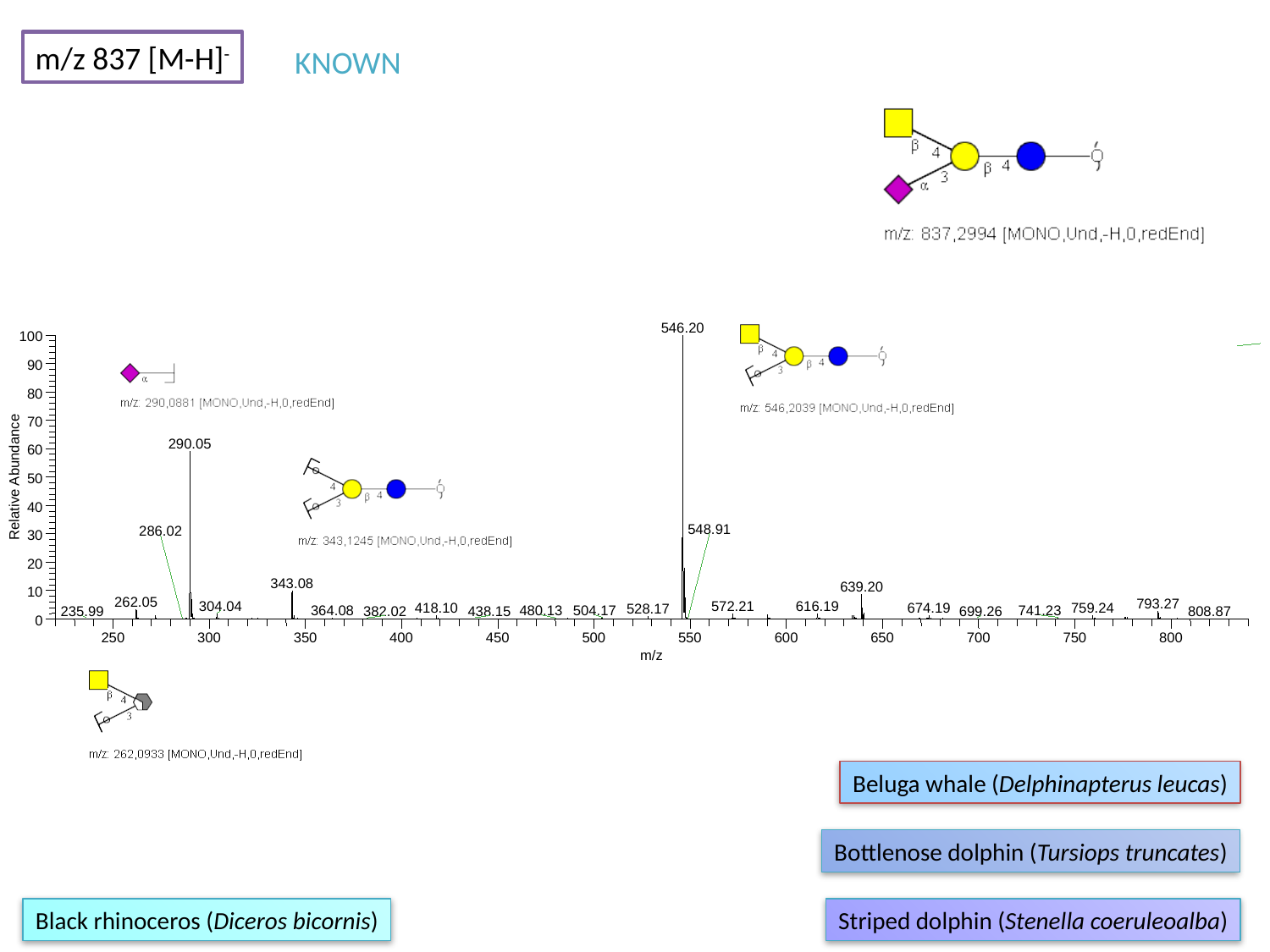

m/z 837 [M-H]^-^
KNOWN
546.20
100
90
80
70
290.05
60
Relative Abundance
50
40
548.91
286.02
30
20
343.08
639.20
10
262.05
793.27
616.19
304.04
572.21
418.10
674.19
759.24
528.17
480.13
504.17
364.08
741.23
235.99
382.02
438.15
699.26
808.87
0
250
300
350
400
450
500
550
600
650
700
750
800
m/z
Beluga whale (Delphinapterus leucas)
Bottlenose dolphin (Tursiops truncates)
Black rhinoceros (Diceros bicornis)
Striped dolphin (Stenella coeruleoalba)

### Slide 53
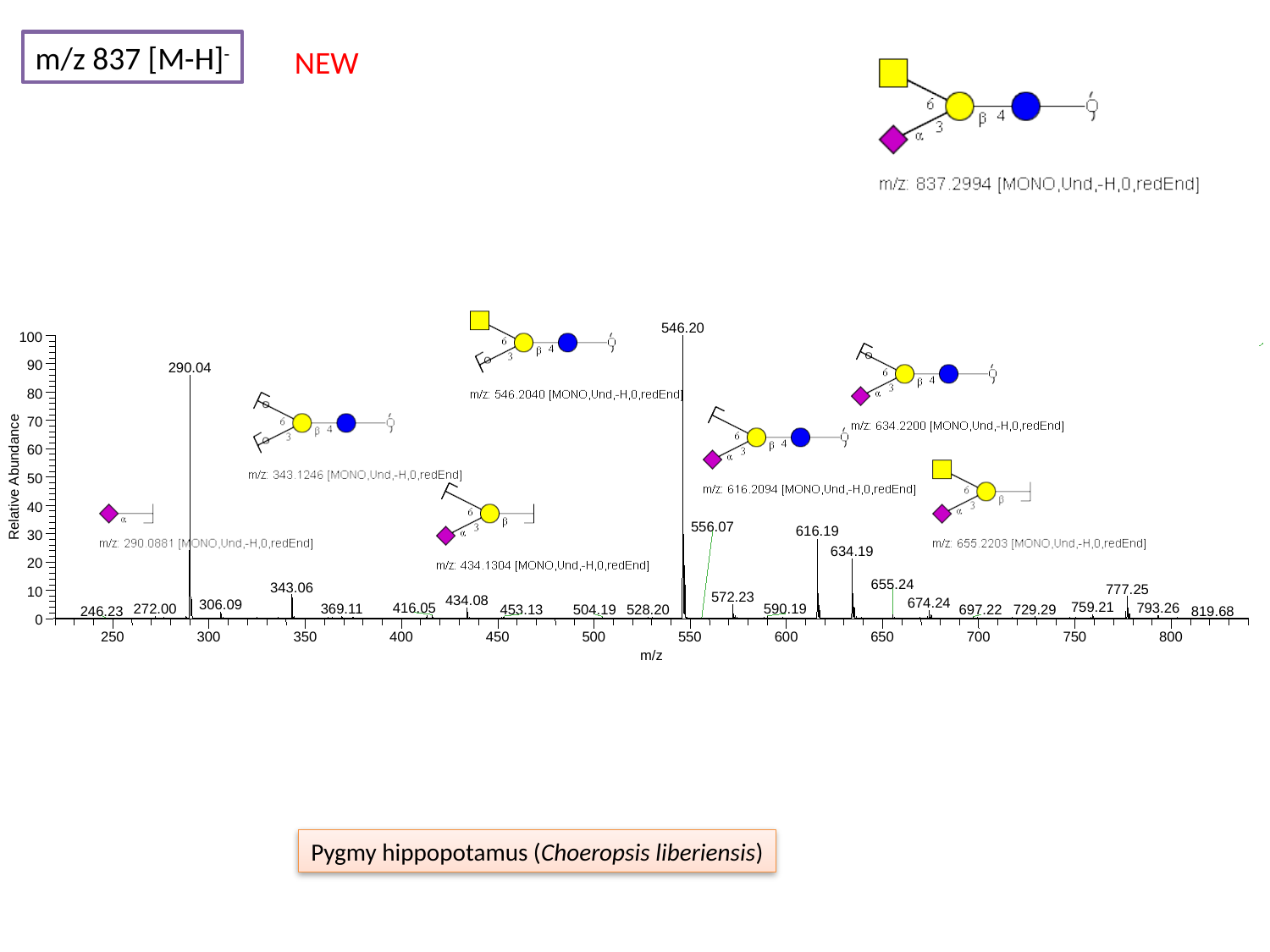

m/z 837 [M-H]^-^
NEW
546.20
100
90
290.04
80
70
60
Relative Abundance
50
40
556.07
616.19
30
634.19
20
655.24
343.06
777.25
10
572.23
434.08
674.24
306.09
759.21
416.05
793.26
272.00
369.11
590.19
729.29
453.13
528.20
504.19
697.22
246.23
819.68
0
250
300
350
400
450
500
550
600
650
700
750
800
m/z
Pygmy hippopotamus (Choeropsis liberiensis)

### Slide 54
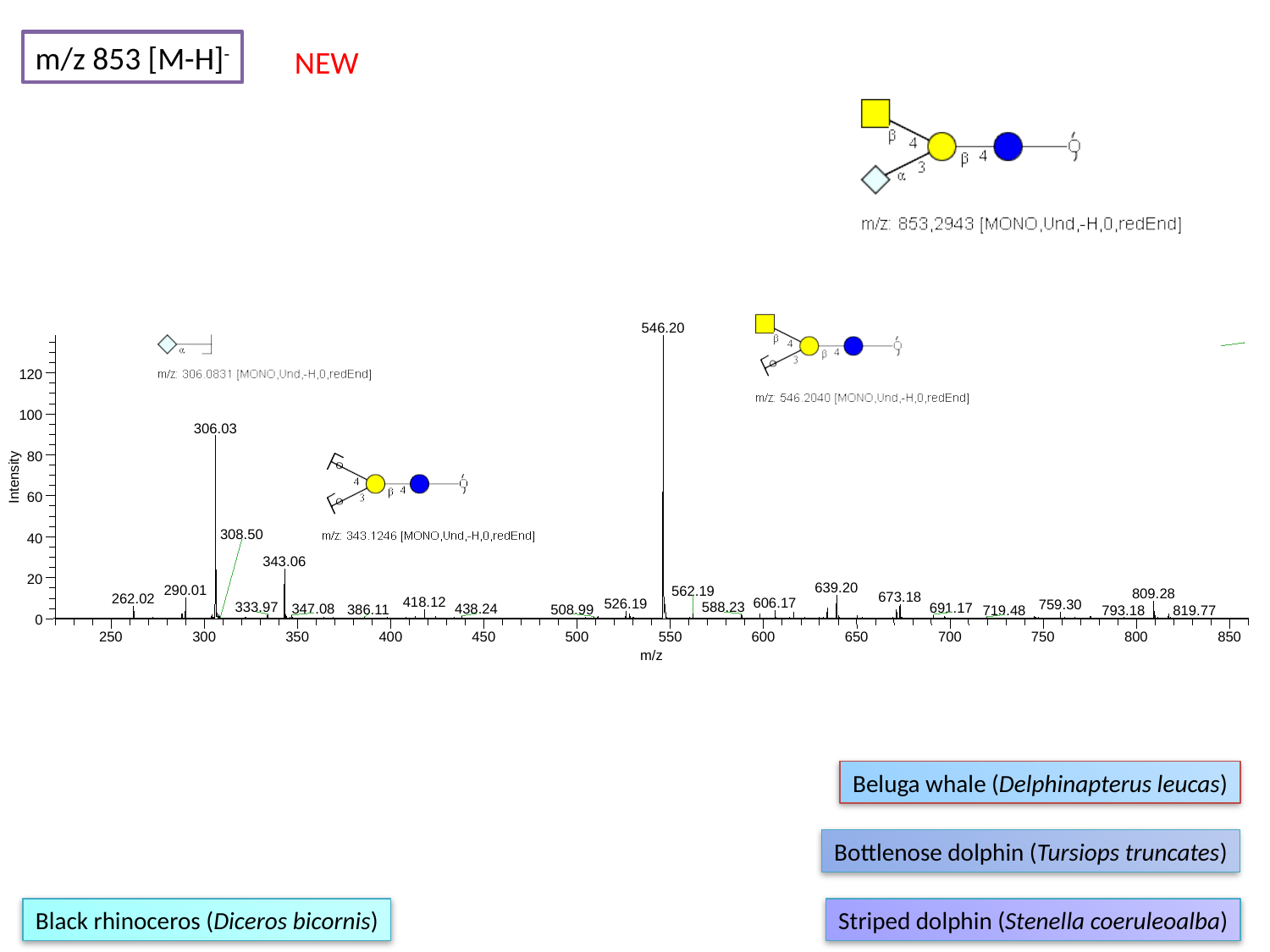

m/z 853 [M-H]^-^
NEW
546.20
120
100
306.03
80
Intensity
60
308.50
40
343.06
20
639.20
290.01
562.19
809.28
673.18
262.02
418.12
606.17
526.19
759.30
588.23
333.97
691.17
347.08
438.24
508.99
386.11
719.48
793.18
819.77
0
250
300
350
400
450
500
550
600
650
700
750
800
850
m/z
Beluga whale (Delphinapterus leucas)
Bottlenose dolphin (Tursiops truncates)
Black rhinoceros (Diceros bicornis)
Striped dolphin (Stenella coeruleoalba)

### Slide 55
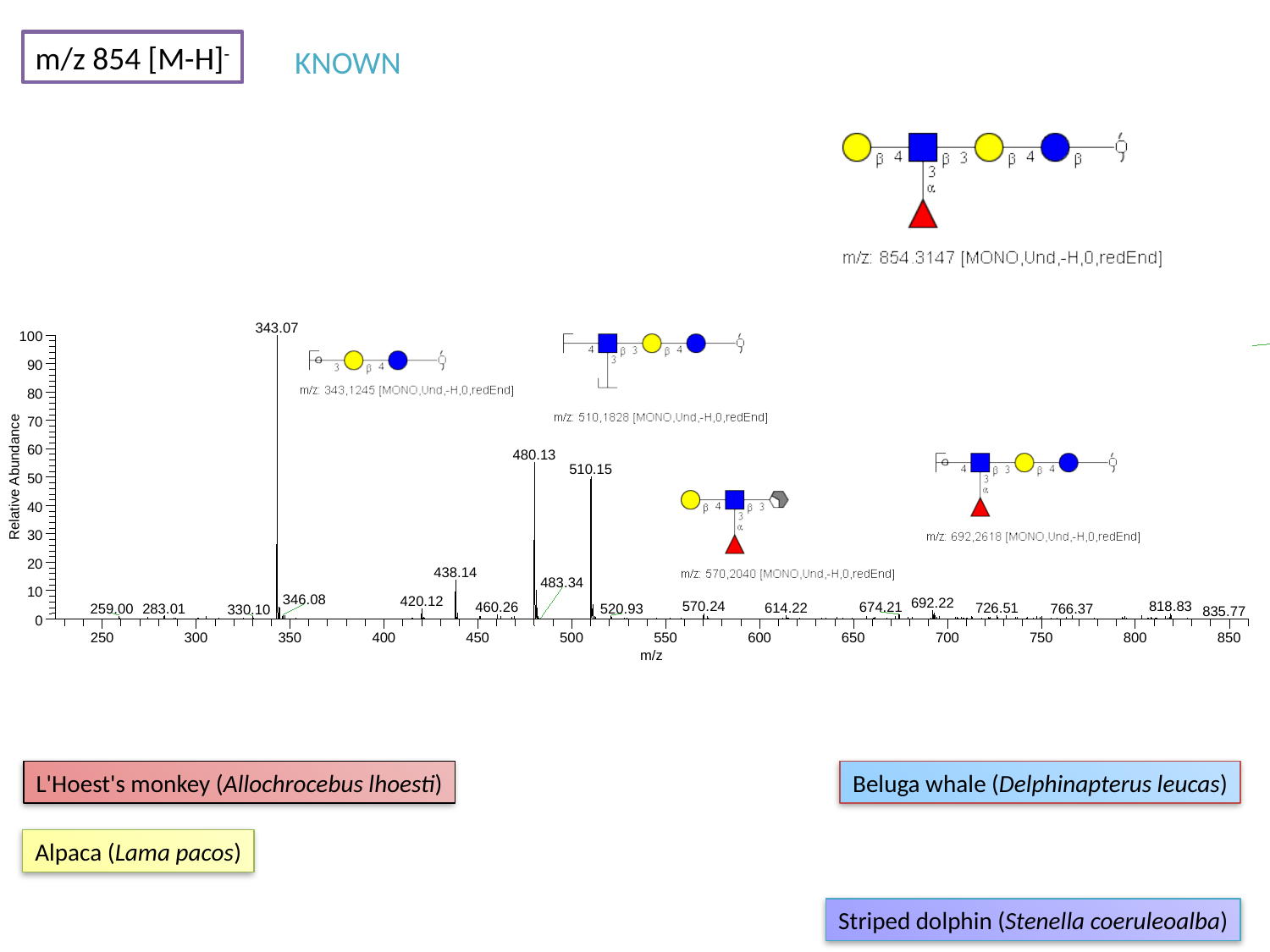

m/z 854 [M-H]^-^
KNOWN
343.07
100
90
80
70
60
480.13
510.15
Relative Abundance
50
40
30
20
438.14
483.34
10
346.08
420.12
692.22
570.24
818.83
460.26
674.21
726.51
614.22
283.01
766.37
259.00
520.93
330.10
835.77
0
250
300
350
400
450
500
550
600
650
700
750
800
850
m/z
L'Hoest's monkey (Allochrocebus lhoesti)
Beluga whale (Delphinapterus leucas)
Alpaca (Lama pacos)
Striped dolphin (Stenella coeruleoalba)

### Slide 56
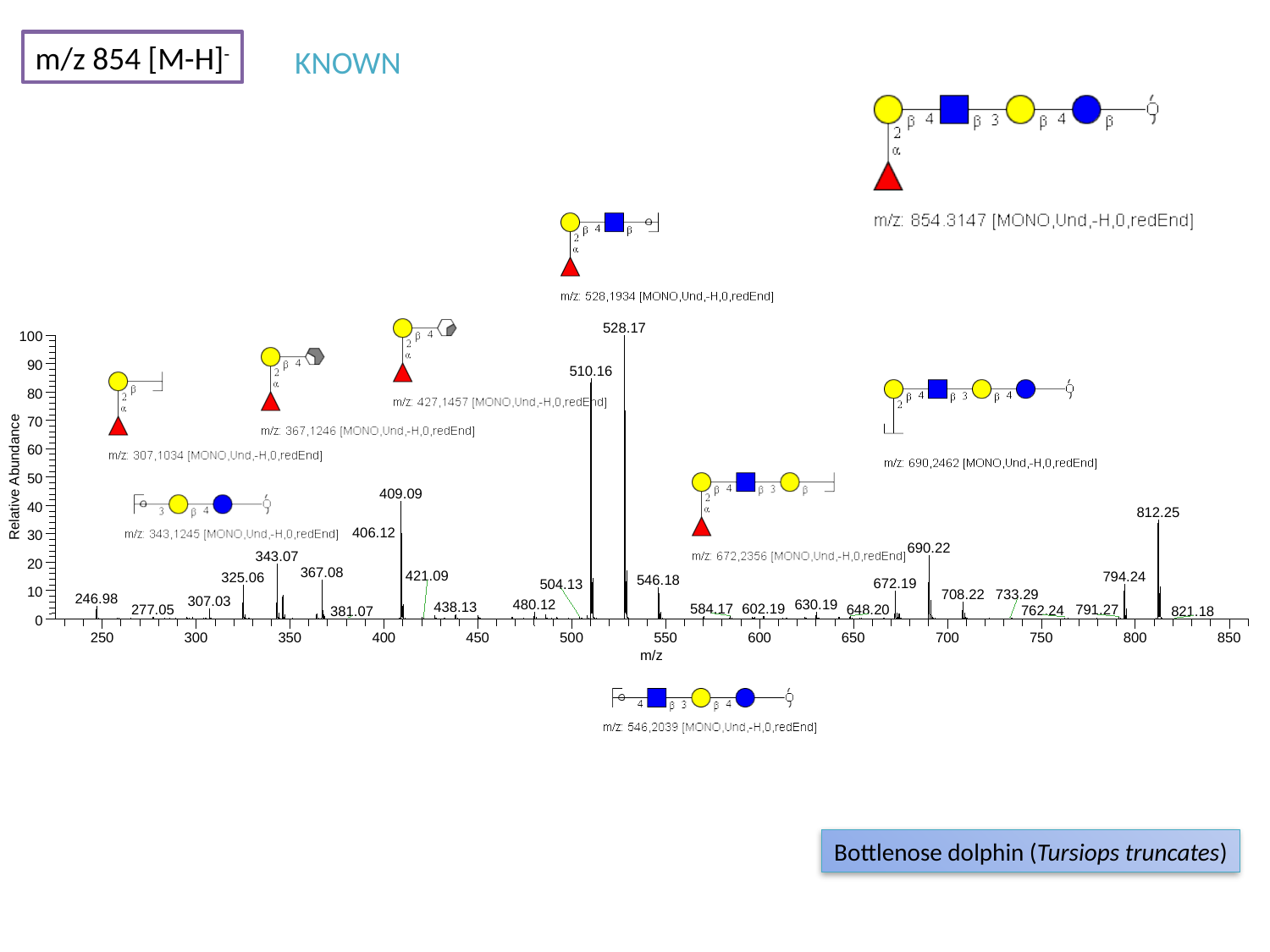

m/z 854 [M-H]^-^
KNOWN
528.17
100
90
510.16
80
70
60
Relative Abundance
50
409.09
40
812.25
406.12
30
690.22
343.07
20
367.08
421.09
794.24
325.06
546.18
672.19
504.13
10
708.22
733.29
246.98
307.03
480.12
630.19
438.13
584.17
602.19
648.20
277.05
791.27
762.24
381.07
821.18
0
250
300
350
400
450
500
550
600
650
700
750
800
850
m/z
Bottlenose dolphin (Tursiops truncates)

### Slide 57
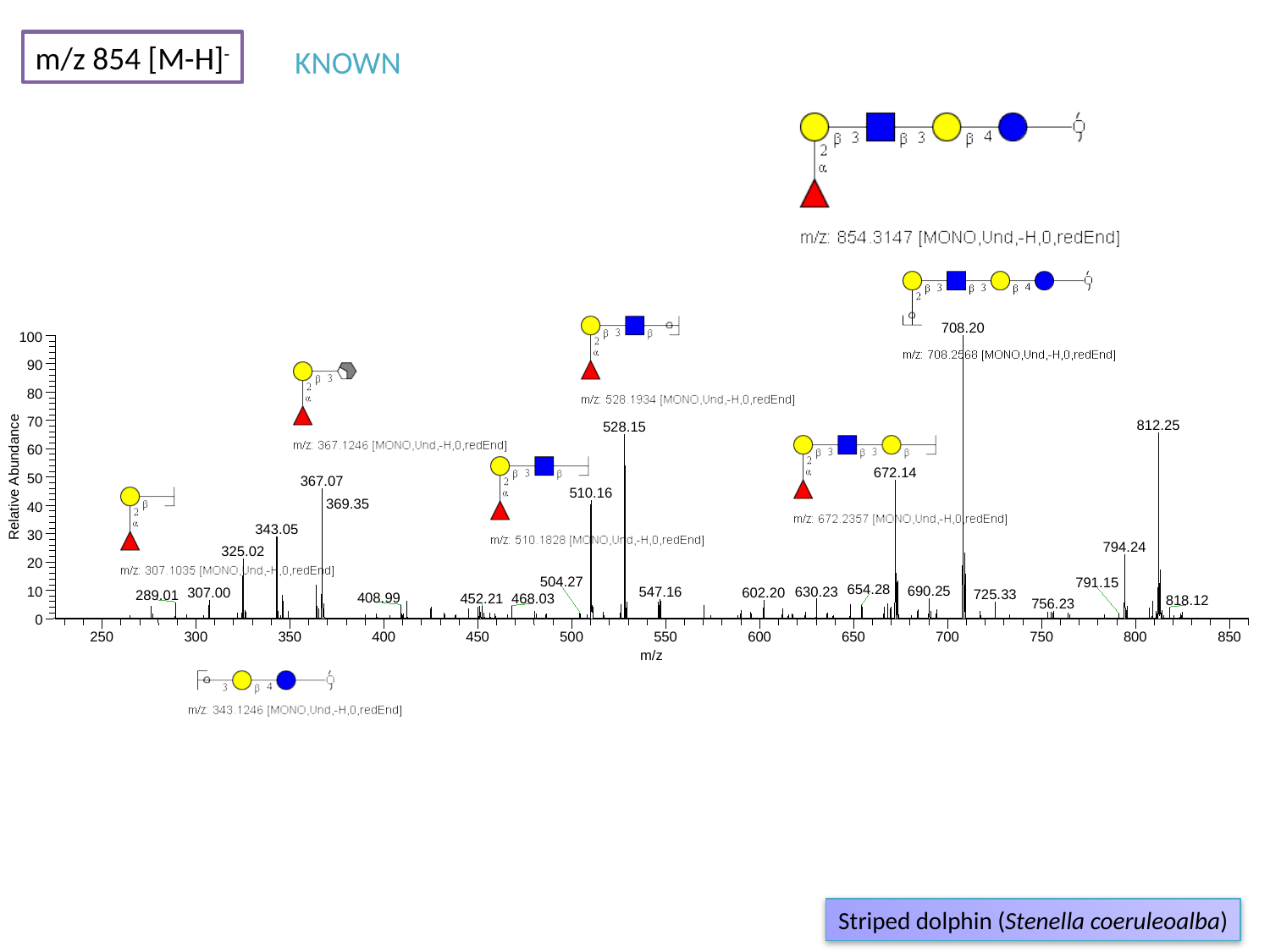

m/z 854 [M-H]^-^
KNOWN
708.20
100
90
80
70
812.25
528.15
60
672.14
Relative Abundance
50
367.07
510.16
369.35
40
343.05
30
794.24
325.02
20
504.27
791.15
654.28
690.25
630.23
10
547.16
307.00
602.20
725.33
289.01
408.99
452.21
468.03
818.12
756.23
0
250
300
350
400
450
500
550
600
650
700
750
800
850
m/z
Striped dolphin (Stenella coeruleoalba)

### Slide 58
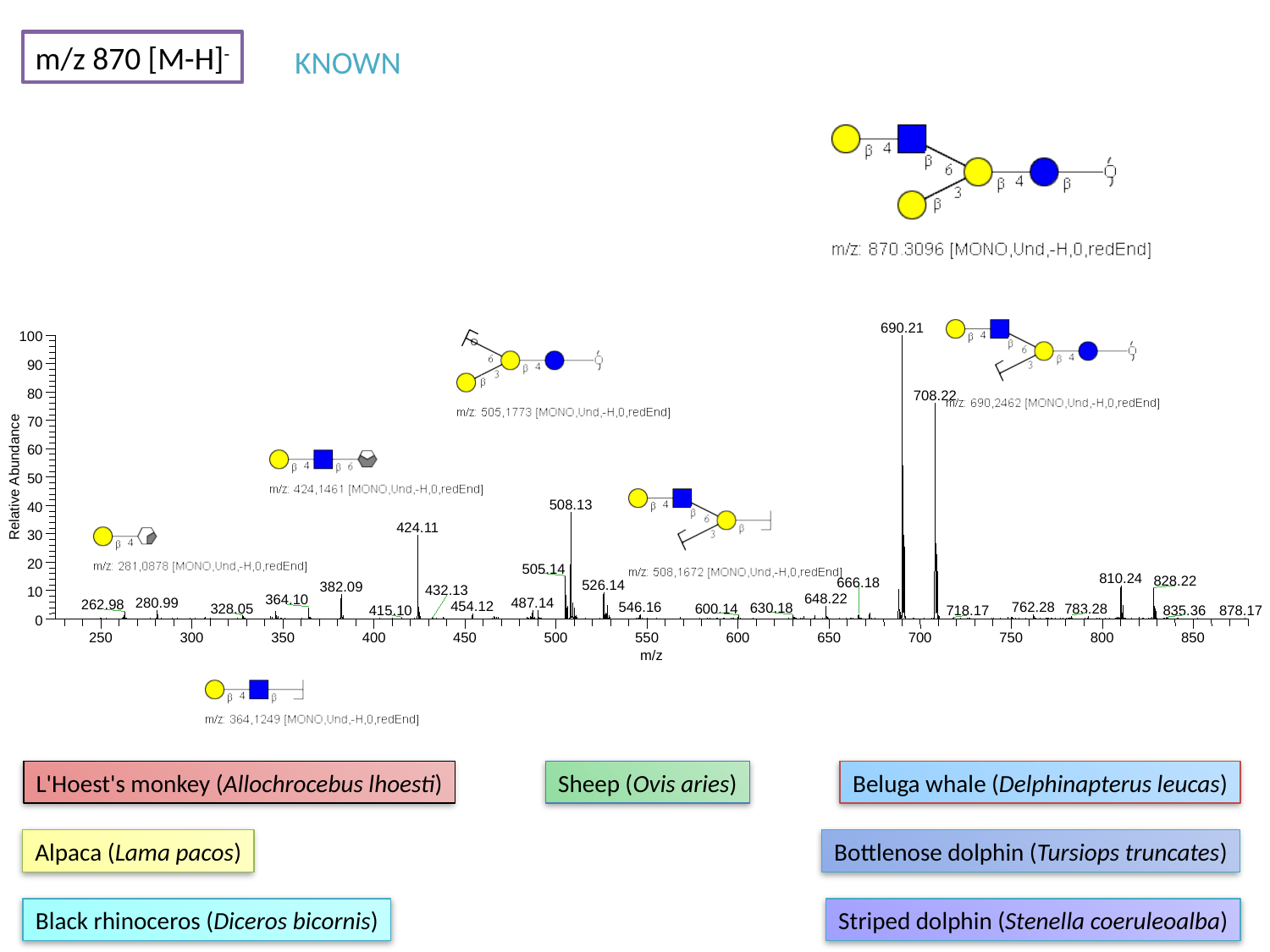

m/z 870 [M-H]^-^
KNOWN
690.21
100
90
80
708.22
70
60
Relative Abundance
50
508.13
40
424.11
30
20
505.14
810.24
828.22
666.18
526.14
382.09
432.13
10
648.22
364.10
280.99
487.14
262.98
454.12
546.16
762.28
630.18
328.05
600.14
783.28
415.10
835.36
718.17
0
250
300
350
400
450
500
550
600
650
700
750
800
850
m/z
878.17
L'Hoest's monkey (Allochrocebus lhoesti)
Sheep (Ovis aries)
Beluga whale (Delphinapterus leucas)
Alpaca (Lama pacos)
Bottlenose dolphin (Tursiops truncates)
Black rhinoceros (Diceros bicornis)
Striped dolphin (Stenella coeruleoalba)

### Slide 59
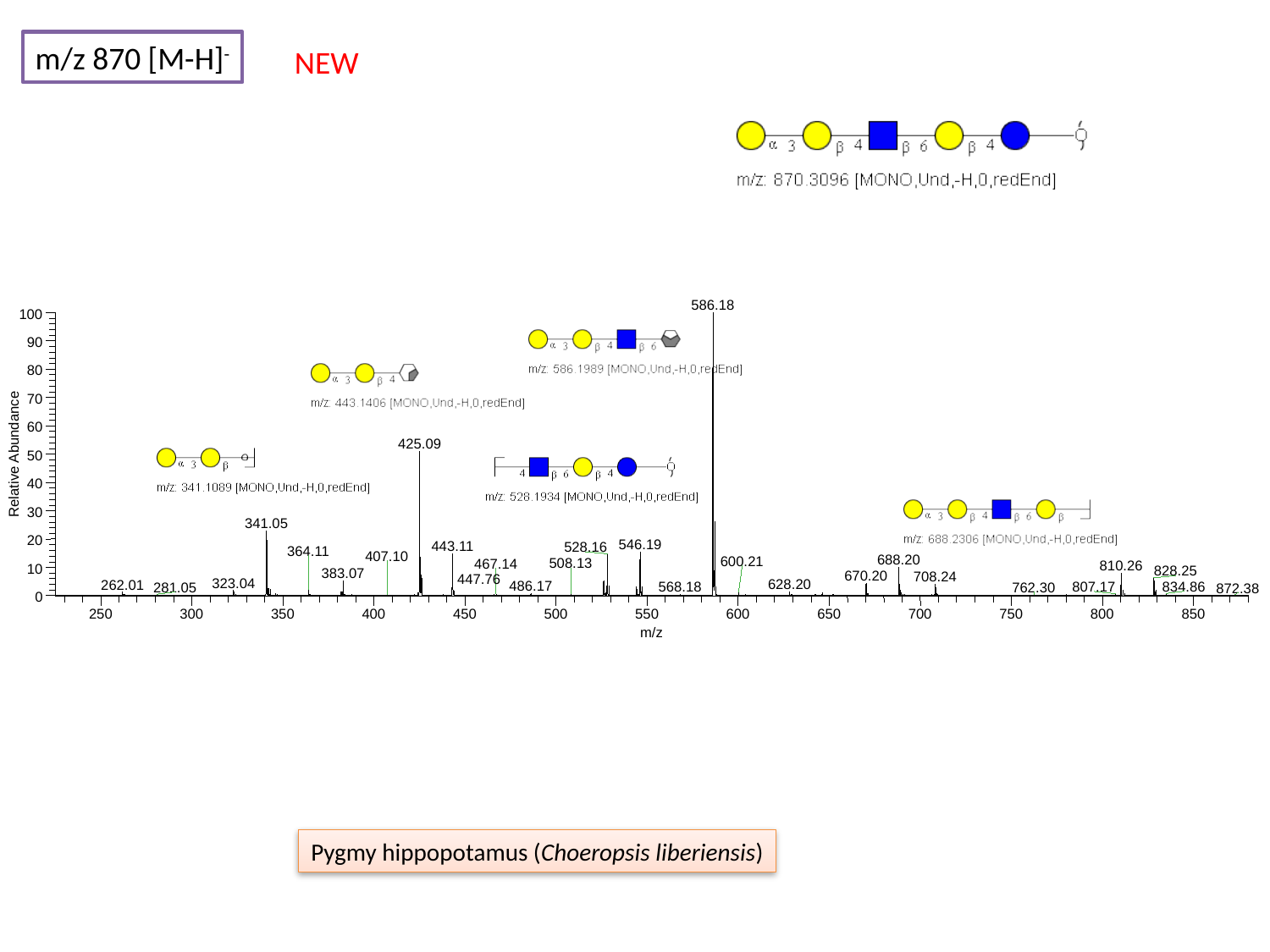

m/z 870 [M-H]^-^
NEW
586.18
100
90
80
70
60
425.09
Relative Abundance
50
40
30
341.05
20
546.19
443.11
528.16
364.11
407.10
688.20
600.21
508.13
467.14
810.26
10
828.25
383.07
670.20
708.24
323.04
628.20
262.01
486.17
568.18
834.86
807.17
281.05
762.30
872.38
0
250
300
350
400
450
500
550
600
650
700
750
800
850
m/z
447.76
Pygmy hippopotamus (Choeropsis liberiensis)

### Slide 60
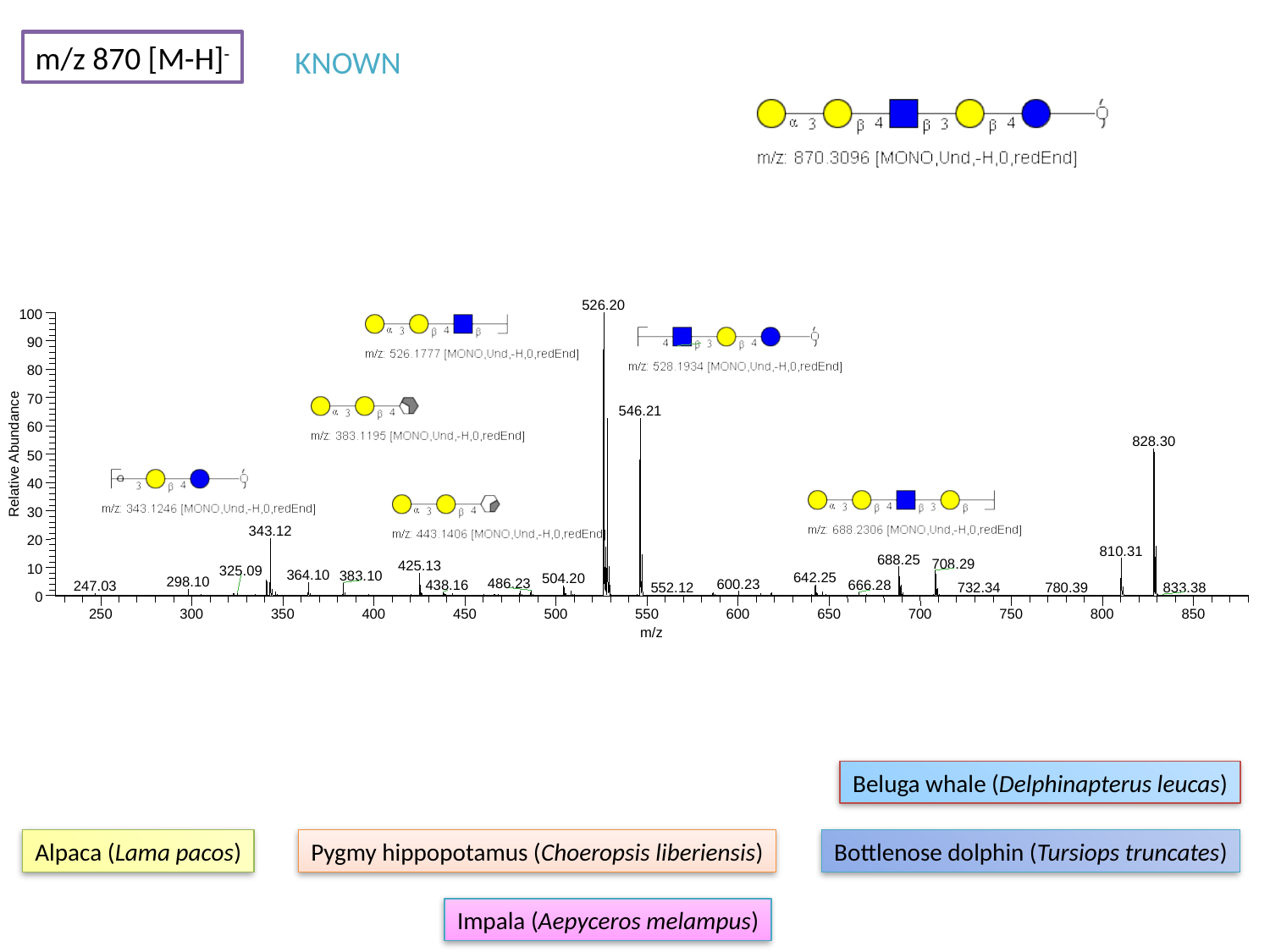

m/z 870 [M-H]^-^
KNOWN
526.20
100
90
80
70
546.21
60
828.30
Relative Abundance
50
40
30
343.12
20
810.31
688.25
708.29
425.13
10
325.09
364.10
383.10
642.25
504.20
298.10
486.23
600.23
438.16
666.28
247.03
732.34
833.38
780.39
552.12
0
250
300
350
400
450
500
550
600
650
700
750
800
850
m/z
Beluga whale (Delphinapterus leucas)
Alpaca (Lama pacos)
Pygmy hippopotamus (Choeropsis liberiensis)
Bottlenose dolphin (Tursiops truncates)
Impala (Aepyceros melampus)

### Slide 61
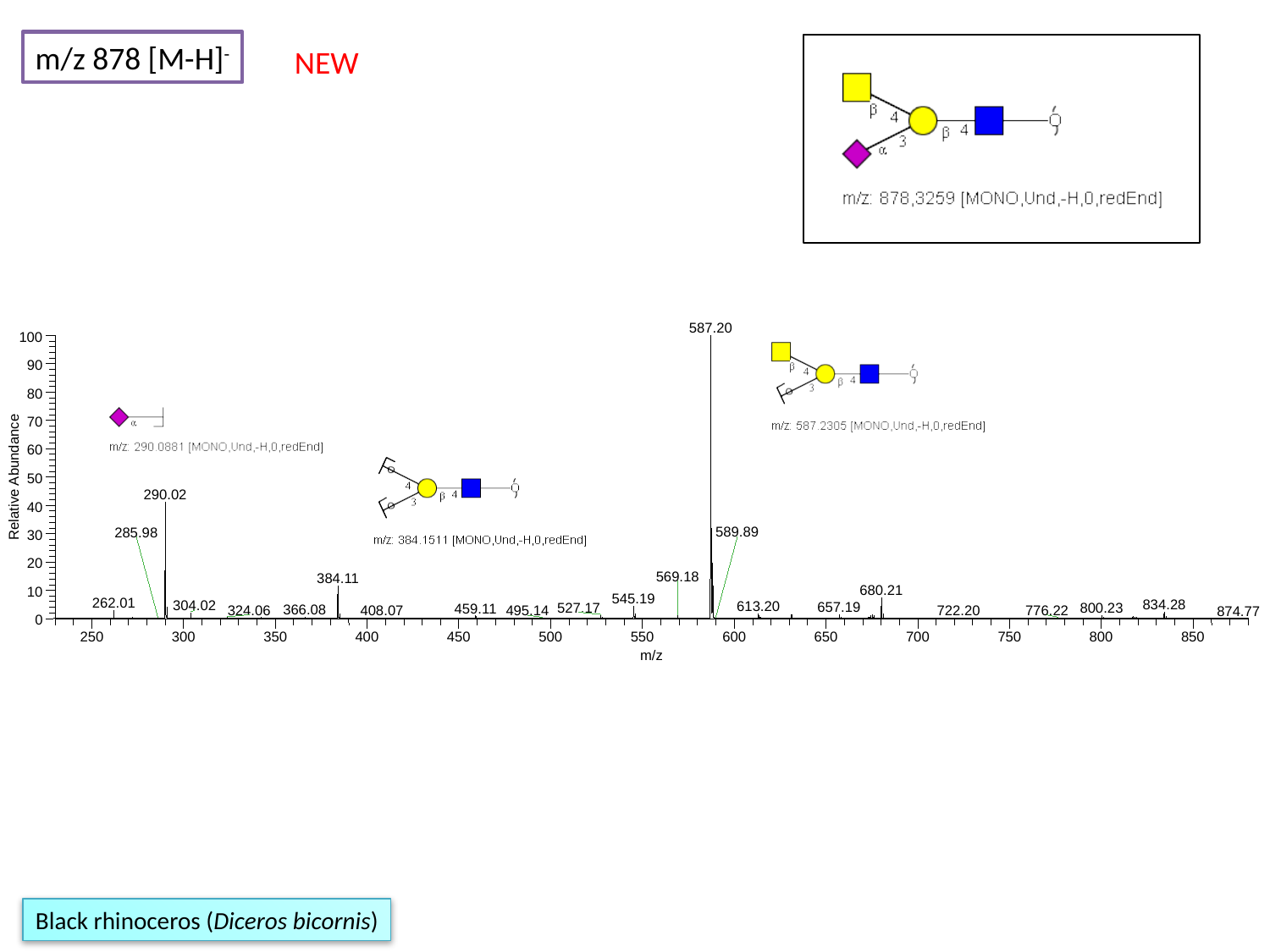

m/z 878 [M-H]^-^
NEW
587.20
100
90
80
70
60
Relative Abundance
50
290.02
40
589.89
285.98
30
20
569.18
384.11
680.21
10
545.19
262.01
834.28
304.02
613.20
657.19
527.17
800.23
459.11
366.08
324.06
722.20
776.22
408.07
495.14
874.77
0
250
300
350
400
450
500
550
600
650
700
750
800
850
m/z
Black rhinoceros (Diceros bicornis)

### Slide 62
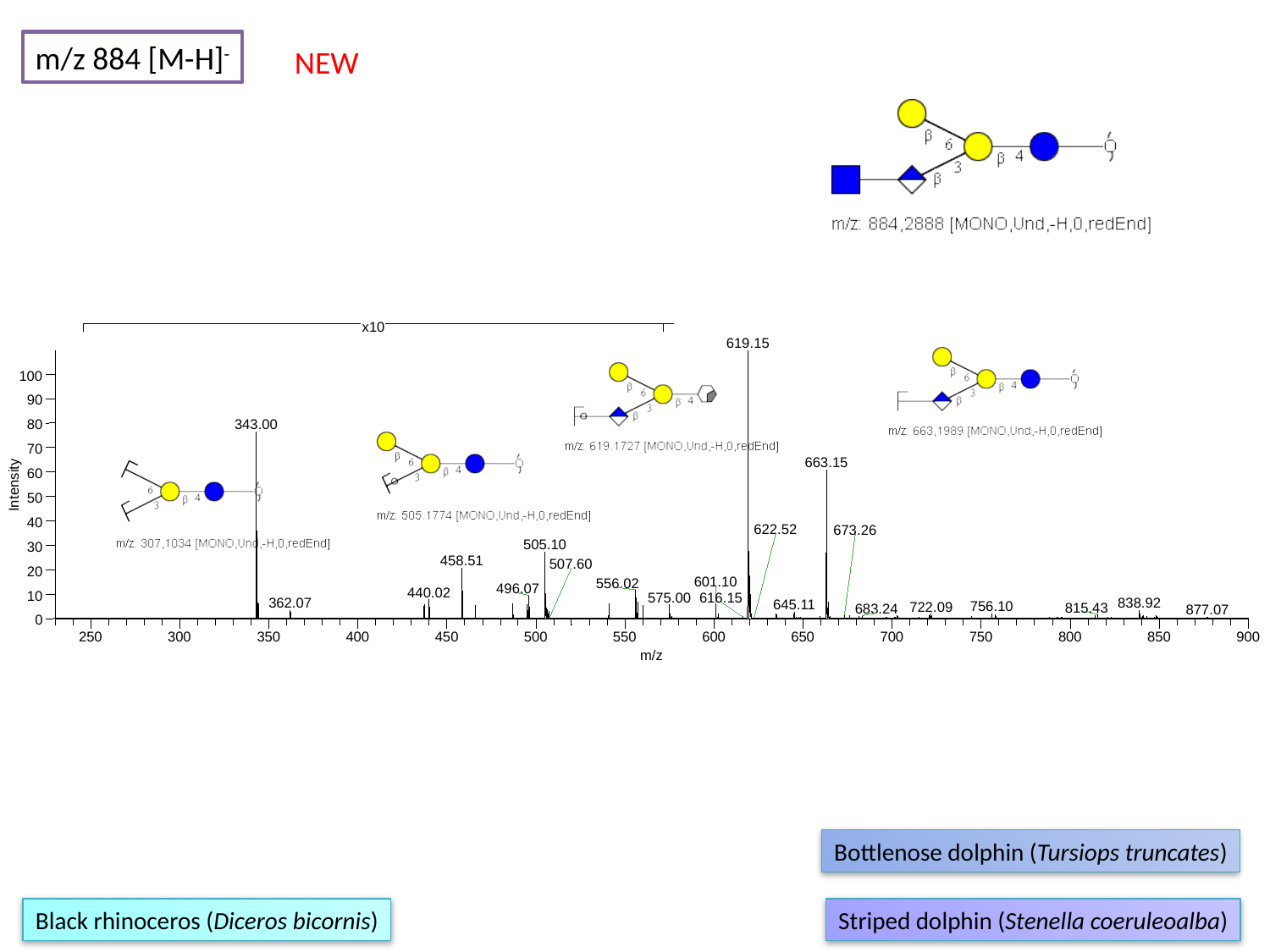

m/z 884 [M-H]^-^
NEW
x10
619.15
100
90
80
343.00
70
663.15
60
Intensity
50
40
622.52
673.26
505.10
30
458.51
507.60
20
601.10
556.02
496.07
440.02
10
575.00
616.15
362.07
838.92
645.11
756.10
722.09
815.43
683.24
877.07
0
250
300
350
400
450
500
550
600
650
700
750
800
850
900
m/z
Bottlenose dolphin (Tursiops truncates)
Black rhinoceros (Diceros bicornis)
Striped dolphin (Stenella coeruleoalba)

### Slide 63
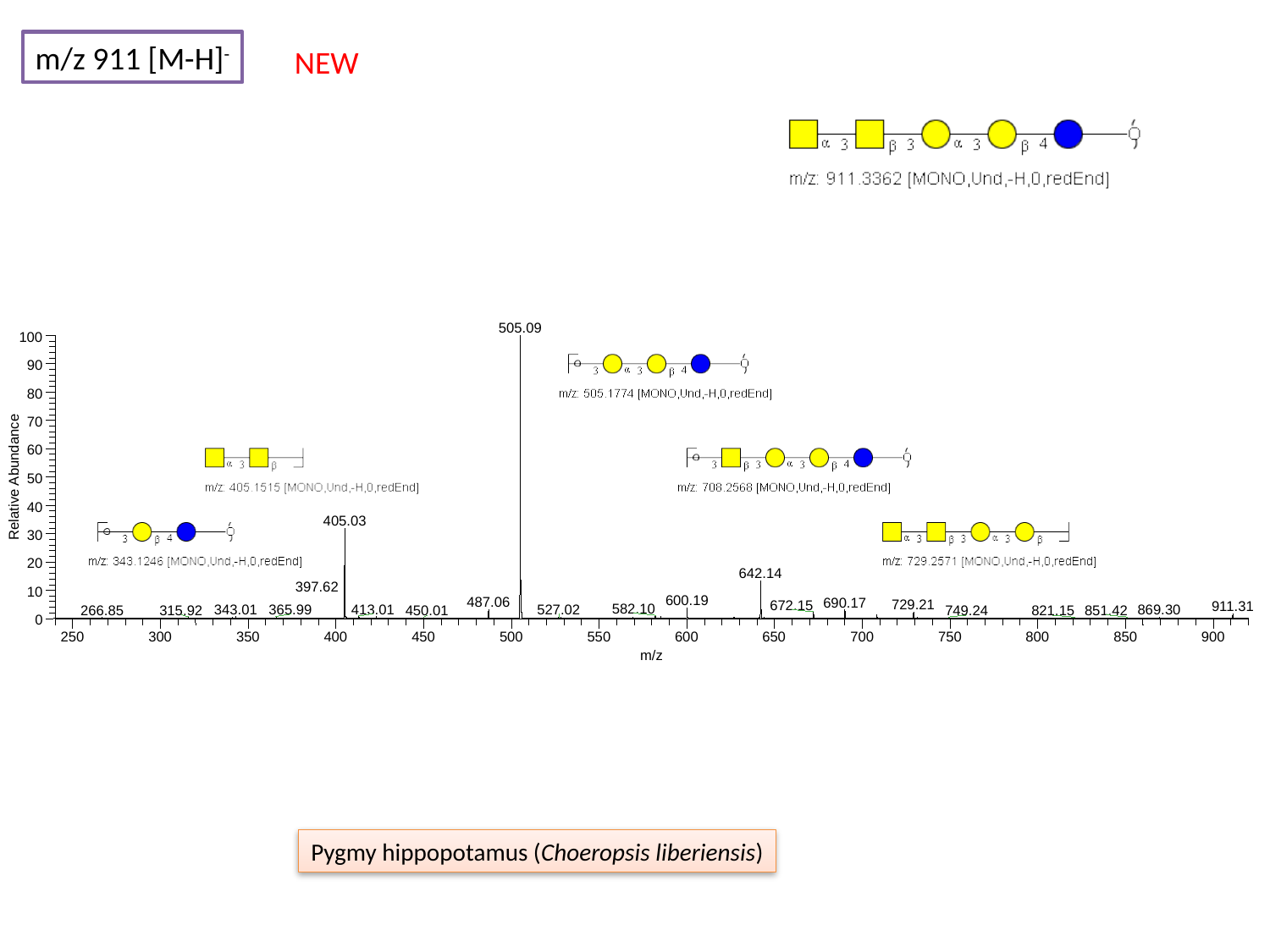

m/z 911 [M-H]^-^
NEW
505.09
100
90
80
70
60
Relative Abundance
50
40
405.03
30
20
642.14
397.62
10
600.19
487.06
690.17
729.21
672.15
911.31
582.10
343.01
413.01
365.99
527.02
869.30
266.85
315.92
851.42
749.24
450.01
821.15
0
250
300
350
400
450
500
550
600
650
700
750
800
850
900
m/z
Pygmy hippopotamus (Choeropsis liberiensis)

### Slide 64
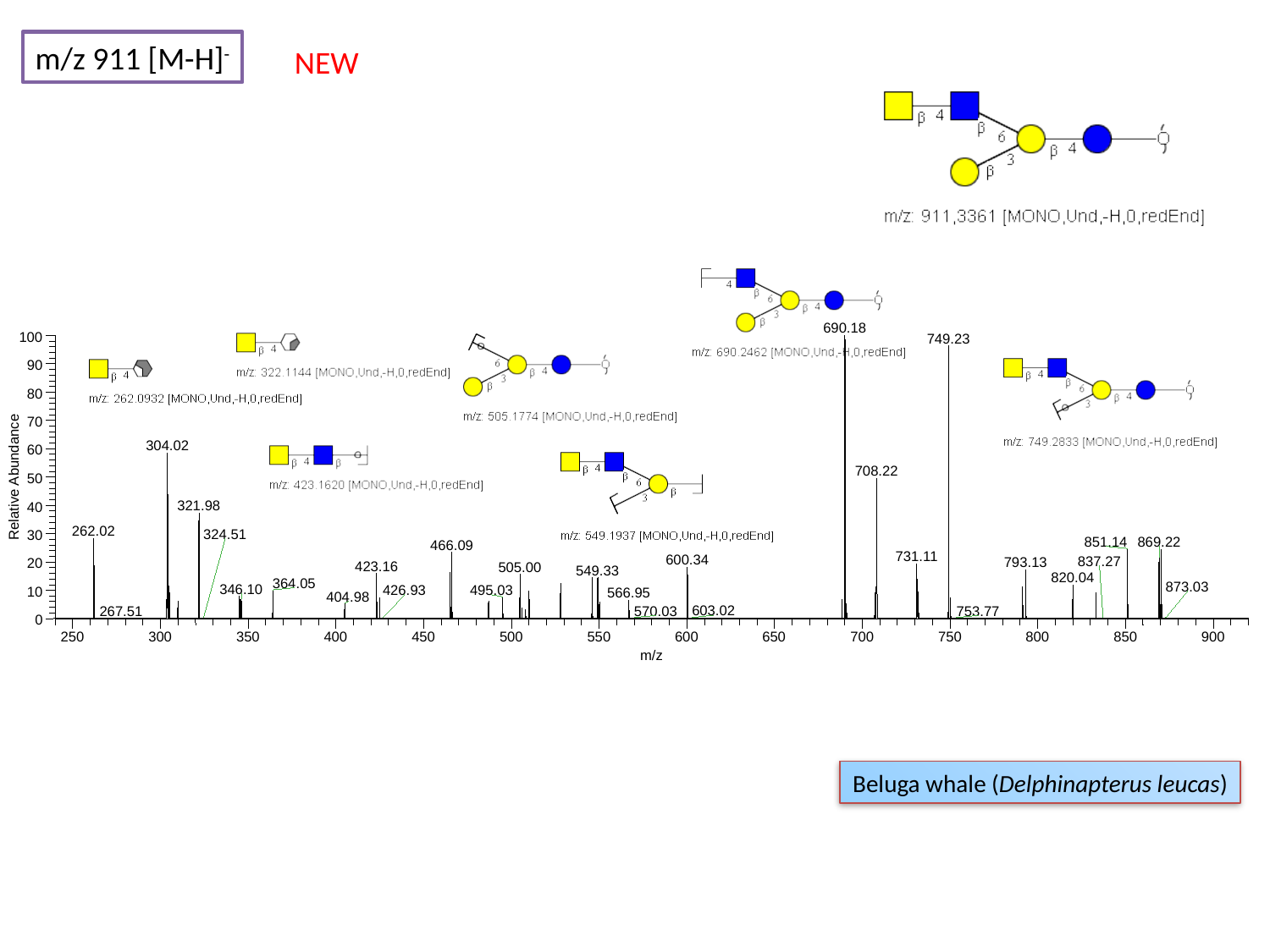

m/z 911 [M-H]^-^
NEW
690.18
100
749.23
90
80
70
304.02
60
708.22
Relative Abundance
50
321.98
40
262.02
324.51
30
869.22
851.14
466.09
731.11
600.34
793.13
20
423.16
505.00
549.33
820.04
364.05
873.03
346.10
426.93
495.03
10
566.95
404.98
603.02
0
250
300
350
400
450
500
550
600
650
700
750
800
850
900
m/z
837.27
267.51
570.03
753.77
Beluga whale (Delphinapterus leucas)

### Slide 65
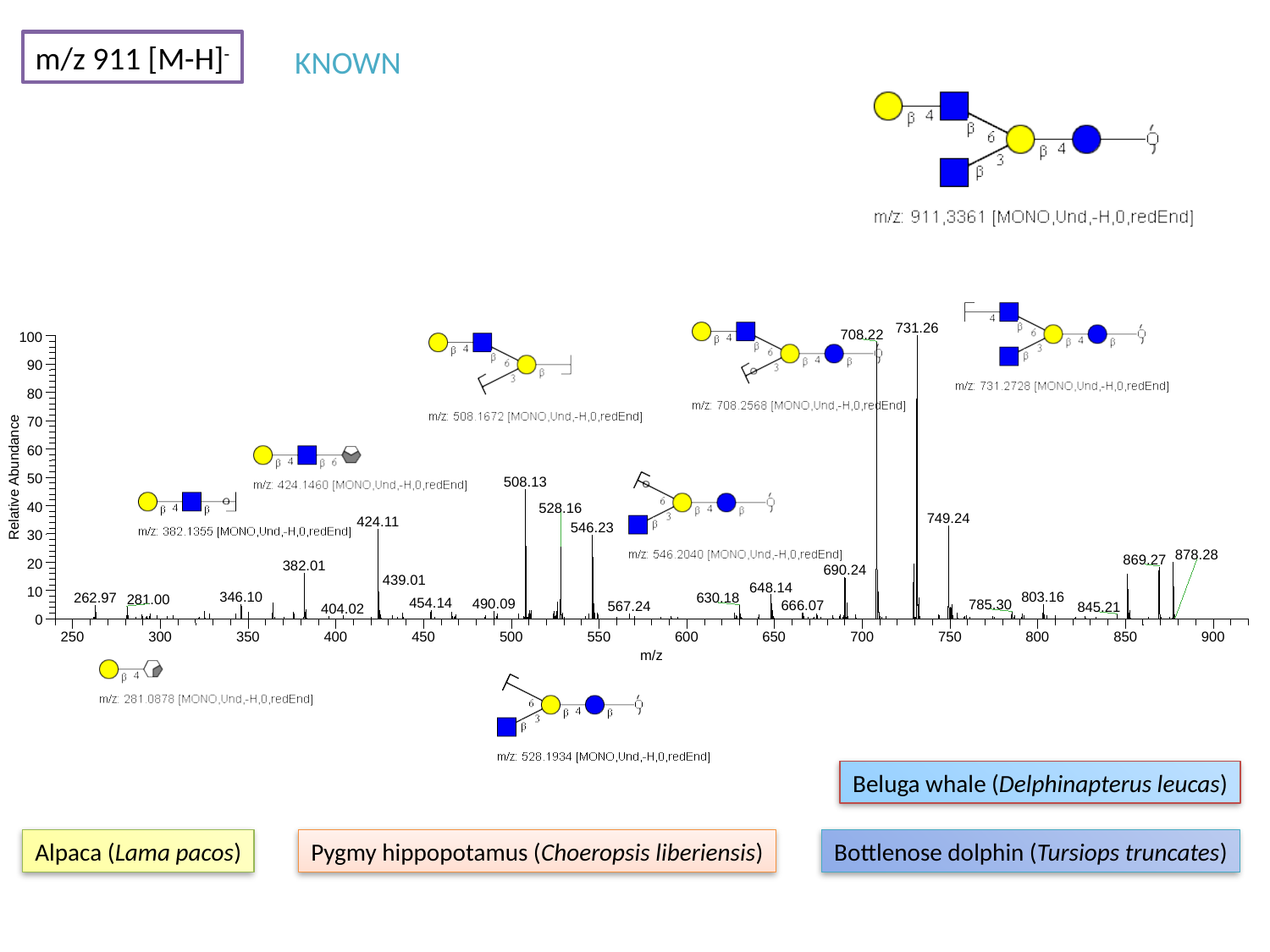

m/z 911 [M-H]^-^
KNOWN
731.26
708.22
100
90
80
70
60
Relative Abundance
50
508.13
40
528.16
749.24
424.11
546.23
30
878.28
869.27
20
382.01
690.24
439.01
648.14
10
346.10
803.16
262.97
630.18
281.00
454.14
490.09
785.30
666.07
567.24
845.21
404.02
0
250
300
350
400
450
500
550
600
650
700
750
800
850
900
m/z
Beluga whale (Delphinapterus leucas)
Alpaca (Lama pacos)
Pygmy hippopotamus (Choeropsis liberiensis)
Bottlenose dolphin (Tursiops truncates)

### Slide 66
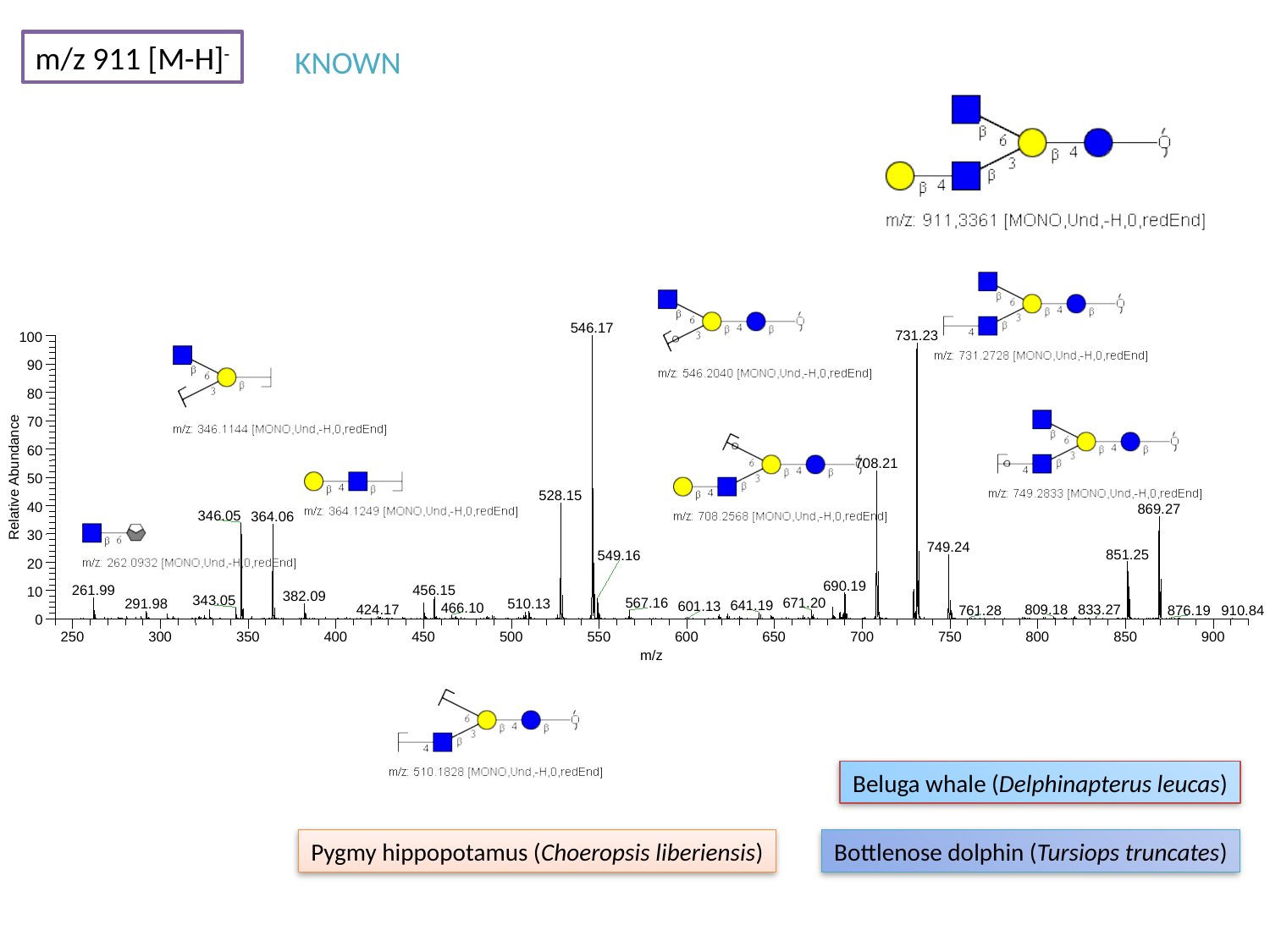

m/z 911 [M-H]^-^
KNOWN
546.17
731.23
100
90
80
70
60
708.21
Relative Abundance
50
528.15
40
869.27
346.05
364.06
30
749.24
851.25
549.16
20
690.19
456.15
261.99
10
382.09
343.05
567.16
671.20
291.98
510.13
641.19
466.10
424.17
833.27
809.18
761.28
0
250
300
350
400
450
500
550
600
650
700
750
800
850
900
m/z
601.13
876.19
910.84
Beluga whale (Delphinapterus leucas)
Pygmy hippopotamus (Choeropsis liberiensis)
Bottlenose dolphin (Tursiops truncates)

### Slide 67
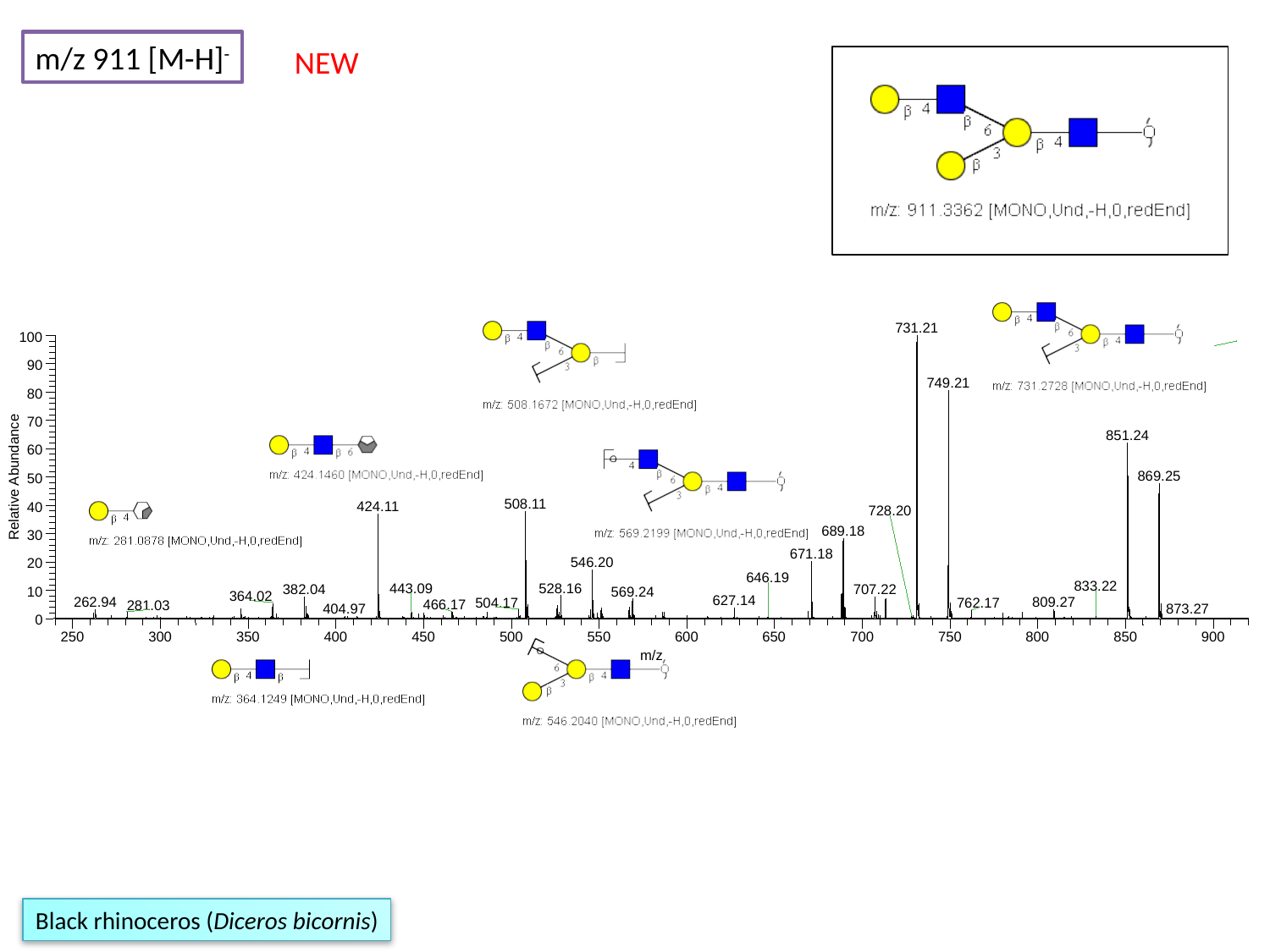

m/z 911 [M-H]^-^
NEW
731.21
100
90
749.21
80
70
851.24
60
869.25
Relative Abundance
50
508.11
424.11
40
728.20
689.18
30
671.18
546.20
20
646.19
833.22
528.16
443.09
707.22
382.04
569.24
10
364.02
627.14
809.27
262.94
504.17
762.17
466.17
281.03
404.97
0
250
300
350
400
450
500
550
600
650
700
750
800
850
900
m/z
873.27
Black rhinoceros (Diceros bicornis)

### Slide 68
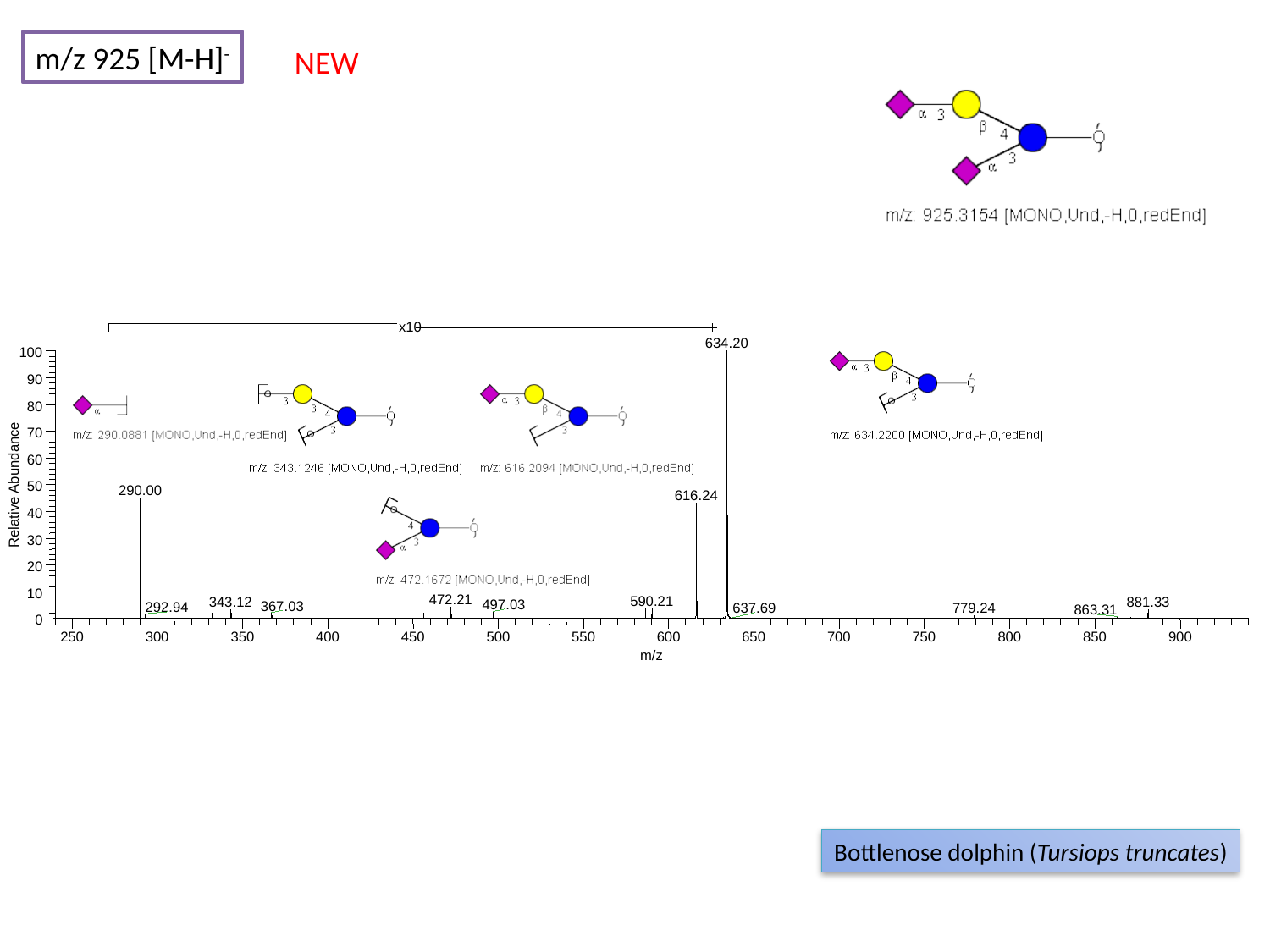

m/z 925 [M-H]^-^
NEW
x10
634.20
100
90
80
70
60
Relative Abundance
50
290.00
616.24
40
30
20
10
472.21
590.21
343.12
881.33
497.03
367.03
292.94
637.69
779.24
863.31
0
250
300
350
400
450
500
550
600
650
700
750
800
850
900
m/z
Bottlenose dolphin (Tursiops truncates)

### Slide 69
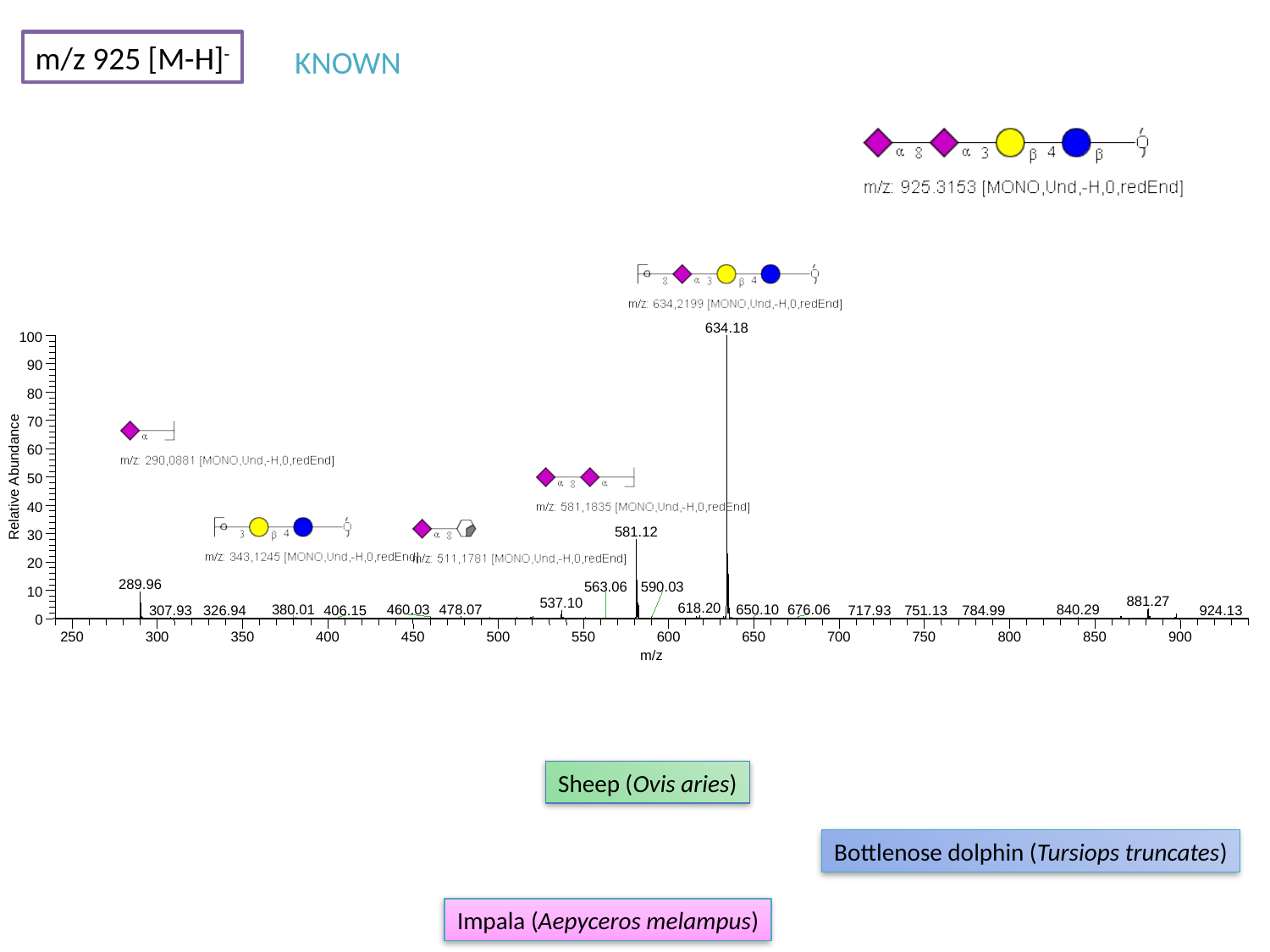

m/z 925 [M-H]^-^
KNOWN
634.18
100
90
80
70
60
Relative Abundance
50
40
581.12
30
20
289.96
563.06
590.03
10
881.27
537.10
618.20
478.07
650.10
676.06
380.01
460.03
840.29
307.93
717.93
784.99
924.13
326.94
406.15
751.13
0
250
300
350
400
450
500
550
600
650
700
750
800
850
900
m/z
Sheep (Ovis aries)
Bottlenose dolphin (Tursiops truncates)
Impala (Aepyceros melampus)

### Slide 70
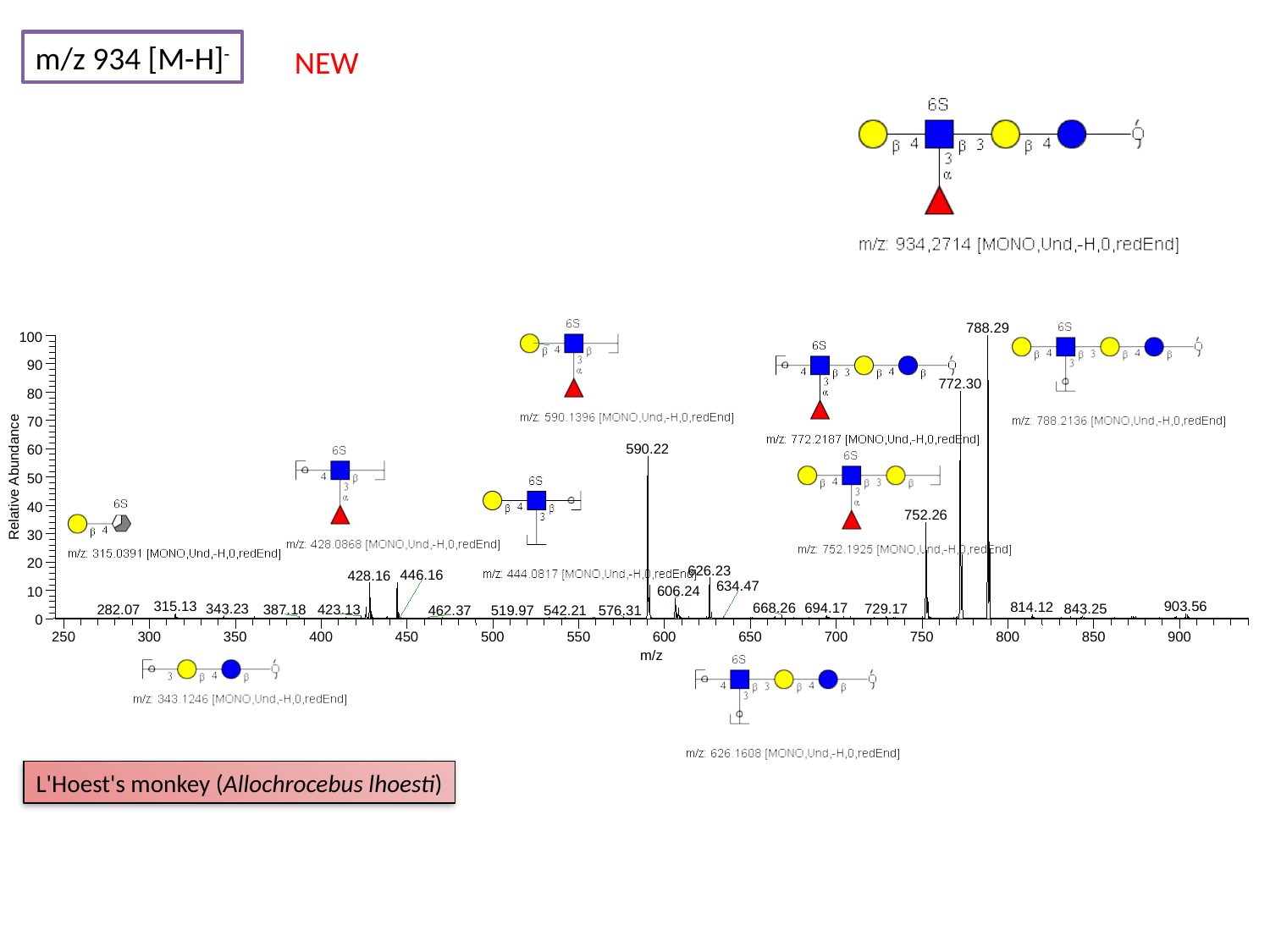

m/z 934 [M-H]^-^
NEW
788.29
100
90
772.30
80
70
590.22
60
Relative Abundance
50
40
752.26
30
20
626.23
446.16
428.16
634.47
606.24
10
315.13
903.56
814.12
668.26
694.17
729.17
343.23
843.25
423.13
282.07
387.18
462.37
542.21
576.31
0
250
300
350
400
450
500
550
600
650
700
750
800
850
900
m/z
519.97
L'Hoest's monkey (Allochrocebus lhoesti)

### Slide 71
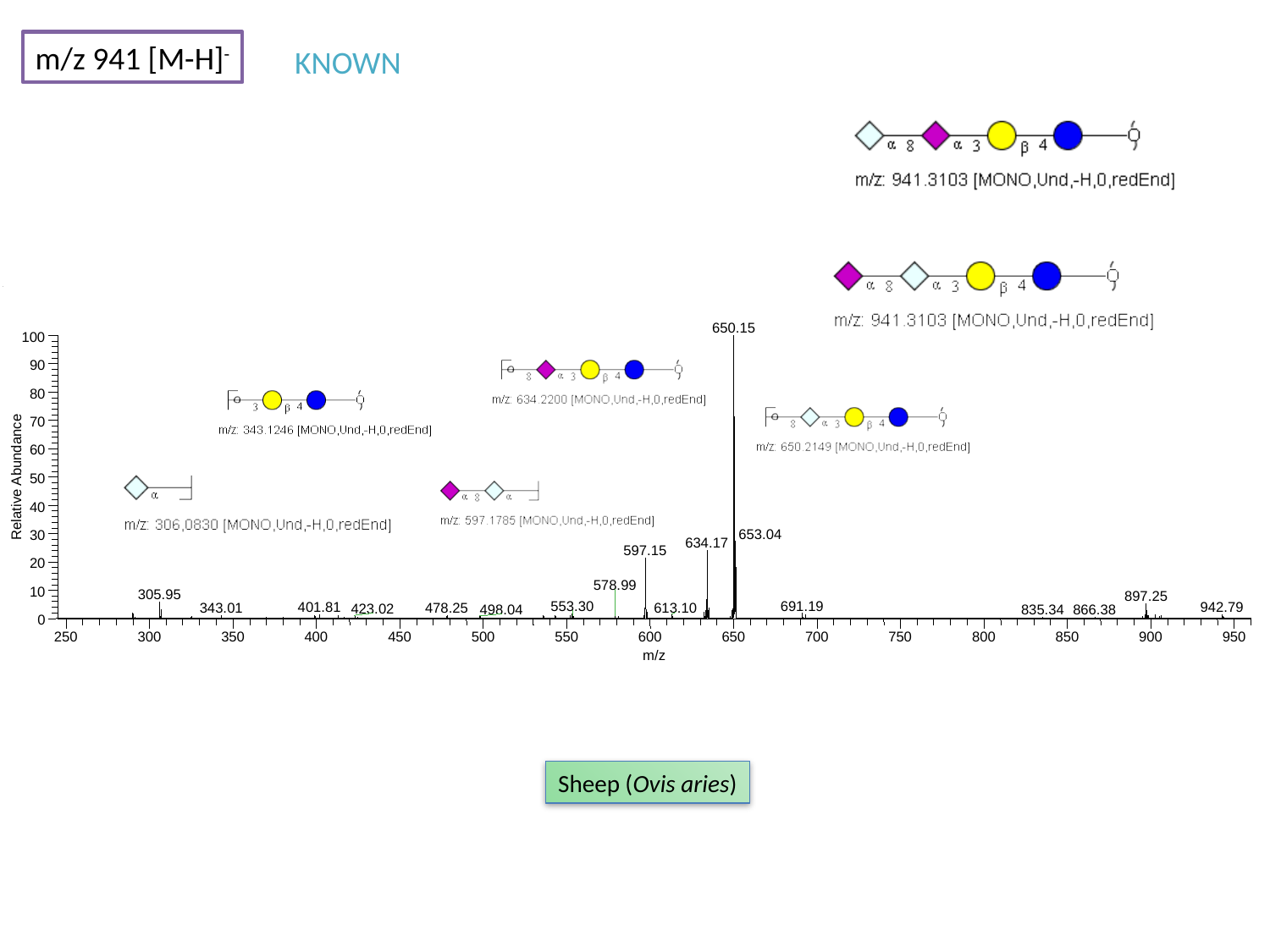

m/z 941 [M-H]^-^
KNOWN
650.15
100
90
80
70
60
Relative Abundance
50
40
653.04
30
634.17
597.15
20
578.99
10
305.95
897.25
691.19
553.30
401.81
942.79
613.10
343.01
478.25
423.02
498.04
835.34
866.38
0
250
300
350
400
450
500
550
600
650
700
750
800
850
900
950
m/z
Sheep (Ovis aries)

### Slide 72
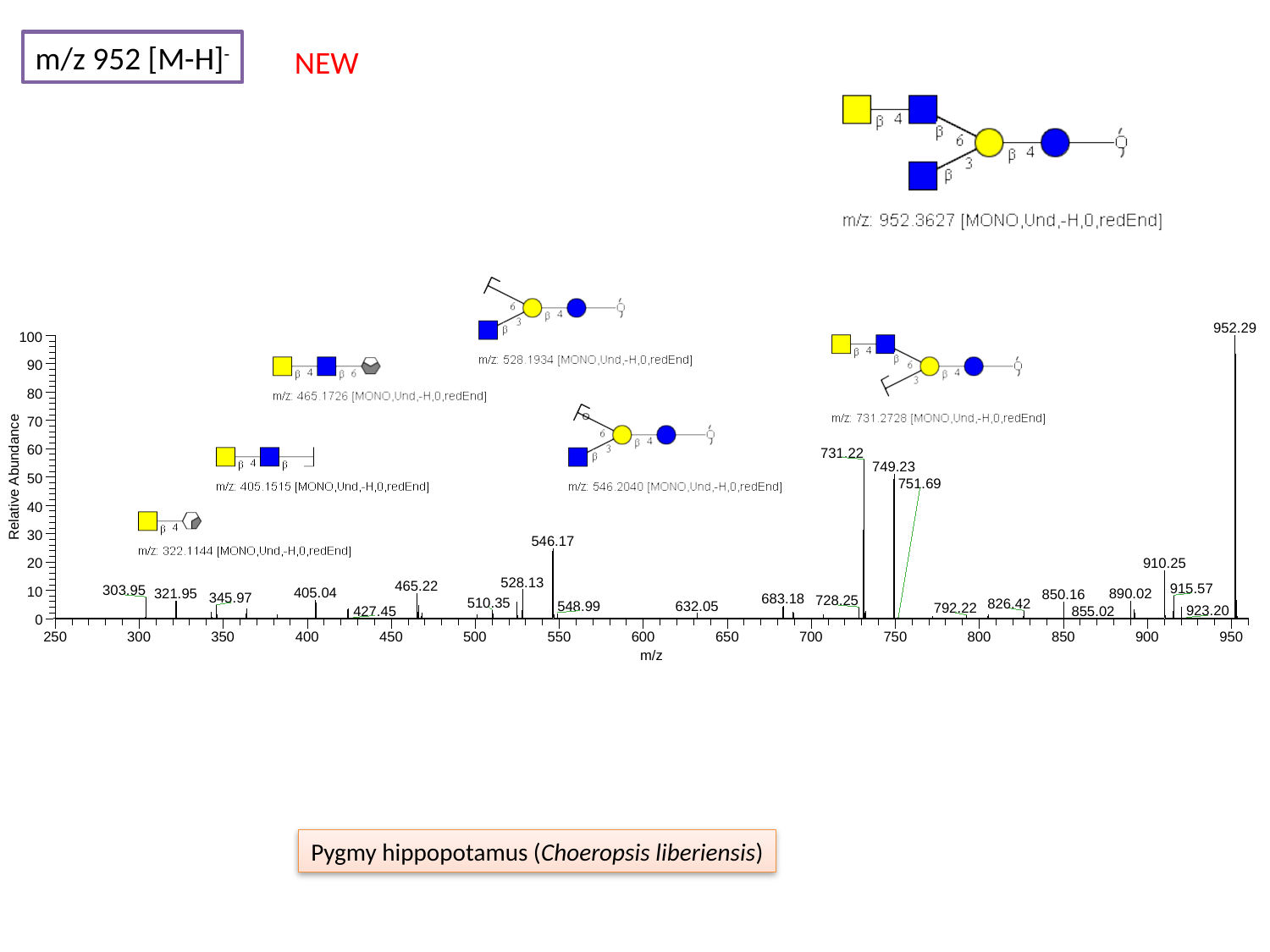

m/z 952 [M-H]^-^
NEW
952.29
100
90
80
70
60
731.22
749.23
Relative Abundance
50
40
30
546.17
910.25
20
528.13
465.22
915.57
303.95
10
405.04
890.02
321.95
850.16
345.97
683.18
728.25
510.35
826.42
632.05
548.99
792.22
0
250
300
350
400
450
500
550
600
650
700
750
800
850
900
950
m/z
751.69
923.20
427.45
855.02
Pygmy hippopotamus (Choeropsis liberiensis)

### Slide 73
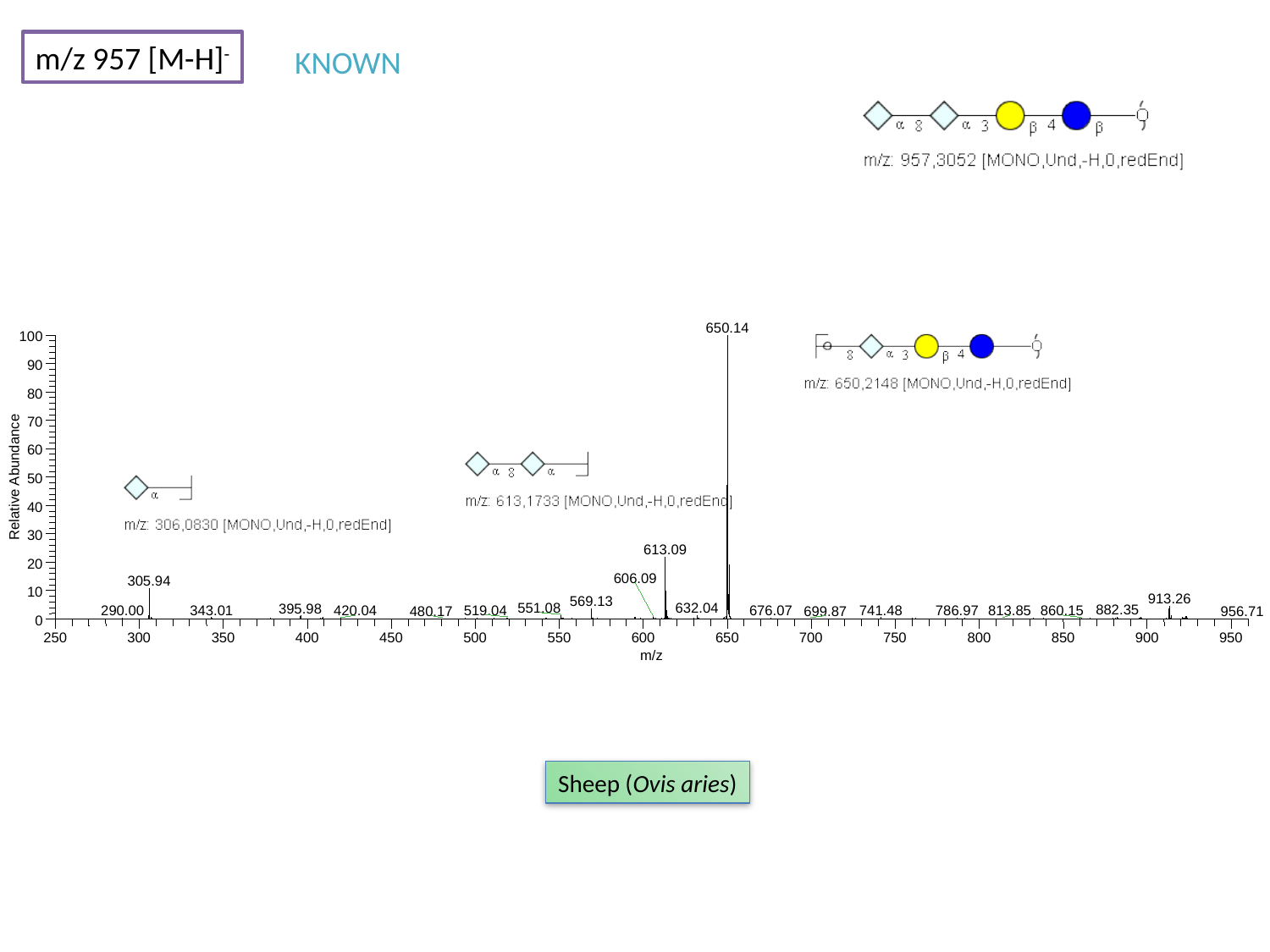

m/z 957 [M-H]^-^
KNOWN
650.14
100
90
80
70
60
Relative Abundance
50
40
30
613.09
20
606.09
305.94
10
913.26
569.13
551.08
632.04
395.98
882.35
343.01
741.48
519.04
290.00
420.04
676.07
786.97
813.85
860.15
480.17
699.87
956.71
0
250
300
350
400
450
500
550
600
650
700
750
800
850
900
950
m/z
Sheep (Ovis aries)

### Slide 74
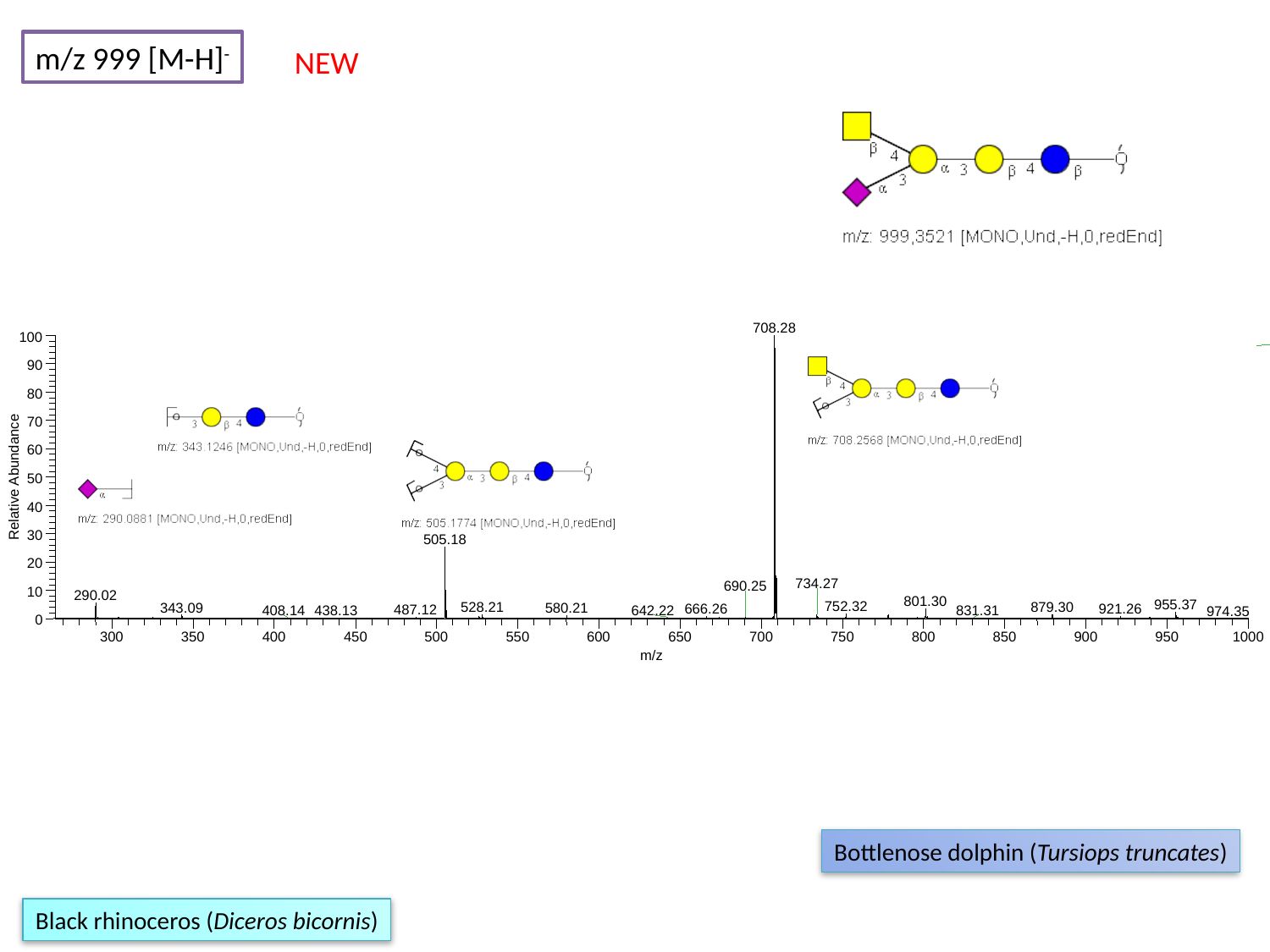

m/z 999 [M-H]^-^
NEW
708.28
100
90
80
70
60
Relative Abundance
50
40
30
505.18
20
734.27
690.25
10
290.02
801.30
955.37
752.32
528.21
879.30
343.09
580.21
921.26
666.26
487.12
438.13
642.22
831.31
408.14
974.35
0
300
350
400
450
500
550
600
650
700
750
800
850
900
950
1000
m/z
Bottlenose dolphin (Tursiops truncates)
Black rhinoceros (Diceros bicornis)

### Slide 75
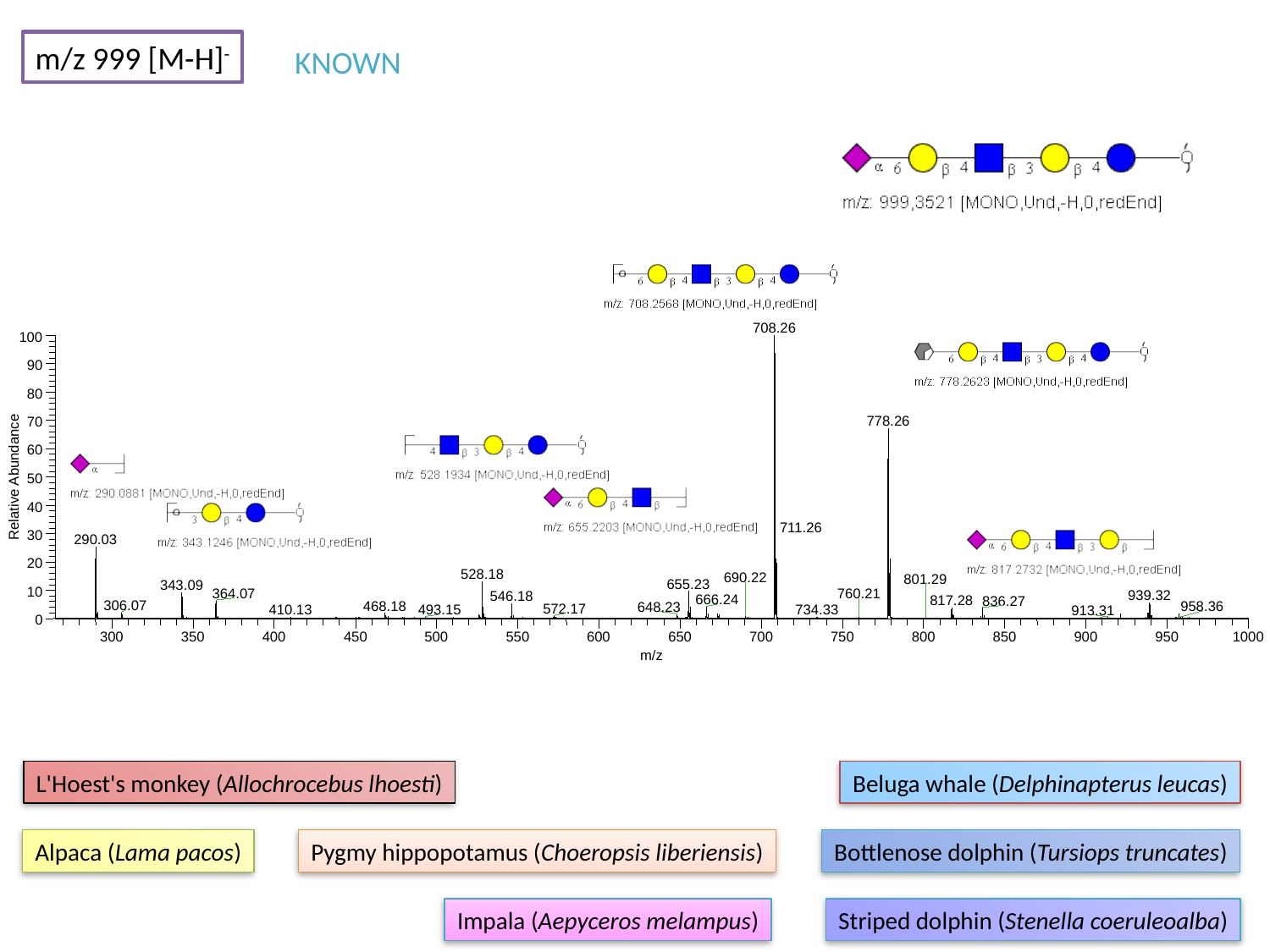

m/z 999 [M-H]^-^
KNOWN
708.26
100
90
80
778.26
70
60
Relative Abundance
50
40
30
290.03
20
528.18
690.22
655.23
343.09
10
364.07
939.32
546.18
666.24
817.28
836.27
306.07
468.18
648.23
572.17
410.13
493.15
734.33
0
300
350
400
450
500
550
600
650
700
750
800
850
900
950
1000
m/z
711.26
801.29
760.21
958.36
913.31
L'Hoest's monkey (Allochrocebus lhoesti)
Beluga whale (Delphinapterus leucas)
Alpaca (Lama pacos)
Pygmy hippopotamus (Choeropsis liberiensis)
Bottlenose dolphin (Tursiops truncates)
Impala (Aepyceros melampus)
Striped dolphin (Stenella coeruleoalba)

### Slide 76
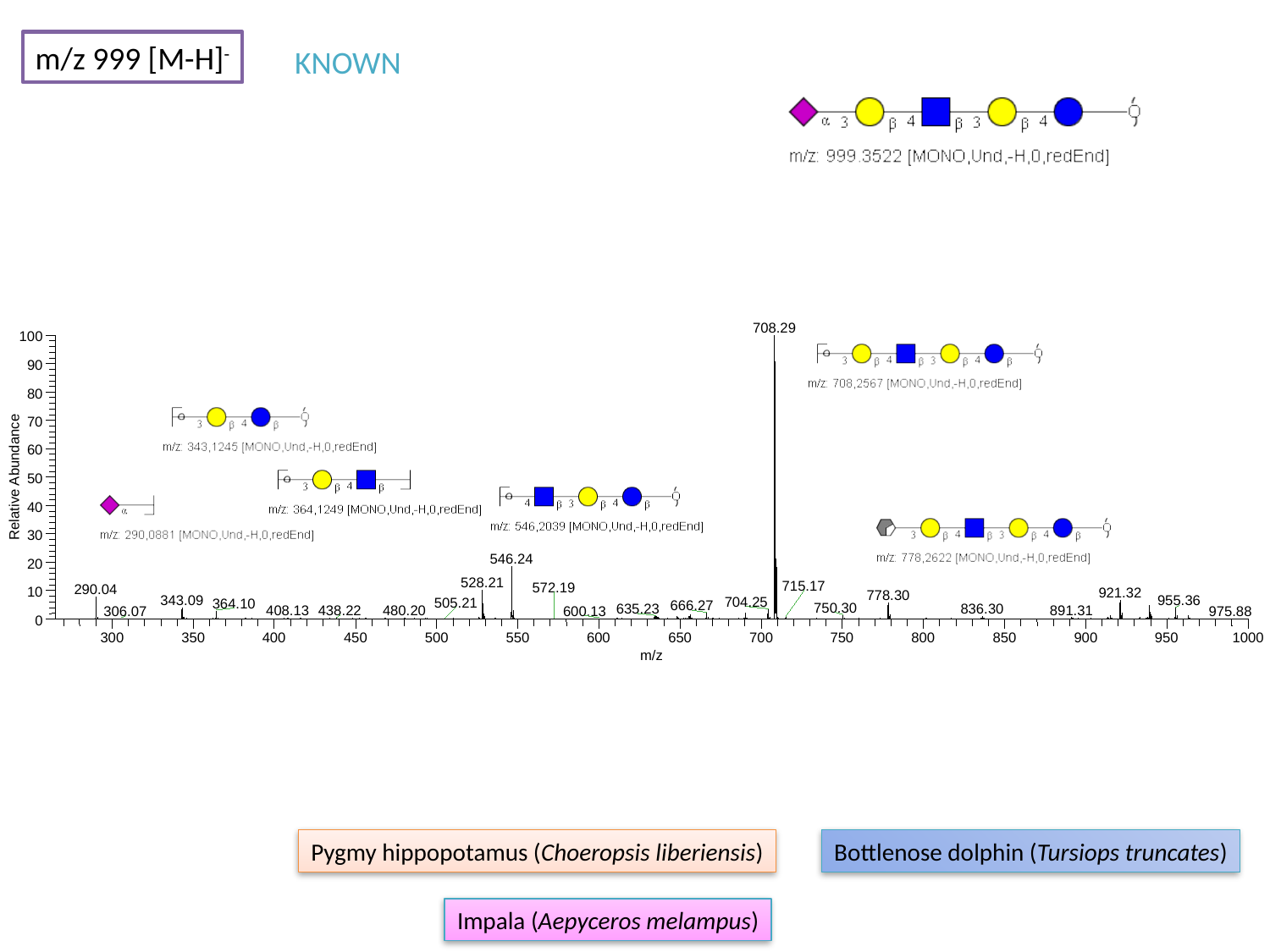

m/z 999 [M-H]^-^
KNOWN
708.29
100
90
80
70
60
Relative Abundance
50
40
30
546.24
20
528.21
715.17
290.04
10
921.32
778.30
343.09
955.36
704.25
364.10
666.27
750.30
836.30
635.23
891.31
480.20
408.13
438.22
306.07
0
300
350
400
450
500
550
600
650
700
750
800
850
900
950
1000
m/z
572.19
505.21
600.13
975.88
Pygmy hippopotamus (Choeropsis liberiensis)
Bottlenose dolphin (Tursiops truncates)
Impala (Aepyceros melampus)

### Slide 77
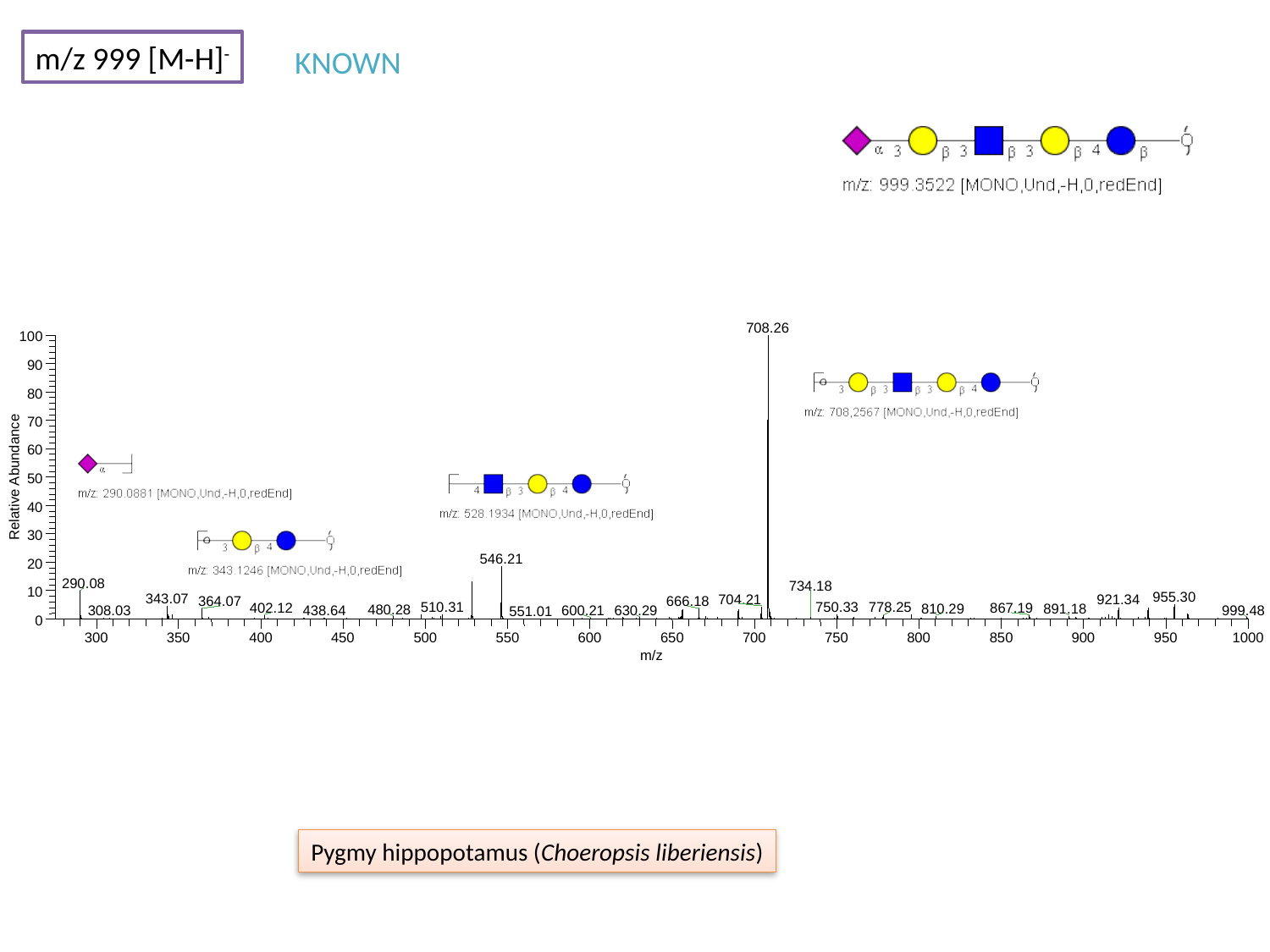

m/z 999 [M-H]^-^
KNOWN
708.26
100
90
80
70
60
Relative Abundance
50
40
30
546.21
20
290.08
734.18
10
955.30
343.07
704.21
921.34
364.07
666.18
510.31
778.25
750.33
402.12
867.19
810.29
891.18
480.28
0
300
350
400
450
500
550
600
650
700
750
800
850
900
950
1000
m/z
630.29
999.48
438.64
600.21
308.03
551.01
Pygmy hippopotamus (Choeropsis liberiensis)

### Slide 78
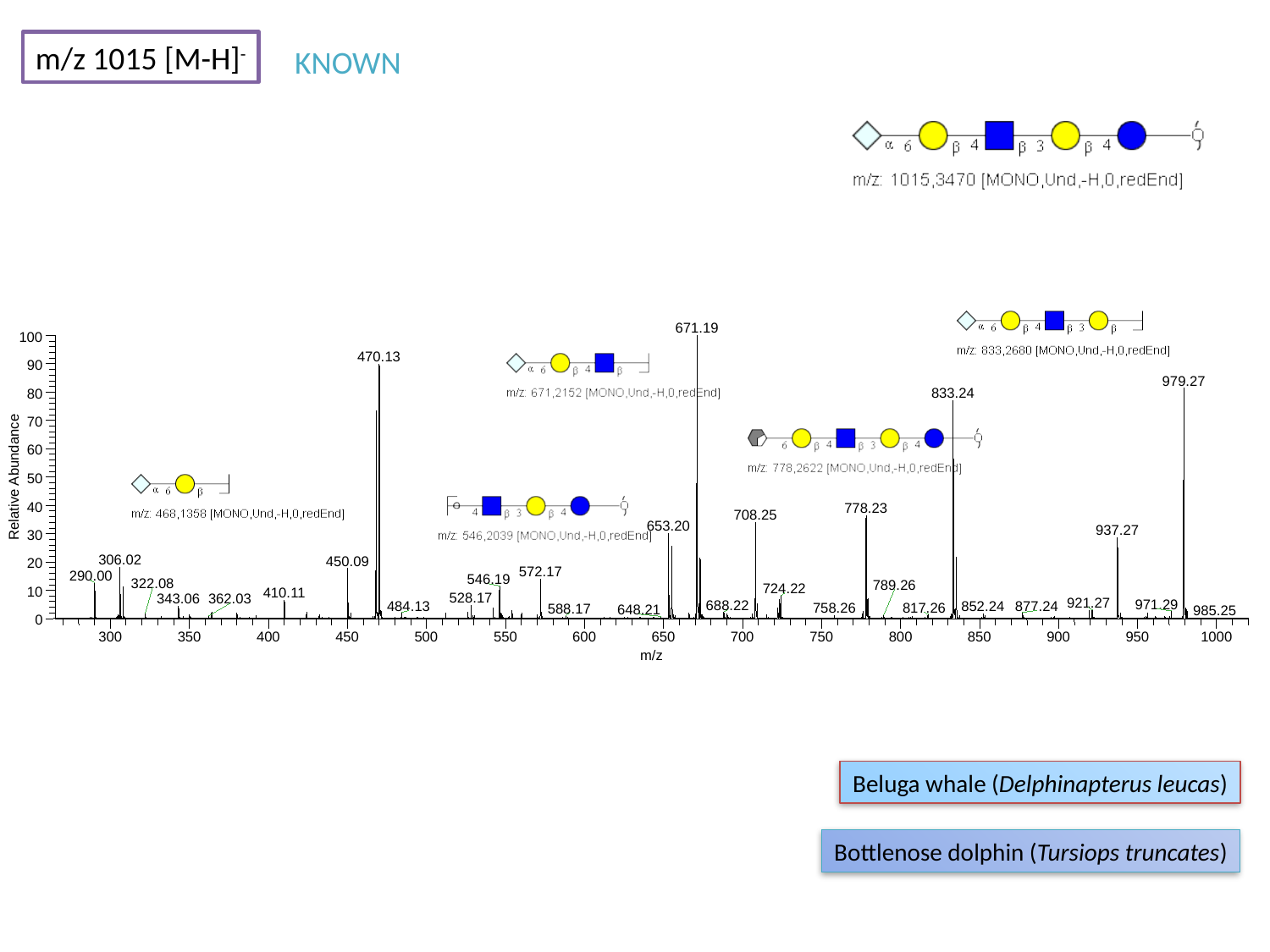

m/z 1015 [M-H]^-^
KNOWN
671.19
100
470.13
90
979.27
833.24
80
70
60
Relative Abundance
50
40
778.23
708.25
653.20
937.27
30
306.02
450.09
20
572.17
290.00
546.19
724.22
10
410.11
528.17
343.06
921.27
971.29
688.22
0
300
350
400
450
500
550
600
650
700
750
800
850
900
950
1000
m/z
322.08
789.26
362.03
484.13
877.24
852.24
817.26
758.26
588.17
648.21
985.25
Beluga whale (Delphinapterus leucas)
Bottlenose dolphin (Tursiops truncates)

### Slide 79
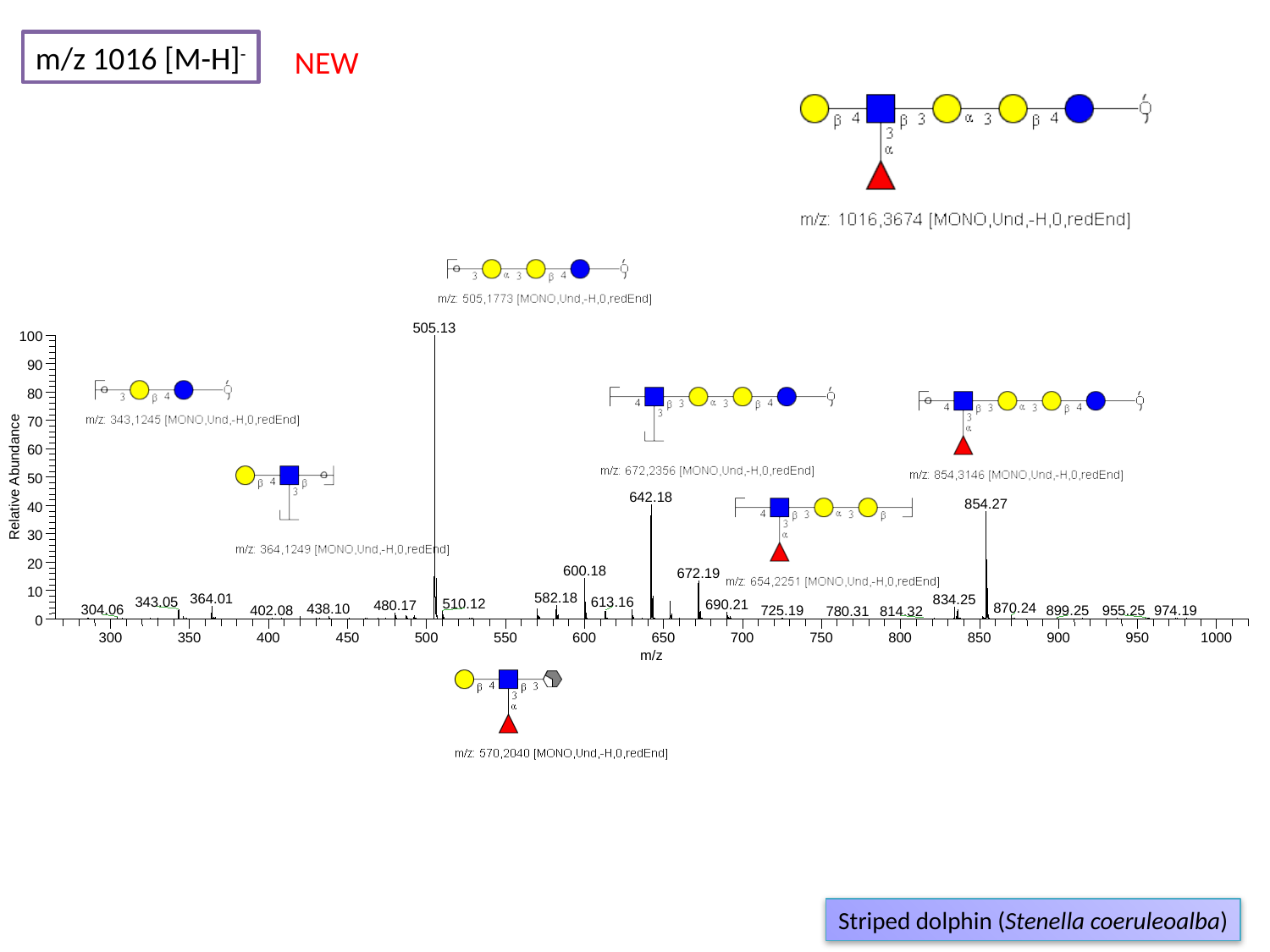

m/z 1016 [M-H]^-^
NEW
505.13
100
90
80
70
60
Relative Abundance
50
642.18
854.27
40
30
20
600.18
672.19
10
582.18
364.01
834.25
343.05
613.16
510.12
690.21
480.17
870.24
438.10
304.06
402.08
974.19
725.19
955.25
0
300
350
400
450
500
550
600
650
700
750
800
850
900
950
1000
m/z
899.25
780.31
814.32
Striped dolphin (Stenella coeruleoalba)

### Slide 80
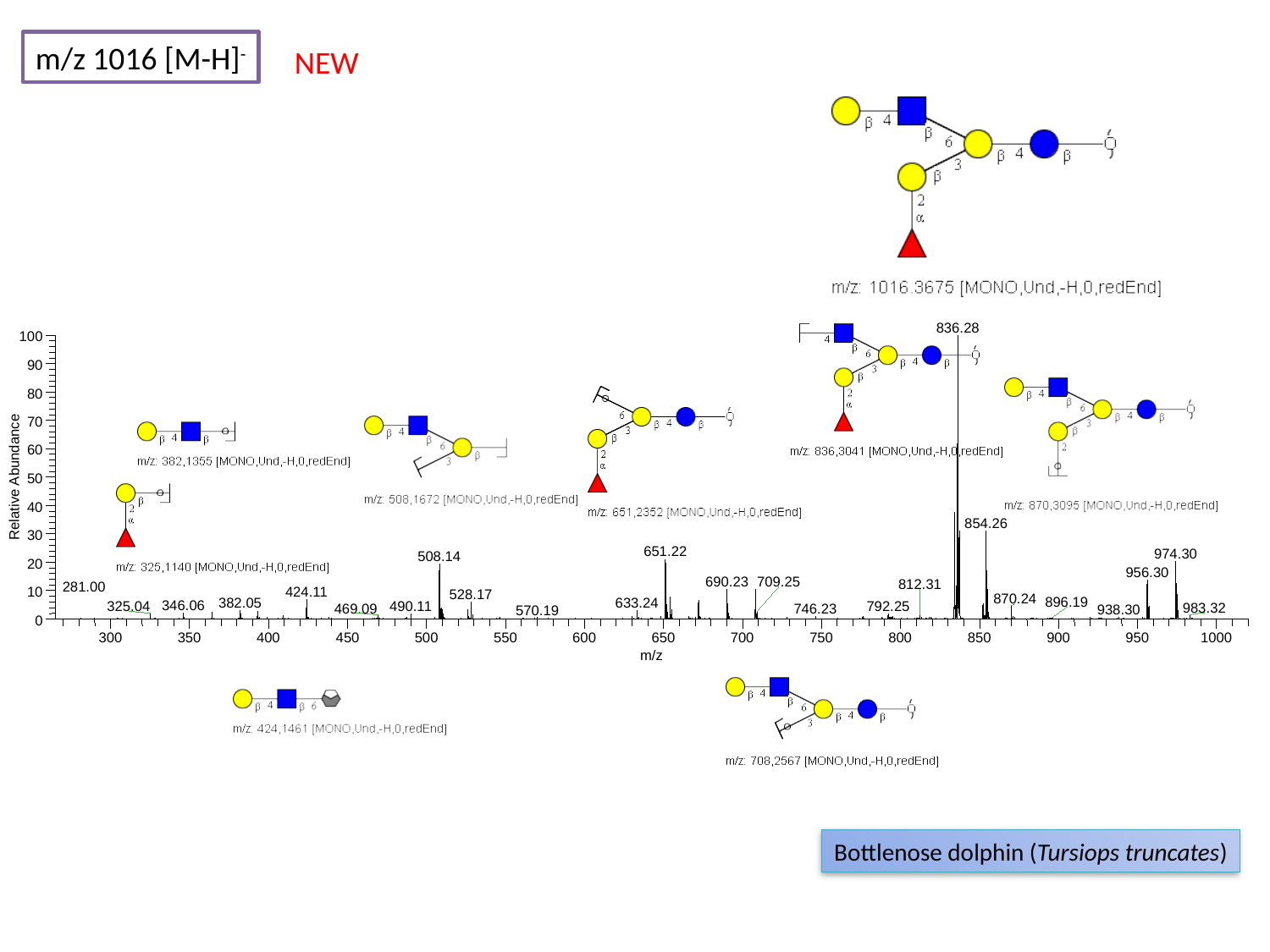

m/z 1016 [M-H]^-^
NEW
836.28
100
90
80
70
60
Relative Abundance
50
40
854.26
30
651.22
974.30
508.14
20
956.30
690.23
709.25
812.31
10
424.11
528.17
870.24
382.05
633.24
346.06
490.11
325.04
792.25
469.09
746.23
938.30
0
300
350
400
450
500
550
600
650
700
750
800
850
900
950
1000
m/z
281.00
896.19
983.32
570.19
Bottlenose dolphin (Tursiops truncates)

### Slide 81
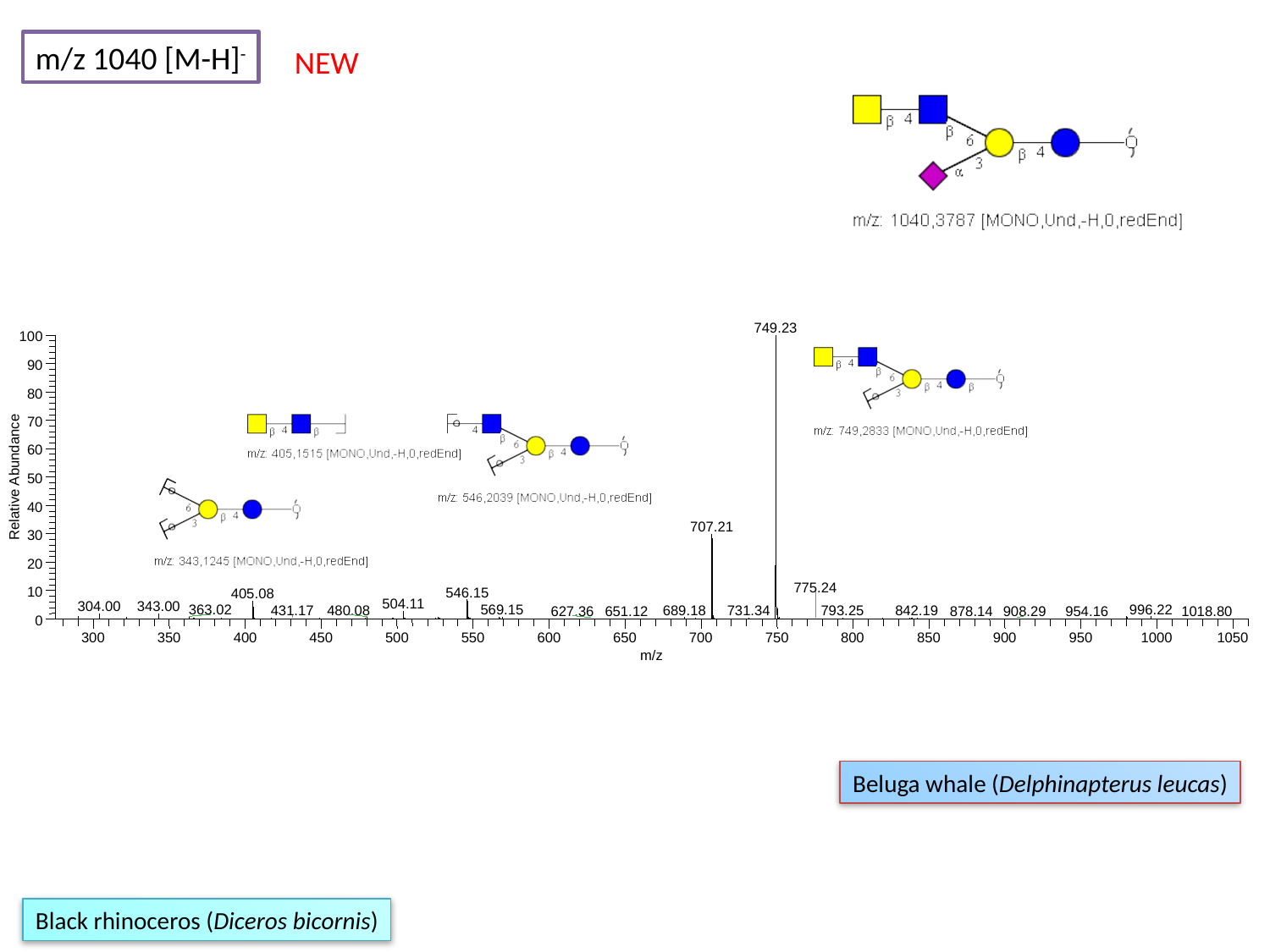

m/z 1040 [M-H]^-^
NEW
749.23
100
90
80
70
60
Relative Abundance
50
40
707.21
30
20
775.24
10
546.15
405.08
504.11
304.00
343.00
996.22
569.15
363.02
480.08
689.18
793.25
431.17
731.34
842.19
878.14
0
300
350
400
450
500
550
600
650
700
750
800
850
900
950
1000
1050
m/z
627.36
651.12
908.29
954.16
1018.80
Beluga whale (Delphinapterus leucas)
Black rhinoceros (Diceros bicornis)

### Slide 82
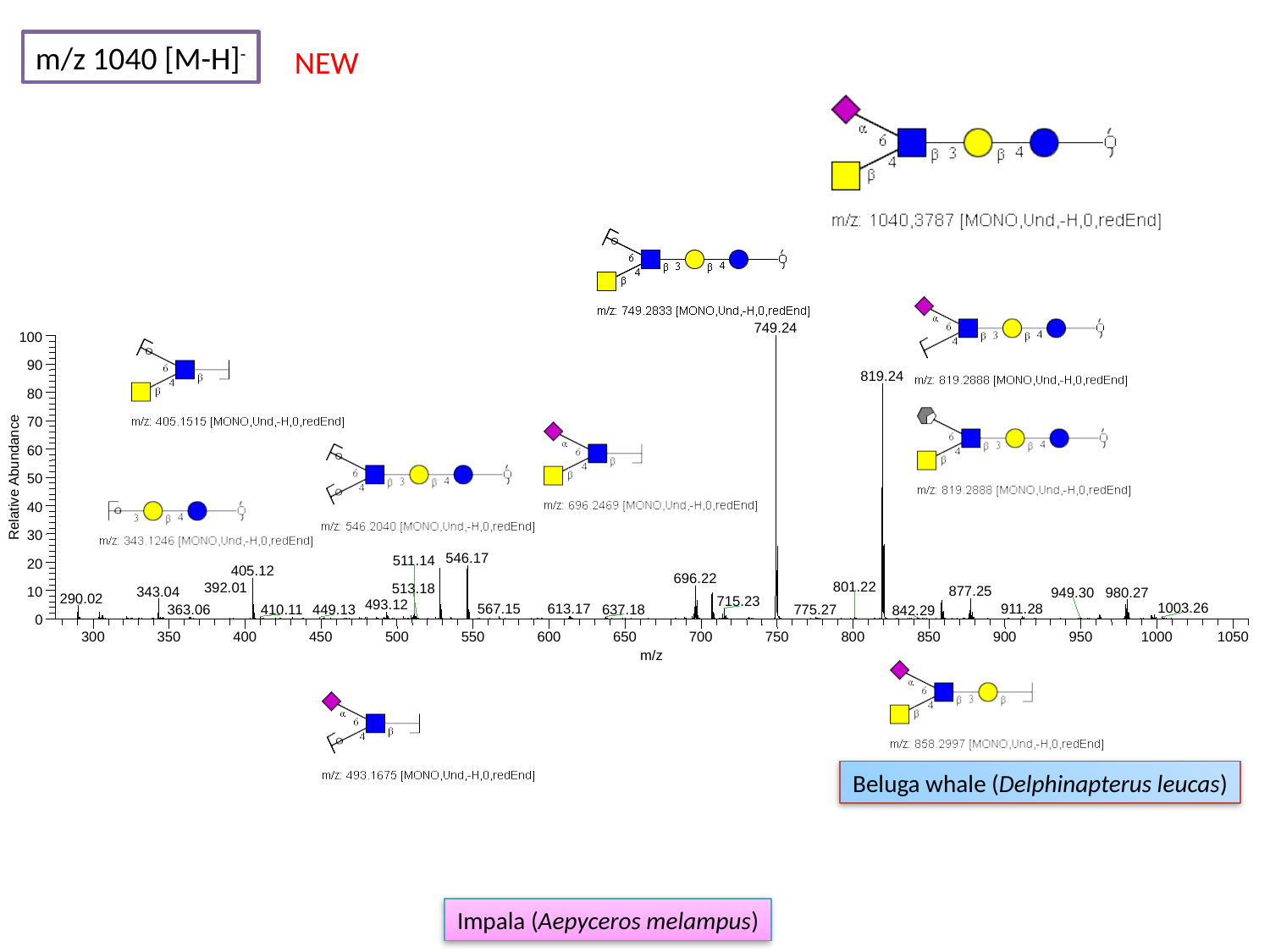

m/z 1040 [M-H]^-^
NEW
749.24
100
90
819.24
80
70
60
Relative Abundance
50
40
30
546.17
511.14
20
405.12
696.22
877.25
343.04
10
980.27
290.02
715.23
493.12
567.15
613.17
911.28
410.11
637.18
775.27
0
300
350
400
450
500
550
600
650
700
750
800
850
900
950
1000
1050
m/z
801.22
392.01
513.18
949.30
1003.26
449.13
363.06
842.29
Beluga whale (Delphinapterus leucas)
Impala (Aepyceros melampus)

### Slide 83
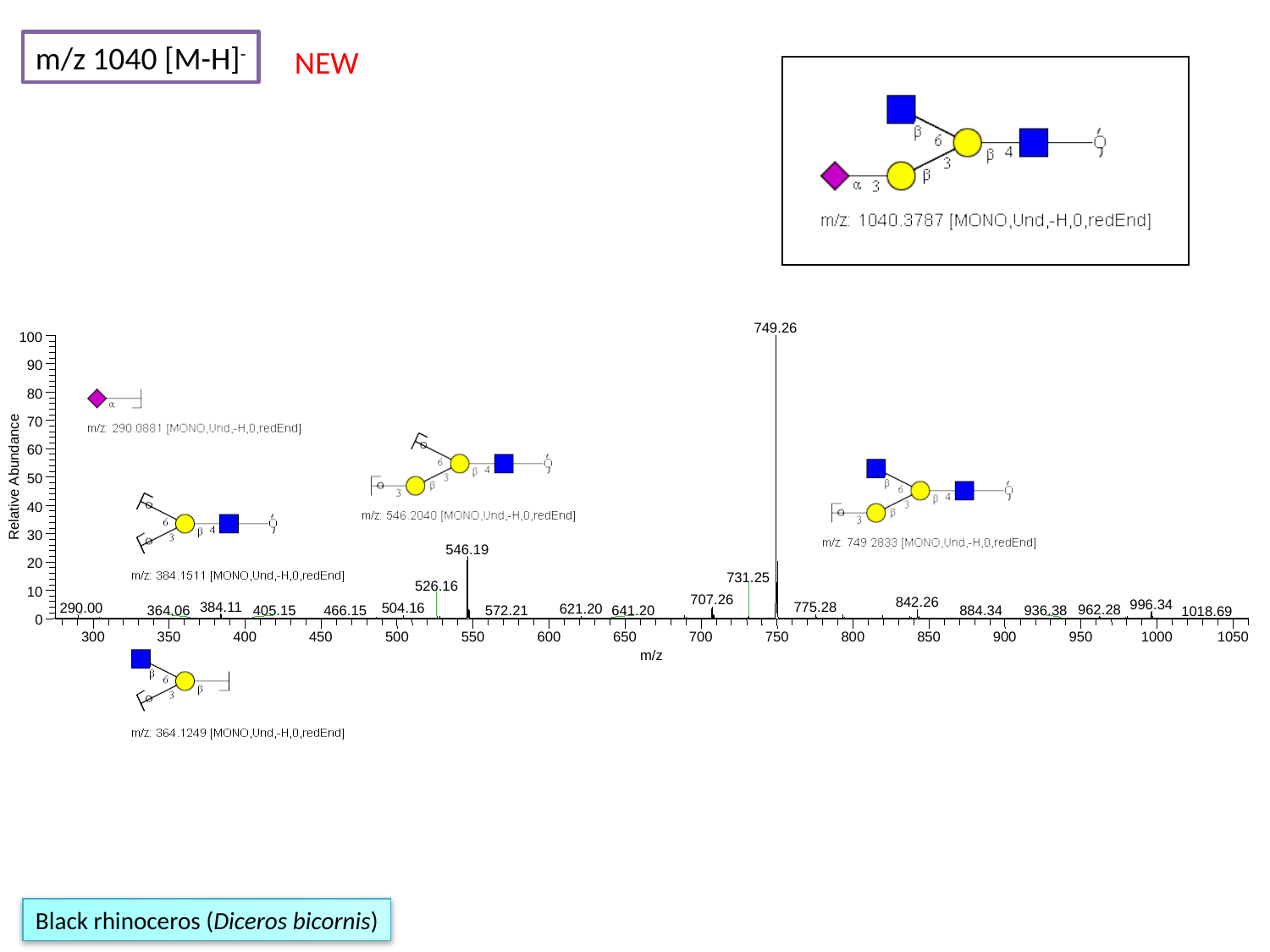

m/z 1040 [M-H]^-^
NEW
749.26
100
90
80
70
60
Relative Abundance
50
40
30
546.19
20
731.25
526.16
10
707.26
842.26
996.34
384.11
775.28
290.00
504.16
621.20
962.28
405.15
466.15
572.21
936.38
0
300
350
400
450
500
550
600
650
700
750
800
850
900
950
1000
1050
m/z
364.06
641.20
884.34
1018.69
Black rhinoceros (Diceros bicornis)

### Slide 84
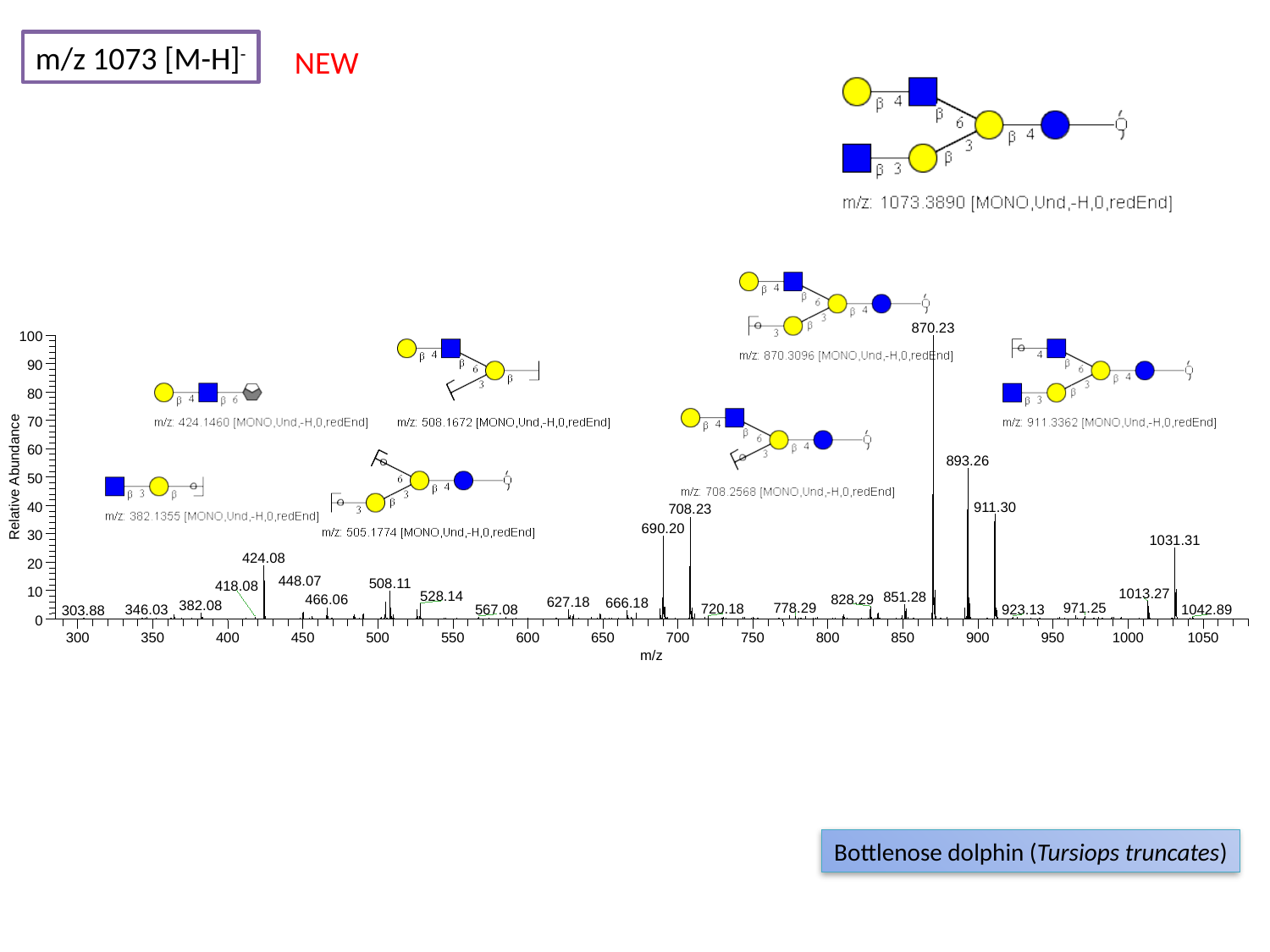

m/z 1073 [M-H]^-^
NEW
870.23
100
90
80
70
60
893.26
Relative Abundance
50
911.30
40
708.23
690.20
30
1031.31
424.08
20
508.11
10
1013.27
528.14
851.28
828.29
466.06
627.18
666.18
382.08
971.25
0
300
350
400
450
500
550
600
650
700
750
800
850
900
950
1000
1050
m/z
448.07
418.08
778.29
720.18
346.03
567.08
923.13
1042.89
303.88
Bottlenose dolphin (Tursiops truncates)

### Slide 85
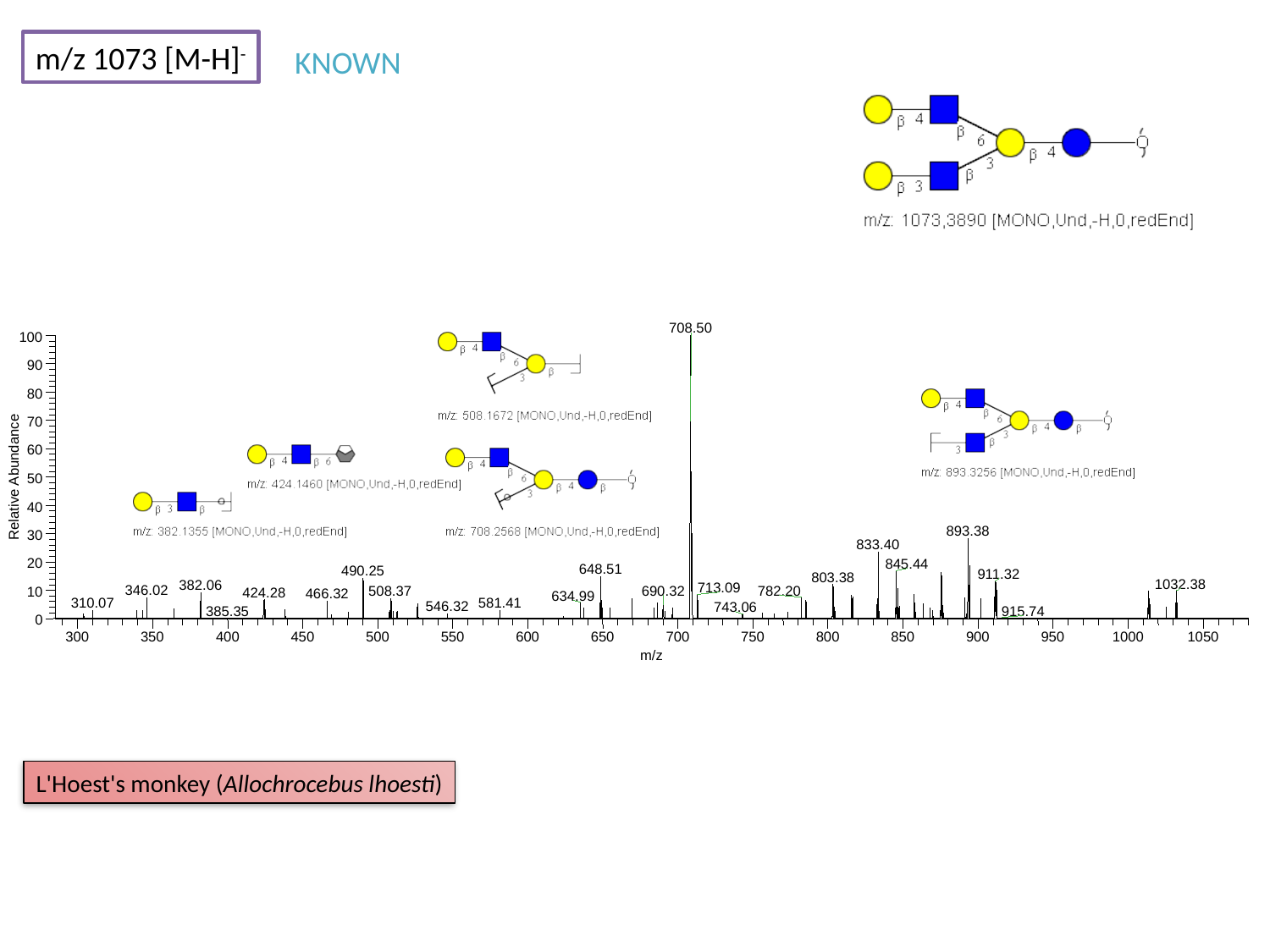

m/z 1073 [M-H]^-^
KNOWN
708.50
100
90
80
70
60
Relative Abundance
50
40
893.38
30
833.40
20
845.44
648.51
490.25
911.32
803.38
1032.38
382.06
713.09
782.20
690.32
10
424.28
0
300
350
400
450
500
550
600
650
700
750
800
850
900
950
1000
1050
m/z
346.02
508.37
466.32
634.99
310.07
581.41
546.32
743.06
385.35
915.74
L'Hoest's monkey (Allochrocebus lhoesti)

### Slide 86
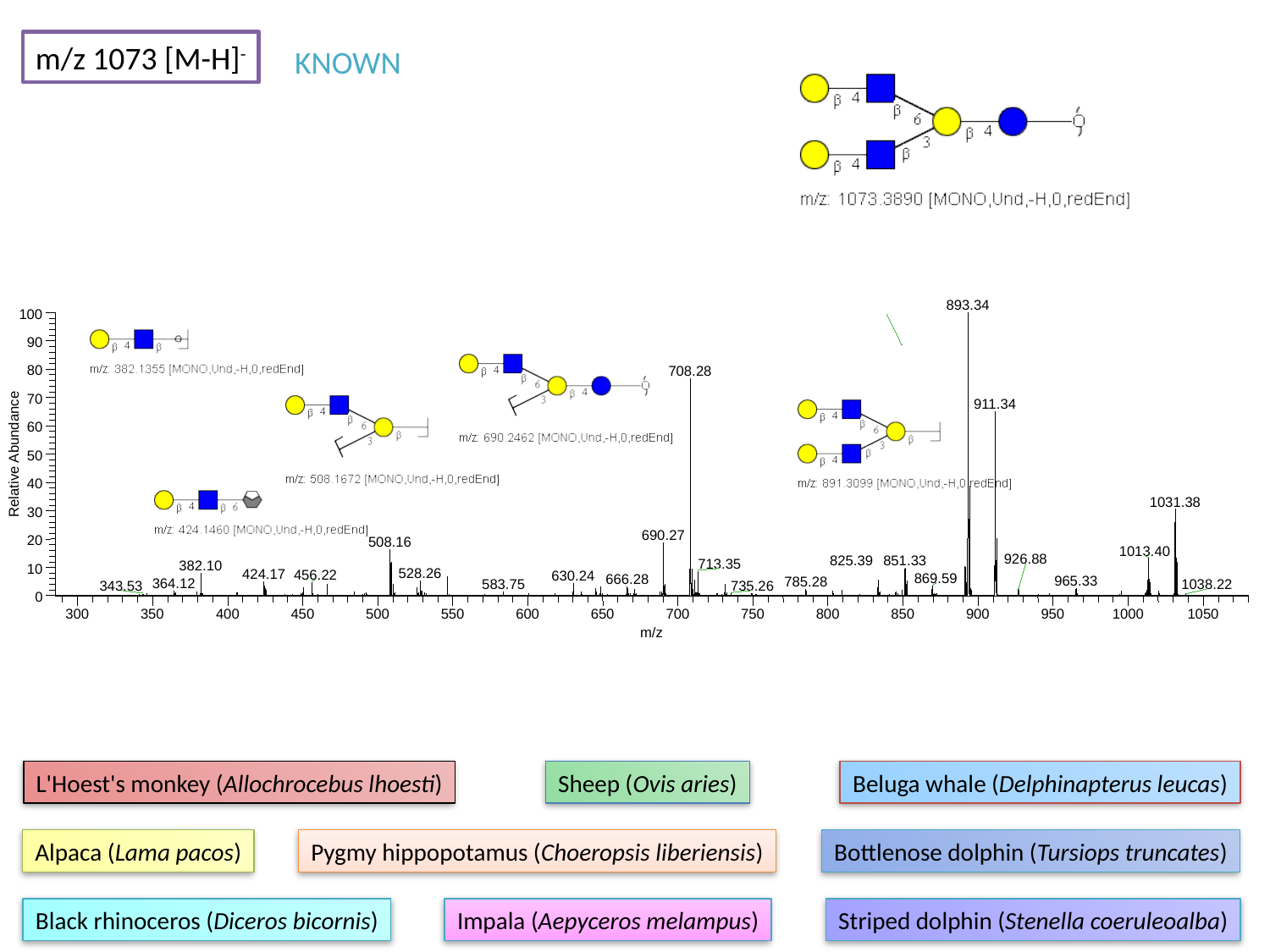

m/z 1073 [M-H]^-^
KNOWN
893.34
100
90
80
708.28
70
911.34
60
Relative Abundance
50
40
1031.38
30
690.27
20
508.16
1013.40
851.33
713.35
382.10
10
528.26
424.17
456.22
630.24
869.59
666.28
965.33
0
300
350
400
450
500
550
600
650
700
750
800
850
900
950
1000
1050
m/z
926.88
825.39
785.28
364.12
583.75
1038.22
343.53
735.26
L'Hoest's monkey (Allochrocebus lhoesti)
Sheep (Ovis aries)
Beluga whale (Delphinapterus leucas)
Alpaca (Lama pacos)
Pygmy hippopotamus (Choeropsis liberiensis)
Bottlenose dolphin (Tursiops truncates)
Black rhinoceros (Diceros bicornis)
Impala (Aepyceros melampus)
Striped dolphin (Stenella coeruleoalba)

### Slide 87
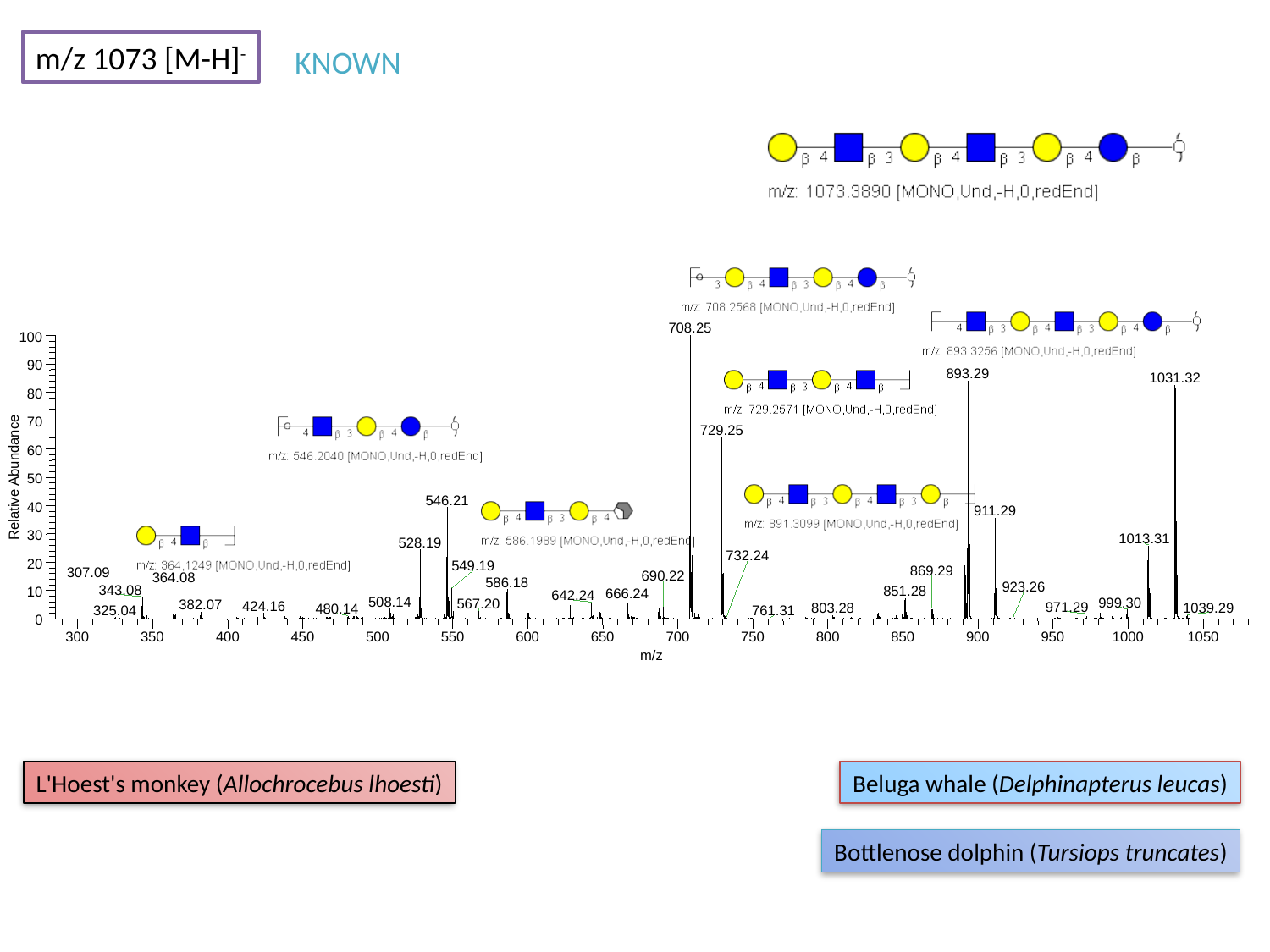

m/z 1073 [M-H]^-^
KNOWN
708.25
100
90
893.29
1031.32
80
70
729.25
60
Relative Abundance
50
546.21
40
911.29
30
1013.31
528.19
20
549.19
690.22
364.08
586.18
343.08
851.28
10
666.24
642.24
0
300
350
400
450
500
550
600
650
700
750
800
850
900
950
1000
1050
m/z
732.24
869.29
307.09
923.26
508.14
999.30
567.20
382.07
424.16
971.29
803.28
1039.29
480.14
325.04
761.31
L'Hoest's monkey (Allochrocebus lhoesti)
Beluga whale (Delphinapterus leucas)
Bottlenose dolphin (Tursiops truncates)

### Slide 88
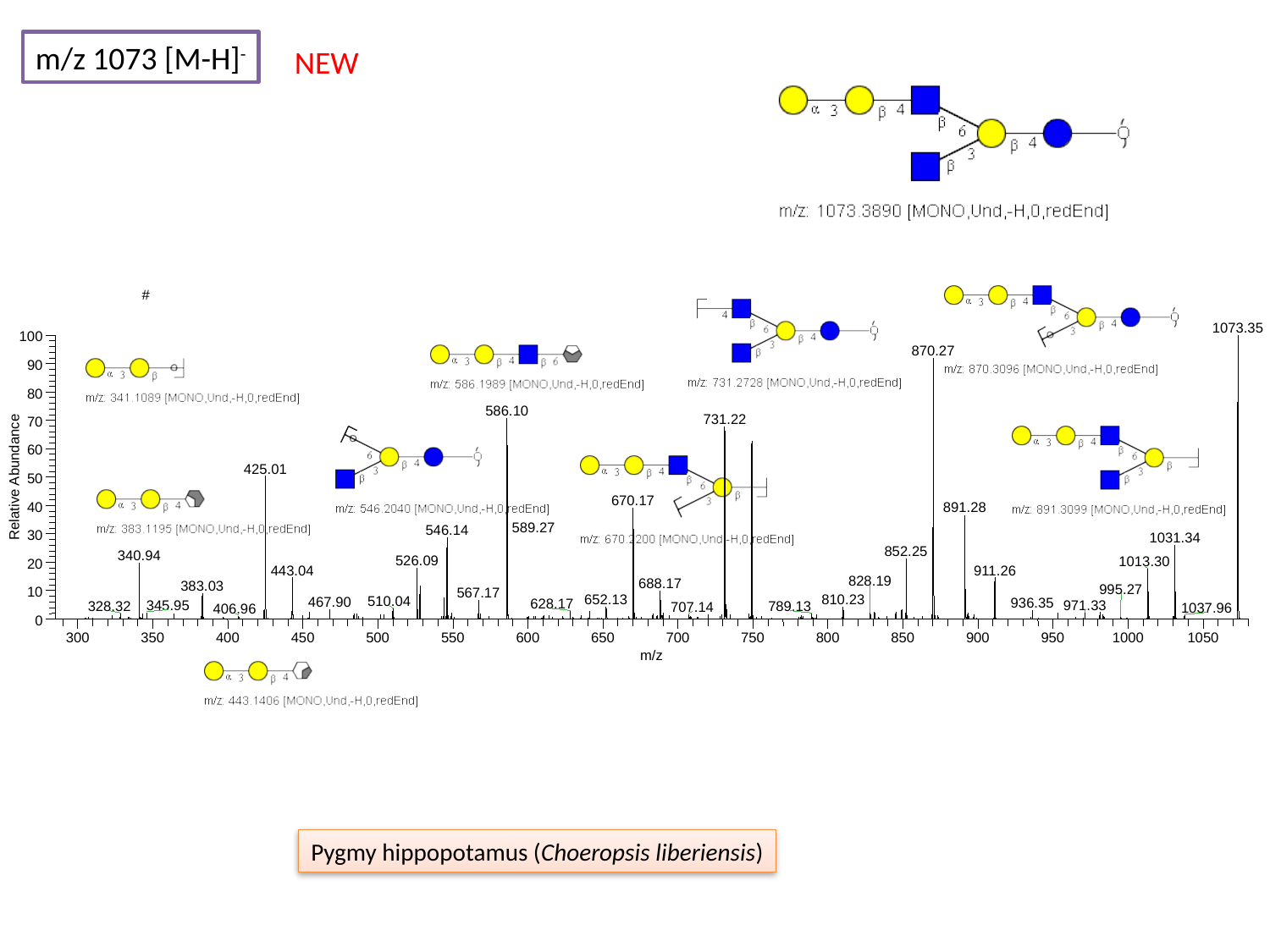

m/z 1073 [M-H]^-^
NEW
#
1073.35
100
870.27
90
80
586.10
731.22
70
60
425.01
Relative Abundance
50
670.17
40
891.28
546.14
30
1031.34
852.25
340.94
526.09
1013.30
20
443.04
911.26
688.17
383.03
10
567.17
810.23
652.13
0
300
350
400
450
500
550
600
650
700
750
800
850
900
950
1000
1050
m/z
589.27
828.19
995.27
510.04
467.90
936.35
628.17
971.33
345.95
328.32
789.13
707.14
1037.96
406.96
Pygmy hippopotamus (Choeropsis liberiensis)

### Slide 89
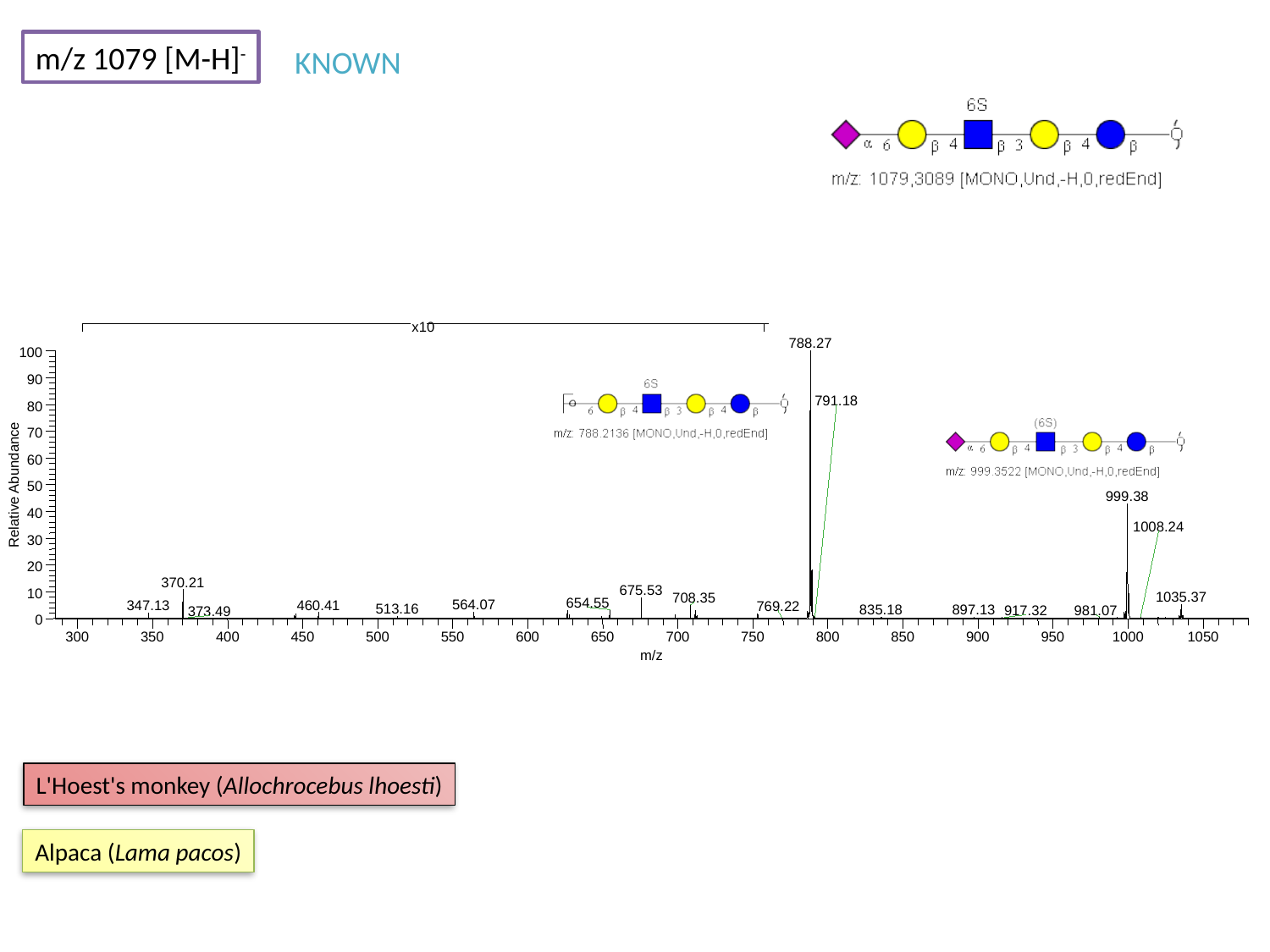

m/z 1079 [M-H]^-^
KNOWN
788.27
100
90
791.18
80
70
60
Relative Abundance
50
999.38
40
1008.24
30
20
370.21
675.53
10
1035.37
708.35
654.55
564.07
347.13
769.22
835.18
897.13
981.07
0
300
350
400
450
500
550
600
650
700
750
800
850
900
950
1000
1050
m/z
x10
460.41
513.16
917.32
373.49
L'Hoest's monkey (Allochrocebus lhoesti)
Alpaca (Lama pacos)

### Slide 90
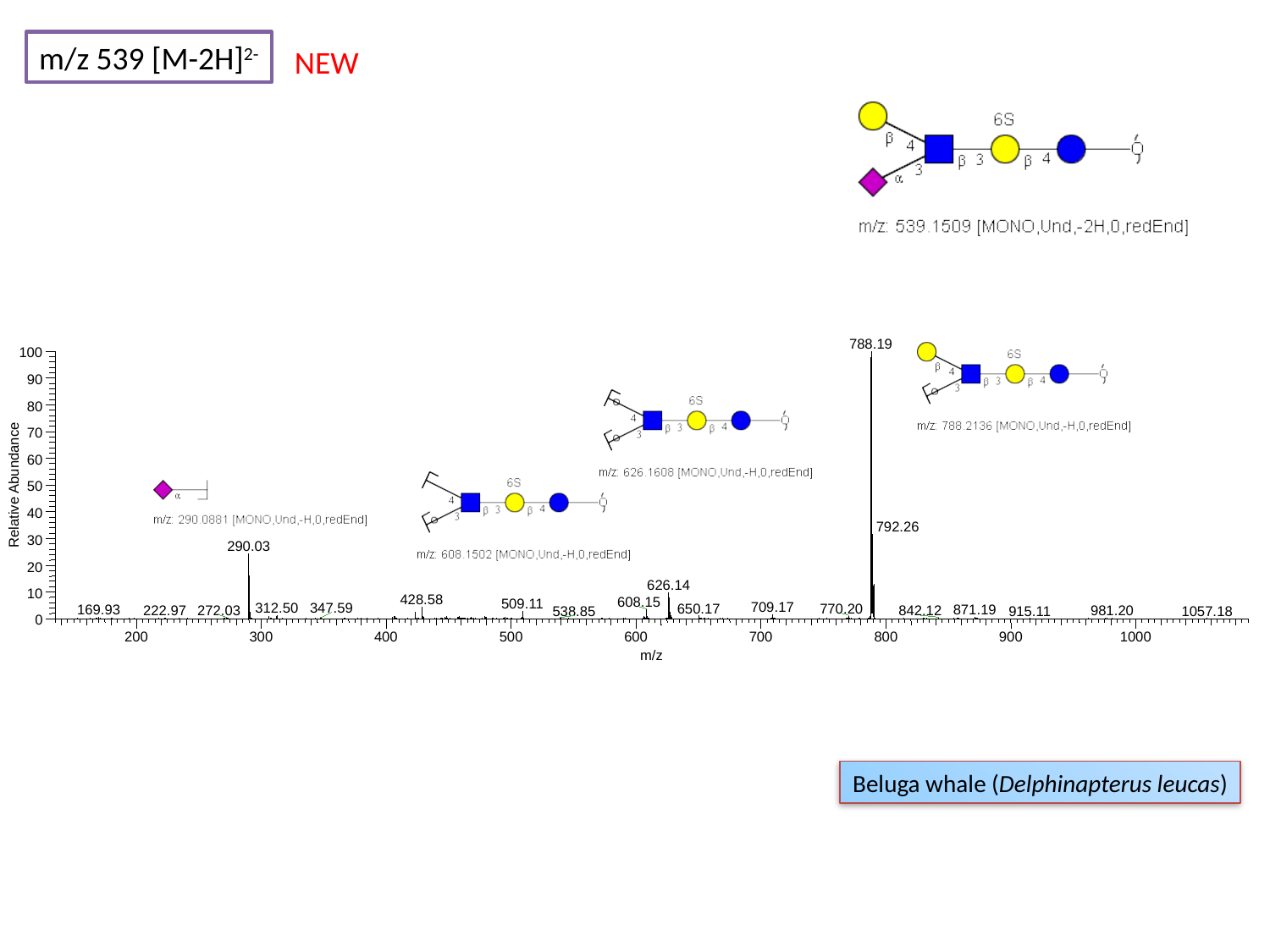

m/z 539 [M-2H]^2-^
NEW
788.19
100
90
80
70
60
Relative Abundance
50
40
792.26
30
290.03
20
626.14
10
428.58
608.15
509.11
709.17
312.50
347.59
650.17
770.20
169.93
871.19
272.03
222.97
842.12
981.20
538.85
915.11
1057.18
0
200
300
400
500
600
700
800
900
1000
m/z
Beluga whale (Delphinapterus leucas)

### Slide 91
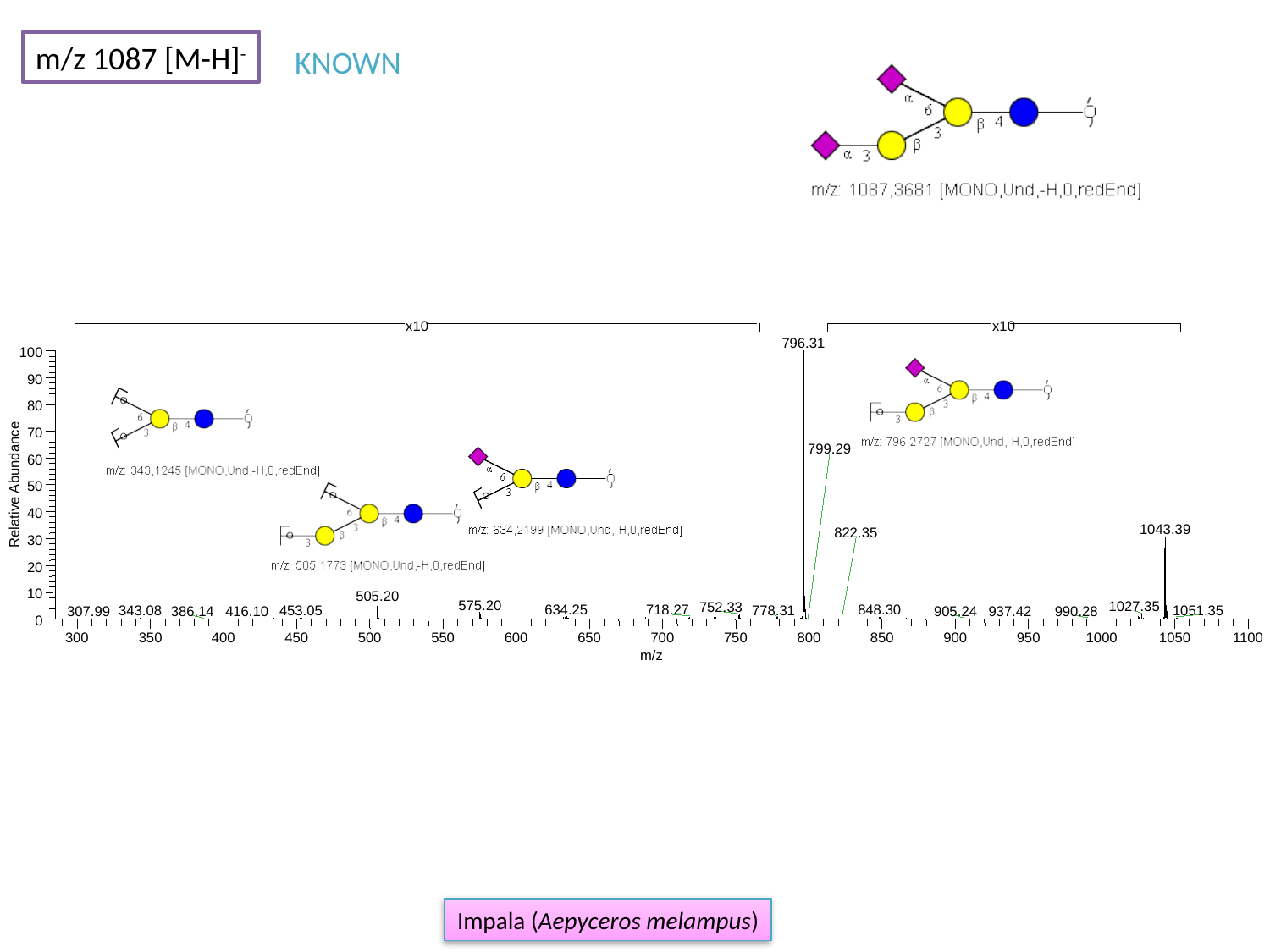

m/z 1087 [M-H]^-^
KNOWN
796.31
100
90
80
70
799.29
60
Relative Abundance
50
40
1043.39
30
20
10
505.20
575.20
1027.35
752.33
634.25
718.27
848.30
778.31
1051.35
453.05
0
300
350
400
450
500
550
600
650
700
750
800
850
900
950
1000
1050
1100
m/z
x10
x10
822.35
343.08
386.14
416.10
905.24
937.42
990.28
307.99
Impala (Aepyceros melampus)

### Slide 92
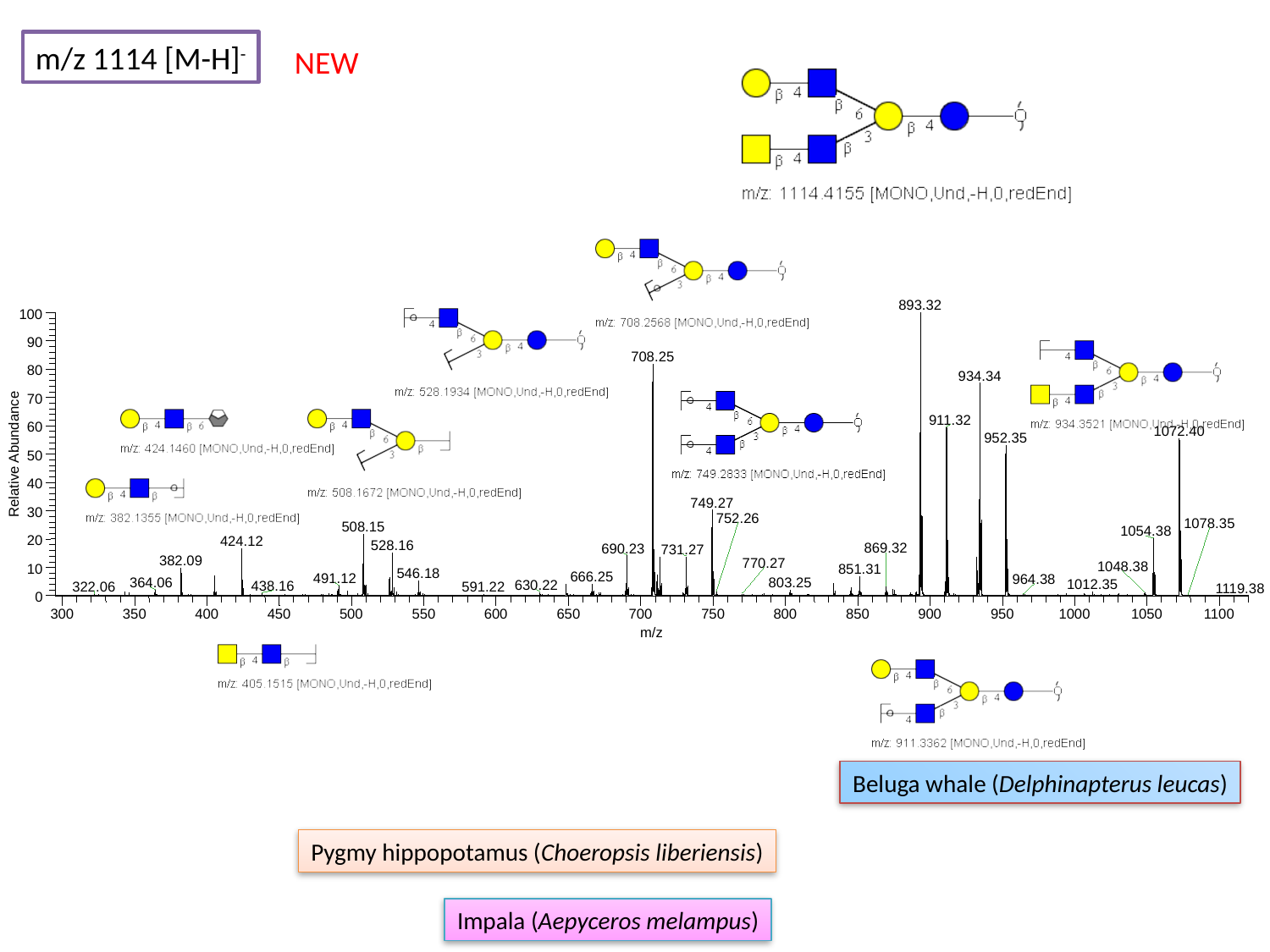

m/z 1114 [M-H]^-^
NEW
893.32
100
90
708.25
80
934.34
70
911.32
60
1072.40
952.35
Relative Abundance
50
40
749.27
30
508.15
1054.38
20
424.12
528.16
690.23
731.27
10
0
300
350
400
450
500
550
600
650
700
750
800
850
900
950
1000
1050
1100
m/z
752.26
1078.35
869.32
382.09
770.27
1048.38
851.31
546.18
666.25
491.12
964.38
364.06
803.25
1012.35
630.22
438.16
322.06
591.22
1119.38
Beluga whale (Delphinapterus leucas)
Pygmy hippopotamus (Choeropsis liberiensis)
Impala (Aepyceros melampus)

### Slide 93
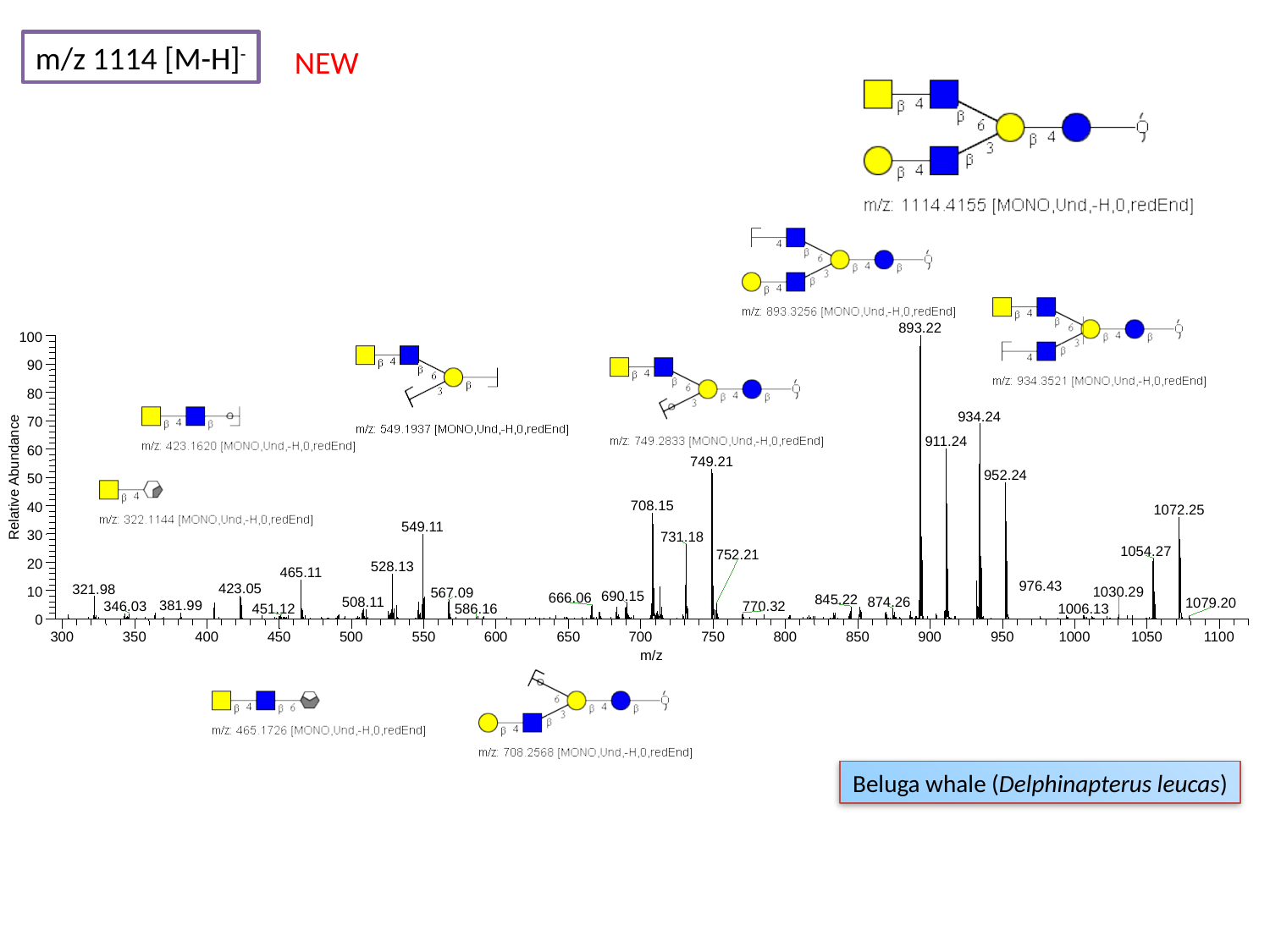

m/z 1114 [M-H]^-^
NEW
893.22
100
90
80
934.24
70
911.24
60
749.21
952.24
Relative Abundance
50
708.15
40
1072.25
549.11
30
731.18
1054.27
20
528.13
465.11
423.05
321.98
10
567.09
0
300
350
400
450
500
550
600
650
700
750
800
850
900
950
1000
1050
1100
m/z
752.21
976.43
1030.29
690.15
666.06
845.22
508.11
874.26
1079.20
381.99
346.03
770.32
451.12
1006.13
586.16
Beluga whale (Delphinapterus leucas)

### Slide 94
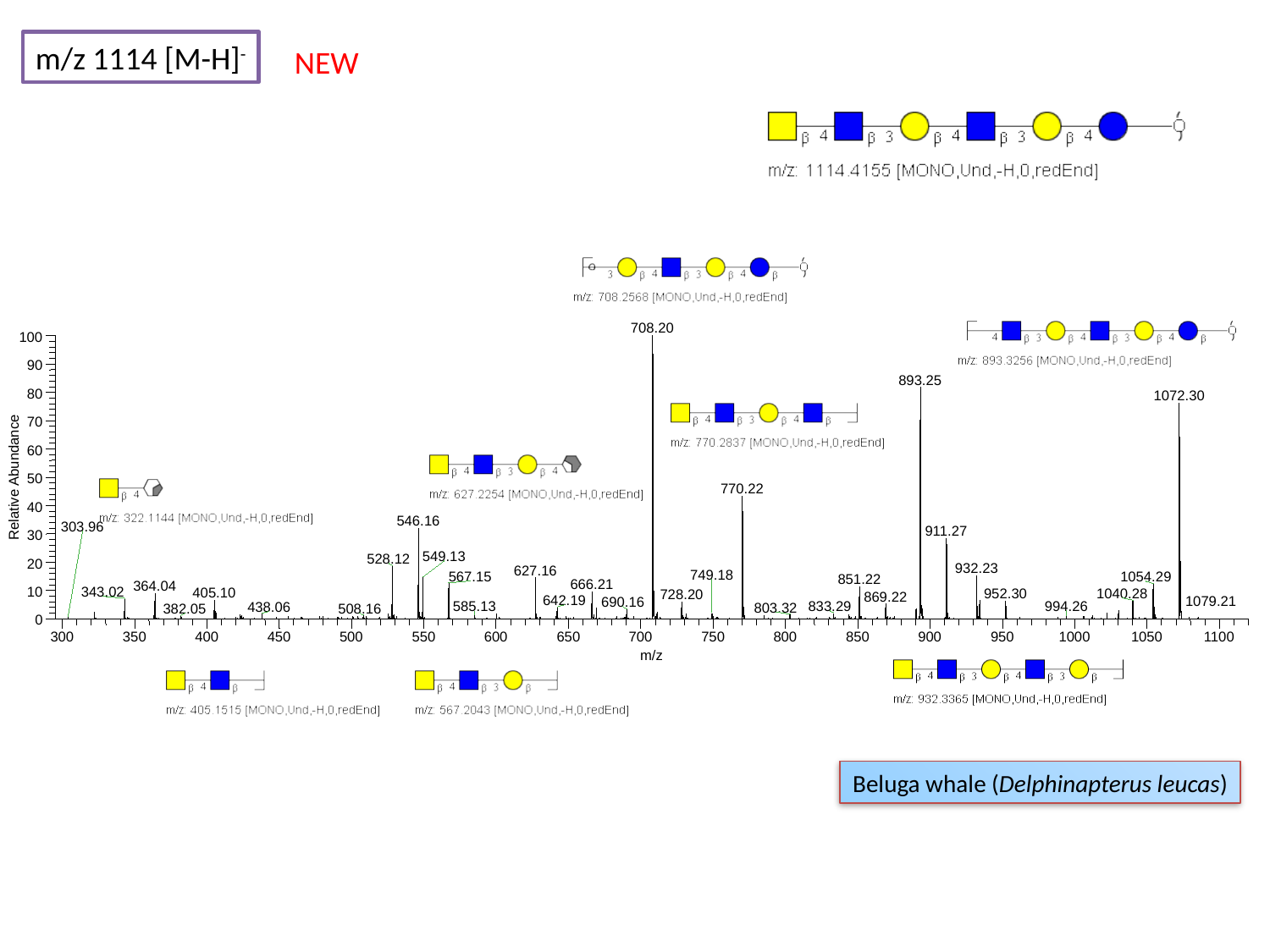

m/z 1114 [M-H]^-^
NEW
708.20
100
90
893.25
80
1072.30
70
60
Relative Abundance
50
770.22
40
546.16
911.27
30
549.13
528.12
20
932.23
627.16
567.15
1054.29
851.22
10
0
300
350
400
450
500
550
600
650
700
750
800
850
900
950
1000
1050
1100
m/z
303.96
749.18
666.21
364.04
343.02
405.10
1040.28
952.30
728.20
869.22
642.19
1079.21
690.16
994.26
833.29
585.13
438.06
803.32
508.16
382.05
Beluga whale (Delphinapterus leucas)

### Slide 95
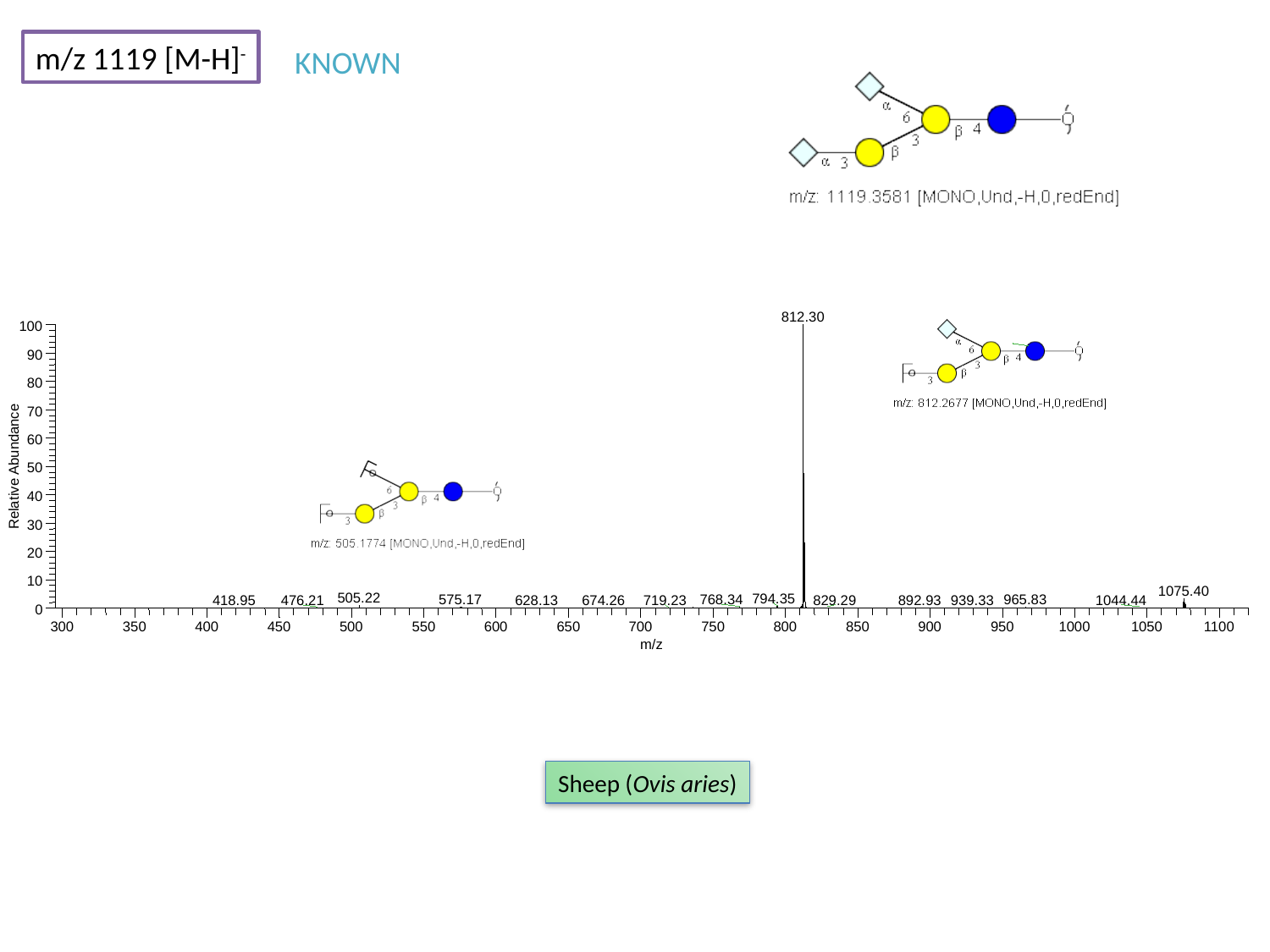

m/z 1119 [M-H]^-^
KNOWN
812.30
100
90
80
70
60
Relative Abundance
50
40
30
20
10
1075.40
505.22
794.35
575.17
768.34
965.83
628.13
674.26
719.23
829.29
1044.44
0
300
350
400
450
500
550
600
650
700
750
800
850
900
950
1000
1050
1100
m/z
892.93
418.95
476.21
939.33
Sheep (Ovis aries)

### Slide 96
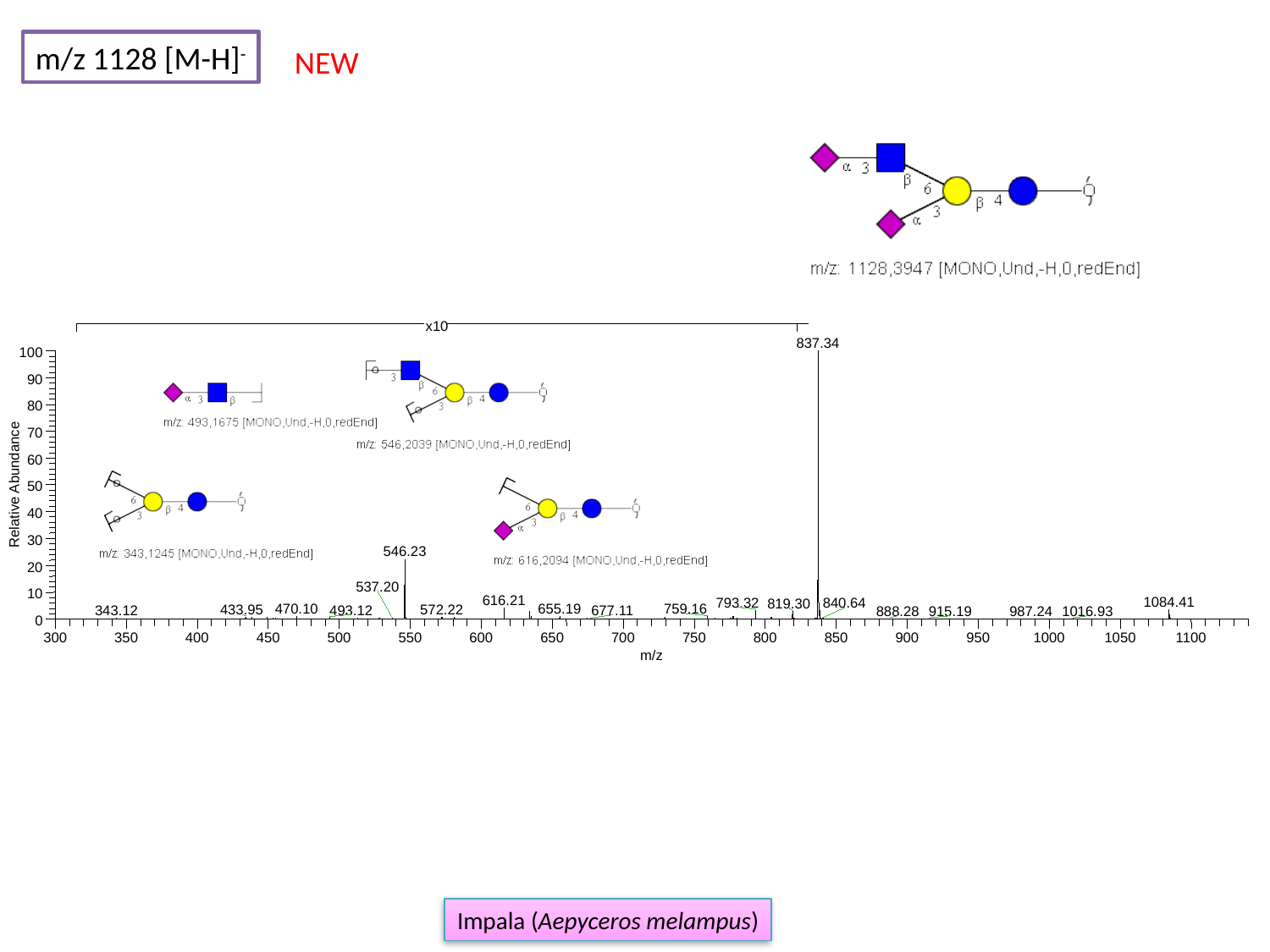

m/z 1128 [M-H]^-^
NEW
837.34
100
90
80
70
60
Relative Abundance
50
40
30
546.23
20
10
616.21
1084.41
793.32
840.64
819.30
655.19
759.16
888.28
0
300
350
400
450
500
550
600
650
700
750
800
850
900
950
1000
1050
1100
m/z
x10
537.20
470.10
433.95
572.22
493.12
677.11
343.12
915.19
987.24
1016.93
Impala (Aepyceros melampus)

### Slide 97
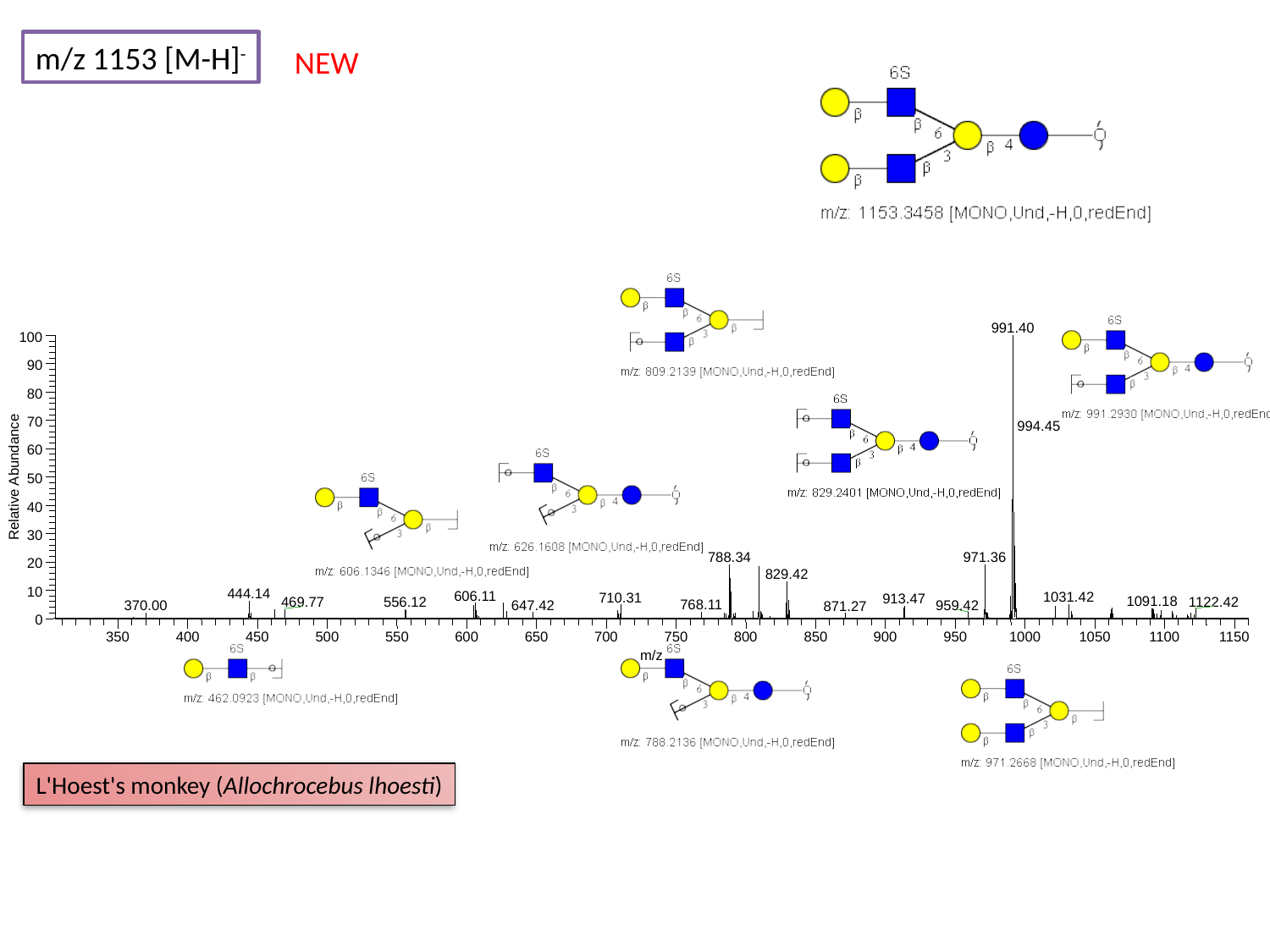

m/z 1153 [M-H]^-^
NEW
991.40
100
90
80
70
60
Relative Abundance
50
40
30
788.34
971.36
20
829.42
10
444.14
606.11
1031.42
710.31
913.47
1091.18
1122.42
469.77
0
350
400
450
500
550
600
650
700
750
800
850
900
950
1000
1050
1100
1150
m/z
994.45
556.12
768.11
647.42
959.42
370.00
871.27
L'Hoest's monkey (Allochrocebus lhoesti)

### Slide 98
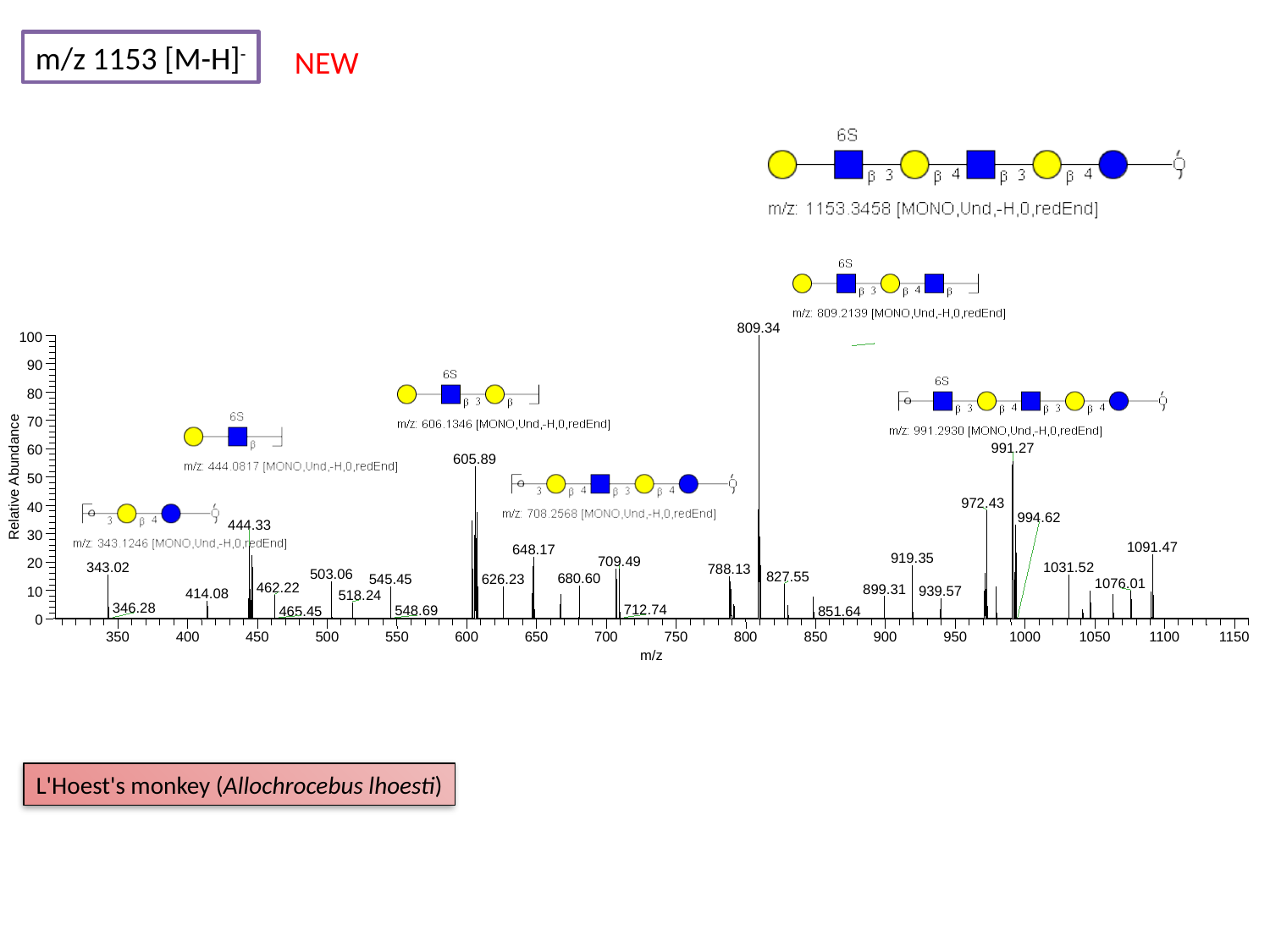

m/z 1153 [M-H]^-^
NEW
809.34
100
90
80
70
991.27
60
605.89
Relative Abundance
50
972.43
40
444.33
30
1091.47
648.17
919.35
709.49
20
788.13
10
0
350
400
450
500
550
600
650
700
750
800
850
900
950
1000
1050
1100
1150
m/z
994.62
1031.52
343.02
503.06
827.55
680.60
626.23
545.45
1076.01
462.22
899.31
939.57
414.08
518.24
346.28
712.74
548.69
465.45
851.64
L'Hoest's monkey (Allochrocebus lhoesti)

### Slide 99
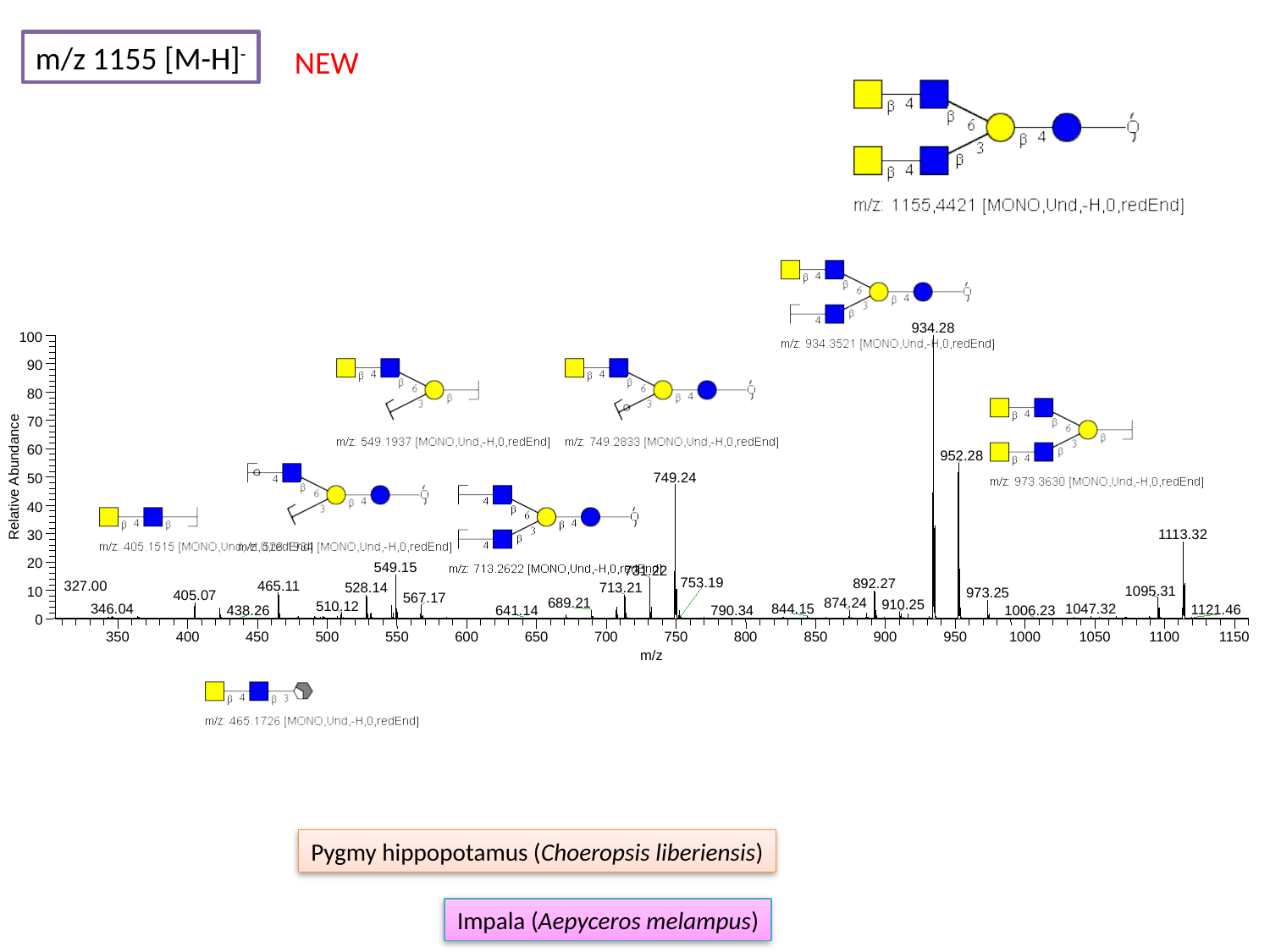

m/z 1155 [M-H]^-^
NEW
934.28
100
90
80
70
60
952.28
Relative Abundance
749.24
50
40
1113.32
30
20
549.15
731.22
892.27
465.11
713.21
1095.31
10
0
350
400
450
500
550
600
650
700
750
800
850
900
950
1000
1050
1100
1150
m/z
753.19
327.00
528.14
973.25
405.07
567.17
689.21
874.24
910.25
510.12
346.04
1047.32
844.15
1121.46
438.26
790.34
641.14
1006.23
Pygmy hippopotamus (Choeropsis liberiensis)
Impala (Aepyceros melampus)

### Slide 100
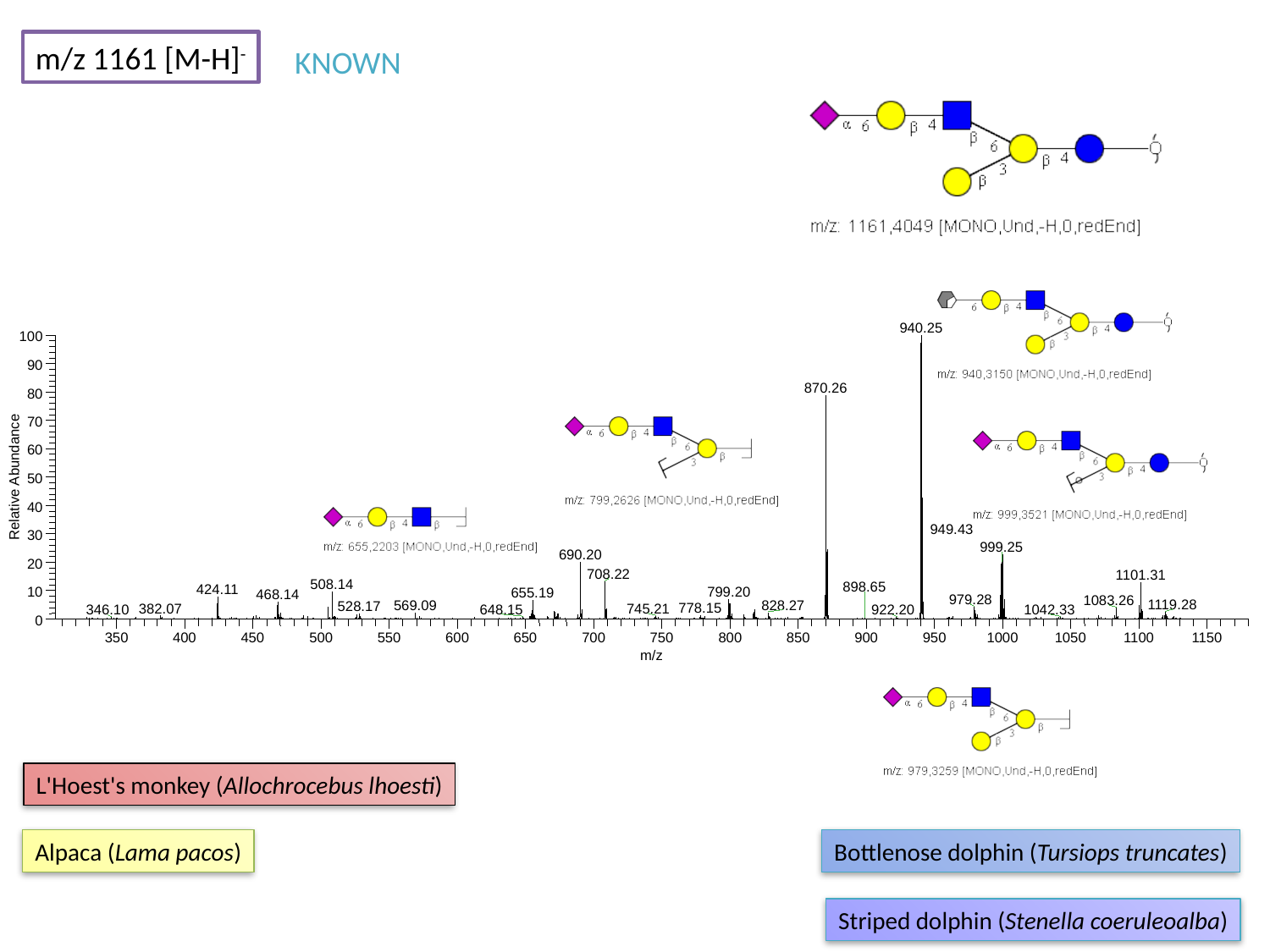

m/z 1161 [M-H]^-^
KNOWN
940.25
100
90
870.26
80
70
60
Relative Abundance
50
40
30
999.25
690.20
20
708.22
1101.31
508.14
424.11
799.20
10
655.19
0
350
400
450
500
550
600
650
700
750
800
850
900
950
1000
1050
1100
1150
m/z
949.43
898.65
468.14
979.28
1083.26
1119.28
828.27
569.09
528.17
778.15
382.07
745.21
648.15
1042.33
346.10
922.20
L'Hoest's monkey (Allochrocebus lhoesti)
Alpaca (Lama pacos)
Bottlenose dolphin (Tursiops truncates)
Striped dolphin (Stenella coeruleoalba)

### Slide 101
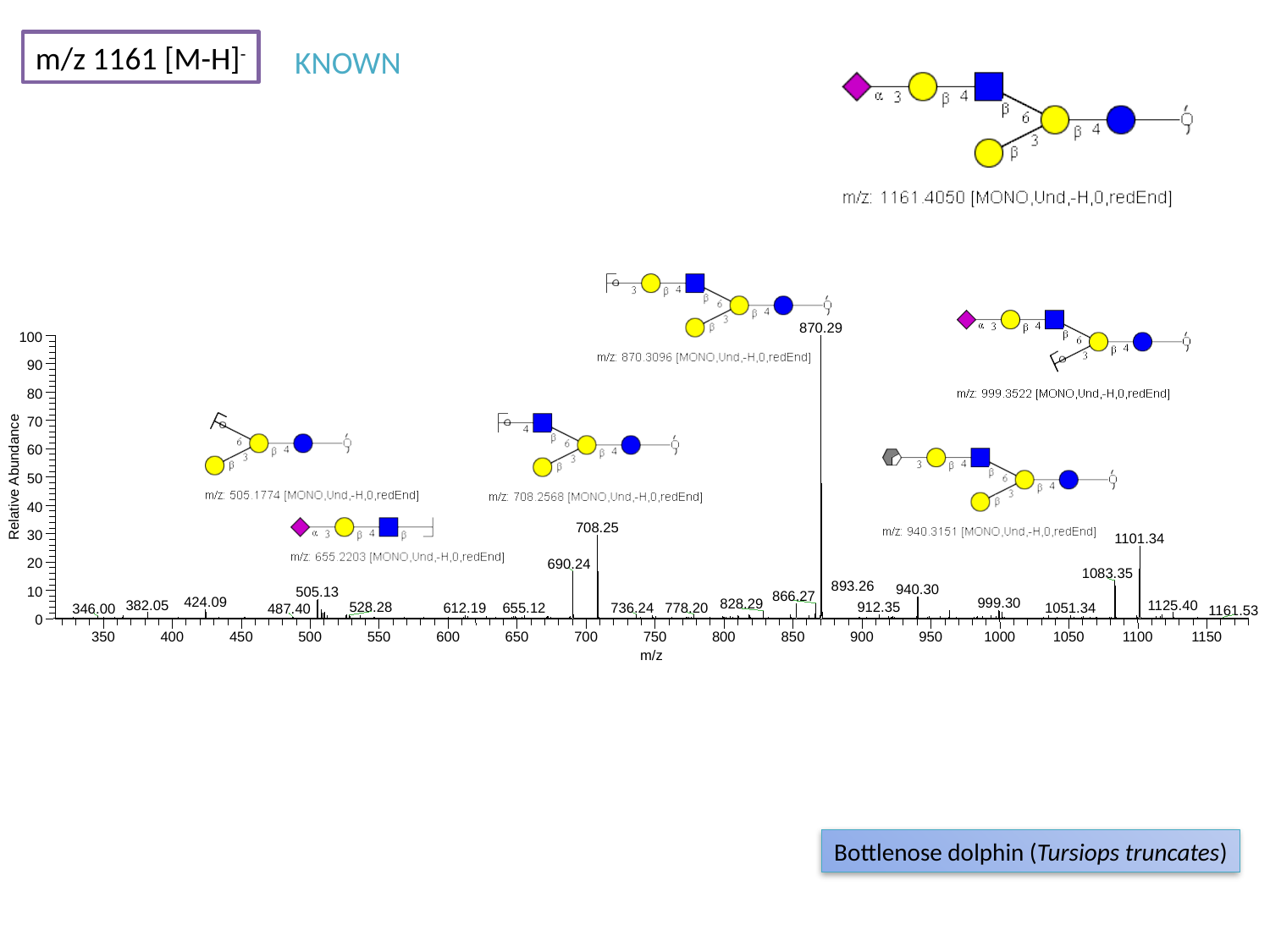

m/z 1161 [M-H]^-^
KNOWN
870.29
100
90
80
70
60
Relative Abundance
50
40
708.25
30
1101.34
20
690.24
1083.35
940.30
10
505.13
866.27
424.09
999.30
0
350
400
450
500
550
600
650
700
750
800
850
900
950
1000
1050
1100
1150
m/z
893.26
828.29
382.05
1125.40
528.28
912.35
736.24
778.20
1051.34
612.19
655.12
346.00
487.40
1161.53
Bottlenose dolphin (Tursiops truncates)

### Slide 102
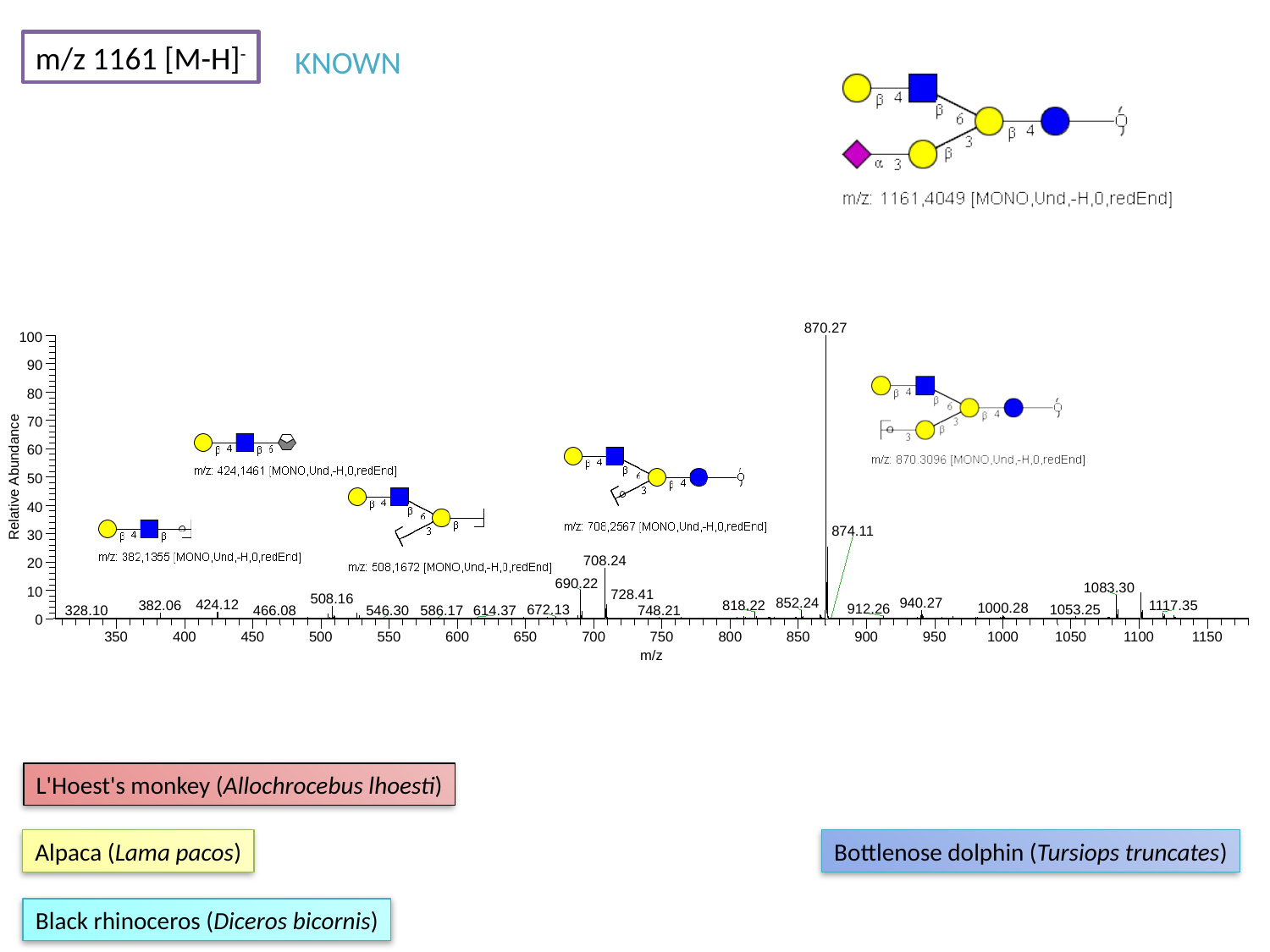

m/z 1161 [M-H]^-^
KNOWN
870.27
100
90
80
70
60
Relative Abundance
50
40
30
708.24
20
690.22
1083.30
10
508.16
852.24
940.27
424.12
1117.35
0
350
400
450
500
550
600
650
700
750
800
850
900
950
1000
1050
1100
1150
m/z
874.11
728.41
818.22
382.06
1000.28
912.26
1053.25
672.13
748.21
546.30
466.08
586.17
614.37
328.10
L'Hoest's monkey (Allochrocebus lhoesti)
Alpaca (Lama pacos)
Bottlenose dolphin (Tursiops truncates)
Black rhinoceros (Diceros bicornis)

### Slide 103
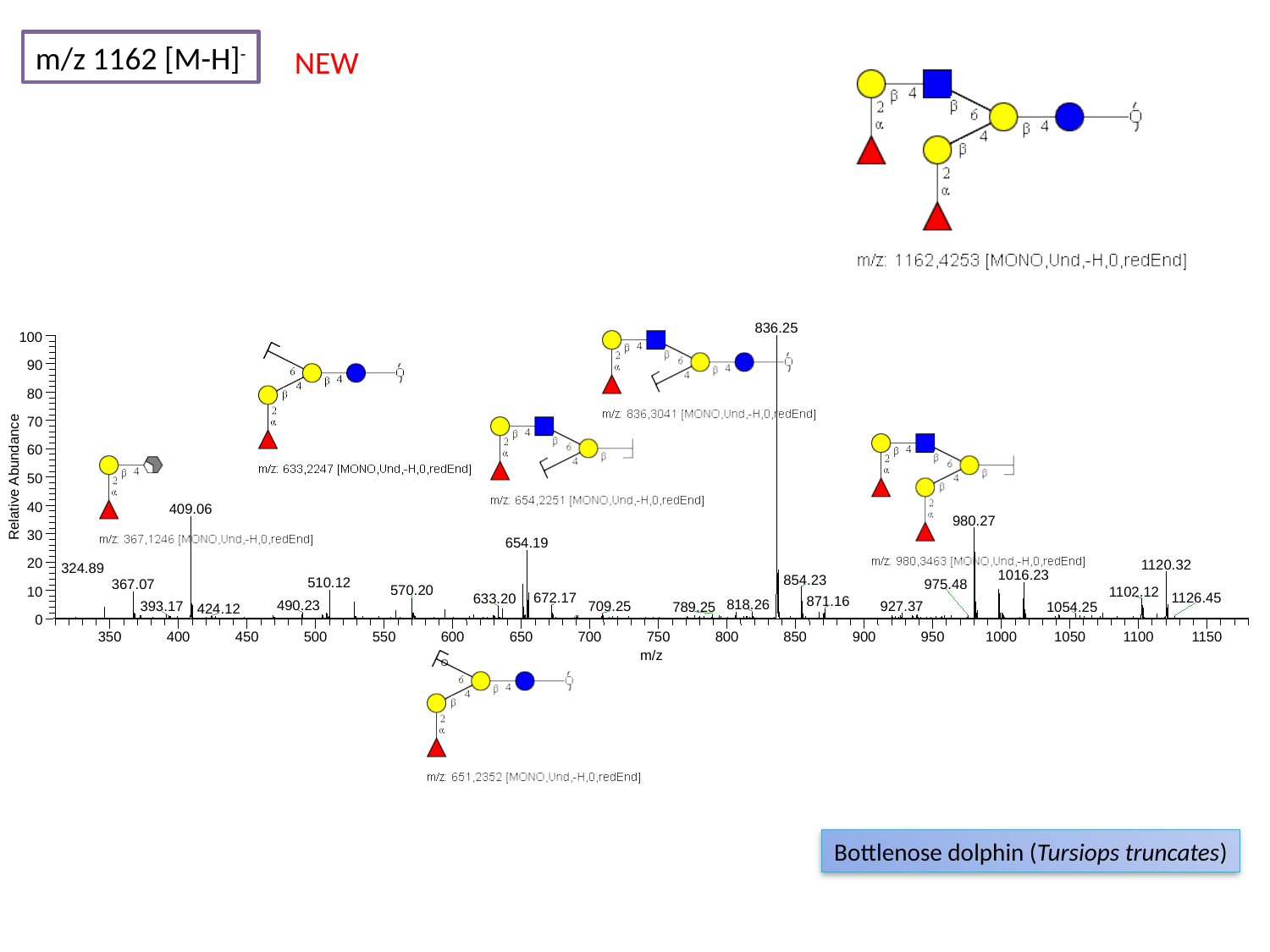

m/z 1162 [M-H]^-^
NEW
836.25
100
90
80
70
60
Relative Abundance
50
40
409.06
980.27
30
654.19
20
1120.32
1016.23
854.23
510.12
367.07
570.20
10
0
350
400
450
500
550
600
650
700
750
800
850
900
950
1000
1050
1100
1150
m/z
324.89
975.48
1102.12
1126.45
672.17
633.20
871.16
818.26
490.23
927.37
709.25
393.17
1054.25
789.25
424.12
Bottlenose dolphin (Tursiops truncates)

### Slide 104
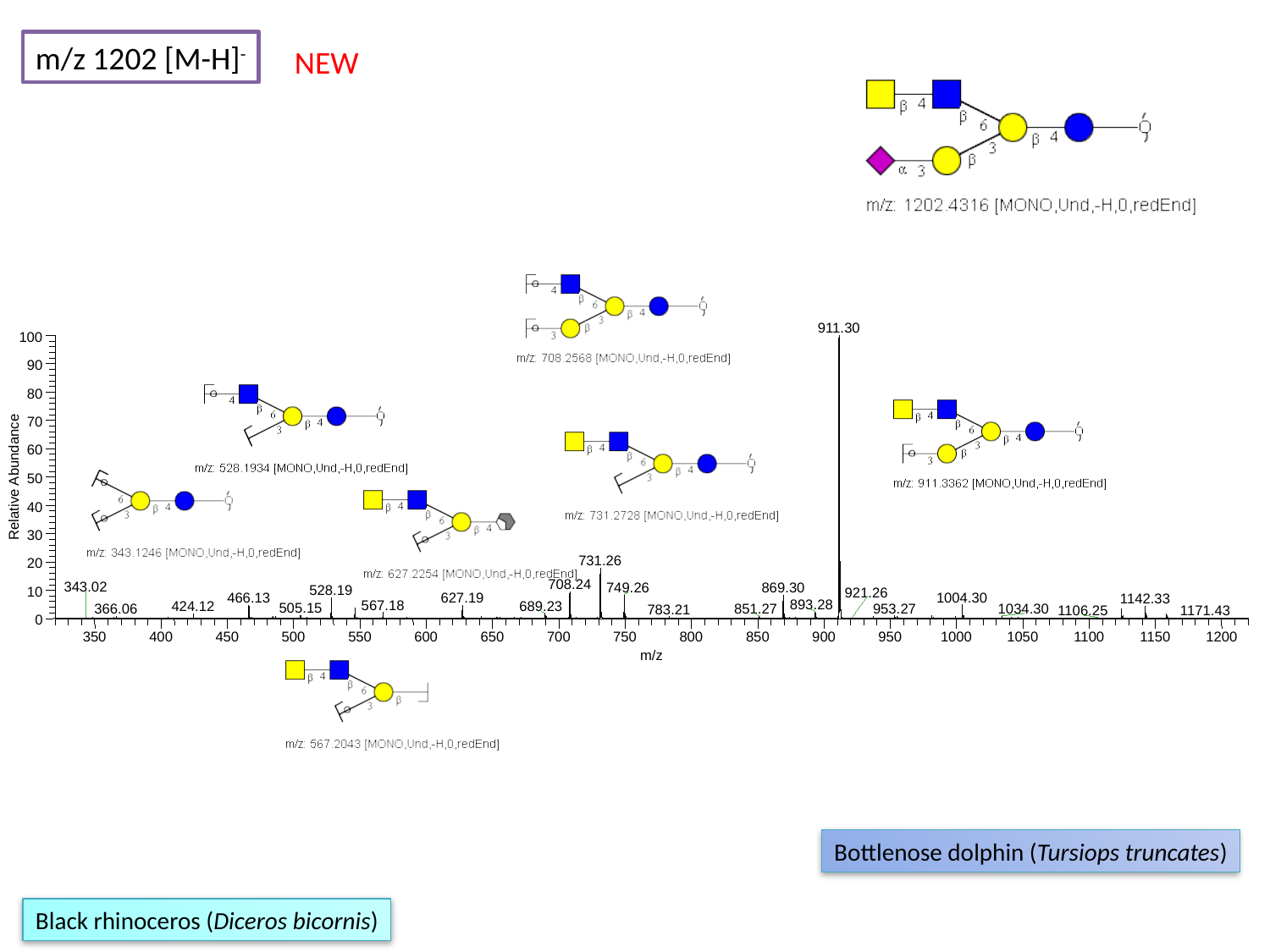

m/z 1202 [M-H]^-^
NEW
911.30
100
90
80
70
60
Relative Abundance
50
40
30
731.26
20
708.24
749.26
869.30
528.19
10
1004.30
0
350
400
450
500
550
600
650
700
750
800
850
900
950
1000
1050
1100
1150
1200
m/z
343.02
921.26
627.19
466.13
1142.33
893.28
567.18
424.12
689.23
505.15
953.27
366.06
851.27
1034.30
783.21
1106.25
1171.43
Bottlenose dolphin (Tursiops truncates)
Black rhinoceros (Diceros bicornis)

### Slide 105
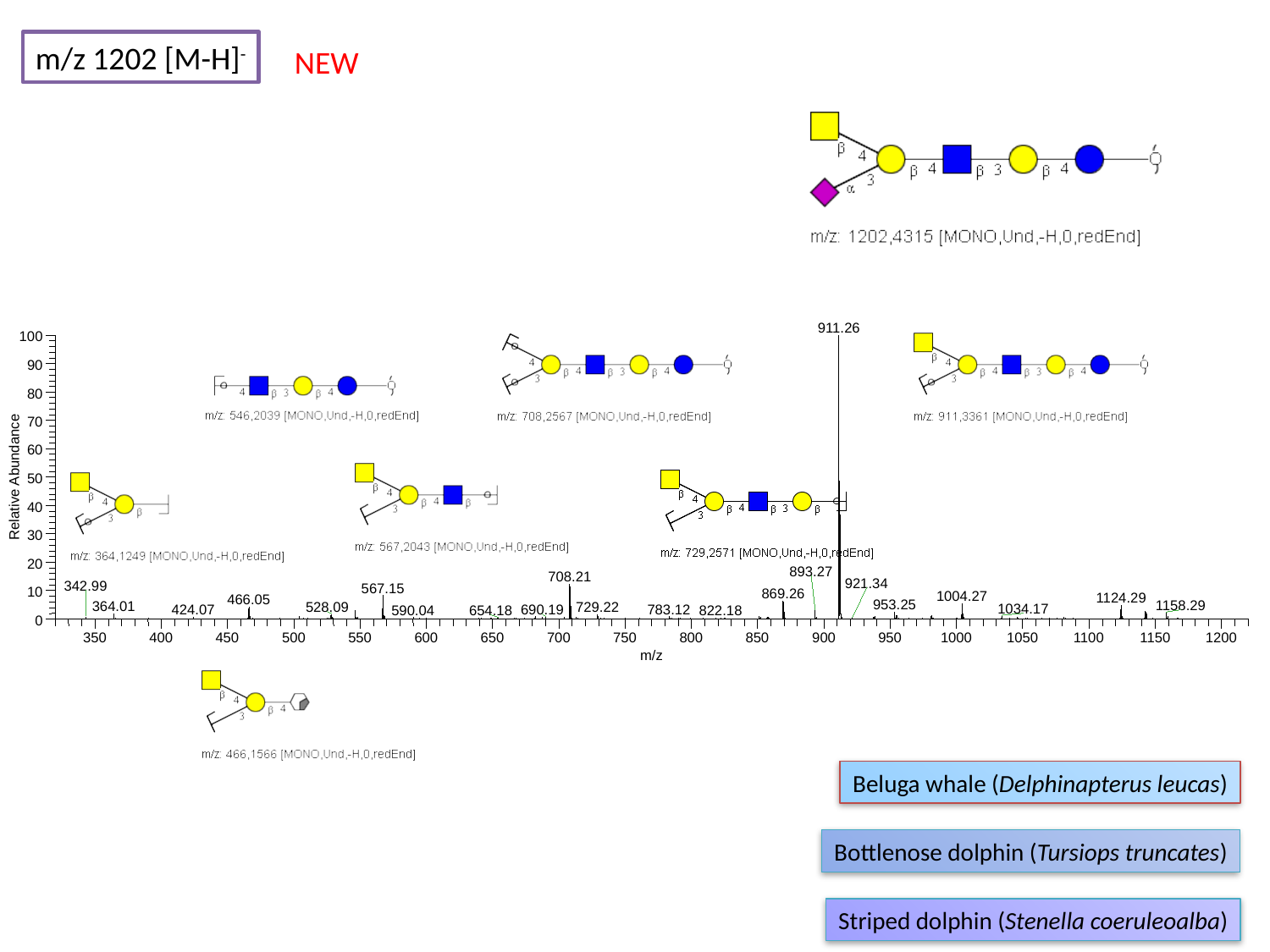

m/z 1202 [M-H]^-^
NEW
911.26
100
90
80
70
60
Relative Abundance
50
40
30
20
893.27
708.21
567.15
10
869.26
1004.27
1124.29
466.05
0
350
400
450
500
550
600
650
700
750
800
850
900
950
1000
1050
1100
1150
1200
m/z
921.34
342.99
953.25
1158.29
364.01
528.09
729.22
1034.17
690.19
783.12
424.07
590.04
654.18
822.18
Beluga whale (Delphinapterus leucas)
Bottlenose dolphin (Tursiops truncates)
Striped dolphin (Stenella coeruleoalba)

### Slide 106
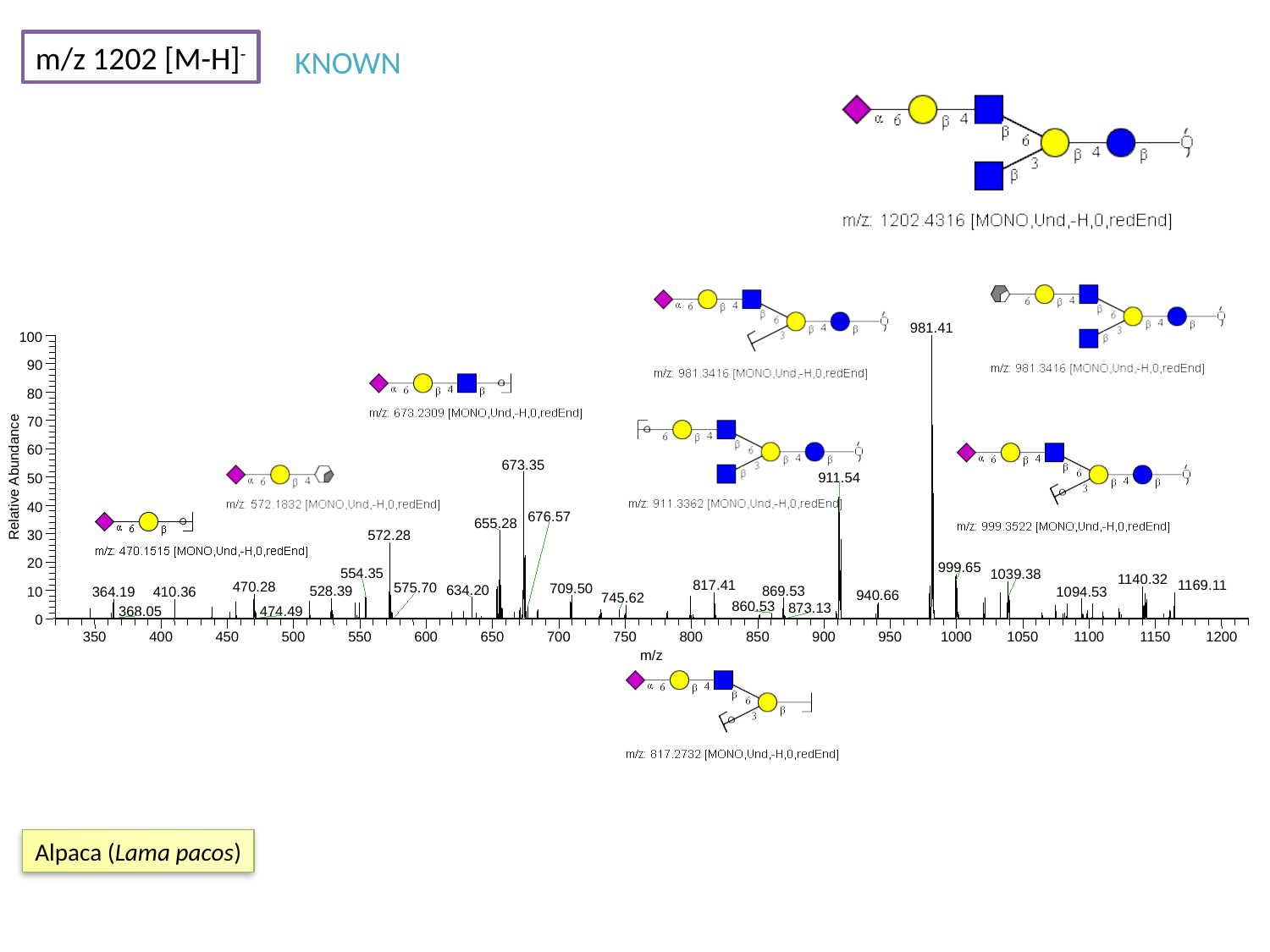

m/z 1202 [M-H]^-^
KNOWN
981.41
100
90
80
70
60
673.35
Relative Abundance
911.54
50
40
655.28
30
572.28
20
999.65
10
0
350
400
450
500
550
600
650
700
750
800
850
900
950
1000
1050
1100
1150
1200
m/z
676.57
554.35
1039.38
1140.32
1169.11
817.41
470.28
575.70
709.50
634.20
869.53
528.39
1094.53
364.19
410.36
940.66
745.62
860.53
873.13
368.05
474.49
Alpaca (Lama pacos)

### Slide 107
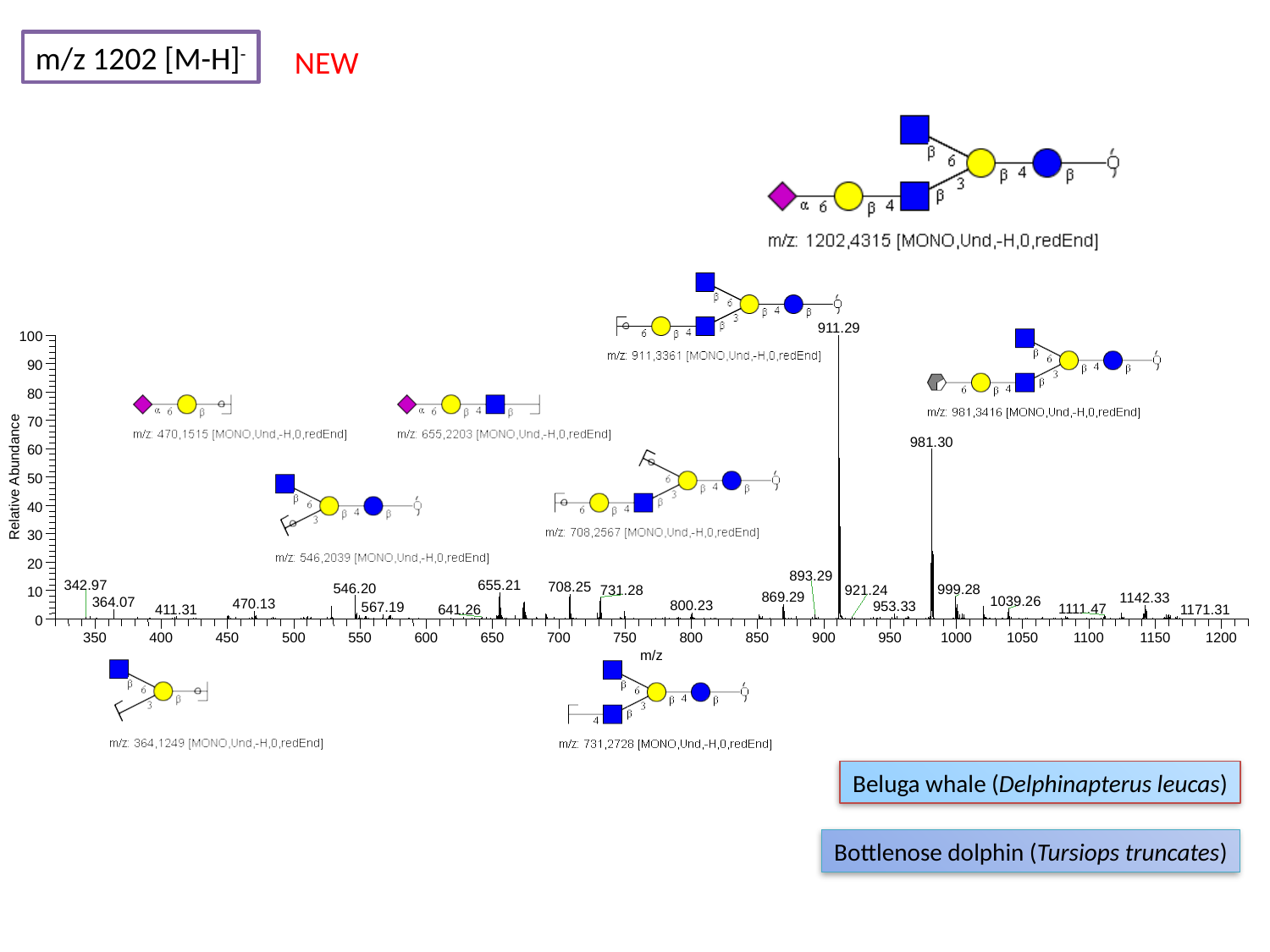

m/z 1202 [M-H]^-^
NEW
911.29
100
90
80
70
981.30
60
Relative Abundance
50
40
30
20
655.21
708.25
546.20
999.28
731.28
10
0
350
400
450
500
550
600
650
700
750
800
850
900
950
1000
1050
1100
1150
1200
m/z
893.29
342.97
921.24
869.29
1142.33
1039.26
364.07
470.13
800.23
953.33
567.19
1111.47
411.31
1171.31
641.26
Beluga whale (Delphinapterus leucas)
Bottlenose dolphin (Tursiops truncates)

### Slide 108
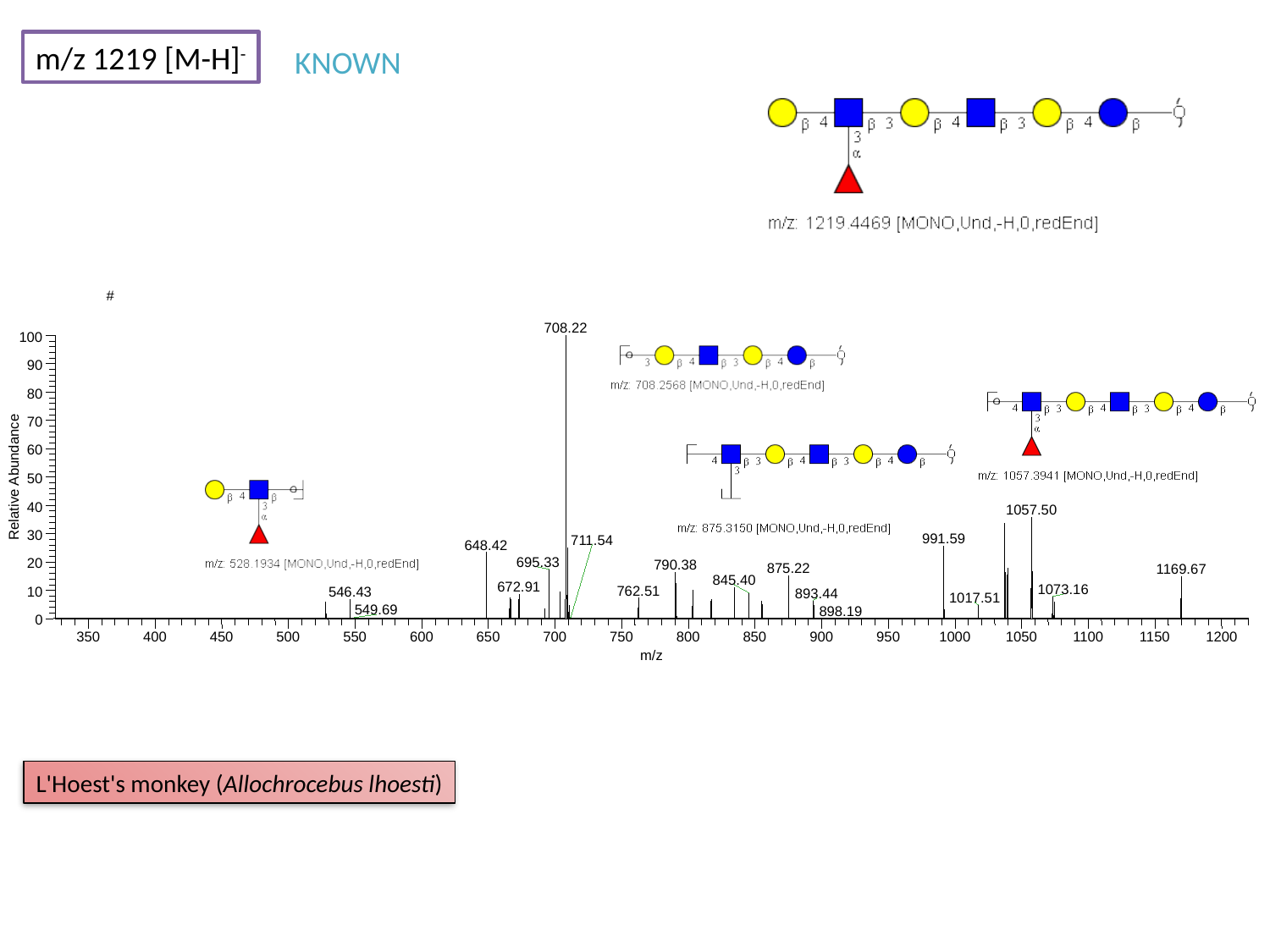

m/z 1219 [M-H]^-^
KNOWN
#
708.22
100
90
80
70
60
Relative Abundance
50
40
1057.50
30
991.59
648.42
695.33
20
790.38
875.22
845.40
10
0
350
400
450
500
550
600
650
700
750
800
850
900
950
1000
1050
1100
1150
1200
m/z
711.54
1169.67
672.91
1073.16
762.51
546.43
893.44
1017.51
549.69
898.19
L'Hoest's monkey (Allochrocebus lhoesti)

### Slide 109
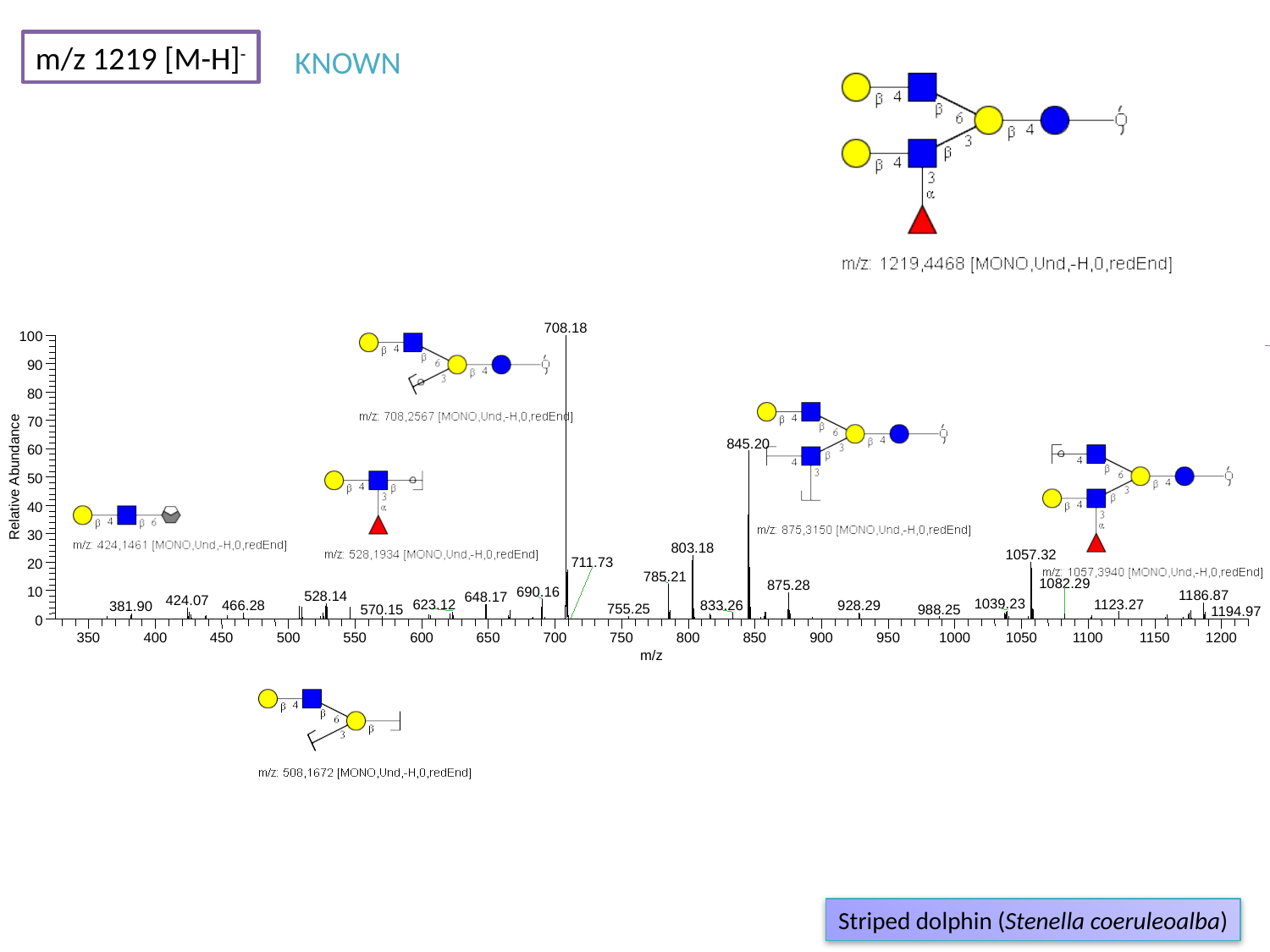

m/z 1219 [M-H]^-^
KNOWN
708.18
100
90
80
70
845.20
60
Relative Abundance
50
40
30
803.18
1057.32
20
785.21
875.28
10
690.16
0
350
400
450
500
550
600
650
700
750
800
850
900
950
1000
1050
1100
1150
1200
m/z
711.73
1082.29
1186.87
528.14
648.17
424.07
1039.23
1123.27
623.12
928.29
466.28
833.26
381.90
755.25
570.15
988.25
1194.97
Striped dolphin (Stenella coeruleoalba)

### Slide 110
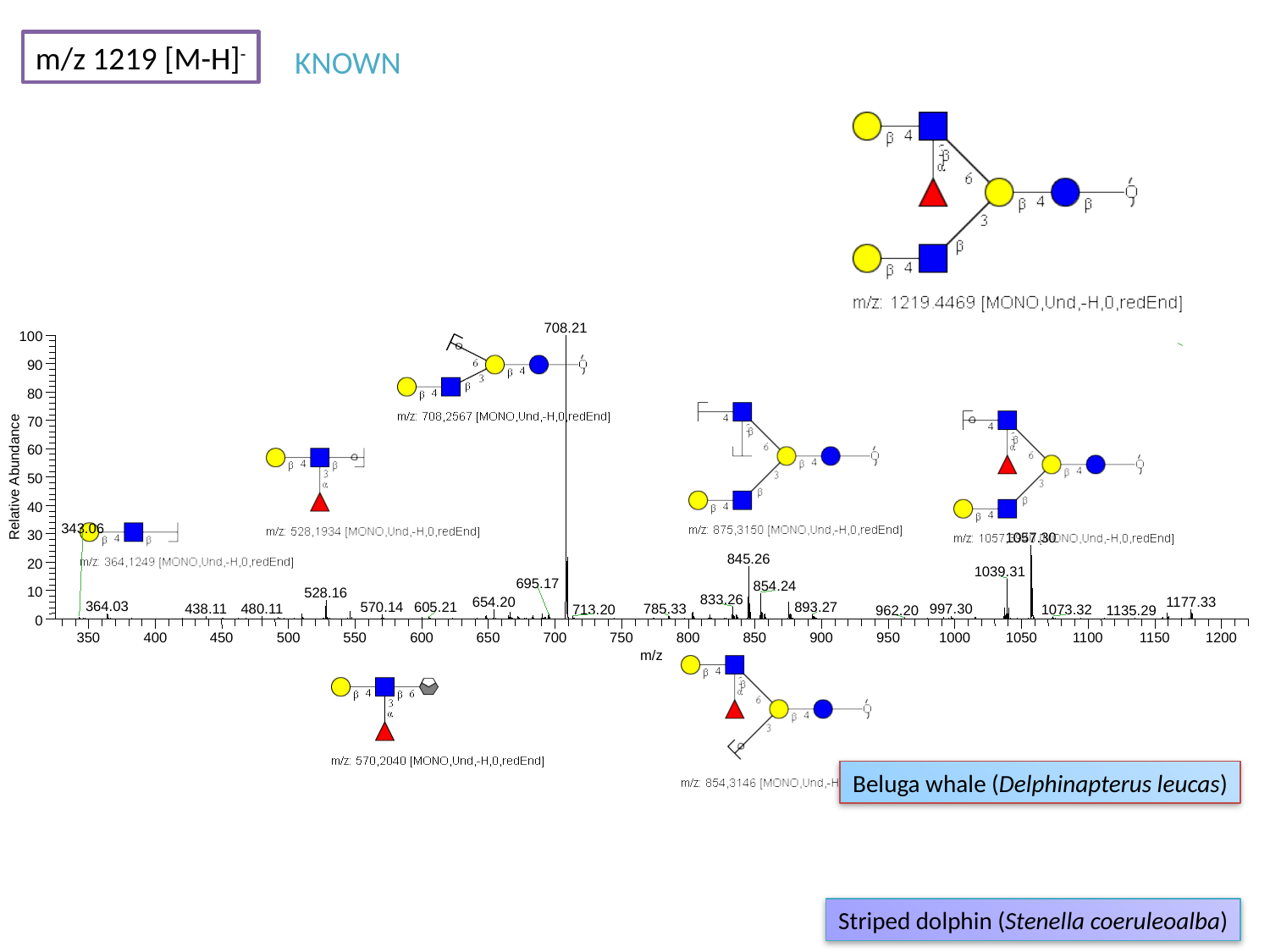

m/z 1219 [M-H]^-^
KNOWN
708.21
100
90
80
70
60
Relative Abundance
50
40
30
1057.30
845.26
20
1039.31
854.24
10
528.16
833.26
0
350
400
450
500
550
600
650
700
750
800
850
900
950
1000
1050
1100
1150
1200
m/z
343.06
695.17
654.20
1177.33
364.03
570.14
605.21
893.27
480.11
438.11
997.30
785.33
713.20
1073.32
962.20
1135.29
Beluga whale (Delphinapterus leucas)
Striped dolphin (Stenella coeruleoalba)

### Slide 111
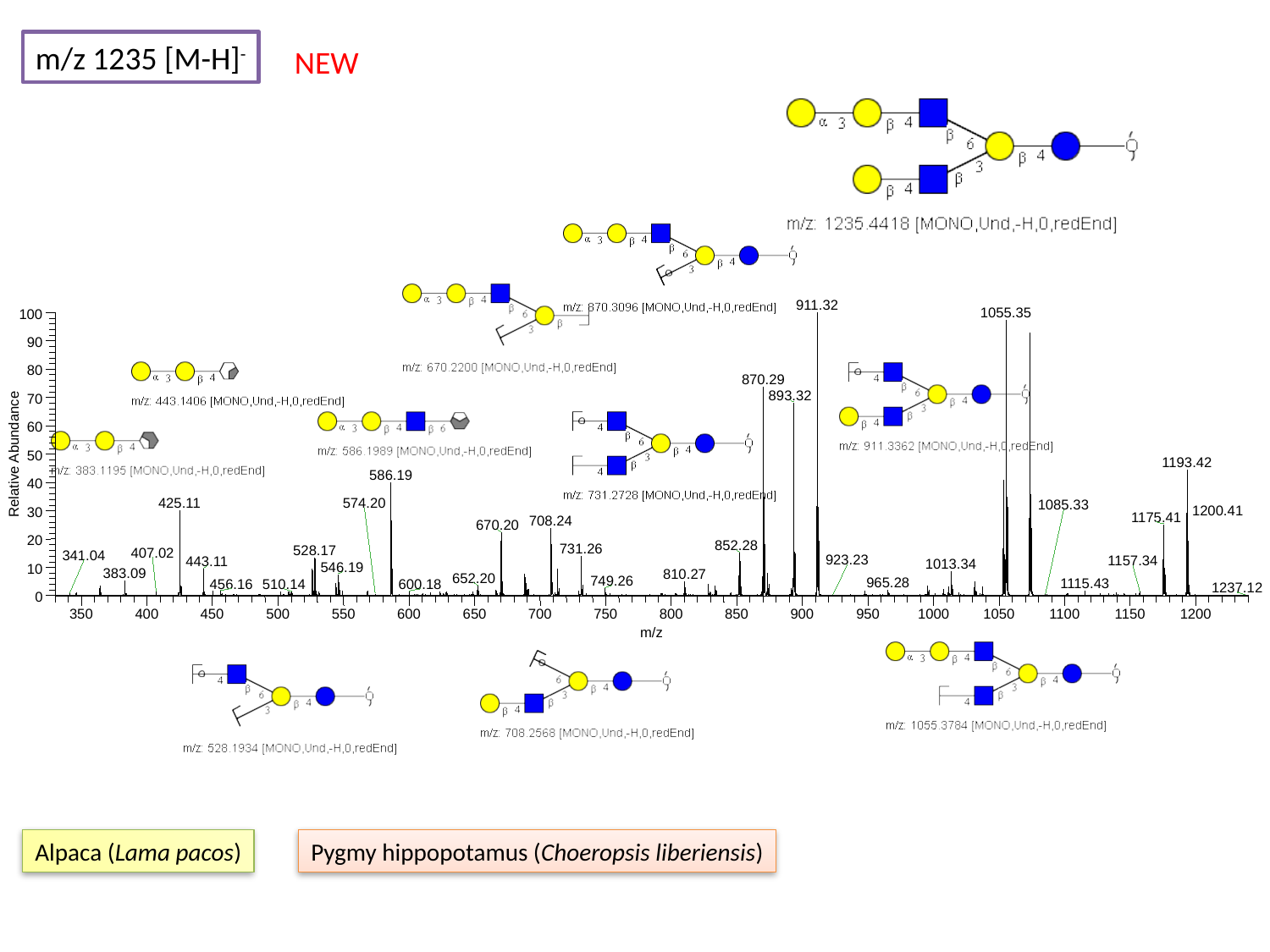

m/z 1235 [M-H]^-^
NEW
911.32
1055.35
100
90
80
870.29
893.32
70
60
Relative Abundance
50
1193.42
586.19
40
30
20
10
0
350
400
450
500
550
600
650
700
750
800
850
900
950
1000
1050
1100
1150
1200
m/z
425.11
574.20
1085.33
1200.41
1175.41
708.24
670.20
852.28
731.26
528.17
407.02
341.04
923.23
1157.34
443.11
1013.34
546.19
383.09
810.27
652.20
749.26
965.28
1115.43
456.16
600.18
510.14
1237.12
Alpaca (Lama pacos)
Pygmy hippopotamus (Choeropsis liberiensis)

### Slide 112
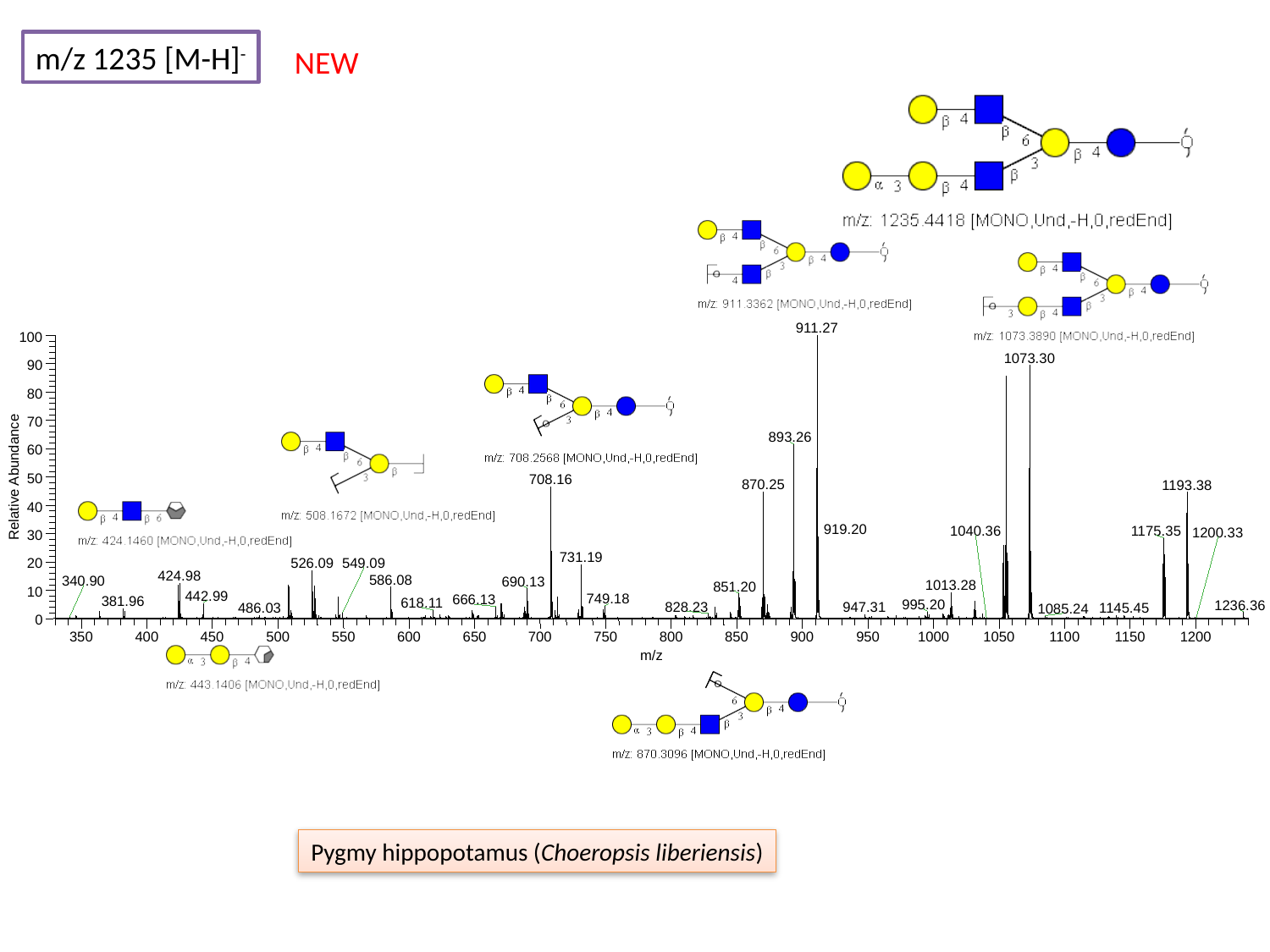

m/z 1235 [M-H]^-^
NEW
911.27
100
1073.30
90
80
70
893.26
60
Relative Abundance
50
708.16
870.25
1193.38
40
30
20
10
0
350
400
450
500
550
600
650
700
750
800
850
900
950
1000
1050
1100
1150
1200
m/z
919.20
1040.36
1175.35
1200.33
731.19
526.09
549.09
424.98
586.08
340.90
690.13
1013.28
851.20
442.99
749.18
666.13
381.96
618.11
995.20
1236.36
828.23
947.31
486.03
1145.45
1085.24
Pygmy hippopotamus (Choeropsis liberiensis)

### Slide 113
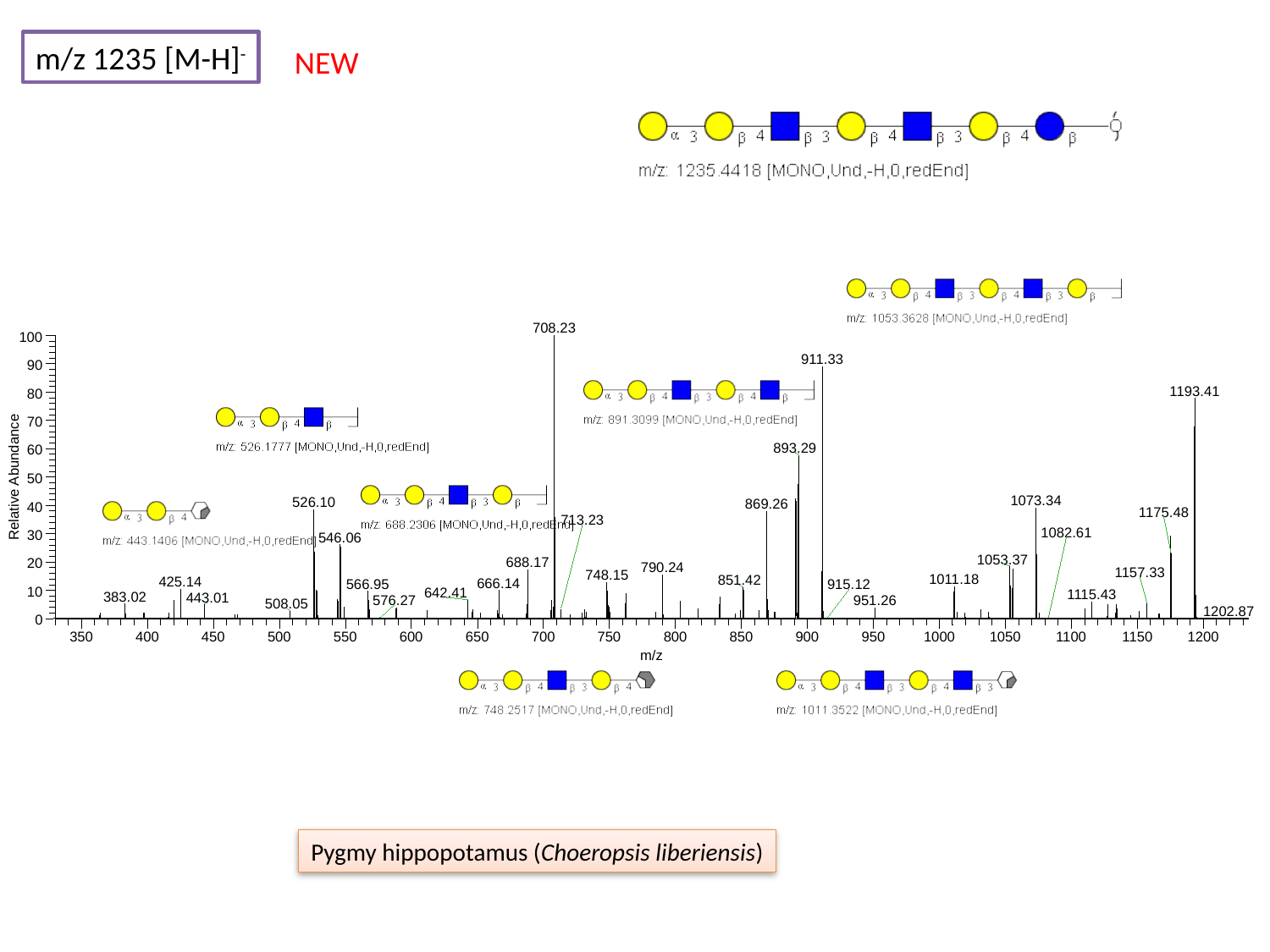

m/z 1235 [M-H]^-^
NEW
708.23
100
911.33
90
1193.41
80
70
893.29
60
Relative Abundance
50
1073.34
526.10
869.26
40
30
20
10
0
350
400
450
500
550
600
650
700
750
800
850
900
950
1000
1050
1100
1150
1200
m/z
1175.48
713.23
1082.61
546.06
1053.37
688.17
790.24
1157.33
748.15
1011.18
851.42
425.14
666.14
566.95
915.12
642.41
1115.43
383.02
443.01
576.27
951.26
508.05
1202.87
Pygmy hippopotamus (Choeropsis liberiensis)

### Slide 114
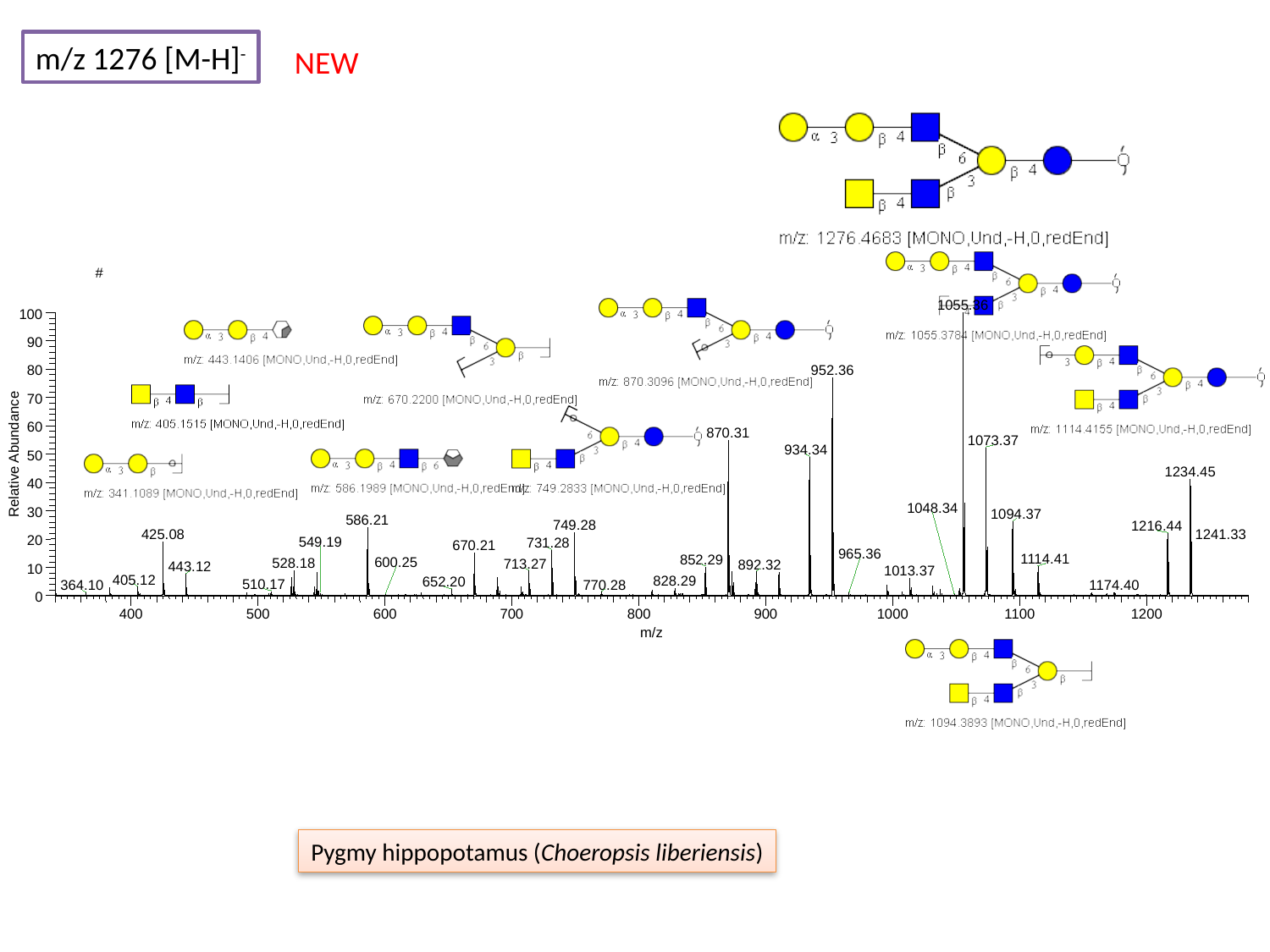

m/z 1276 [M-H]^-^
NEW
#
400
1055.36
100
90
952.36
80
70
60
870.31
1073.37
934.34
Relative Abundance
50
1234.45
40
1048.34
30
1094.37
586.21
749.28
1216.44
1241.33
425.08
20
549.19
731.28
670.21
965.36
1114.41
852.29
600.25
528.18
713.27
892.32
443.12
10
1013.37
405.12
828.29
652.20
510.17
770.28
1174.40
364.10
0
500
600
700
800
900
1000
1100
1200
m/z
Pygmy hippopotamus (Choeropsis liberiensis)

### Slide 115
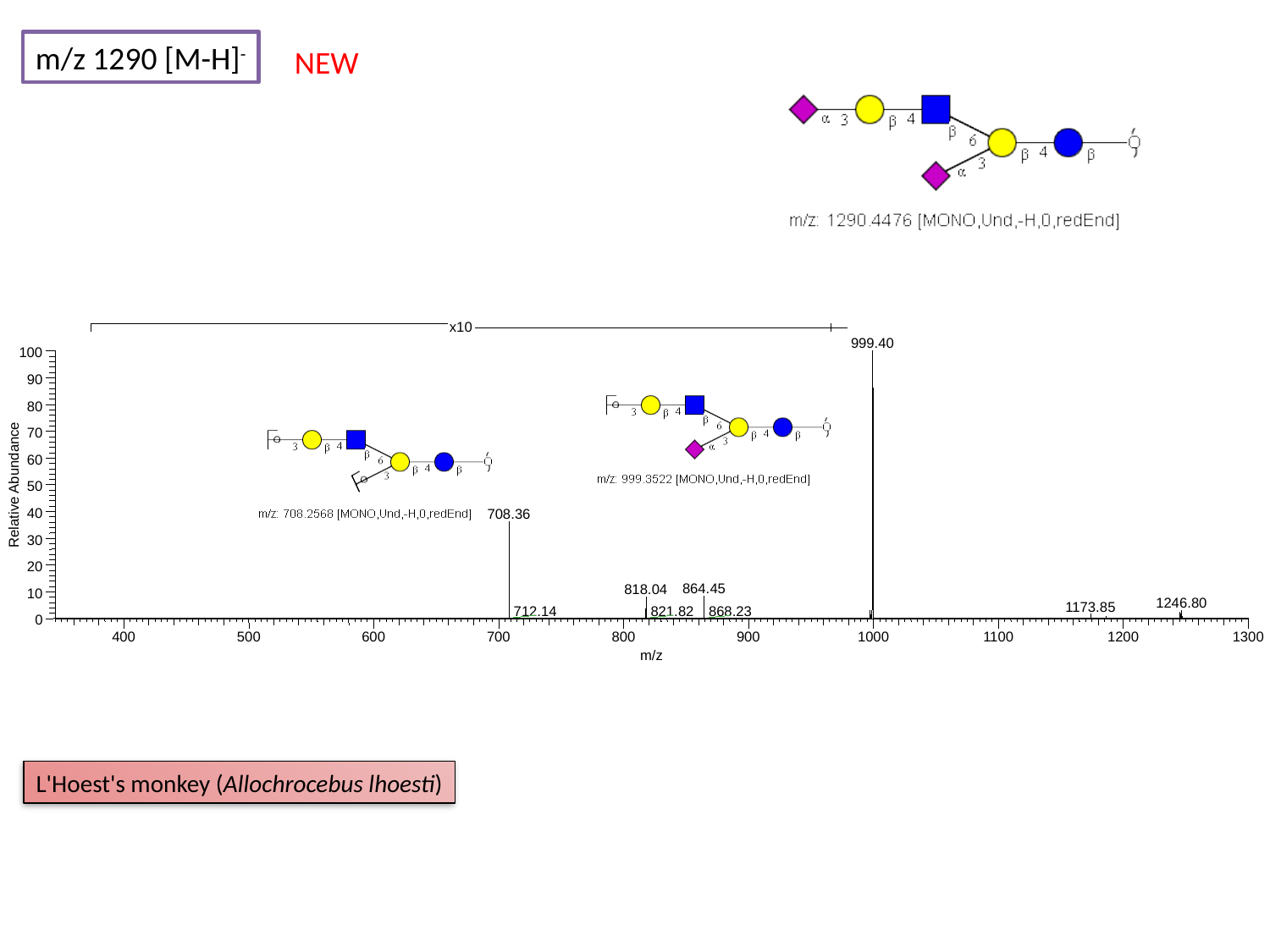

m/z 1290 [M-H]^-^
NEW
x10
999.40
100
90
80
70
60
Relative Abundance
50
40
708.36
30
20
864.45
818.04
10
1246.80
1173.85
712.14
821.82
868.23
0
400
500
600
700
800
900
1000
1100
1200
1300
m/z
L'Hoest's monkey (Allochrocebus lhoesti)

### Slide 116
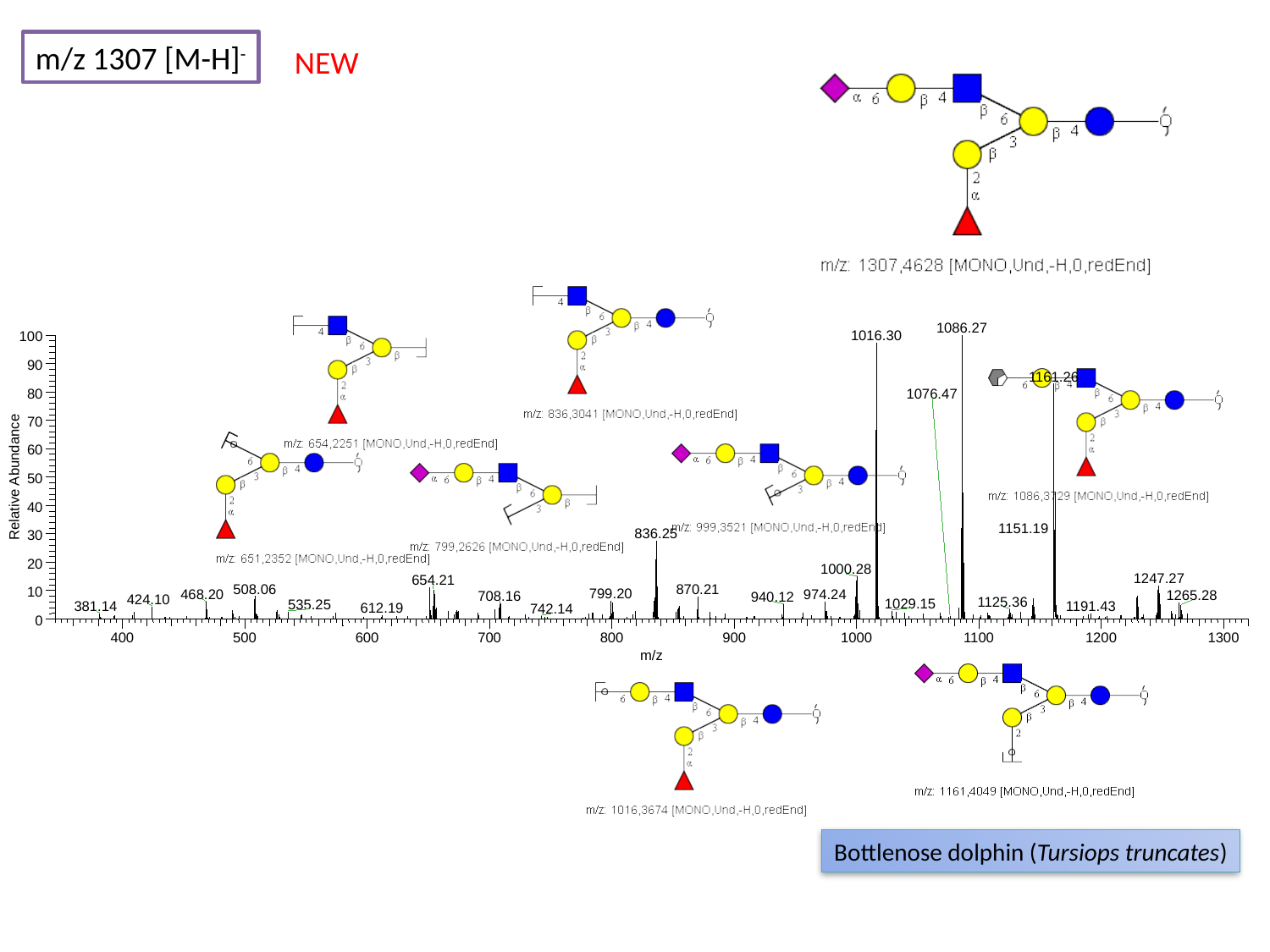

m/z 1307 [M-H]^-^
NEW
1086.27
1016.30
100
90
1161.26
80
1076.47
70
60
Relative Abundance
50
40
1151.19
836.25
30
20
1000.28
1247.27
654.21
508.06
870.21
10
799.20
468.20
974.24
1265.28
708.16
940.12
424.10
1125.36
1029.15
535.25
381.14
1191.43
612.19
742.14
0
400
500
600
700
800
900
1000
1100
1200
1300
m/z
Bottlenose dolphin (Tursiops truncates)

### Slide 117
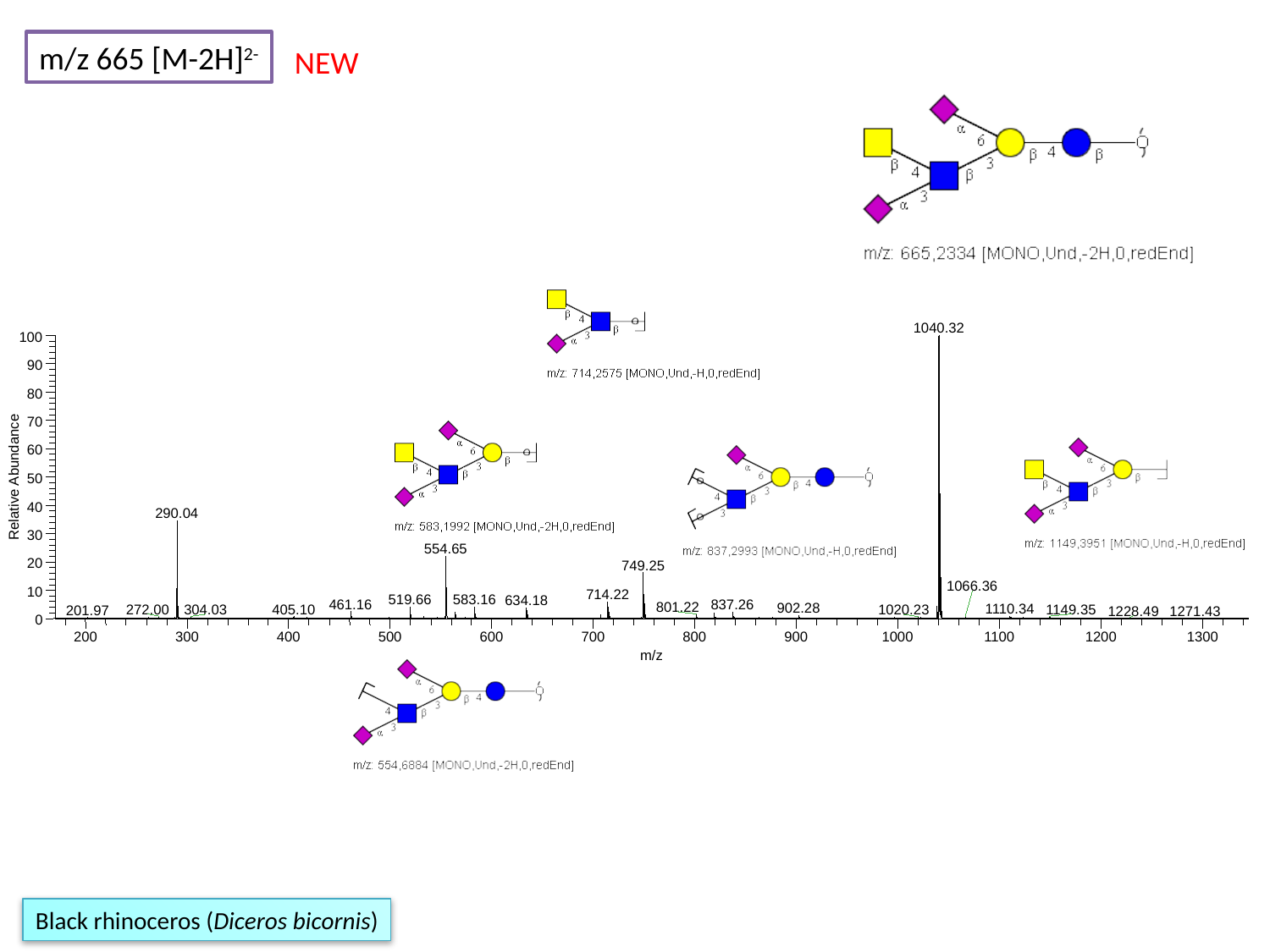

m/z 665 [M-2H]^2-^
NEW
1040.32
100
90
80
70
60
Relative Abundance
50
40
290.04
30
554.65
20
749.25
1066.36
10
714.22
583.16
519.66
634.18
461.16
837.26
801.22
902.28
1110.34
405.10
272.00
1149.35
304.03
1020.23
201.97
1228.49
1271.43
0
200
300
400
500
600
700
800
900
1000
1100
1200
1300
m/z
Black rhinoceros (Diceros bicornis)

### Slide 118
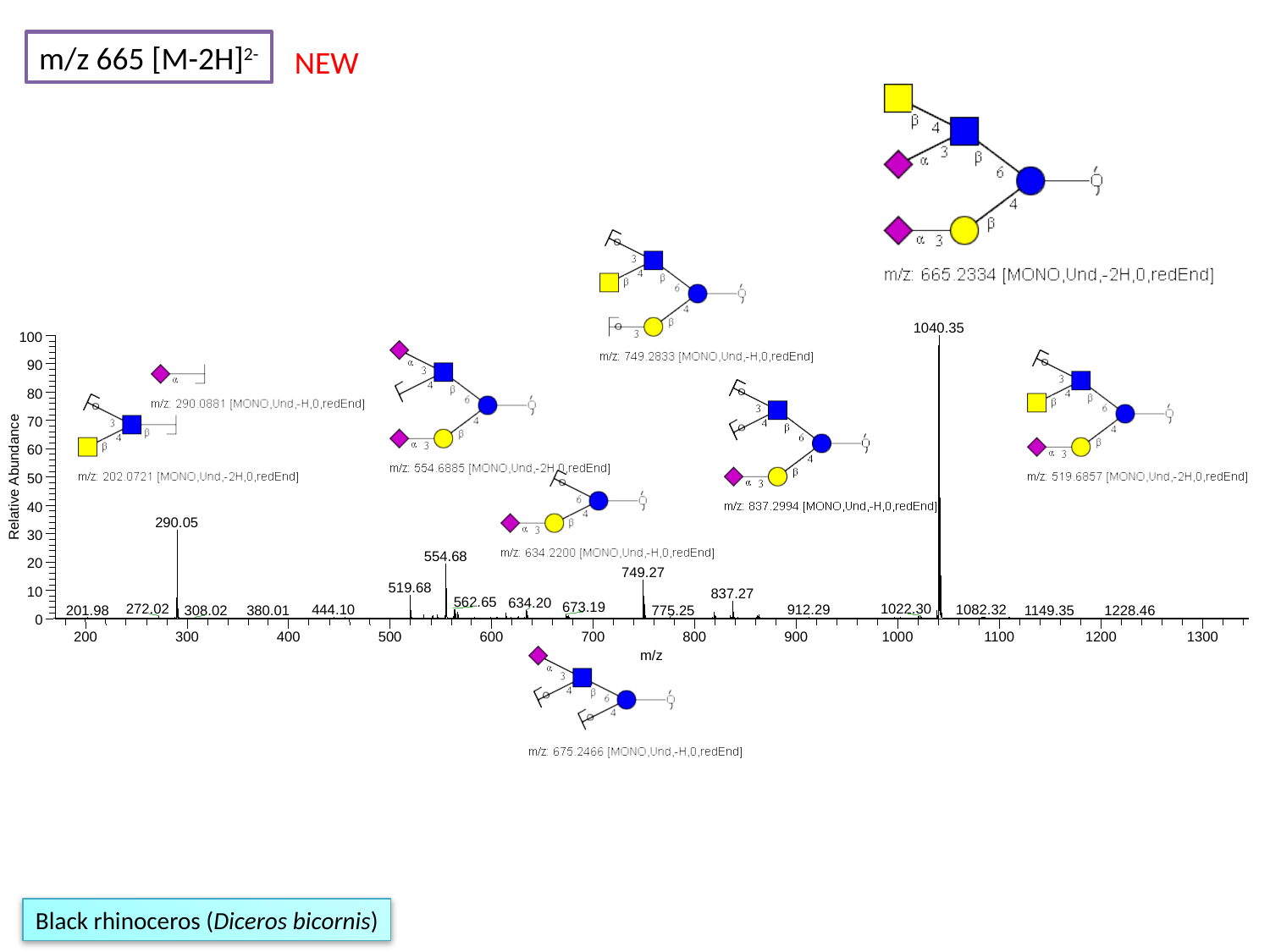

m/z 665 [M-2H]^2-^
NEW
1040.35
100
90
80
70
60
Relative Abundance
50
40
290.05
30
554.68
20
749.27
519.68
10
837.27
562.65
634.20
673.19
272.02
1022.30
1082.32
444.10
912.29
201.98
775.25
308.02
380.01
1149.35
1228.46
0
200
300
400
500
600
700
800
900
1000
1100
1200
1300
m/z
Black rhinoceros (Diceros bicornis)

### Slide 119
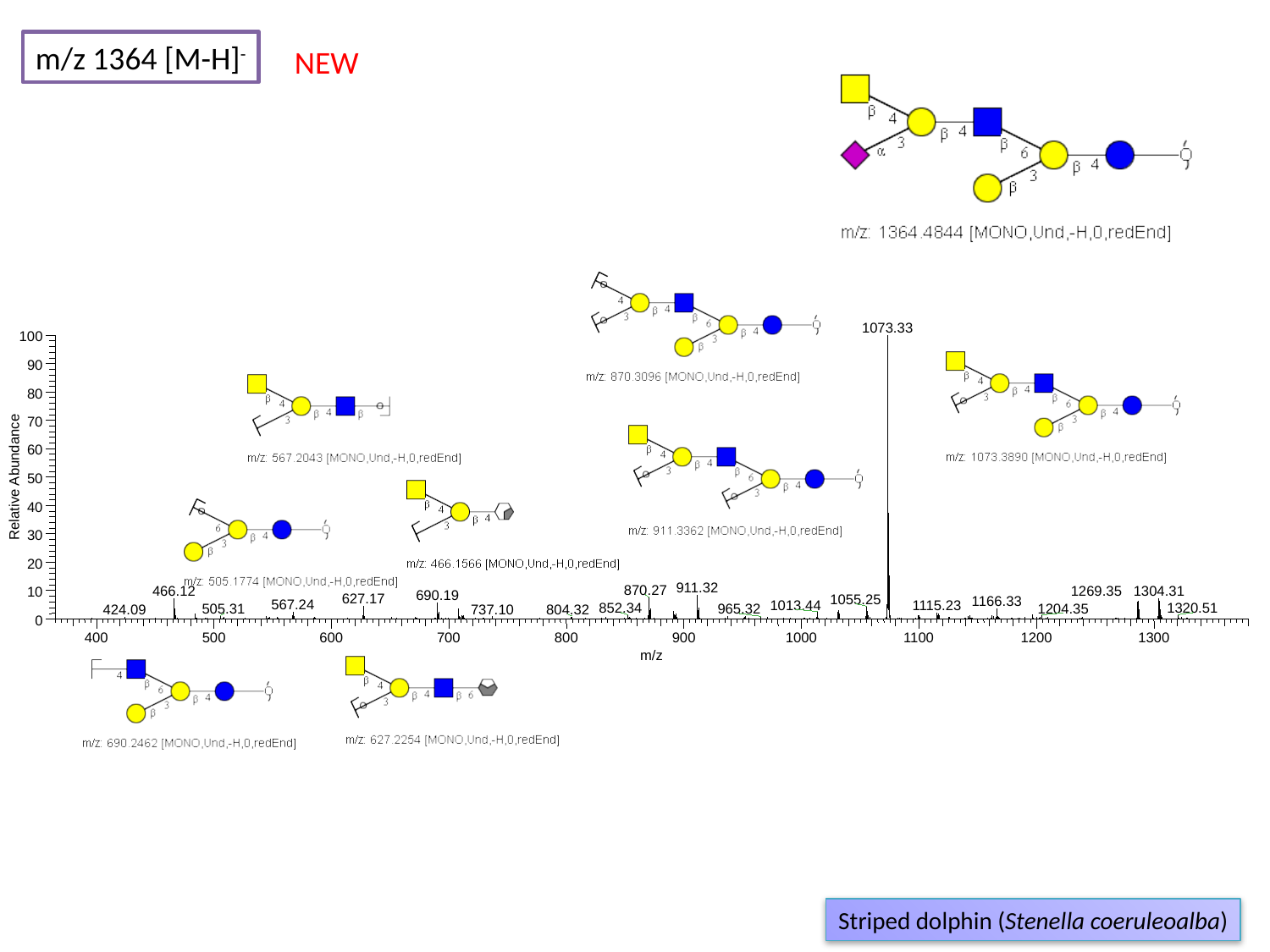

m/z 1364 [M-H]^-^
NEW
1073.33
100
90
80
70
60
Relative Abundance
50
40
30
20
911.32
870.27
1269.35
1304.31
466.12
10
690.19
627.17
1055.25
1166.33
567.24
1013.44
1115.23
852.34
1320.51
1204.35
505.31
965.32
737.10
424.09
804.32
0
400
500
600
700
800
900
1000
1100
1200
1300
m/z
Striped dolphin (Stenella coeruleoalba)

### Slide 120
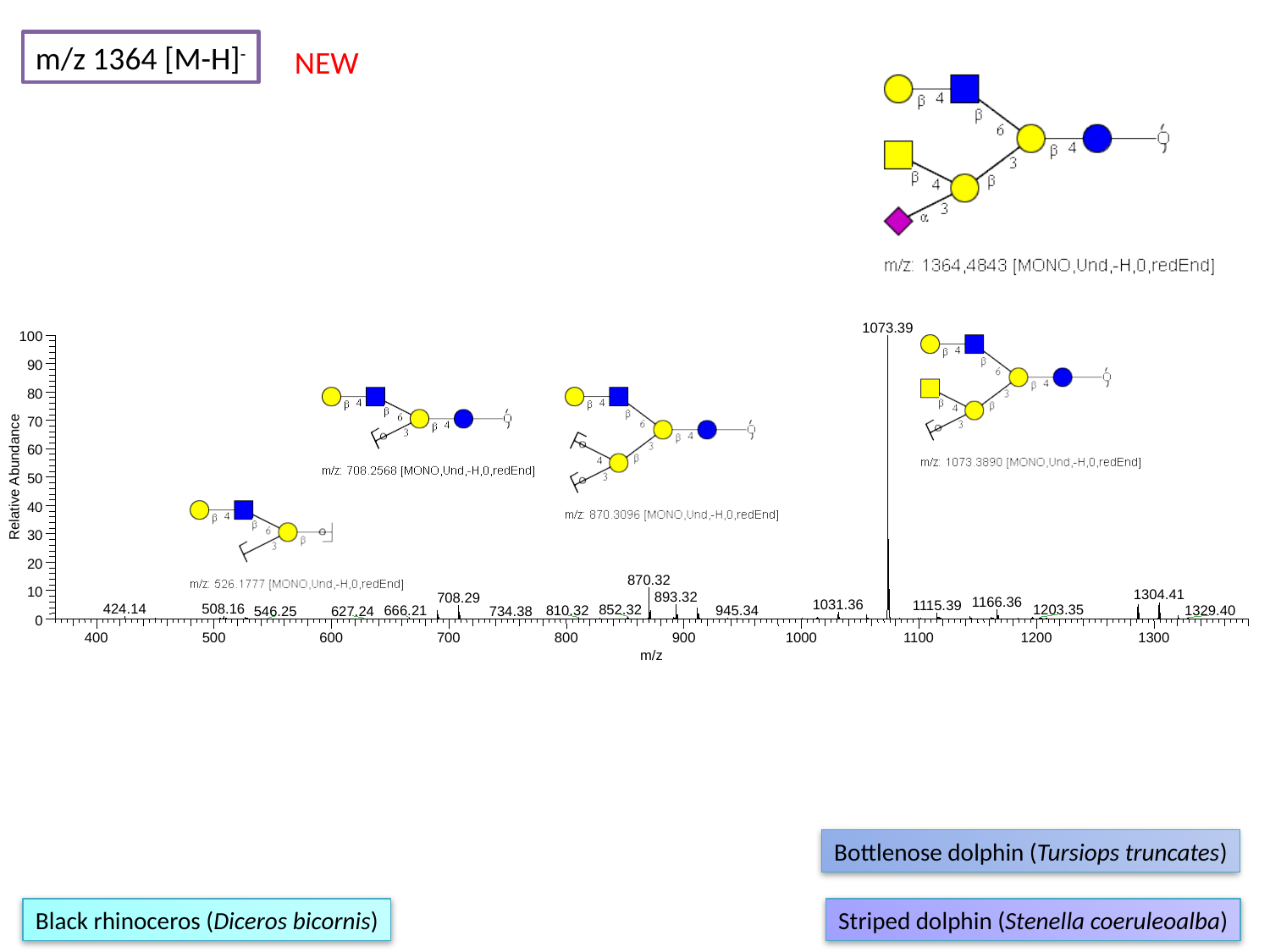

m/z 1364 [M-H]^-^
NEW
1073.39
100
90
80
70
60
Relative Abundance
50
40
30
20
870.32
10
1304.41
893.32
708.29
1166.36
1031.36
1115.39
508.16
424.14
1203.35
852.32
666.21
810.32
945.34
1329.40
546.25
627.24
734.38
0
400
500
600
700
800
900
1000
1100
1200
1300
m/z
Bottlenose dolphin (Tursiops truncates)
Black rhinoceros (Diceros bicornis)
Striped dolphin (Stenella coeruleoalba)

### Slide 121
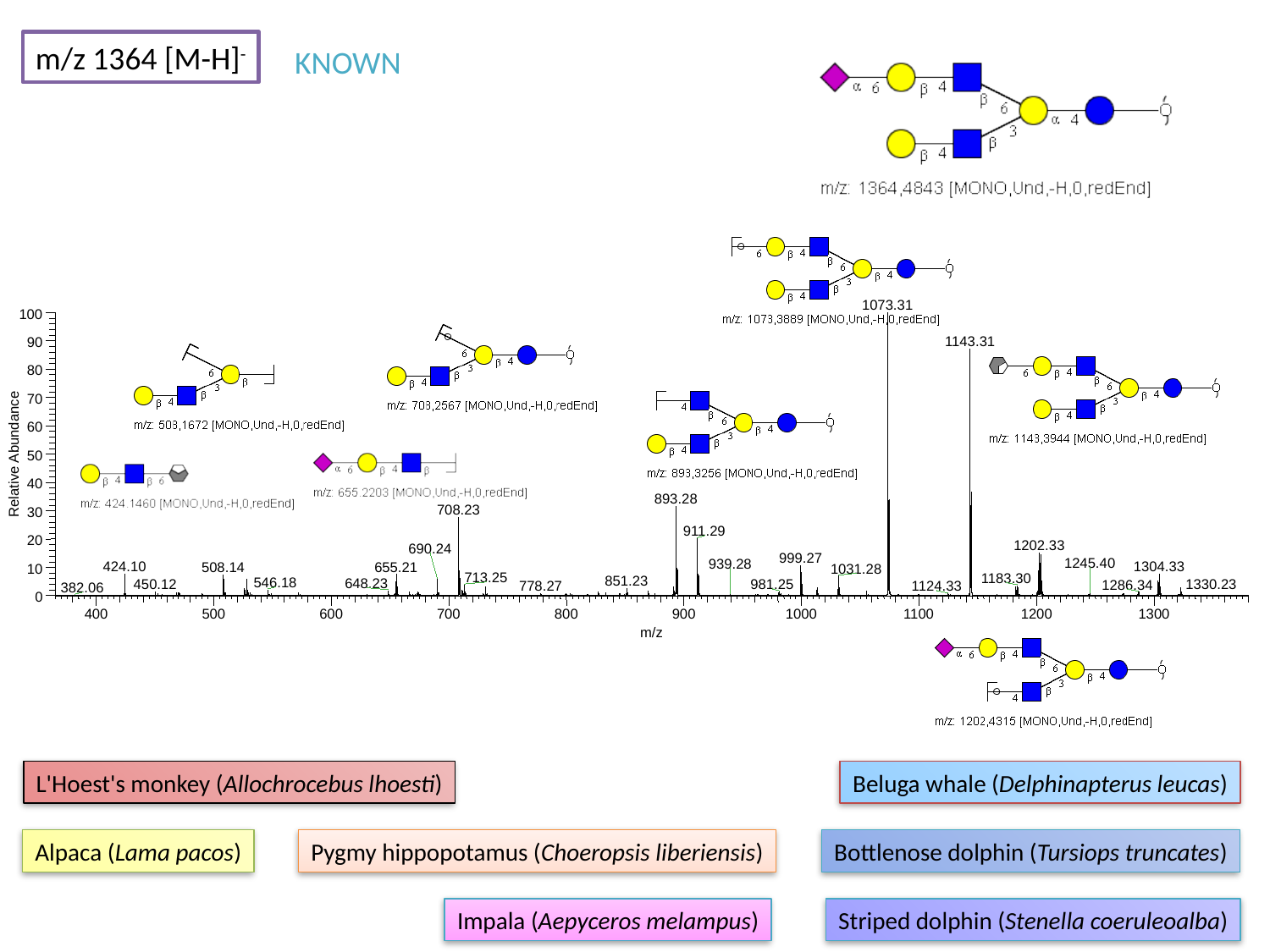

m/z 1364 [M-H]^-^
KNOWN
1073.31
100
1143.31
90
80
70
60
Relative Abundance
50
40
893.28
708.23
30
911.29
20
1202.33
690.24
999.27
1245.40
939.28
424.10
1304.33
655.21
508.14
1031.28
10
713.25
1183.30
851.23
546.18
648.23
1330.23
450.12
981.25
1286.34
778.27
1124.33
382.06
0
400
500
600
700
800
900
1000
1100
1200
1300
m/z
L'Hoest's monkey (Allochrocebus lhoesti)
Beluga whale (Delphinapterus leucas)
Alpaca (Lama pacos)
Pygmy hippopotamus (Choeropsis liberiensis)
Bottlenose dolphin (Tursiops truncates)
Impala (Aepyceros melampus)
Striped dolphin (Stenella coeruleoalba)

### Slide 122
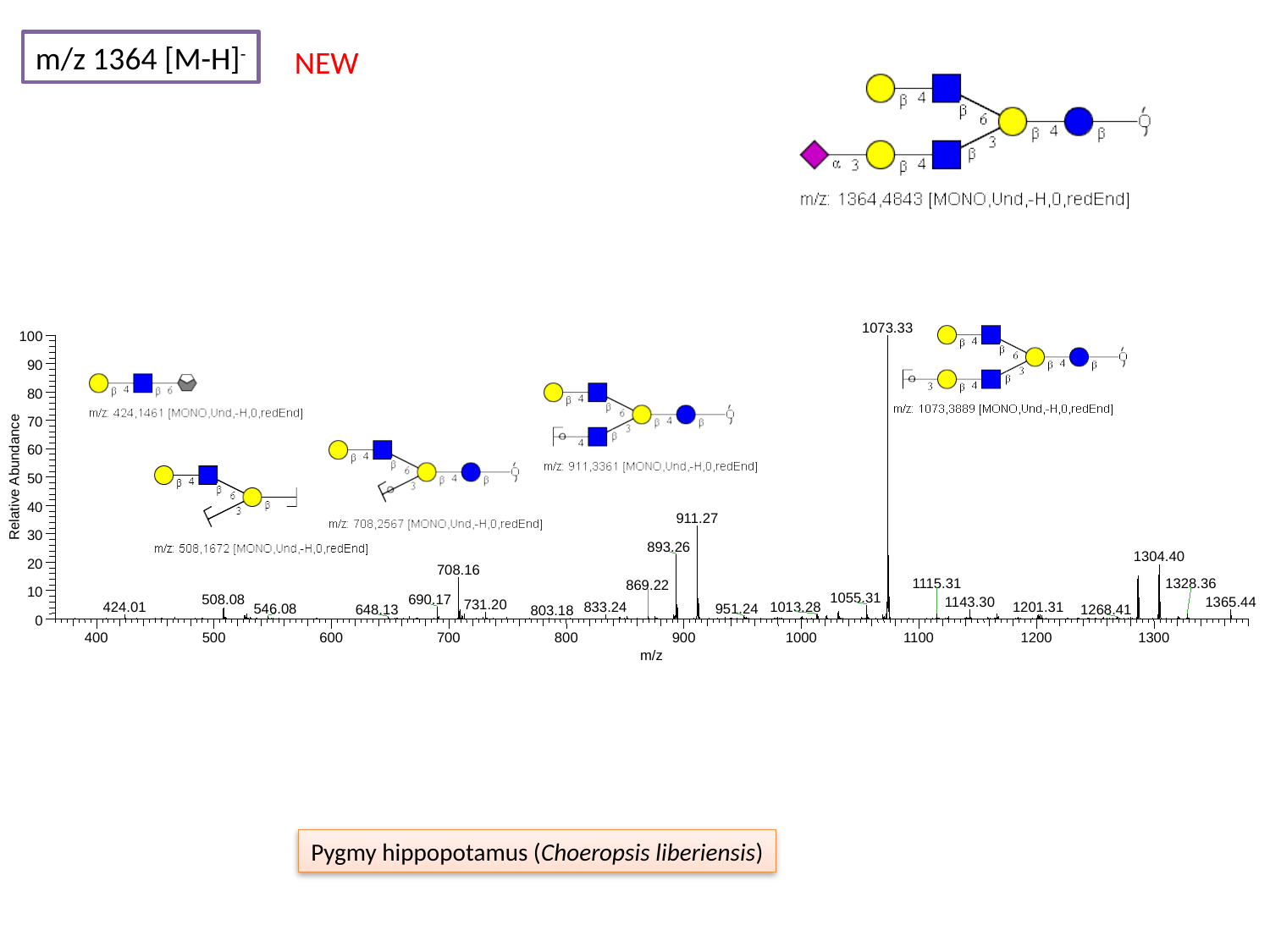

m/z 1364 [M-H]^-^
NEW
1073.33
100
90
80
70
60
Relative Abundance
50
40
911.27
30
893.26
1304.40
20
708.16
1328.36
1115.31
869.22
10
1055.31
690.17
508.08
1143.30
1365.44
731.20
1013.28
833.24
1201.31
424.01
546.08
951.24
648.13
1268.41
803.18
0
400
500
600
700
800
900
1000
1100
1200
1300
m/z
Pygmy hippopotamus (Choeropsis liberiensis)

### Slide 123
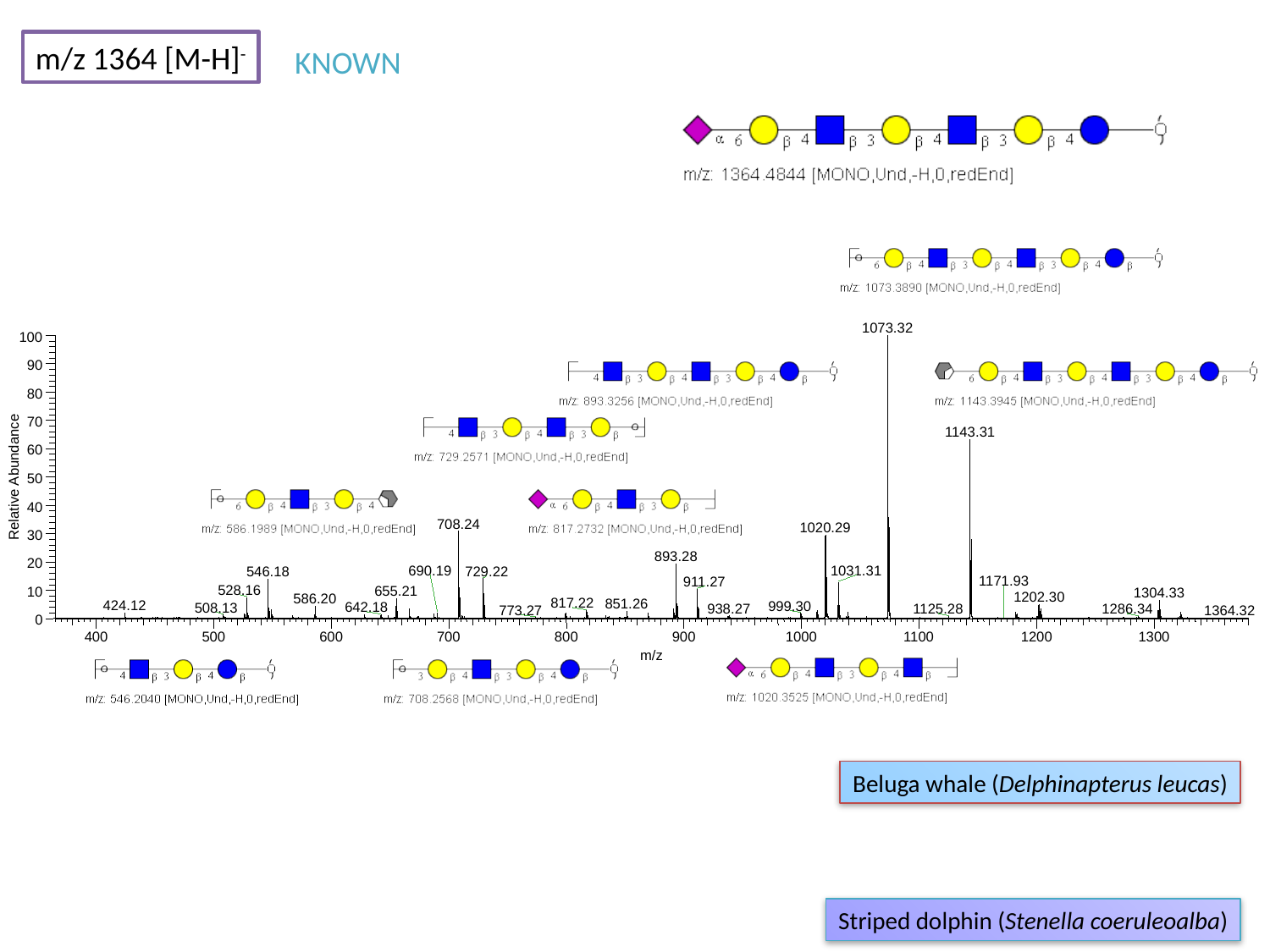

m/z 1364 [M-H]^-^
KNOWN
1073.32
100
90
80
70
1143.31
60
Relative Abundance
50
40
708.24
1020.29
30
893.28
20
1031.31
690.19
729.22
546.18
1171.93
911.27
528.16
655.21
10
1304.33
1202.30
586.20
817.22
851.26
424.12
999.30
642.18
508.13
938.27
1125.28
1286.34
773.27
1364.32
0
400
500
600
700
800
900
1000
1100
1200
1300
m/z
Beluga whale (Delphinapterus leucas)
Striped dolphin (Stenella coeruleoalba)

### Slide 124
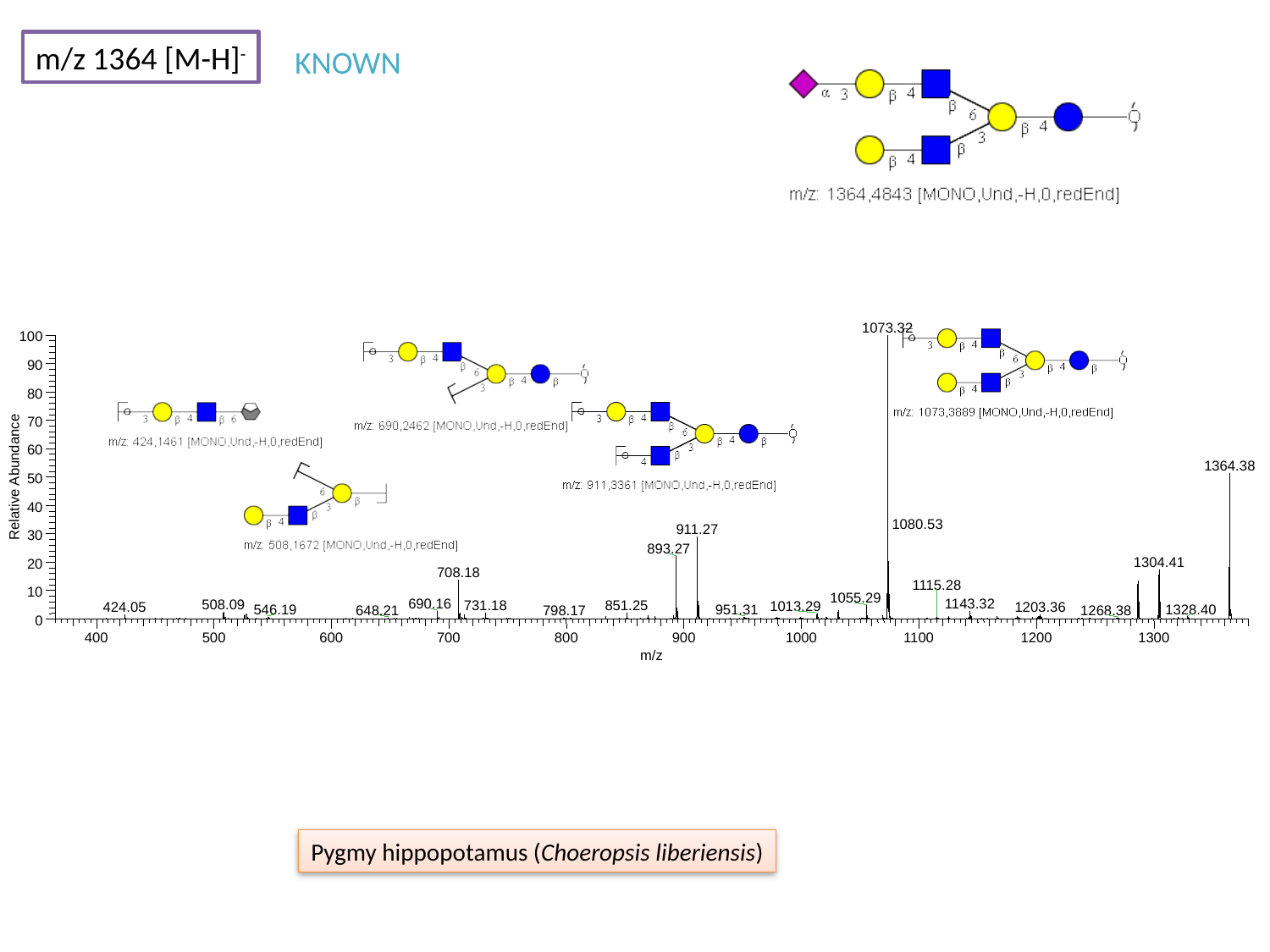

m/z 1364 [M-H]^-^
KNOWN
1073.32
100
90
80
70
60
1364.38
Relative Abundance
50
40
1080.53
911.27
30
893.27
1304.41
20
708.18
1115.28
10
1055.29
1143.32
690.16
508.09
731.18
851.25
1013.29
424.05
1203.36
1328.40
546.19
951.31
648.21
1268.38
798.17
0
400
500
600
700
800
900
1000
1100
1200
1300
m/z
Pygmy hippopotamus (Choeropsis liberiensis)

### Slide 125
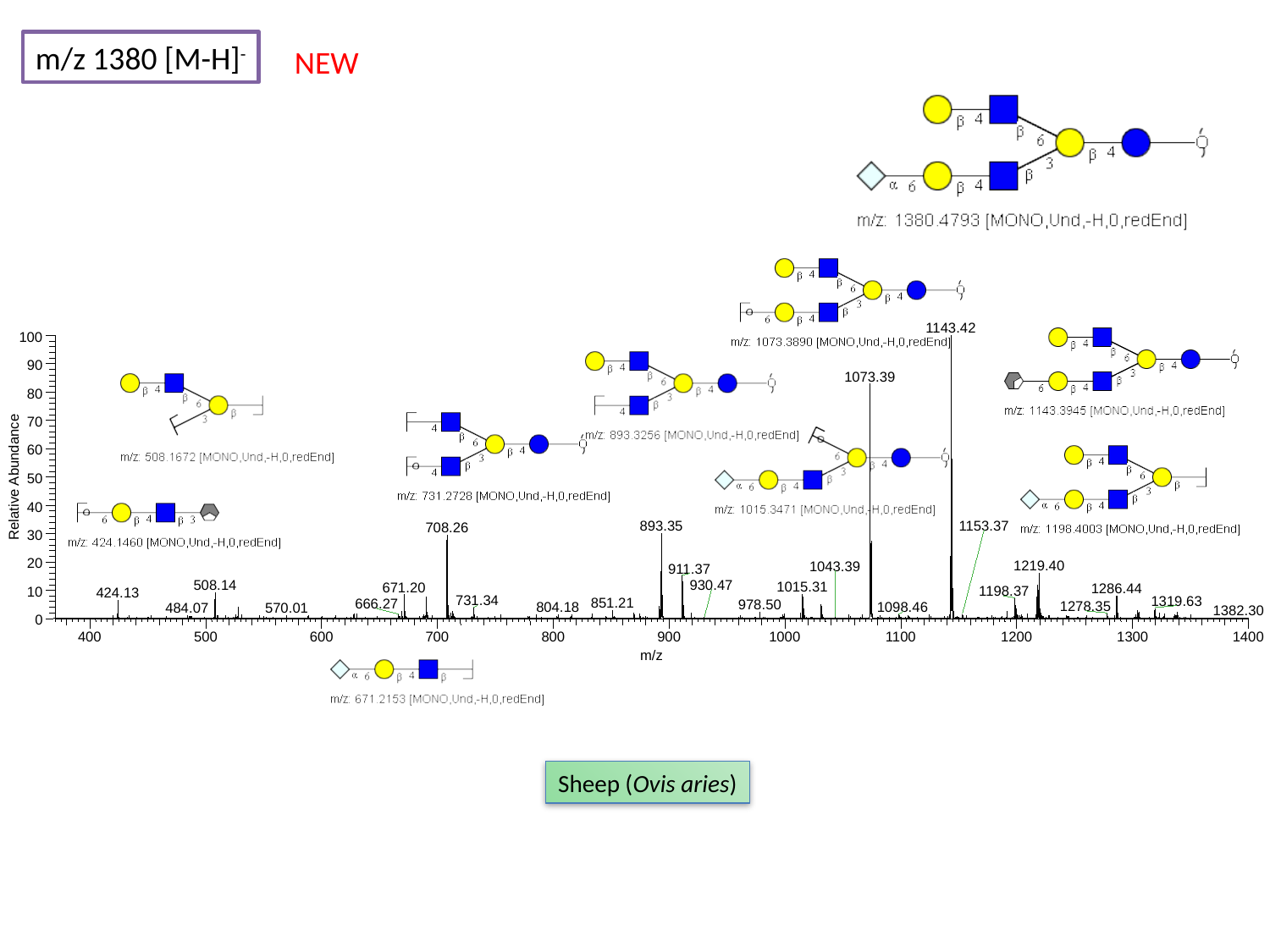

m/z 1380 [M-H]^-^
NEW
1143.42
100
90
1073.39
80
70
60
Relative Abundance
50
40
1153.37
893.35
708.26
30
20
1219.40
1043.39
911.37
930.47
508.14
1015.31
671.20
1286.44
1198.37
10
424.13
731.34
1319.63
851.21
666.27
978.50
1278.35
804.18
1098.46
484.07
570.01
1382.30
0
400
500
600
700
800
900
1000
1100
1200
1300
1400
m/z
Sheep (Ovis aries)

### Slide 126
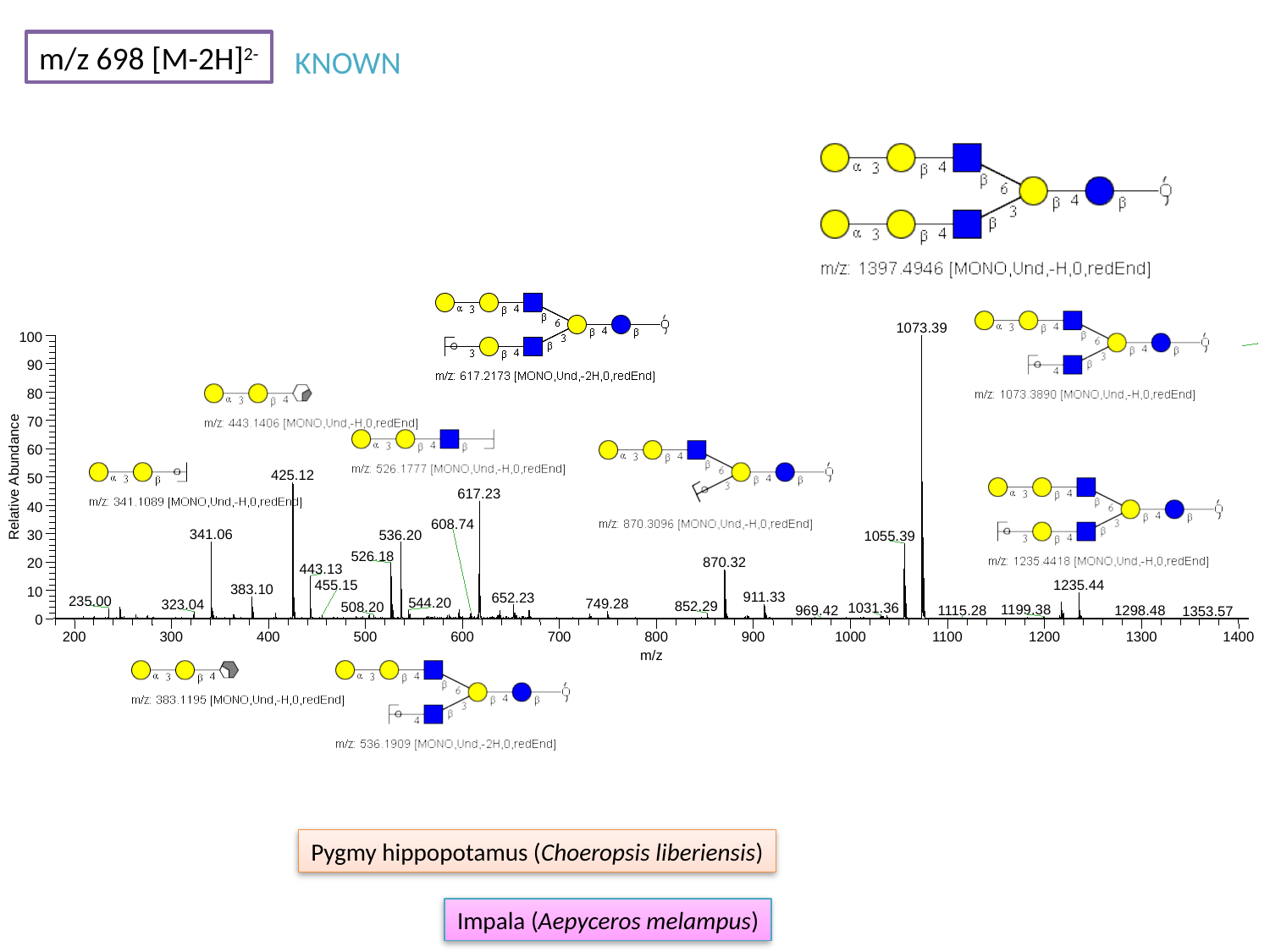

m/z 698 [M-2H]^2-^
KNOWN
1073.39
100
90
80
70
60
425.12
Relative Abundance
50
617.23
40
608.74
341.06
536.20
30
1055.39
526.18
870.32
20
443.13
455.15
1235.44
383.10
10
911.33
652.23
235.00
544.20
749.28
323.04
852.29
508.20
1031.36
1199.38
969.42
1298.48
1115.28
1353.57
0
200
300
400
500
600
700
800
900
1000
1100
1200
1300
1400
m/z
Pygmy hippopotamus (Choeropsis liberiensis)
Impala (Aepyceros melampus)

### Slide 127
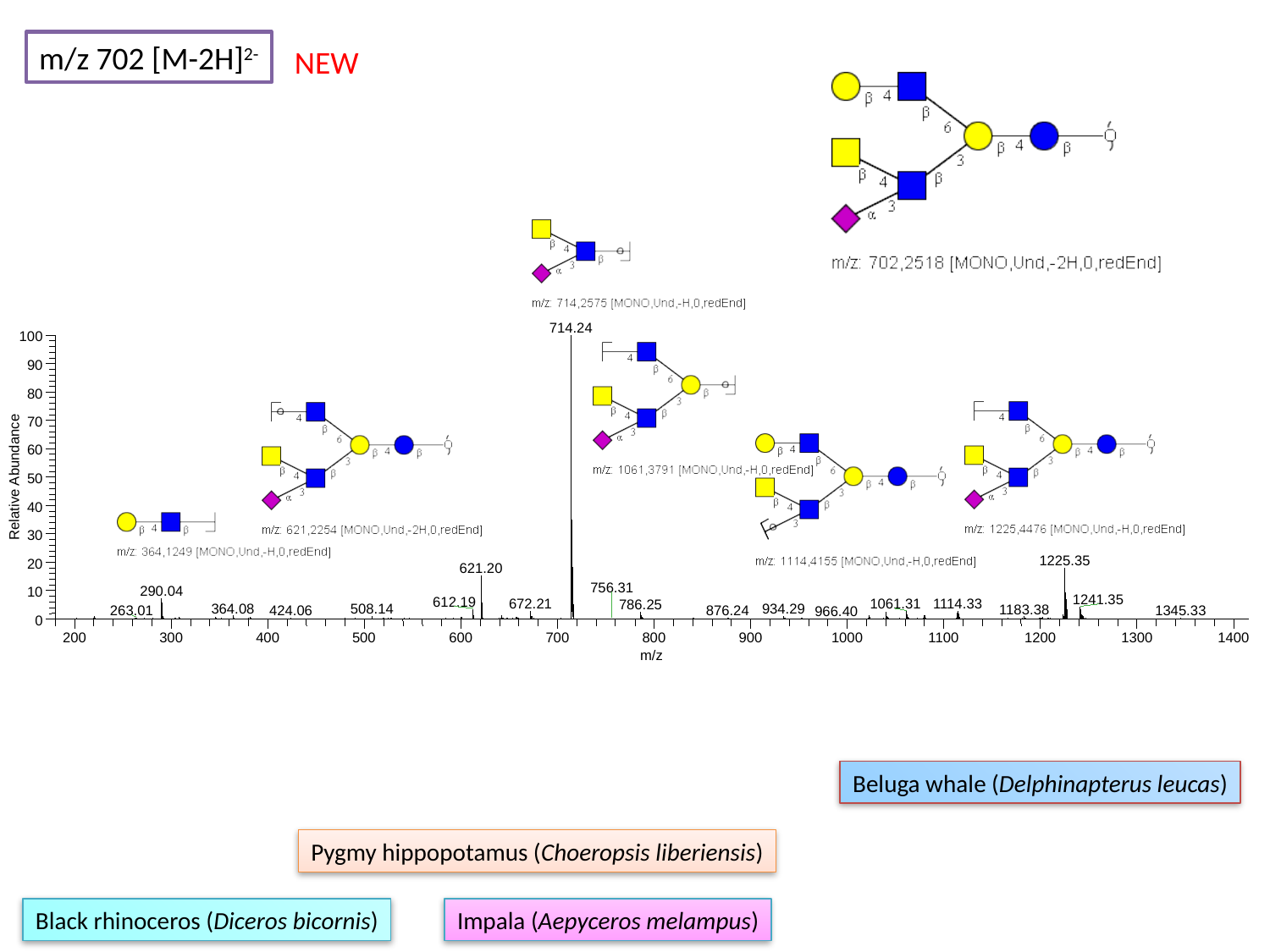

m/z 702 [M-2H]^2-^
NEW
714.24
100
90
80
70
60
Relative Abundance
50
40
30
1225.35
20
621.20
756.31
290.04
10
1241.35
612.19
672.21
1114.33
1061.31
786.25
364.08
934.29
508.14
1183.38
263.01
1345.33
424.06
876.24
966.40
0
200
300
400
500
600
700
800
900
1000
1100
1200
1300
1400
m/z
Beluga whale (Delphinapterus leucas)
Pygmy hippopotamus (Choeropsis liberiensis)
Black rhinoceros (Diceros bicornis)
Impala (Aepyceros melampus)

### Slide 128
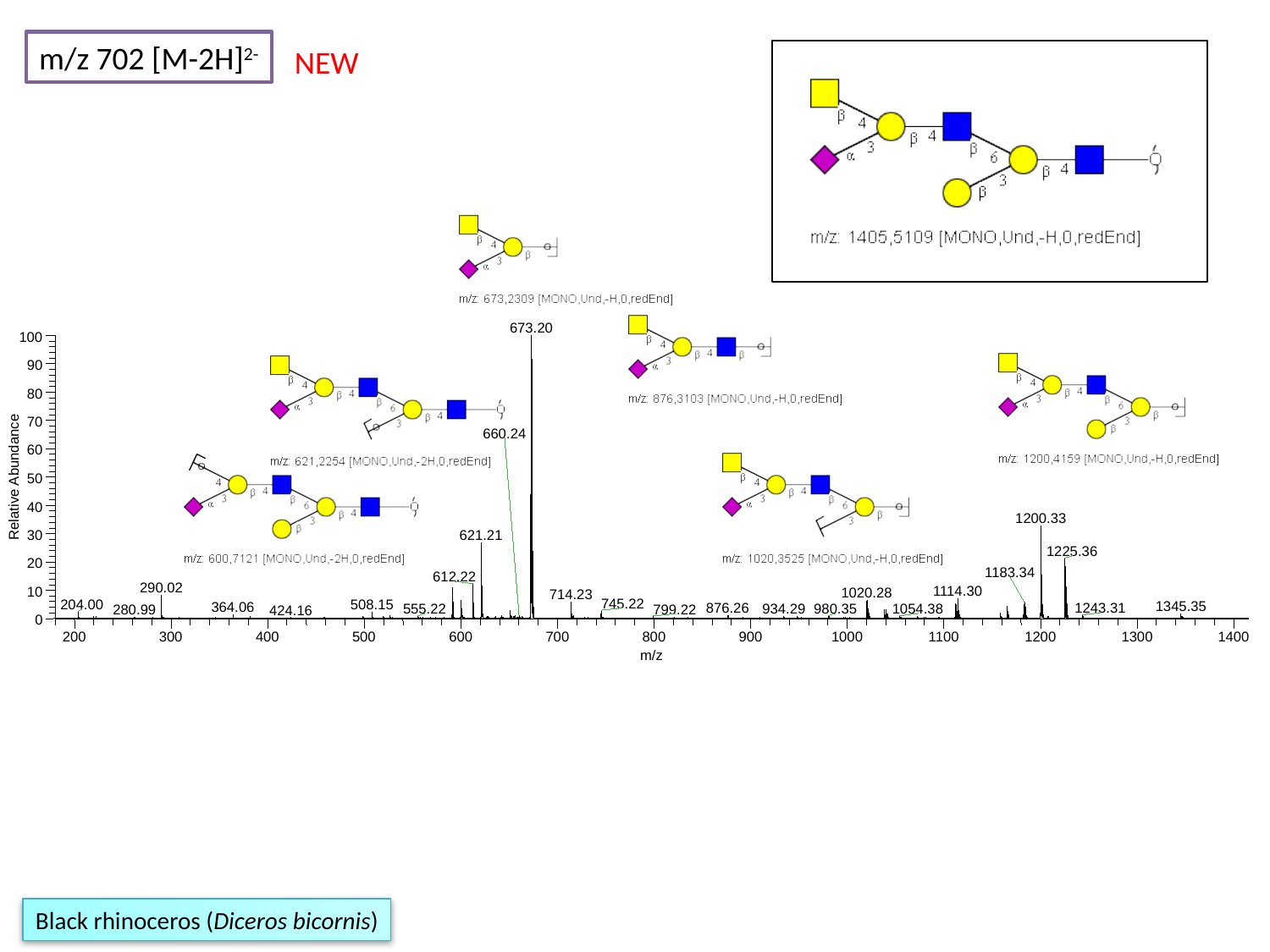

m/z 702 [M-2H]^2-^
NEW
673.20
100
90
80
70
660.24
60
Relative Abundance
50
40
1200.33
30
621.21
1225.36
20
1183.34
612.22
290.02
1114.30
10
1020.28
714.23
745.22
204.00
508.15
1345.35
364.06
876.26
1243.31
934.29
980.35
555.22
1054.38
799.22
280.99
424.16
0
200
300
400
500
600
700
800
900
1000
1100
1200
1300
1400
m/z
Black rhinoceros (Diceros bicornis)

### Slide 129
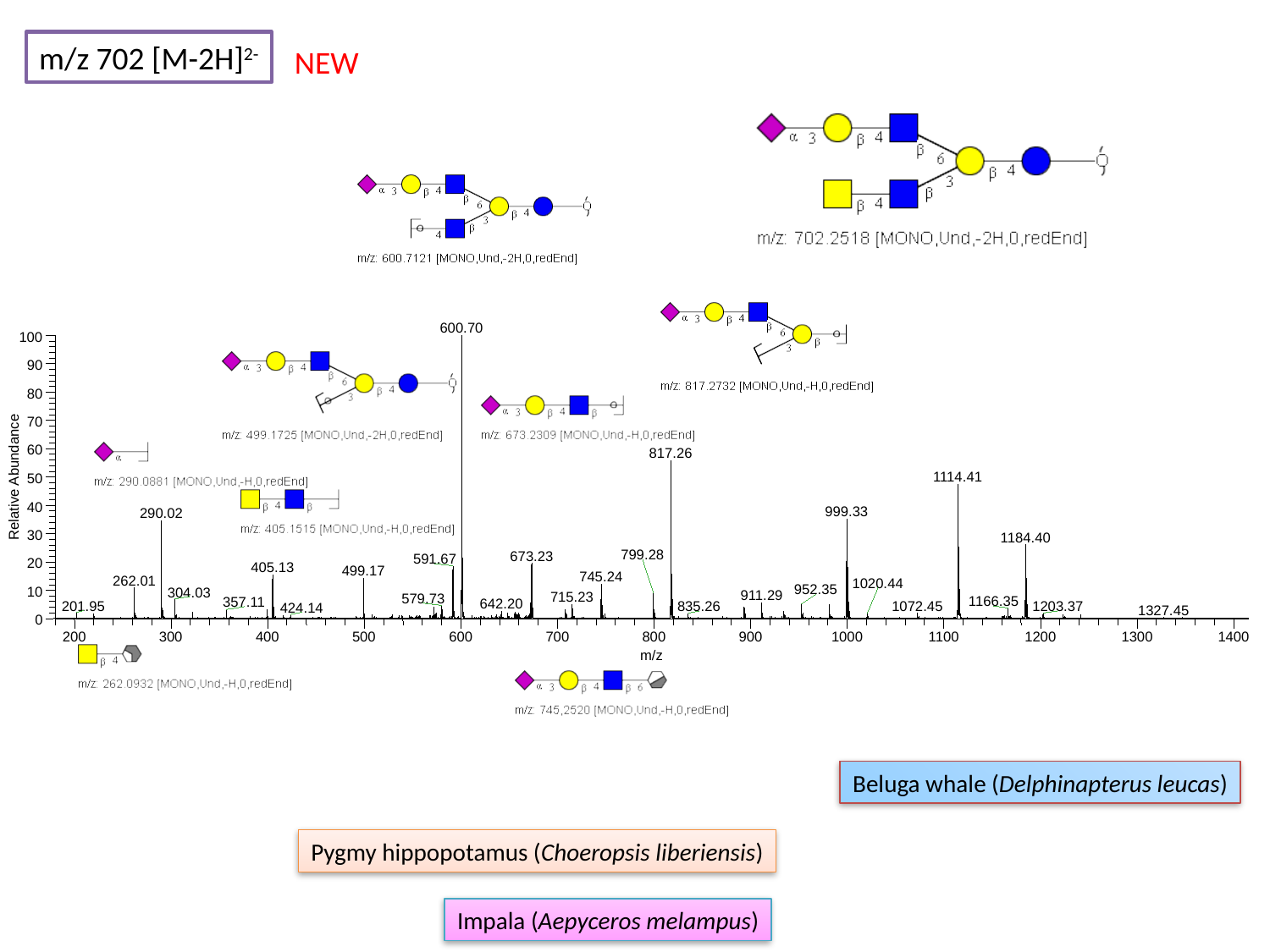

m/z 702 [M-2H]^2-^
NEW
600.70
100
90
80
70
60
817.26
Relative Abundance
1114.41
50
40
999.33
290.02
30
1184.40
799.28
673.23
591.67
20
405.13
499.17
745.24
262.01
1020.44
952.35
10
304.03
911.29
715.23
579.73
1166.35
357.11
642.20
201.95
1072.45
1203.37
835.26
424.14
1327.45
0
200
300
400
500
600
700
800
900
1000
1100
1200
1300
1400
m/z
Beluga whale (Delphinapterus leucas)
Pygmy hippopotamus (Choeropsis liberiensis)
Impala (Aepyceros melampus)

### Slide 130
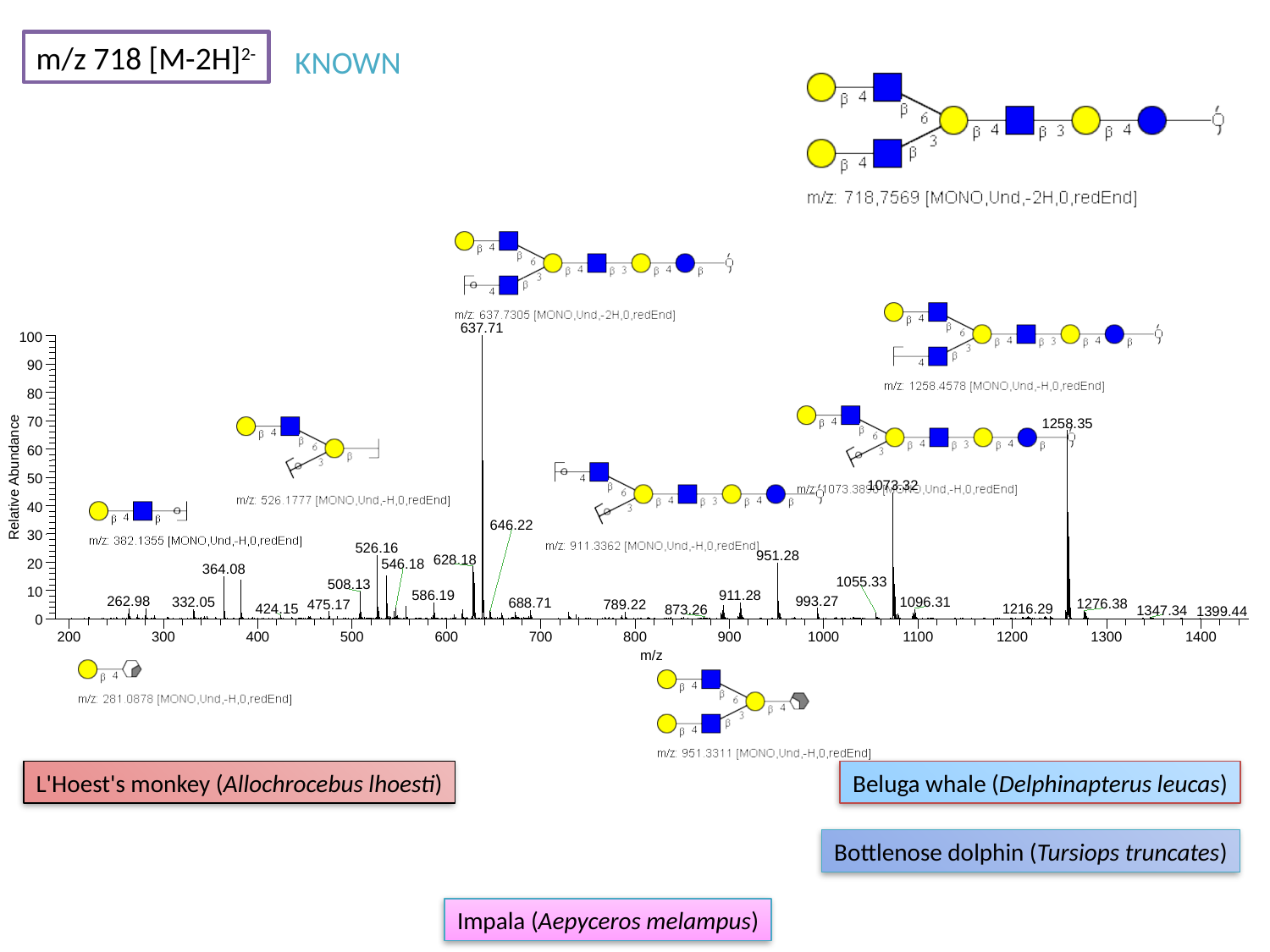

m/z 718 [M-2H]^2-^
KNOWN
637.71
100
90
80
70
1258.35
60
Relative Abundance
50
1073.32
40
646.22
30
526.16
951.28
628.18
20
546.18
364.08
1055.33
508.13
10
586.19
911.28
993.27
262.98
1096.31
332.05
688.71
1276.38
475.17
789.22
424.15
1216.29
873.26
1347.34
1399.44
0
200
300
400
500
600
700
800
900
1000
1100
1200
1300
1400
m/z
L'Hoest's monkey (Allochrocebus lhoesti)
Beluga whale (Delphinapterus leucas)
Bottlenose dolphin (Tursiops truncates)
Impala (Aepyceros melampus)

### Slide 131
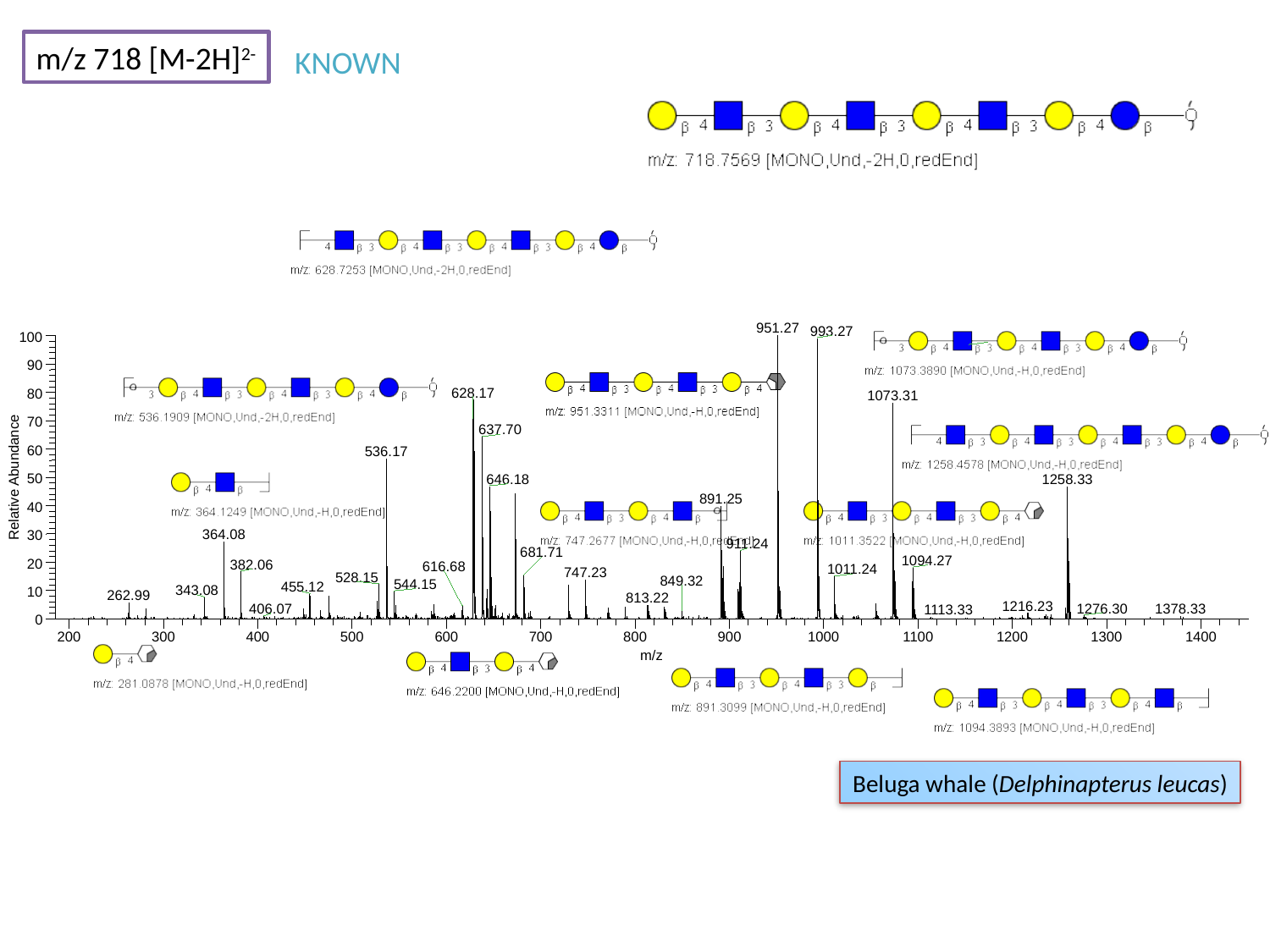

m/z 718 [M-2H]^2-^
KNOWN
951.27
993.27
100
90
628.17
80
1073.31
70
637.70
60
536.17
Relative Abundance
50
646.18
1258.33
891.25
40
364.08
30
911.24
681.71
1094.27
20
382.06
616.68
1011.24
747.23
528.15
849.32
544.15
455.12
343.08
10
262.99
813.22
1216.23
1276.30
0
200
300
400
500
600
700
800
900
1000
1100
1200
1300
1400
m/z
1378.33
406.07
1113.33
Beluga whale (Delphinapterus leucas)

### Slide 132
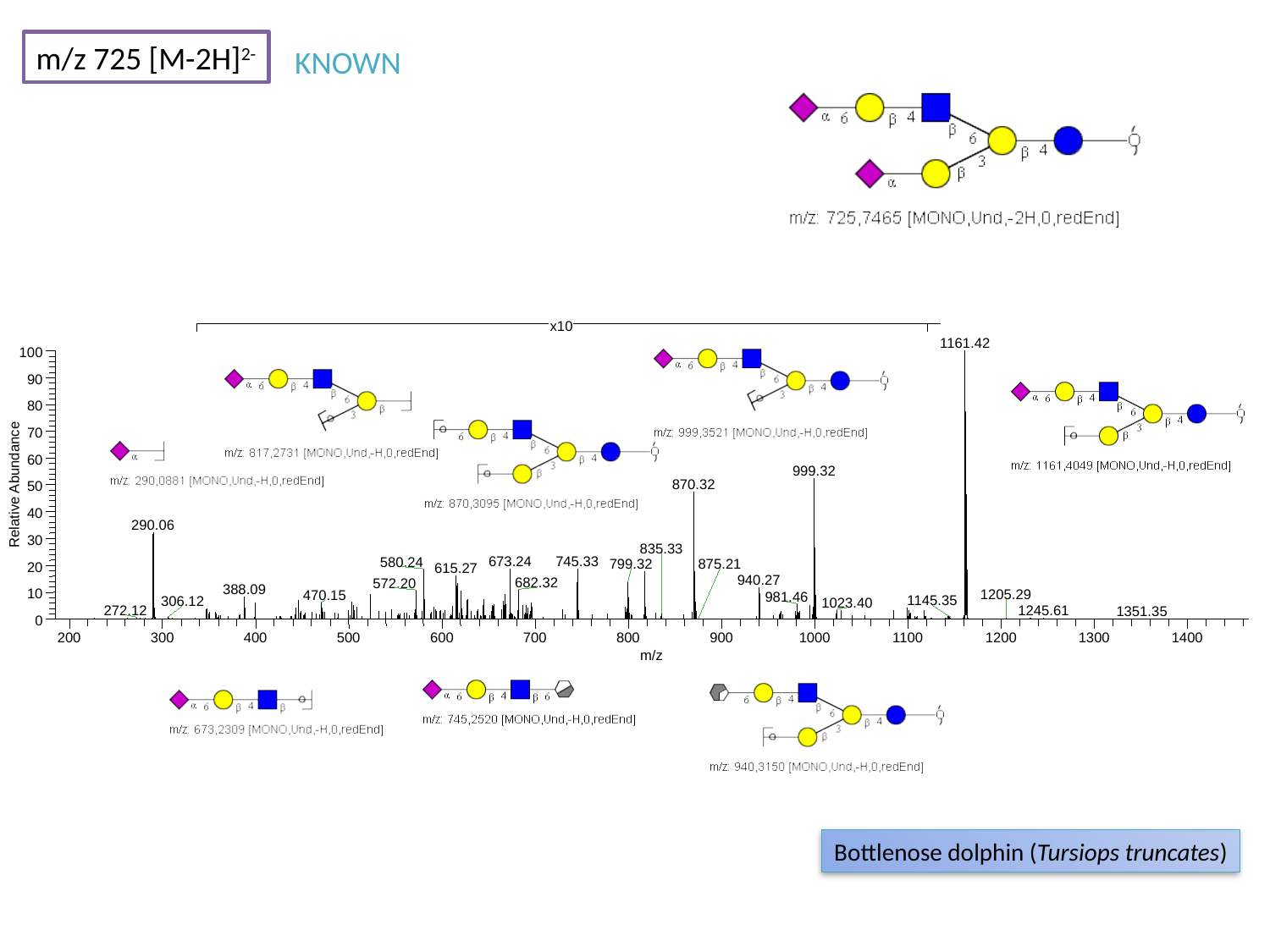

m/z 725 [M-2H]^2-^
KNOWN
x10
1161.42
100
90
80
70
60
999.32
Relative Abundance
870.32
50
40
290.06
30
835.33
673.24
745.33
580.24
875.21
799.32
20
615.27
940.27
682.32
572.20
388.09
10
1205.29
470.15
981.46
1145.35
306.12
1023.40
272.12
1245.61
1351.35
0
200
300
400
500
600
700
800
900
1000
1100
1200
1300
1400
m/z
Bottlenose dolphin (Tursiops truncates)

### Slide 133
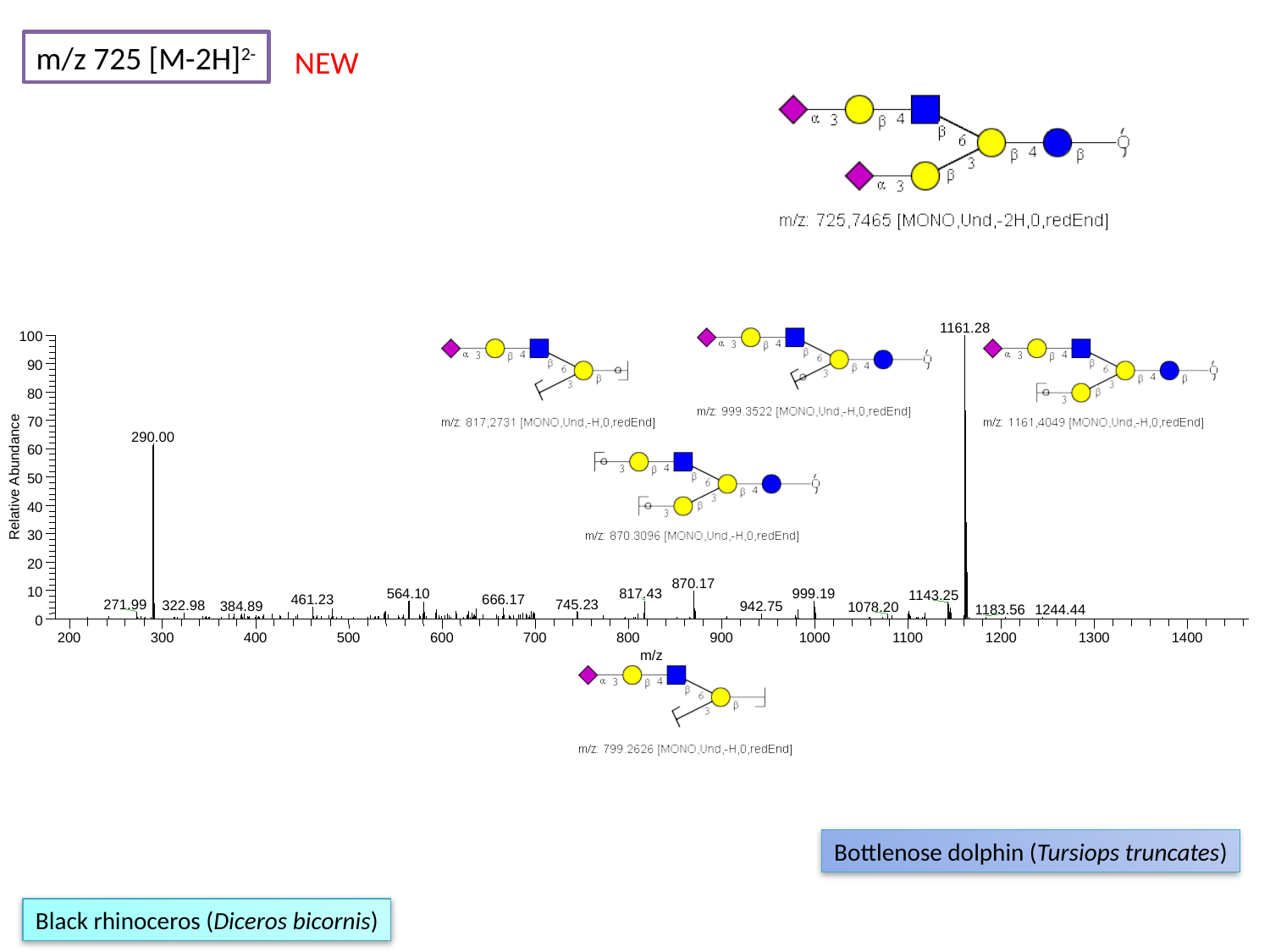

m/z 725 [M-2H]^2-^
NEW
1161.28
100
90
80
70
290.00
60
Relative Abundance
50
40
30
20
870.17
10
817.43
564.10
999.19
1143.25
461.23
666.17
745.23
271.99
322.98
384.89
942.75
1078.20
1183.56
1244.44
0
200
300
400
500
600
700
800
900
1000
1100
1200
1300
1400
m/z
Bottlenose dolphin (Tursiops truncates)
Black rhinoceros (Diceros bicornis)

### Slide 134
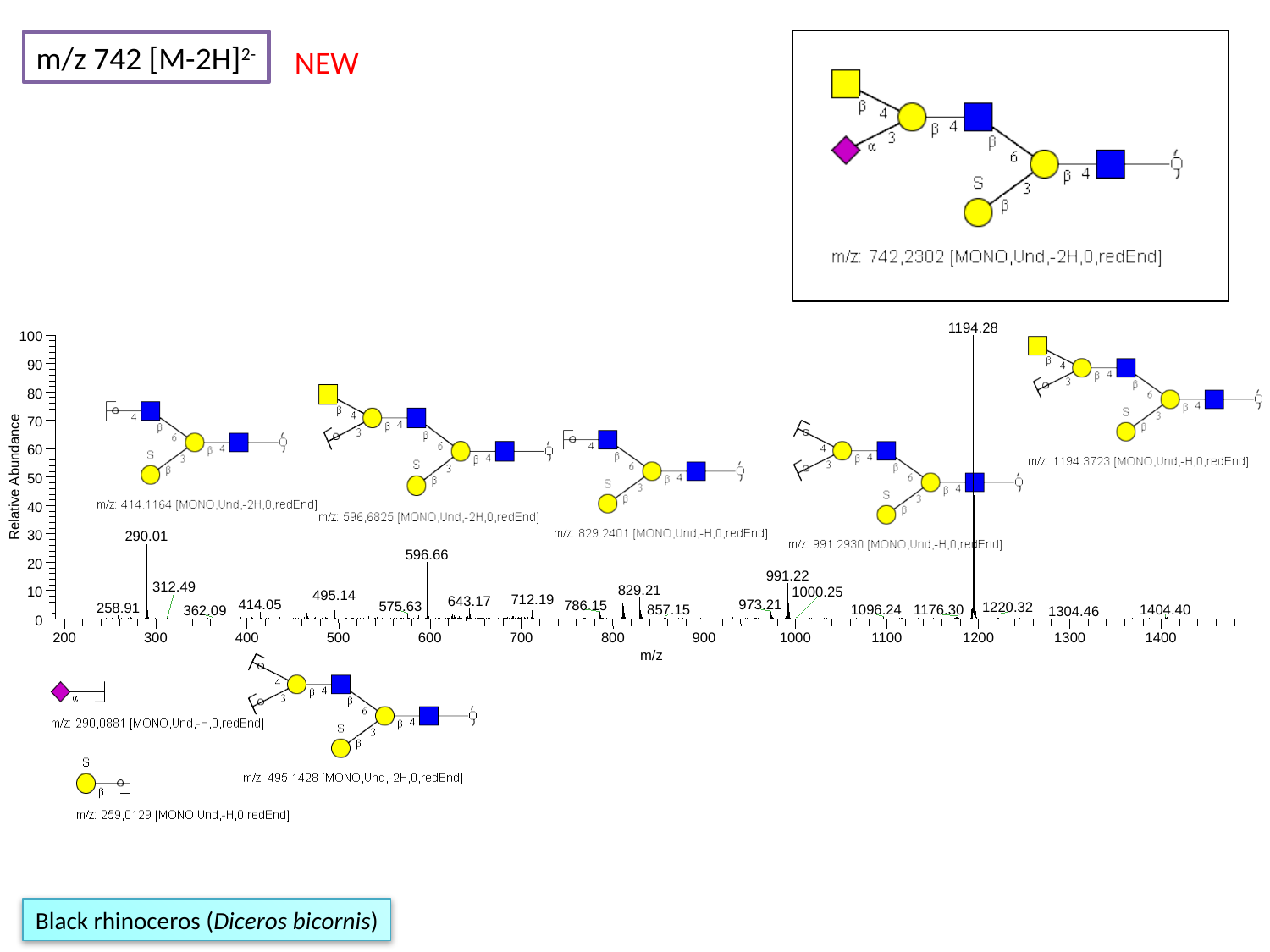

m/z 742 [M-2H]^2-^
NEW
1194.28
100
90
80
70
60
Relative Abundance
50
40
30
290.01
596.66
20
991.22
312.49
829.21
1000.25
10
495.14
712.19
643.17
414.05
973.21
786.15
575.63
1220.32
258.91
857.15
1176.30
1404.40
1096.24
362.09
1304.46
0
200
300
400
500
600
700
800
900
1000
1100
1200
1300
1400
m/z
Black rhinoceros (Diceros bicornis)

### Slide 135
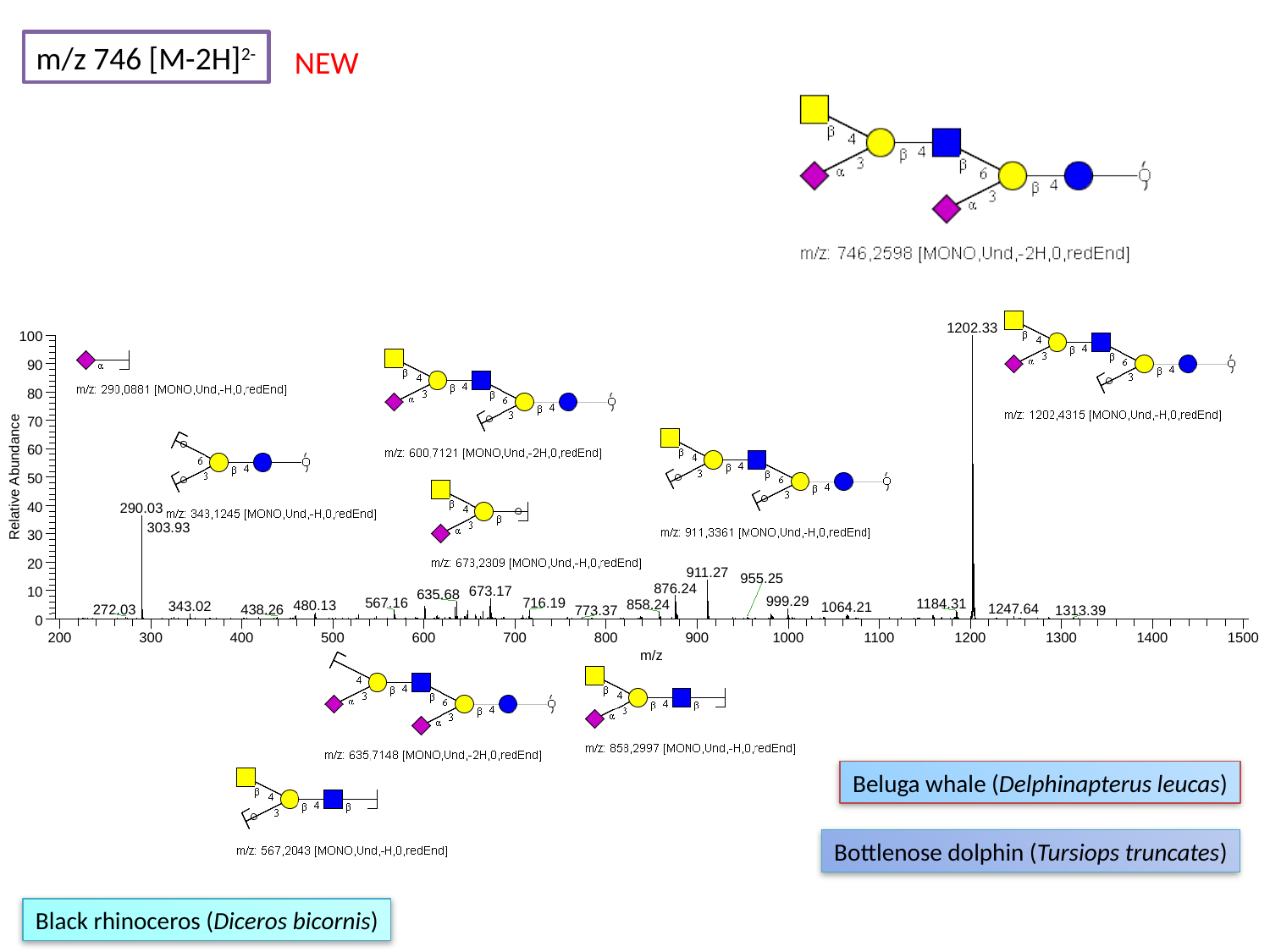

m/z 746 [M-2H]^2-^
NEW
1202.33
100
90
80
70
60
Relative Abundance
50
40
290.03
303.93
30
20
911.27
955.25
876.24
673.17
10
635.68
999.29
716.19
567.16
1184.31
858.24
480.13
343.02
1064.21
1247.64
272.03
438.26
773.37
1313.39
0
200
300
400
500
600
700
800
900
1000
1100
1200
1300
1400
1500
m/z
Beluga whale (Delphinapterus leucas)
Bottlenose dolphin (Tursiops truncates)
Black rhinoceros (Diceros bicornis)

### Slide 136
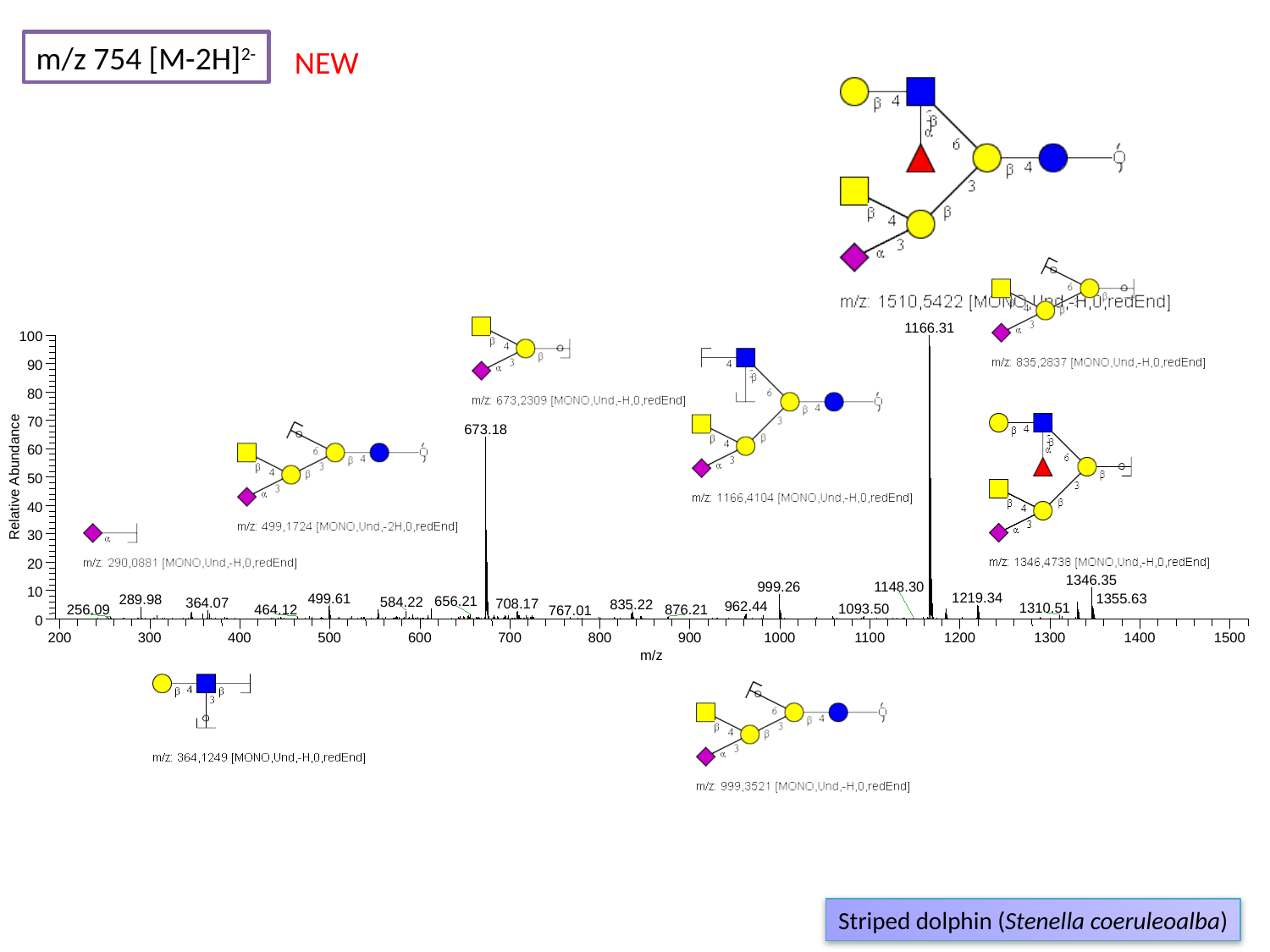

m/z 754 [M-2H]^2-^
NEW
1166.31
100
90
80
70
673.18
60
Relative Abundance
50
40
30
20
1346.35
1148.30
999.26
10
1219.34
1355.63
499.61
289.98
656.21
584.22
364.07
708.17
835.22
962.44
1310.51
1093.50
256.09
464.12
876.21
767.01
0
200
300
400
500
600
700
800
900
1000
1100
1200
1300
1400
1500
m/z
Striped dolphin (Stenella coeruleoalba)

### Slide 137
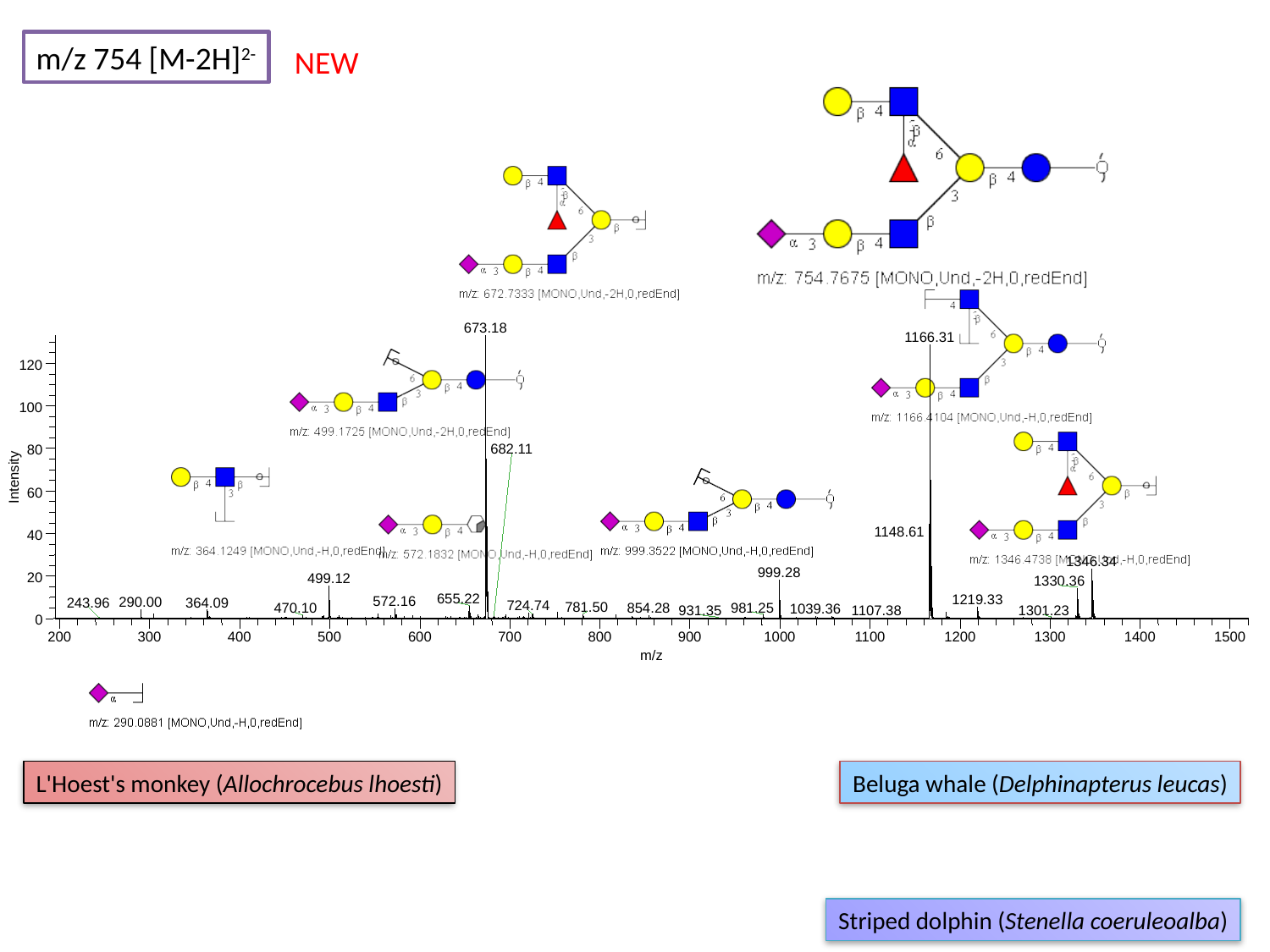

m/z 754 [M-2H]^2-^
NEW
673.18
1166.31
120
100
682.11
80
Intensity
60
1148.61
40
1346.34
999.28
20
499.12
1330.36
655.22
1219.33
572.16
290.00
364.09
243.96
724.74
781.50
981.25
470.10
854.28
1039.36
1301.23
931.35
1107.38
0
200
300
400
500
600
700
800
900
1000
1100
1200
1300
1400
1500
m/z
L'Hoest's monkey (Allochrocebus lhoesti)
Beluga whale (Delphinapterus leucas)
Striped dolphin (Stenella coeruleoalba)

### Slide 138
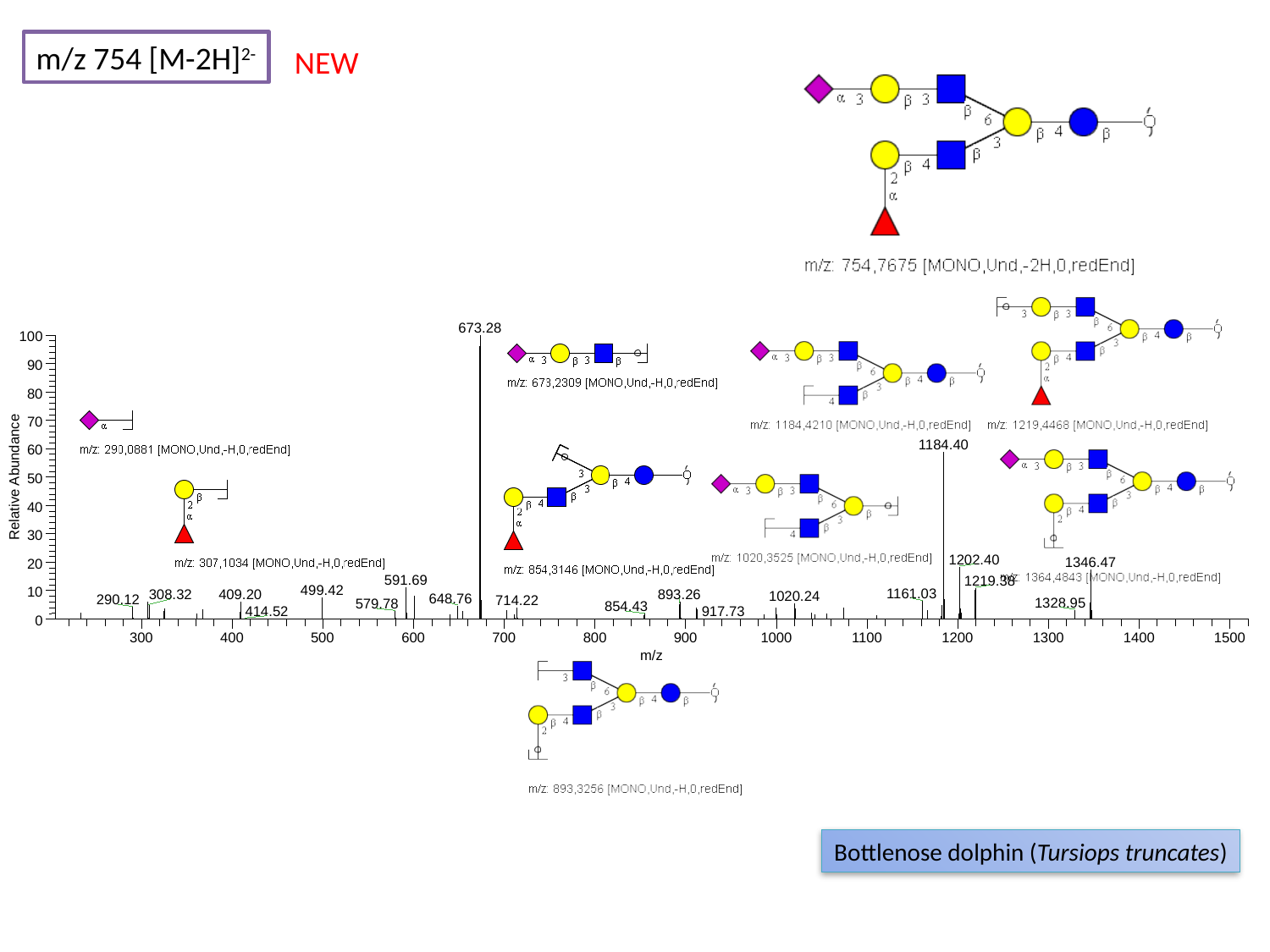

m/z 754 [M-2H]^2-^
NEW
673.28
100
90
80
70
1184.40
60
Relative Abundance
50
40
30
1202.40
1346.47
20
591.69
1219.38
499.42
10
1161.03
308.32
409.20
893.26
1020.24
648.76
290.12
714.22
1328.95
579.78
854.43
414.52
917.73
0
300
400
500
600
700
800
900
1000
1100
1200
1300
1400
1500
m/z
Bottlenose dolphin (Tursiops truncates)

### Slide 139
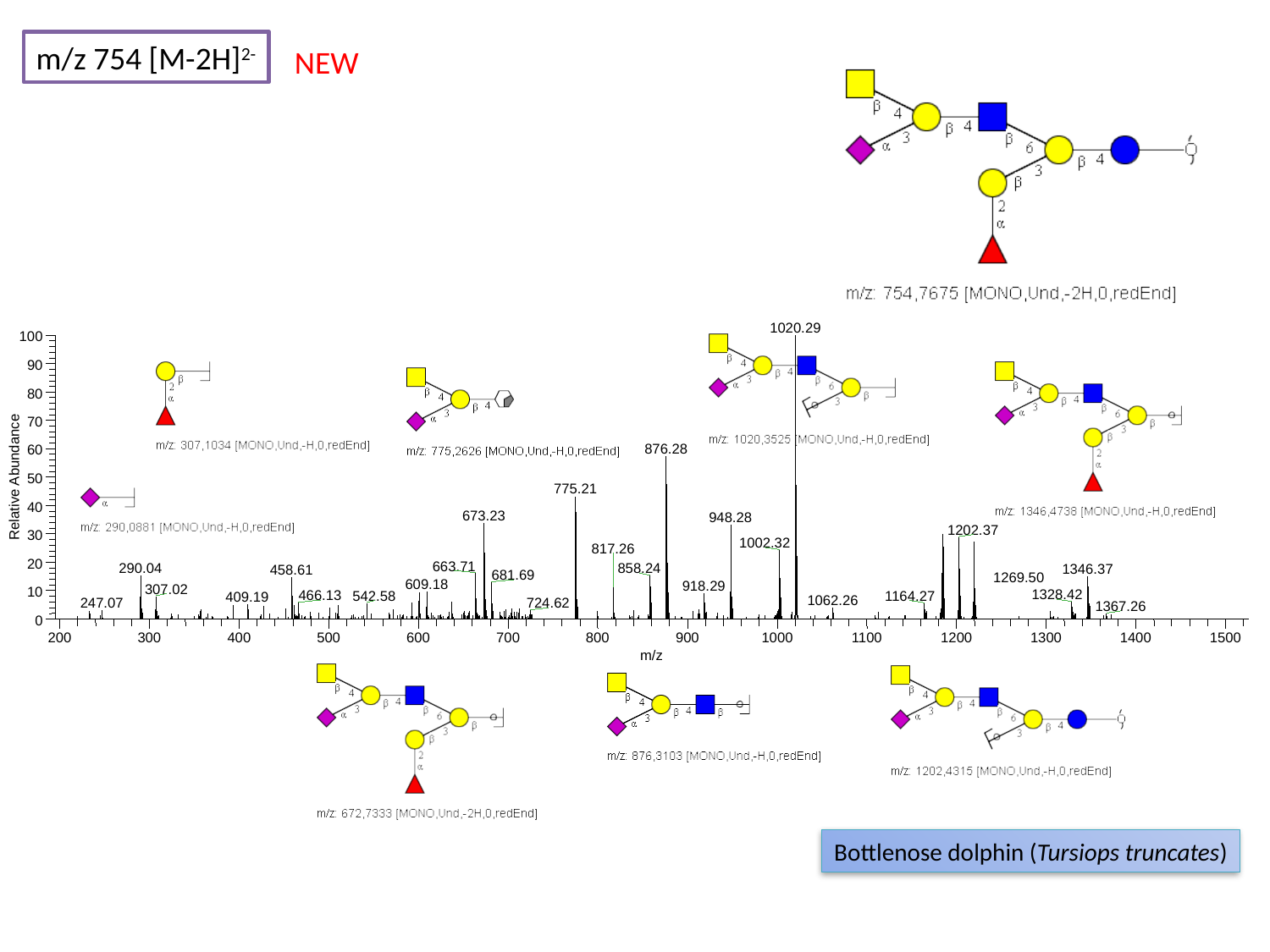

m/z 754 [M-2H]^2-^
NEW
1020.29
100
90
80
70
876.28
60
Relative Abundance
50
775.21
40
673.23
948.28
1202.37
30
1002.32
817.26
20
663.71
290.04
858.24
1346.37
458.61
681.69
609.18
918.29
307.02
10
1328.42
466.13
542.58
1164.27
409.19
1062.26
247.07
724.62
0
200
300
400
500
600
700
800
900
1000
1100
1200
1300
1400
1500
m/z
1269.50
1367.26
Bottlenose dolphin (Tursiops truncates)

### Slide 140
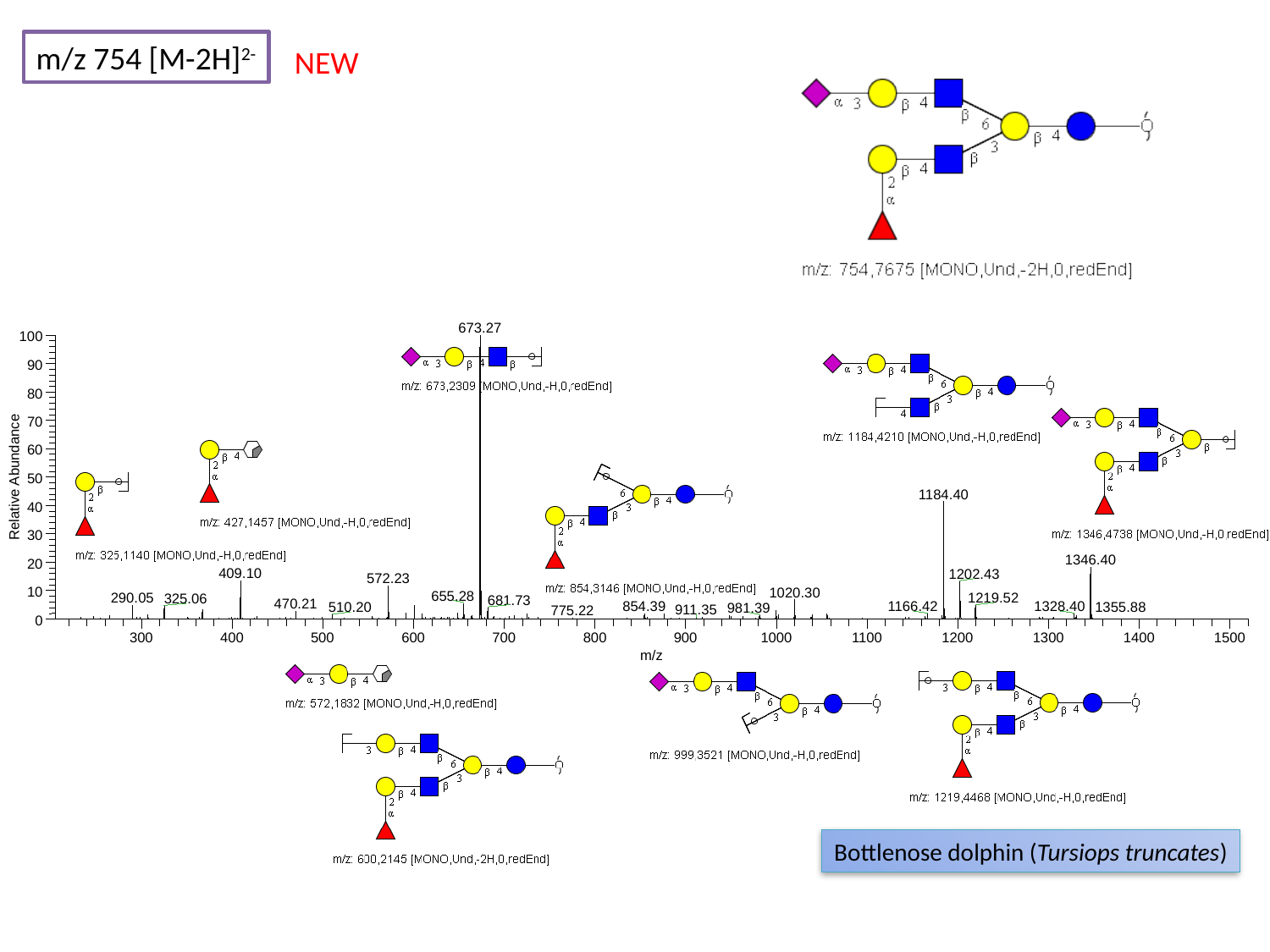

m/z 754 [M-2H]^2-^
NEW
673.27
100
90
80
70
60
Relative Abundance
50
1184.40
40
30
1346.40
20
409.10
1202.43
572.23
10
1020.30
655.28
290.05
1219.52
325.06
681.73
470.21
854.39
1166.42
1328.40
1355.88
510.20
981.39
911.35
775.22
0
300
400
500
600
700
800
900
1000
1100
1200
1300
1400
1500
m/z
Bottlenose dolphin (Tursiops truncates)

### Slide 141
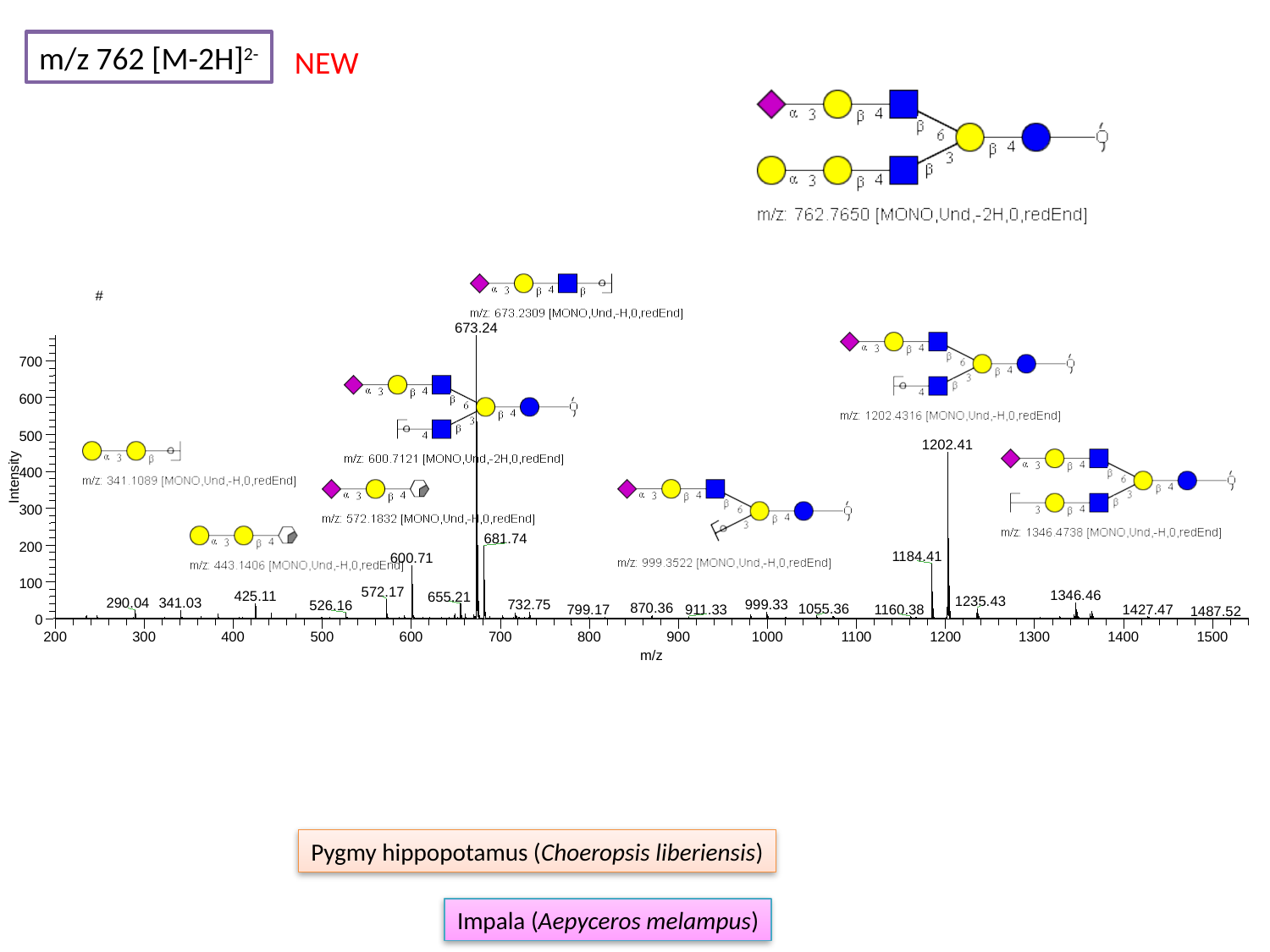

m/z 762 [M-2H]^2-^
NEW
#
673.24
700
600
500
1202.41
400
Intensity
300
681.74
200
1184.41
600.71
100
572.17
1346.46
425.11
655.21
1235.43
341.03
290.04
732.75
999.33
526.16
870.36
1055.36
1160.38
1427.47
911.33
799.17
1487.52
0
200
300
400
500
600
700
800
900
1000
1100
1200
1300
1400
1500
m/z
Pygmy hippopotamus (Choeropsis liberiensis)
Impala (Aepyceros melampus)

### Slide 142
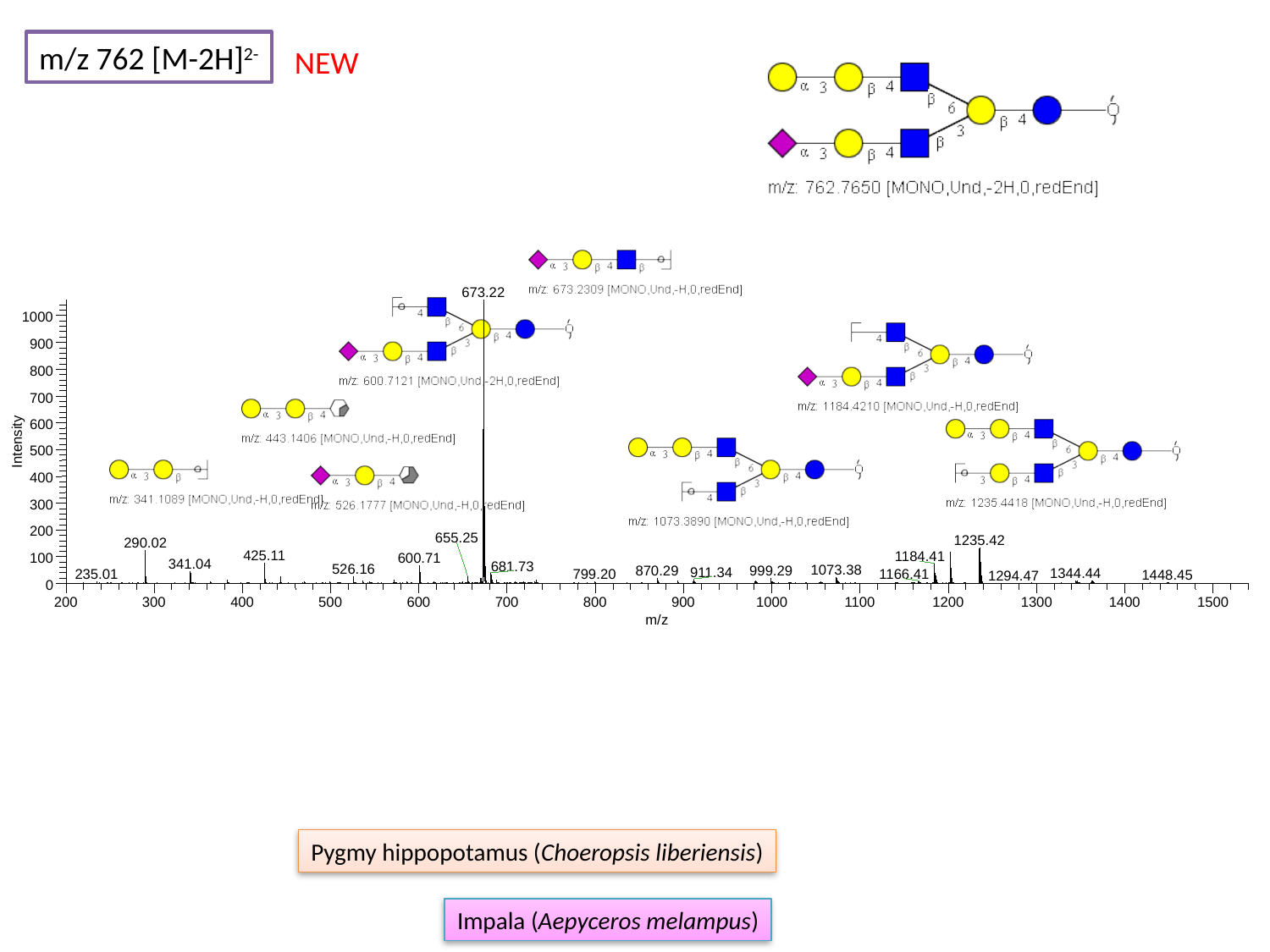

m/z 762 [M-2H]^2-^
NEW
673.22
1000
900
800
700
600
Intensity
500
400
300
200
655.25
1235.42
290.02
425.11
1184.41
600.71
100
341.04
681.73
526.16
1073.38
870.29
999.29
911.34
1344.44
1166.41
235.01
799.20
1448.45
1294.47
0
200
300
400
500
600
700
800
900
1000
1100
1200
1300
1400
1500
m/z
Pygmy hippopotamus (Choeropsis liberiensis)
Impala (Aepyceros melampus)

### Slide 143
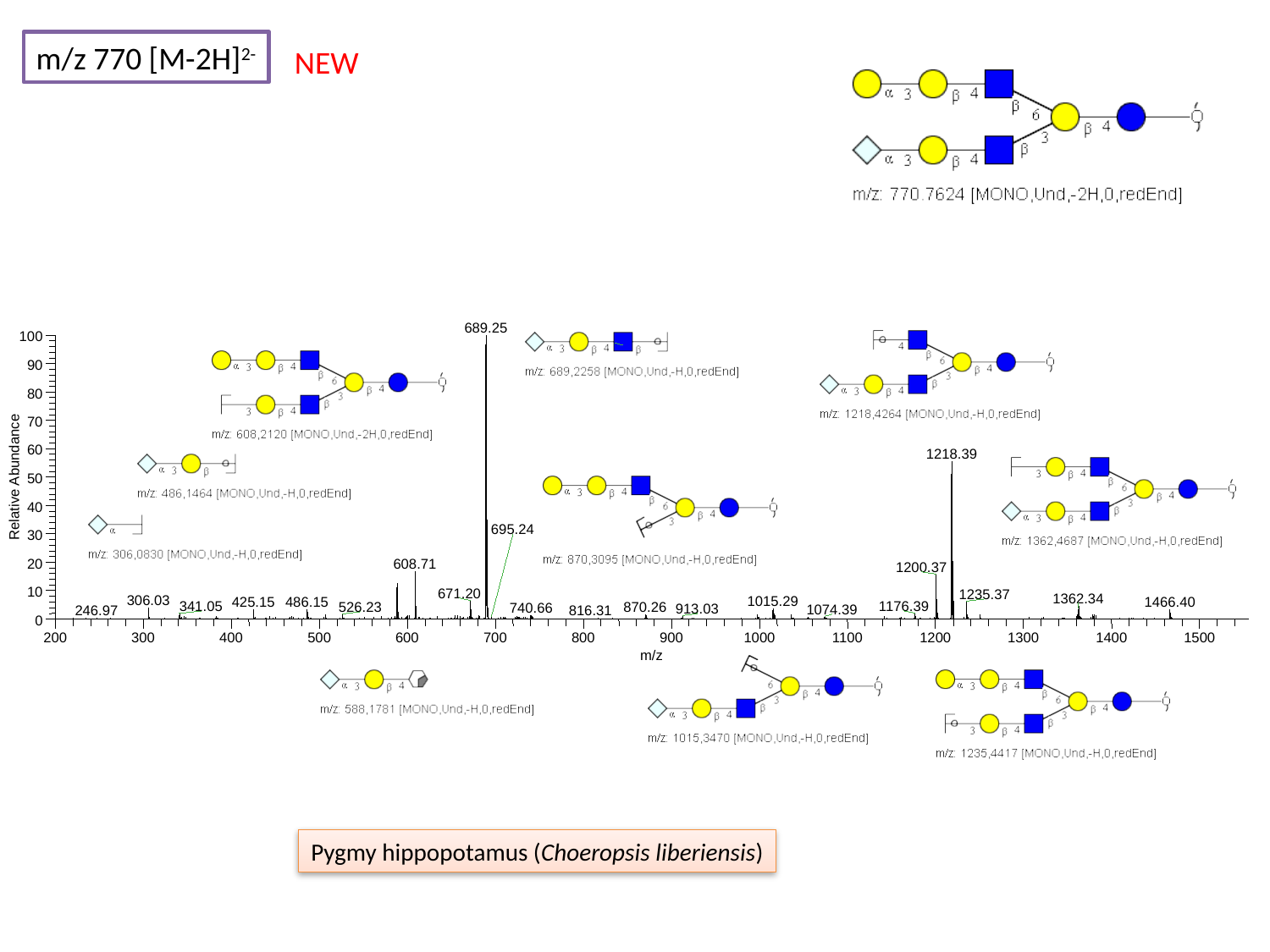

m/z 770 [M-2H]^2-^
NEW
689.25
100
90
80
70
60
1218.39
Relative Abundance
50
40
695.24
30
20
608.71
1200.37
10
671.20
1235.37
1362.34
306.03
1015.29
486.15
1466.40
425.15
1176.39
341.05
526.23
870.26
740.66
913.03
1074.39
246.97
816.31
0
200
300
400
500
600
700
800
900
1000
1100
1200
1300
1400
1500
m/z
Pygmy hippopotamus (Choeropsis liberiensis)

### Slide 144
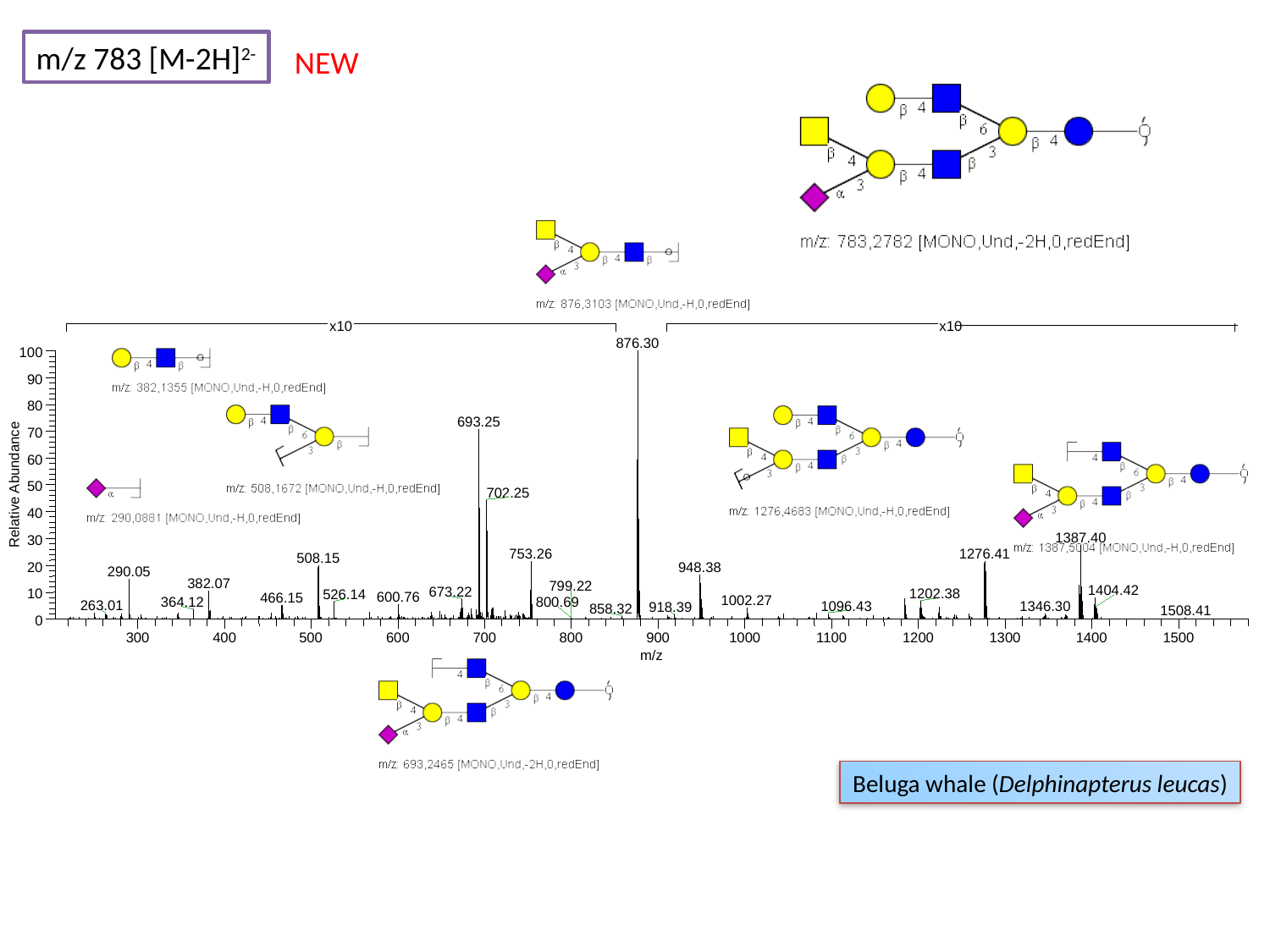

m/z 783 [M-2H]^2-^
NEW
876.30
100
90
80
693.25
70
60
Relative Abundance
50
702.25
40
1387.40
30
1276.41
753.26
508.15
20
948.38
290.05
382.07
799.22
1404.42
673.22
10
1202.38
526.14
600.76
466.15
1002.27
364.12
263.01
1096.43
1346.30
918.39
858.32
1508.41
0
300
400
500
600
700
800
900
1000
1100
1200
1300
1400
1500
m/z
x10
x10
800.69
Beluga whale (Delphinapterus leucas)

### Slide 145
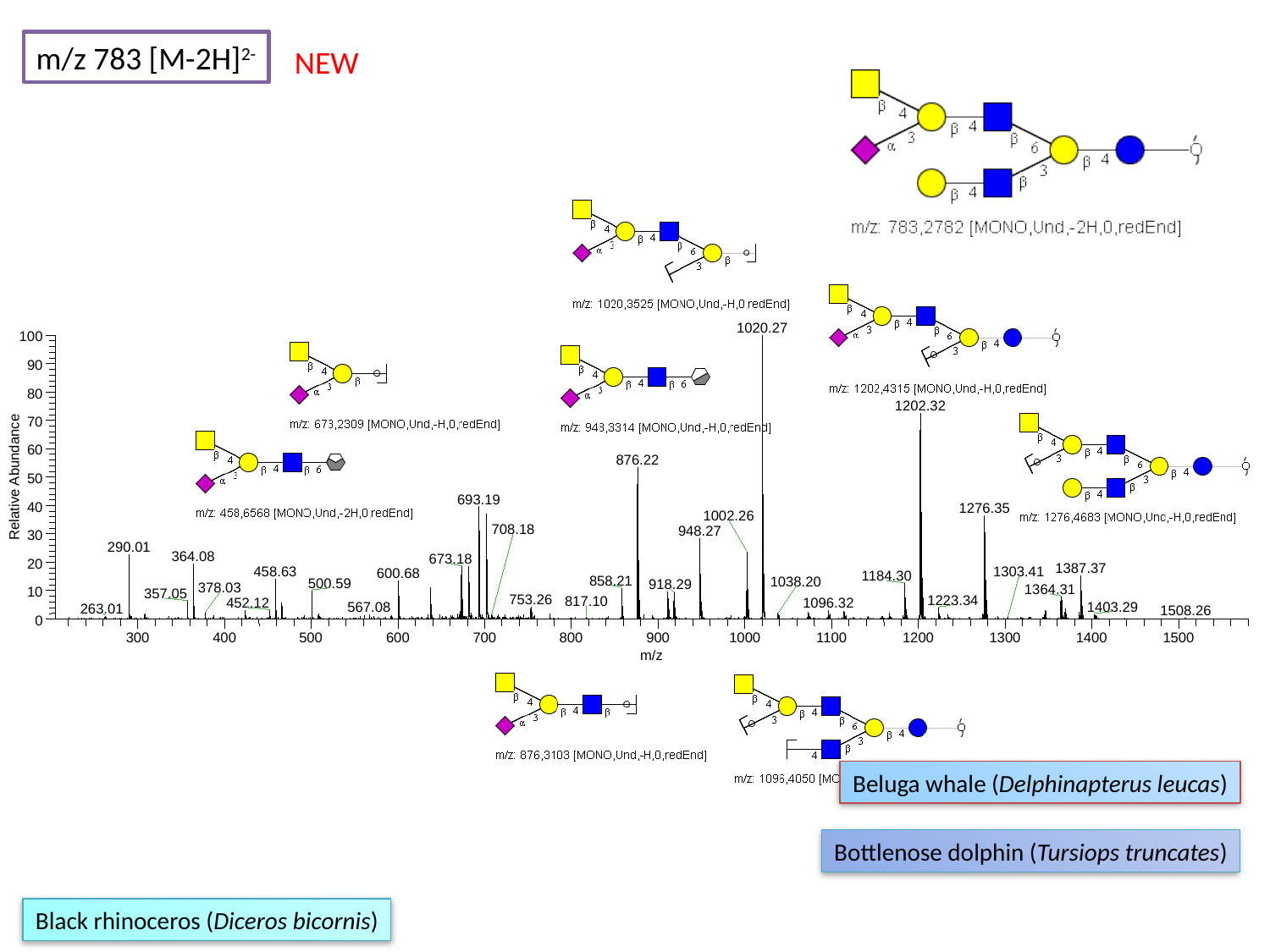

m/z 783 [M-2H]^2-^
NEW
1020.27
100
90
80
1202.32
70
60
876.22
Relative Abundance
50
693.19
40
1276.35
1002.26
948.27
30
290.01
364.08
673.18
20
1387.37
458.63
600.68
1184.30
858.21
500.59
918.29
1364.31
10
357.05
753.26
1223.34
452.12
1096.32
1403.29
0
300
400
500
600
700
800
900
1000
1100
1200
1300
1400
1500
m/z
708.18
1303.41
1038.20
378.03
817.10
567.08
263.01
1508.26
Beluga whale (Delphinapterus leucas)
Bottlenose dolphin (Tursiops truncates)
Black rhinoceros (Diceros bicornis)

### Slide 146
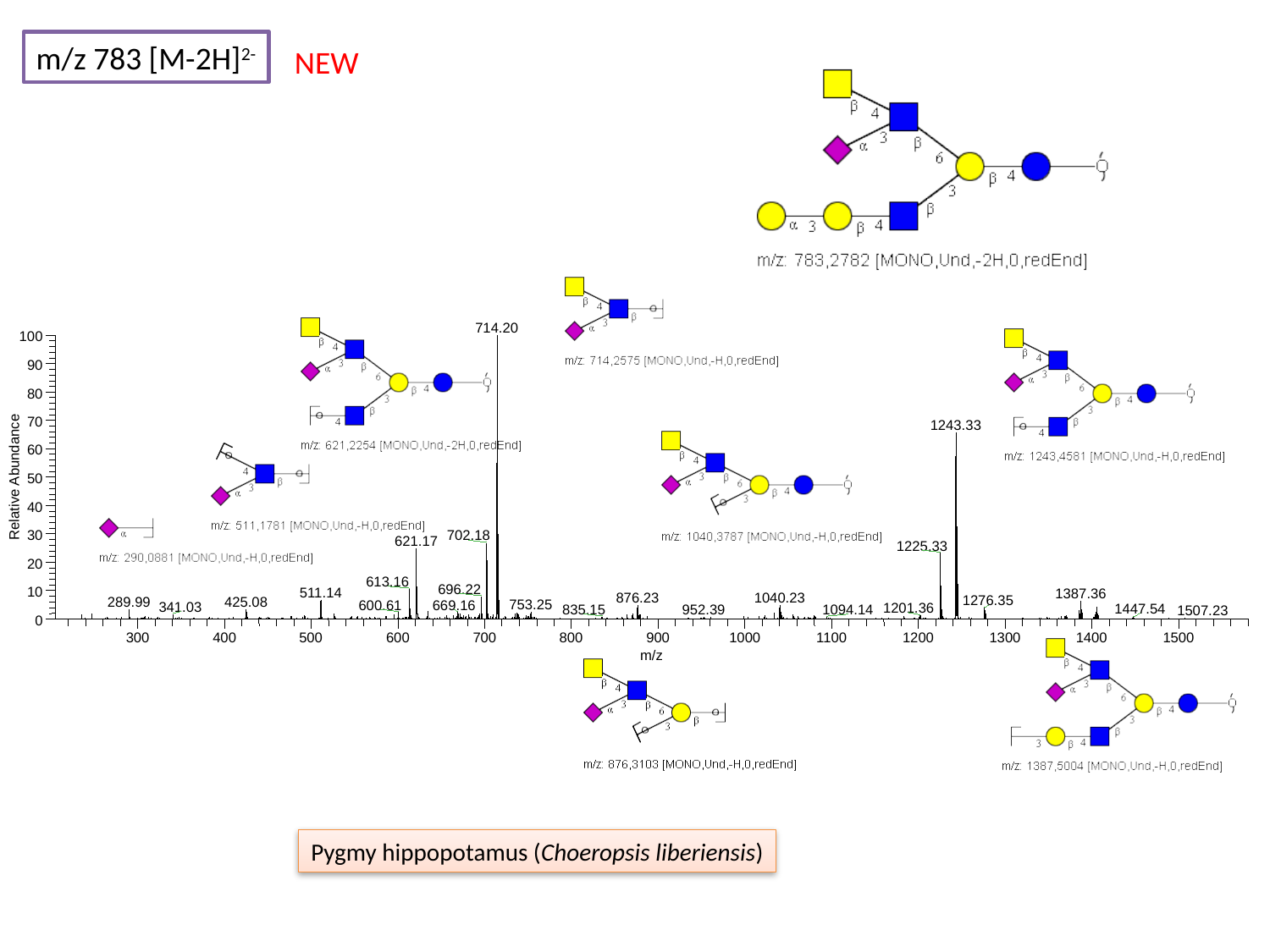

m/z 783 [M-2H]^2-^
NEW
714.20
100
90
80
70
1243.33
60
Relative Abundance
50
40
30
702.18
621.17
1225.33
20
613.16
696.22
10
511.14
1387.36
876.23
1040.23
1276.35
289.99
425.08
753.25
600.61
669.16
341.03
1201.36
1447.54
835.15
952.39
1094.14
0
300
400
500
600
700
800
900
1000
1100
1200
1300
1400
1500
m/z
1507.23
Pygmy hippopotamus (Choeropsis liberiensis)

### Slide 147
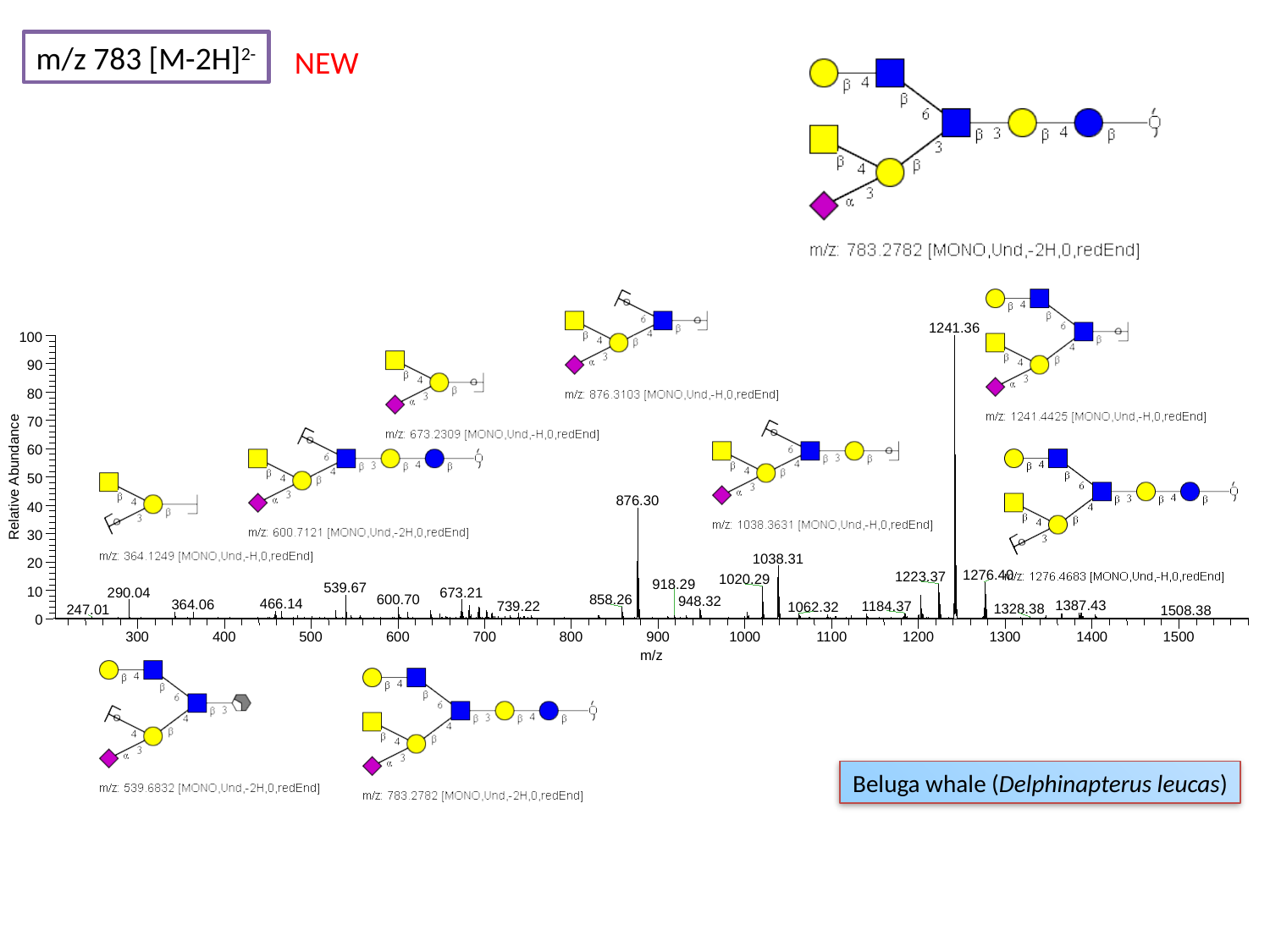

m/z 783 [M-2H]^2-^
NEW
1241.36
100
90
80
70
60
Relative Abundance
50
876.30
40
30
1038.31
20
1276.40
1223.37
1020.29
918.29
539.67
10
673.21
290.04
600.70
858.26
948.32
466.14
364.06
1387.43
739.22
1184.37
1062.32
1328.38
247.01
1508.38
0
300
400
500
600
700
800
900
1000
1100
1200
1300
1400
1500
m/z
Beluga whale (Delphinapterus leucas)

### Slide 148
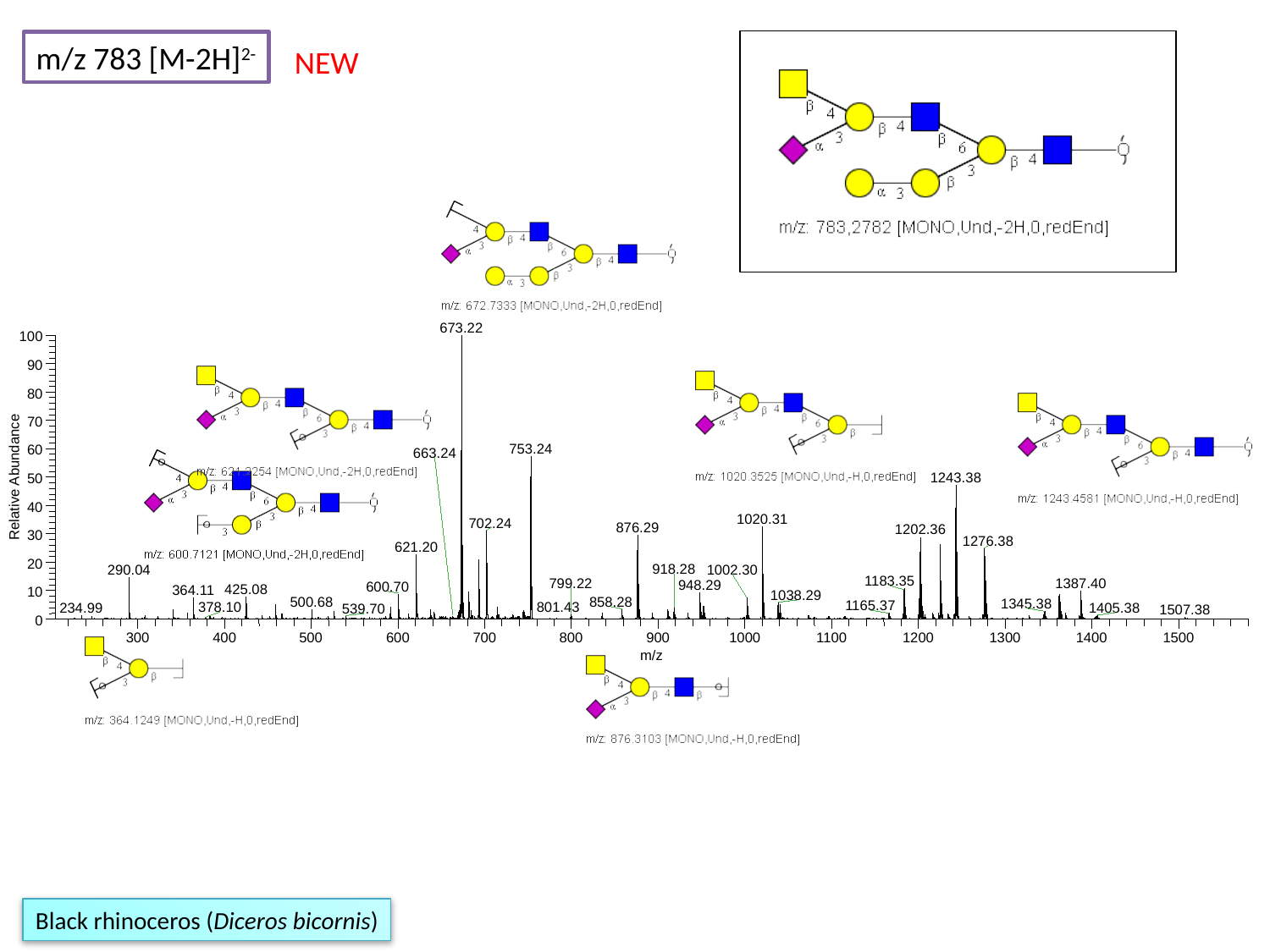

m/z 783 [M-2H]^2-^
NEW
673.22
100
90
80
70
753.24
60
Relative Abundance
1243.38
50
40
1020.31
702.24
876.29
1202.36
30
1276.38
621.20
20
918.28
290.04
1002.30
1183.35
799.22
1387.40
948.29
600.70
425.08
364.11
10
1038.29
500.68
858.28
1345.38
1165.37
0
300
400
500
600
700
800
900
1000
1100
1200
1300
1400
1500
m/z
663.24
801.43
378.10
234.99
1405.38
539.70
1507.38
Black rhinoceros (Diceros bicornis)

### Slide 149
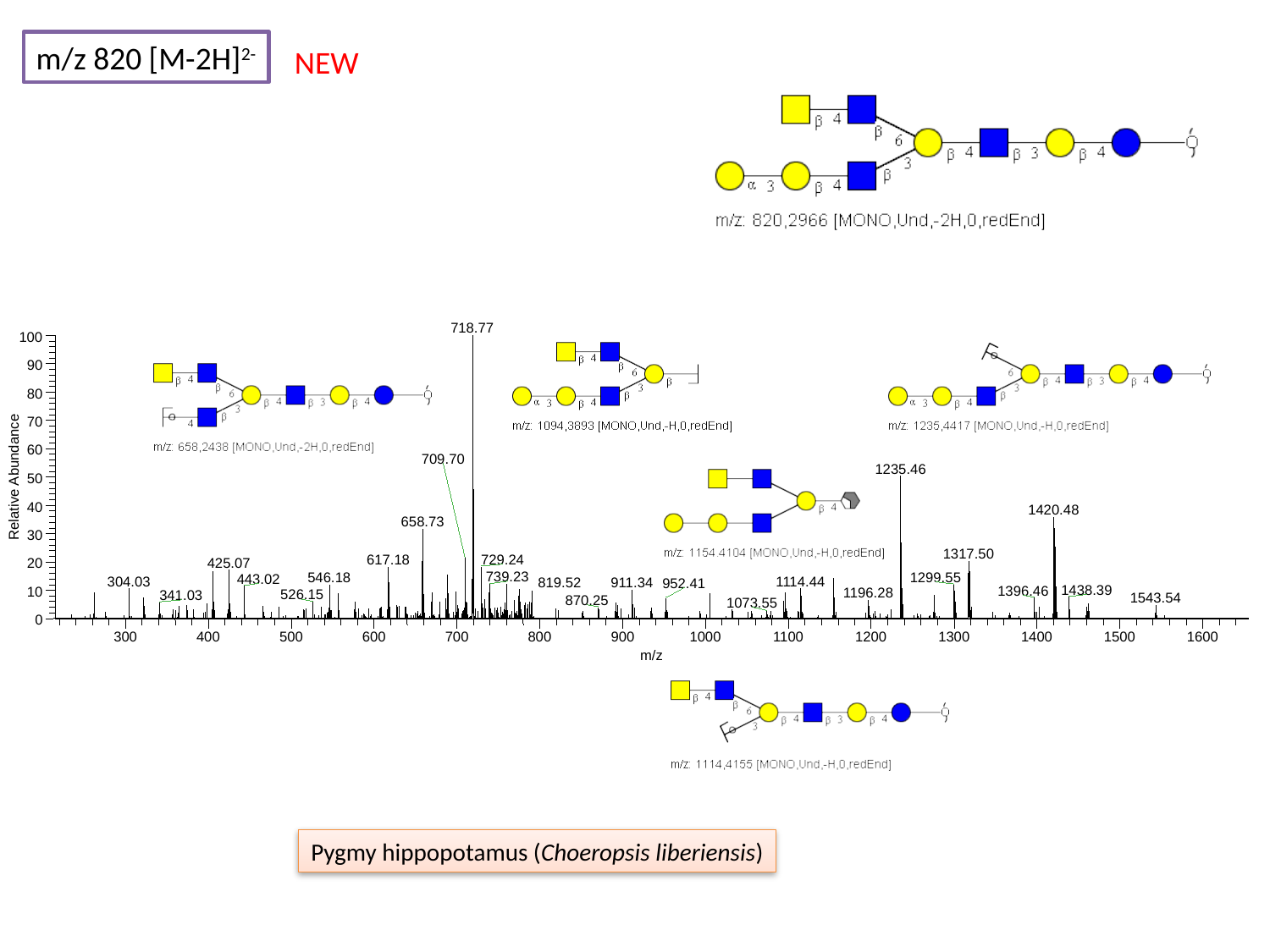

m/z 820 [M-2H]^2-^
NEW
718.77
100
90
80
70
60
709.70
1235.46
Relative Abundance
50
40
1420.48
658.73
30
1317.50
617.18
729.24
425.07
20
739.23
546.18
1299.55
443.02
304.03
1114.44
911.34
952.41
1438.39
1396.46
10
526.15
341.03
0
300
400
500
600
700
800
900
1000
1100
1200
1300
1400
1500
1600
m/z
819.52
1196.28
1543.54
870.25
1073.55
Pygmy hippopotamus (Choeropsis liberiensis)

### Slide 150
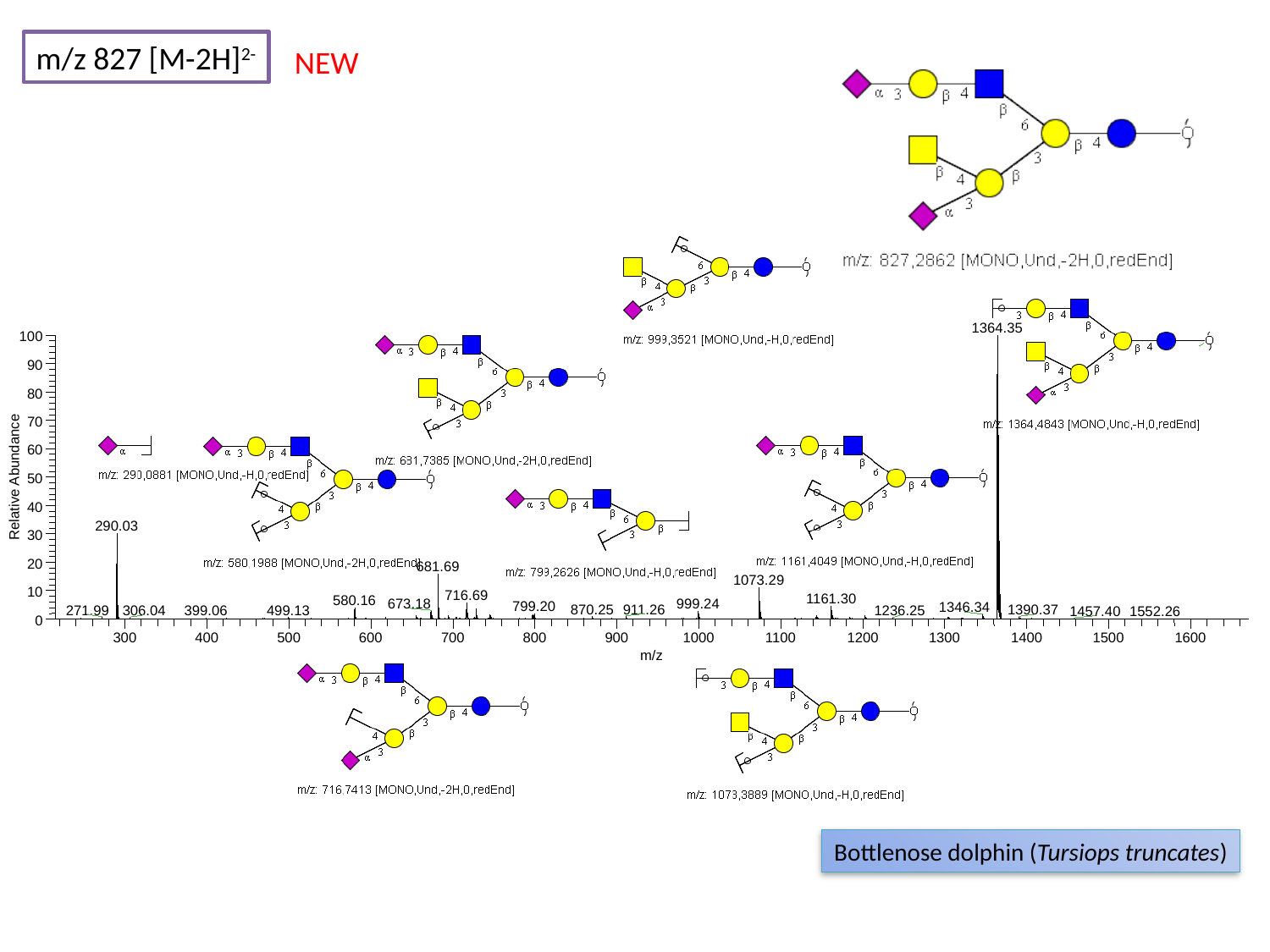

m/z 827 [M-2H]^2-^
NEW
1364.35
100
90
80
70
60
Relative Abundance
50
40
290.03
30
20
681.69
1073.29
10
716.69
1161.30
580.16
673.18
999.24
799.20
1346.34
870.25
911.26
1390.37
271.99
499.13
306.04
399.06
1236.25
1457.40
1552.26
0
300
400
500
600
700
800
900
1000
1100
1200
1300
1400
1500
1600
m/z
Bottlenose dolphin (Tursiops truncates)

### Slide 151
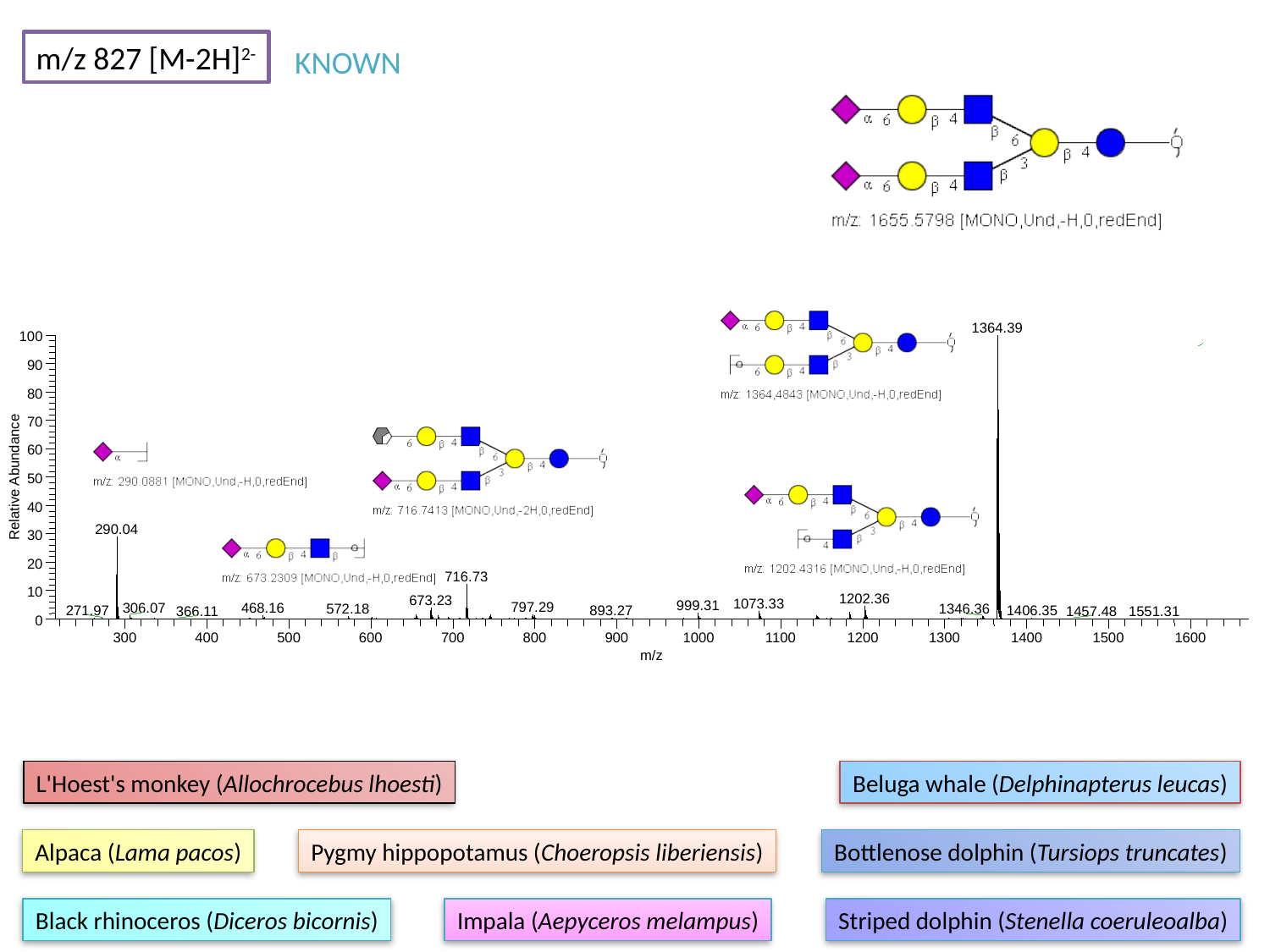

m/z 827 [M-2H]^2-^
KNOWN
1364.39
100
90
80
70
60
Relative Abundance
50
40
290.04
30
20
716.73
10
1202.36
673.23
1073.33
999.31
797.29
306.07
468.16
572.18
1346.36
271.97
893.27
1406.35
366.11
1457.48
1551.31
0
300
400
500
600
700
800
900
1000
1100
1200
1300
1400
1500
1600
m/z
L'Hoest's monkey (Allochrocebus lhoesti)
Beluga whale (Delphinapterus leucas)
Alpaca (Lama pacos)
Pygmy hippopotamus (Choeropsis liberiensis)
Bottlenose dolphin (Tursiops truncates)
Black rhinoceros (Diceros bicornis)
Impala (Aepyceros melampus)
Striped dolphin (Stenella coeruleoalba)

### Slide 152
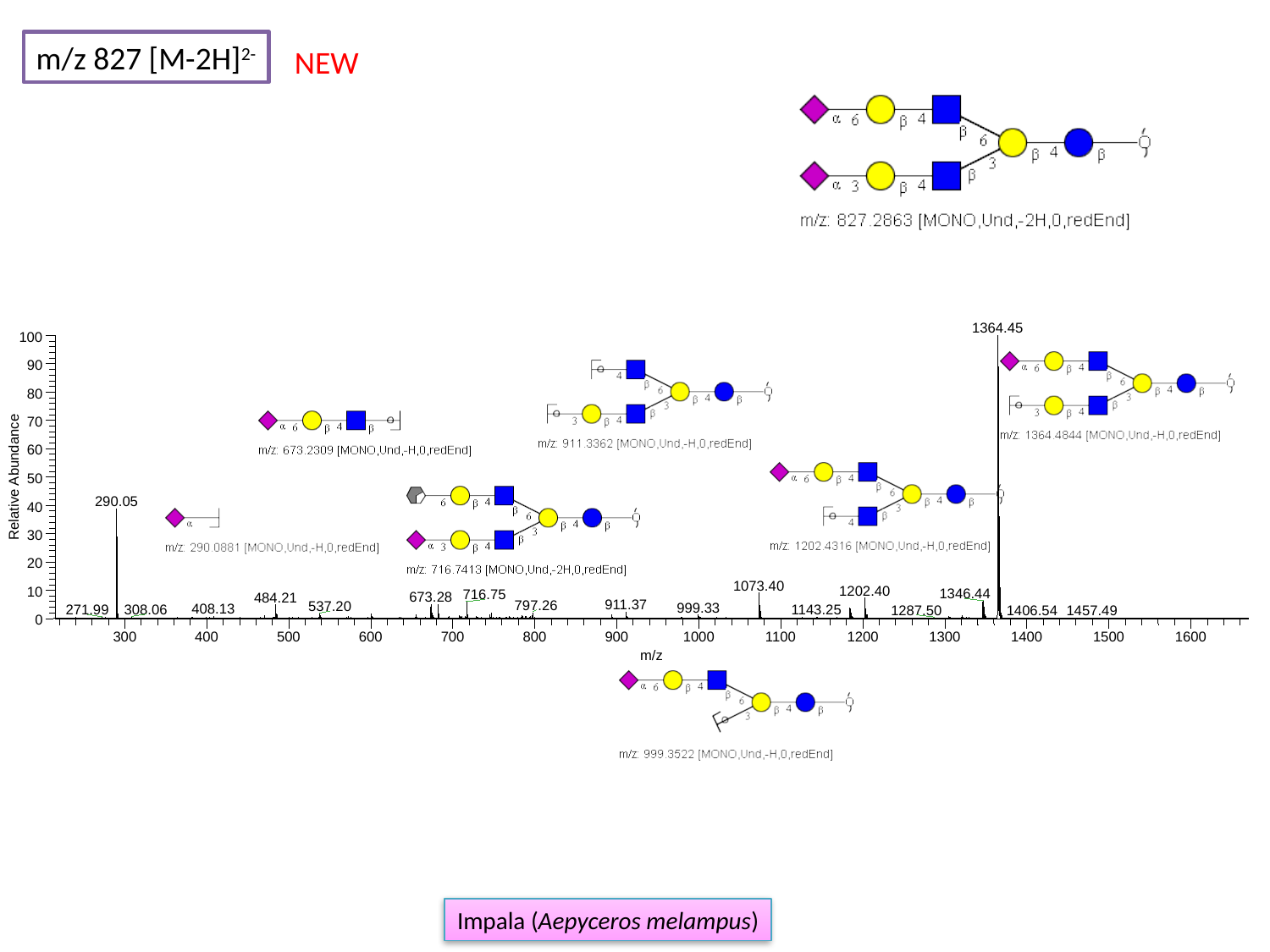

m/z 827 [M-2H]^2-^
NEW
1364.45
100
90
80
70
60
Relative Abundance
50
290.05
40
30
20
1073.40
1202.40
10
1346.44
716.75
673.28
484.21
911.37
797.26
537.20
999.33
408.13
308.06
271.99
1143.25
1287.50
1457.49
1406.54
0
300
400
500
600
700
800
900
1000
1100
1200
1300
1400
1500
1600
m/z
Impala (Aepyceros melampus)

### Slide 153
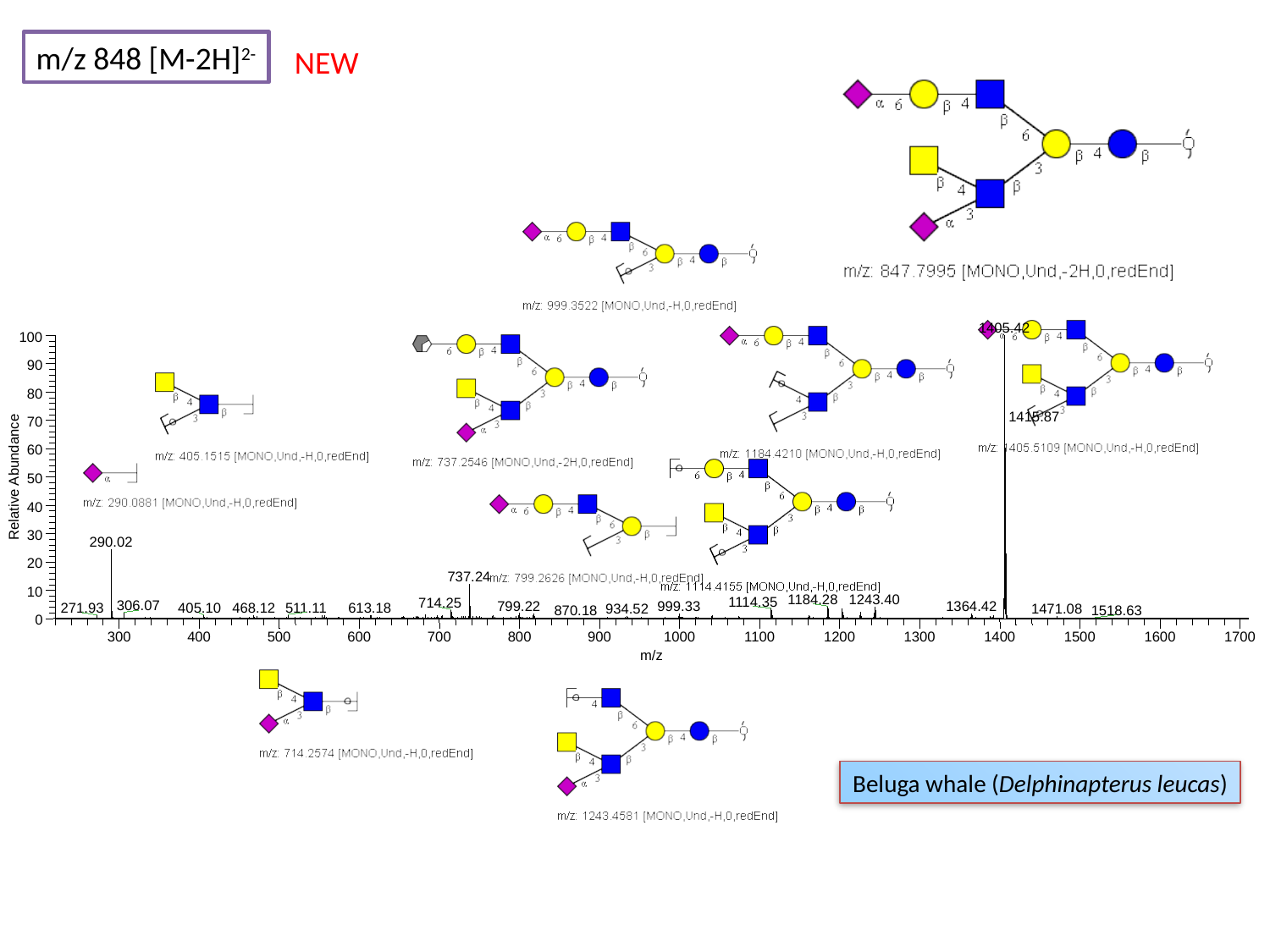

m/z 848 [M-2H]^2-^
NEW
1405.42
100
90
80
70
60
Relative Abundance
50
40
30
290.02
20
737.24
10
1184.28
1243.40
1114.35
714.25
306.07
799.22
999.33
1364.42
468.12
511.11
271.93
405.10
613.18
1471.08
934.52
1518.63
870.18
0
300
400
500
600
700
800
900
1000
1100
1200
1300
1400
1500
1600
1700
m/z
1415.87
Beluga whale (Delphinapterus leucas)

### Slide 154
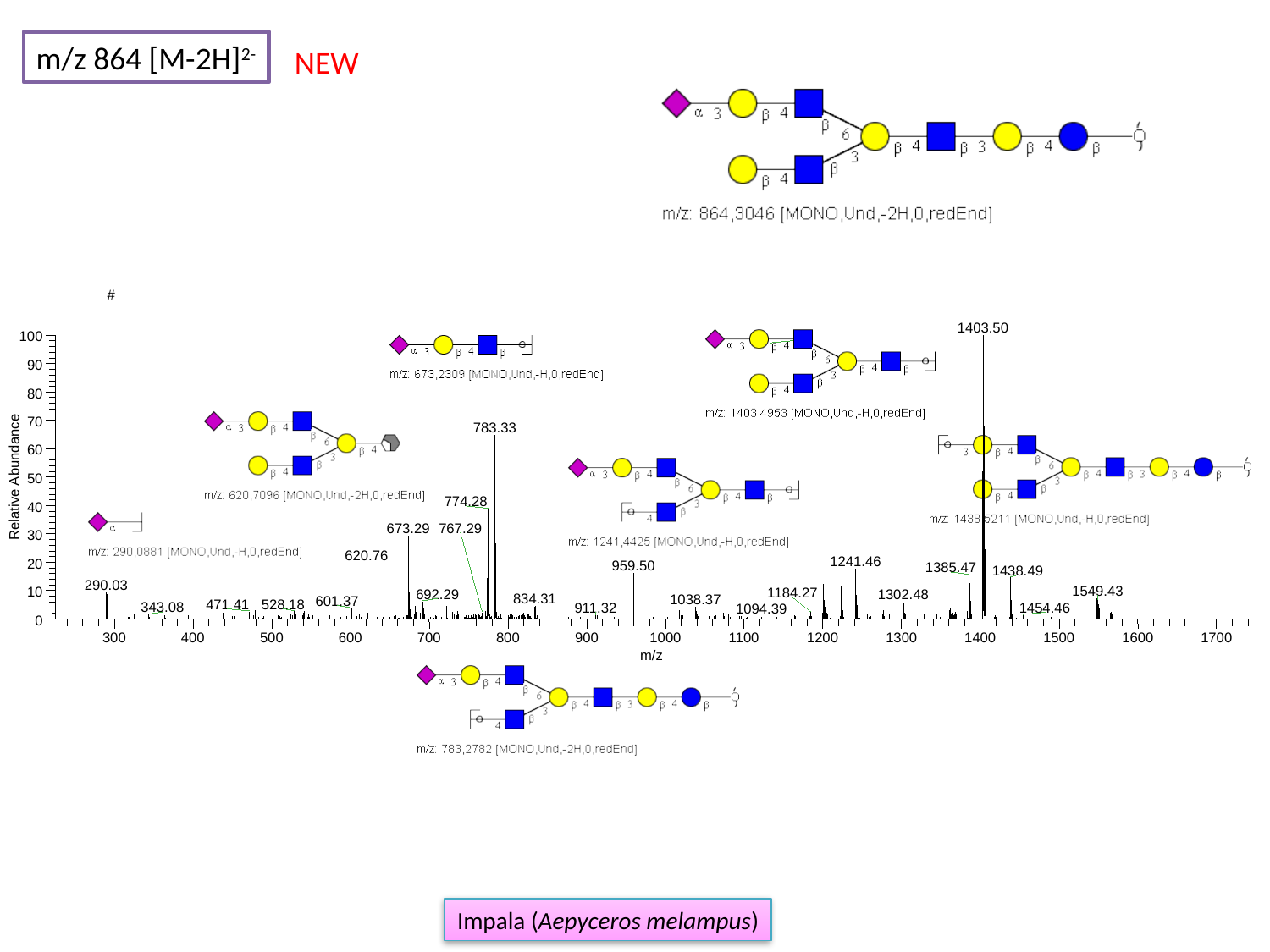

m/z 864 [M-2H]^2-^
NEW
#
1403.50
100
90
80
70
783.33
60
Relative Abundance
50
774.28
40
673.29
30
620.76
1241.46
20
959.50
1385.47
1438.49
290.03
1549.43
10
1184.27
692.29
1302.48
834.31
1038.37
601.37
528.18
0
300
400
500
600
700
800
900
1000
1100
1200
1300
1400
1500
1600
1700
m/z
767.29
471.41
343.08
911.32
1454.46
1094.39
Impala (Aepyceros melampus)

### Slide 155
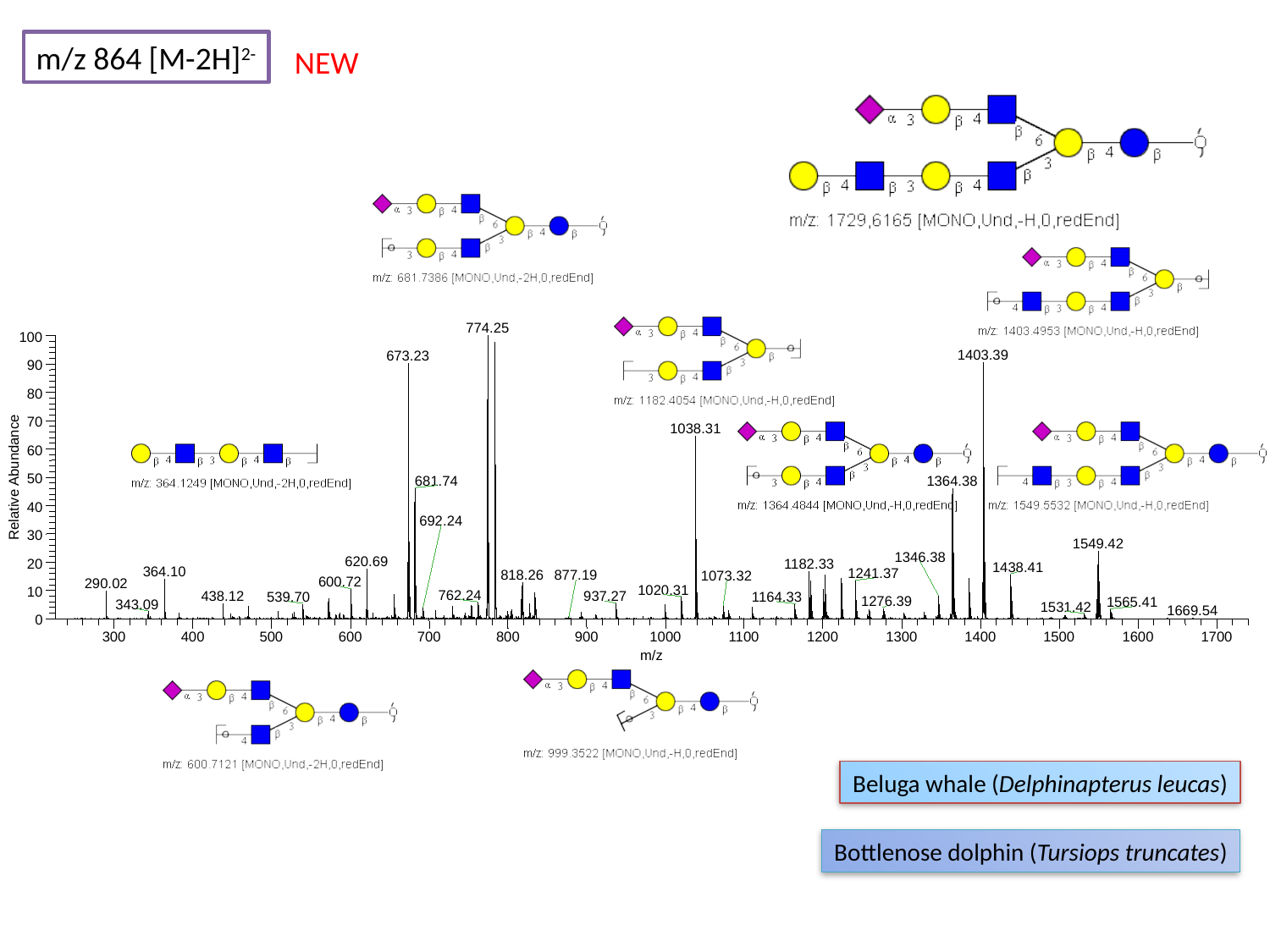

m/z 864 [M-2H]^2-^
NEW
774.25
100
1403.39
673.23
90
80
70
1038.31
60
Relative Abundance
50
681.74
1364.38
40
30
1549.42
1346.38
620.69
20
1182.33
1438.41
364.10
1241.37
818.26
600.72
290.02
1020.31
10
937.27
1164.33
0
300
400
500
600
700
800
900
1000
1100
1200
1300
1400
1500
1600
1700
m/z
692.24
877.19
1073.32
762.24
438.12
539.70
1276.39
1565.41
343.09
1531.42
1669.54
Beluga whale (Delphinapterus leucas)
Bottlenose dolphin (Tursiops truncates)

### Slide 156
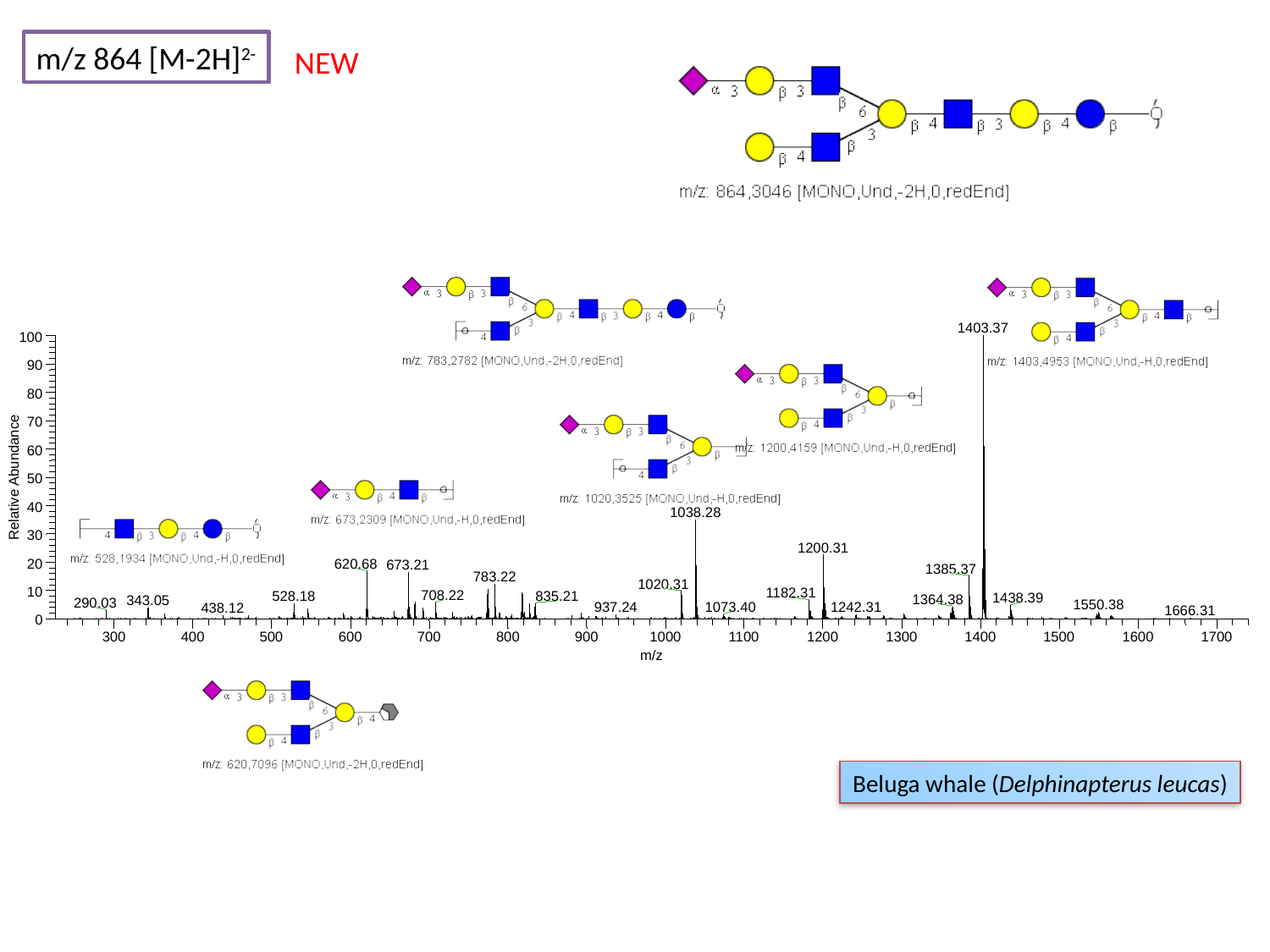

m/z 864 [M-2H]^2-^
NEW
1403.37
100
90
80
70
60
Relative Abundance
50
40
1038.28
30
1200.31
20
620.68
673.21
1385.37
783.22
1020.31
10
1182.31
708.22
528.18
835.21
1438.39
1364.38
343.05
290.03
1550.38
0
300
400
500
600
700
800
900
1000
1100
1200
1300
1400
1500
1600
1700
m/z
1073.40
1242.31
937.24
438.12
1666.31
Beluga whale (Delphinapterus leucas)

### Slide 157
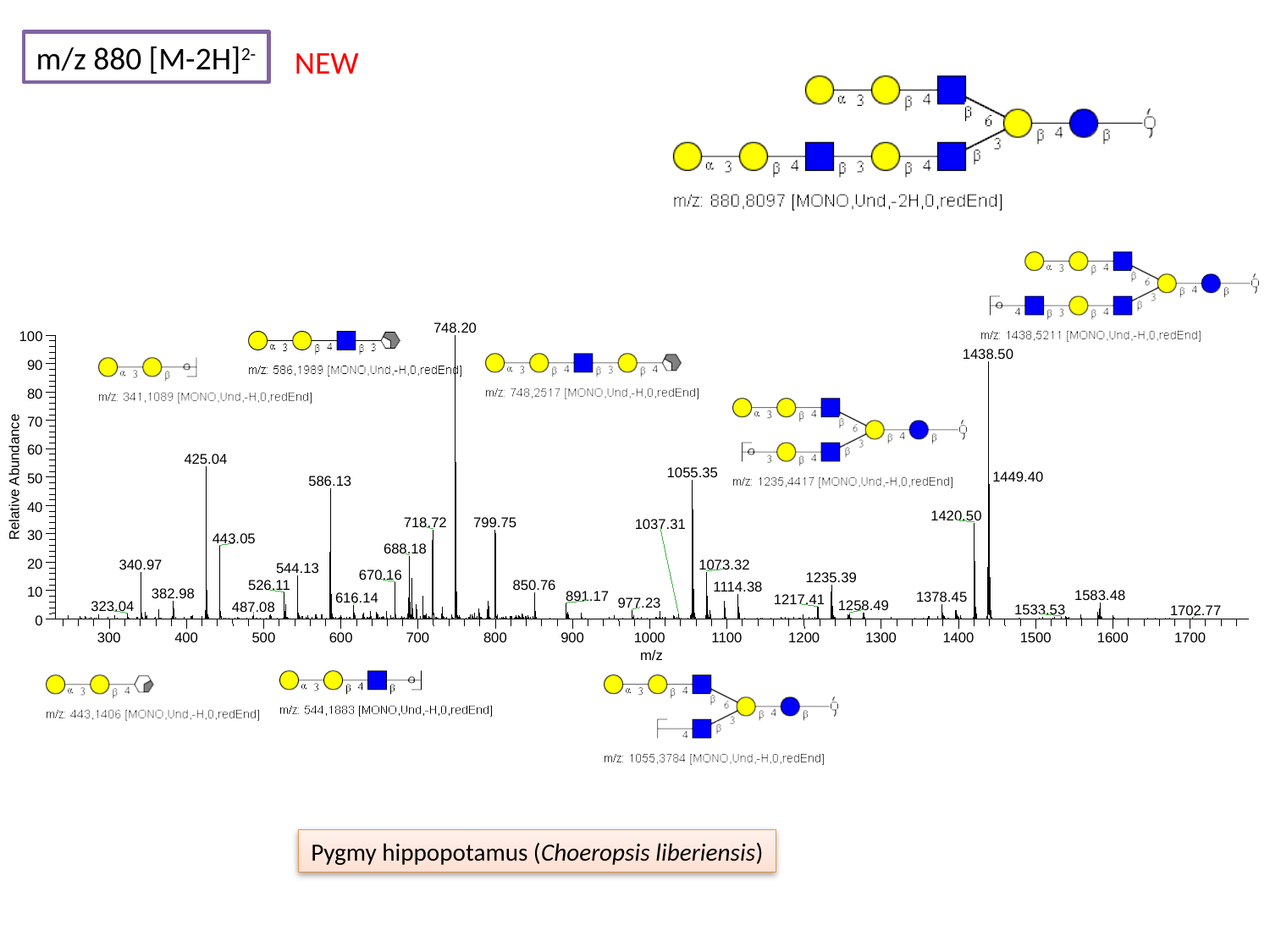

m/z 880 [M-2H]^2-^
NEW
748.20
100
1438.50
90
80
70
60
425.04
1055.35
Relative Abundance
50
586.13
40
1420.50
718.72
799.75
30
443.05
688.18
20
340.97
1073.32
544.13
670.16
1235.39
526.11
850.76
1114.38
10
0
300
400
500
600
700
800
900
1000
1100
1200
1300
1400
1500
1600
1700
m/z
1449.40
1037.31
382.98
1583.48
891.17
1378.45
616.14
1217.41
977.23
1258.49
323.04
487.08
1533.53
1702.77
Pygmy hippopotamus (Choeropsis liberiensis)

### Slide 158
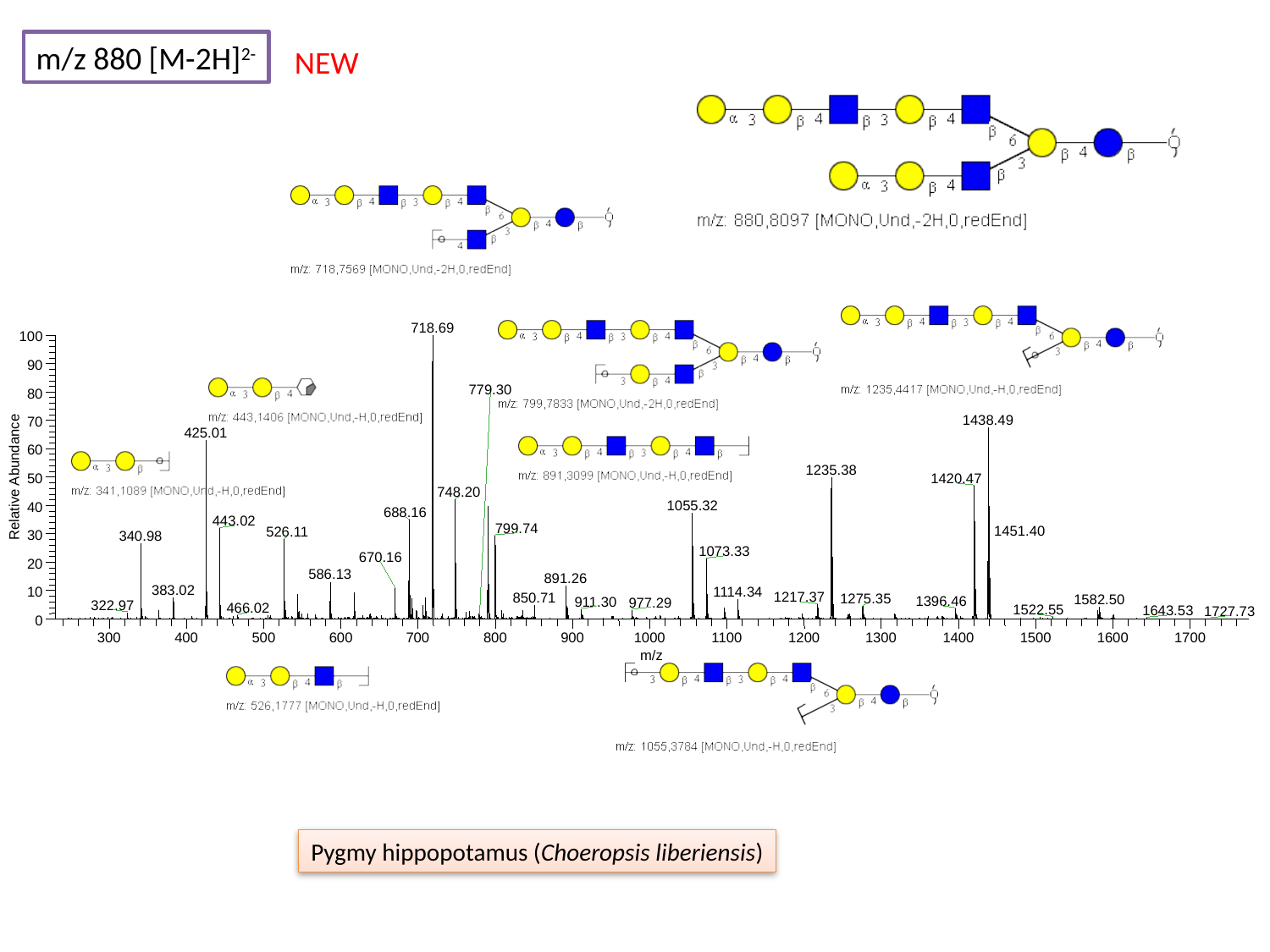

m/z 880 [M-2H]^2-^
NEW
718.69
100
90
80
1438.49
70
425.01
60
1235.38
Relative Abundance
50
1420.47
748.20
1055.32
40
688.16
443.02
799.74
526.11
30
340.98
1073.33
670.16
20
586.13
891.26
1114.34
10
0
300
400
500
600
700
800
900
1000
1100
1200
1300
1400
1500
1600
1700
m/z
779.30
1451.40
383.02
1217.37
850.71
1275.35
1582.50
1396.46
911.30
977.29
322.97
466.02
1522.55
1643.53
1727.73
Pygmy hippopotamus (Choeropsis liberiensis)

### Slide 159
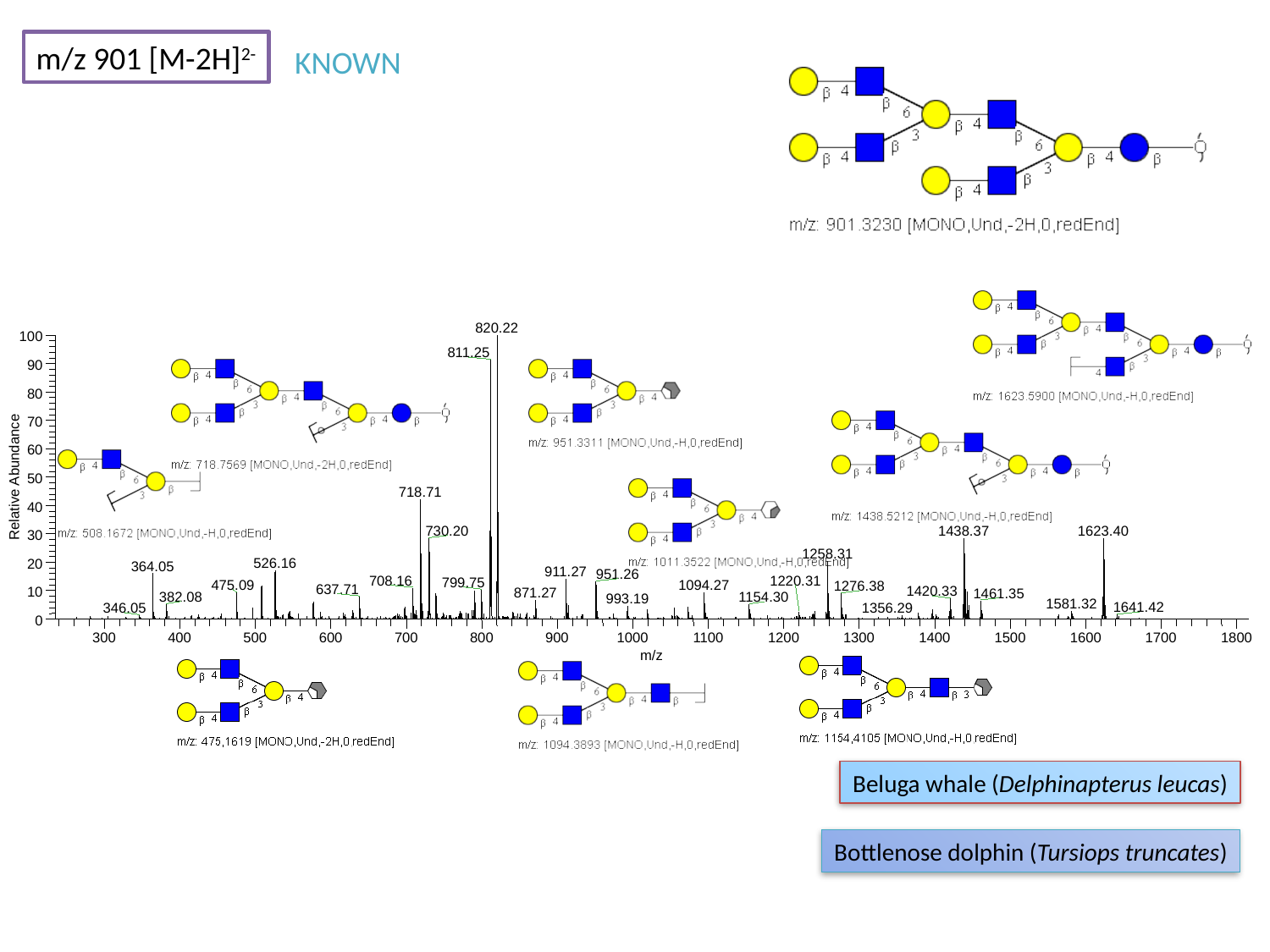

m/z 901 [M-2H]^2-^
KNOWN
820.22
100
811.25
90
80
70
60
Relative Abundance
50
718.71
40
730.20
1623.40
1438.37
30
1258.31
526.16
20
364.05
911.27
951.26
708.16
799.75
1094.27
1276.38
1420.33
10
0
300
400
500
600
700
800
900
1000
1100
1200
1300
1400
1500
1600
1700
1800
m/z
1220.31
475.09
637.71
871.27
1461.35
382.08
1154.30
993.19
1581.32
1641.42
1356.29
346.05
Beluga whale (Delphinapterus leucas)
Bottlenose dolphin (Tursiops truncates)

### Slide 160
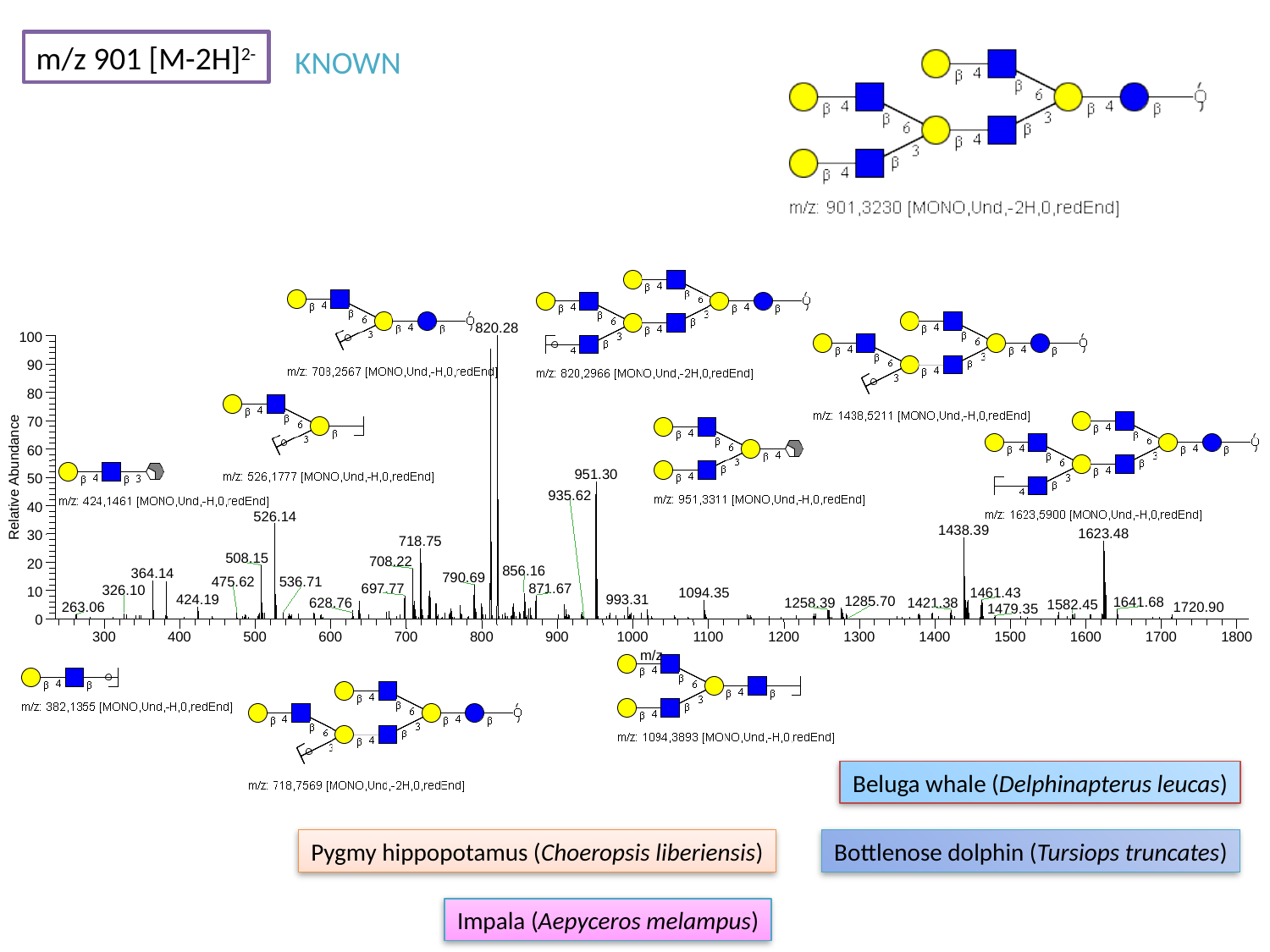

m/z 901 [M-2H]^2-^
KNOWN
820.28
100
90
80
70
60
951.30
Relative Abundance
50
40
526.14
1438.39
1623.48
30
718.75
508.15
708.22
20
856.16
364.14
790.69
697.77
871.67
10
1094.35
1461.43
0
300
400
500
600
700
800
900
1000
1100
1200
1300
1400
1500
1600
1700
1800
m/z
935.62
536.71
475.62
326.10
424.19
993.31
1285.70
1641.68
1258.39
1421.38
628.76
1582.45
263.06
1720.90
1479.35
Beluga whale (Delphinapterus leucas)
Pygmy hippopotamus (Choeropsis liberiensis)
Bottlenose dolphin (Tursiops truncates)
Impala (Aepyceros melampus)

### Slide 161
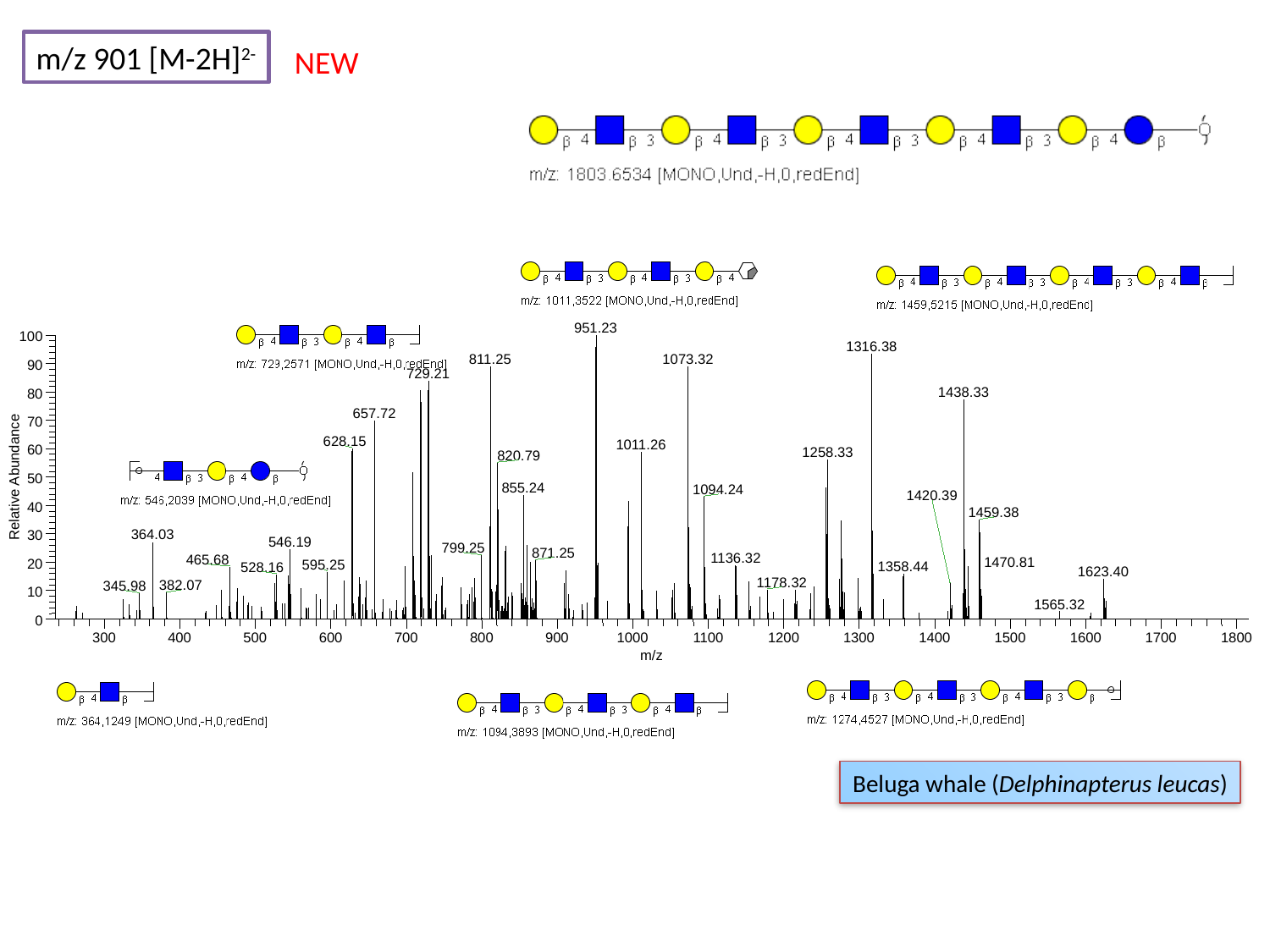

m/z 901 [M-2H]^2-^
NEW
951.23
100
1316.38
811.25
1073.32
90
729.21
1438.33
80
657.72
70
628.15
1011.26
60
1258.33
820.79
Relative Abundance
50
855.24
1094.24
40
1459.38
364.03
30
546.19
799.25
20
10
0
300
400
500
600
700
800
900
1000
1100
1200
1300
1400
1500
1600
1700
1800
m/z
1420.39
871.25
1136.32
465.68
1470.81
595.25
1358.44
528.16
1623.40
1178.32
382.07
345.98
1565.32
Beluga whale (Delphinapterus leucas)

### Slide 162
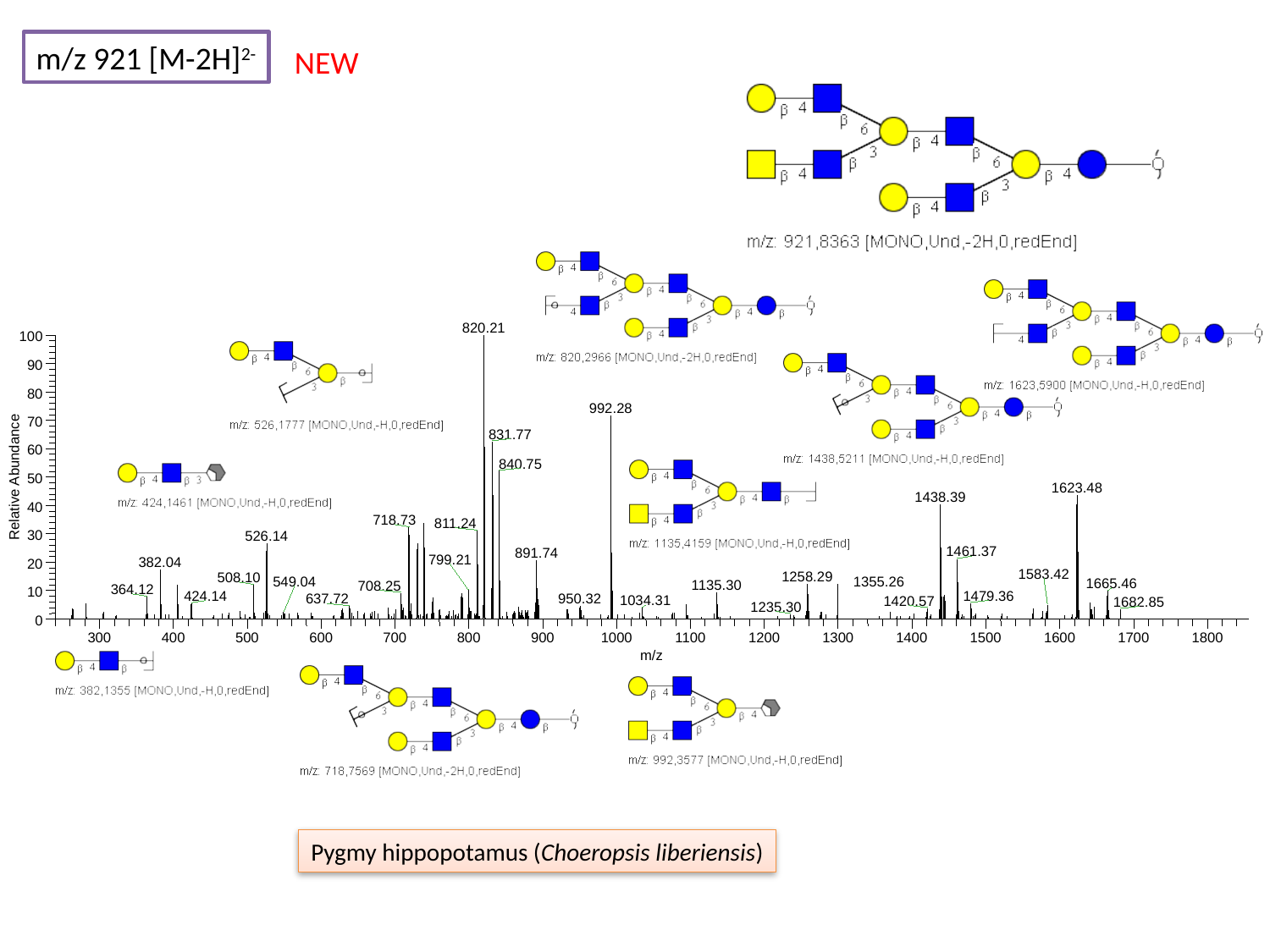

m/z 921 [M-2H]^2-^
NEW
820.21
100
90
80
992.28
70
831.77
60
840.75
Relative Abundance
50
1623.48
1438.39
40
718.73
811.24
30
526.14
1461.37
891.74
382.04
20
508.10
1665.46
10
0
300
400
500
600
700
800
900
1000
1100
1200
1300
1400
1500
1600
1700
1800
m/z
799.21
1583.42
1258.29
549.04
1355.26
1135.30
708.25
364.12
1479.36
424.14
637.72
950.32
1034.31
1420.57
1682.85
1235.30
Pygmy hippopotamus (Choeropsis liberiensis)

### Slide 163
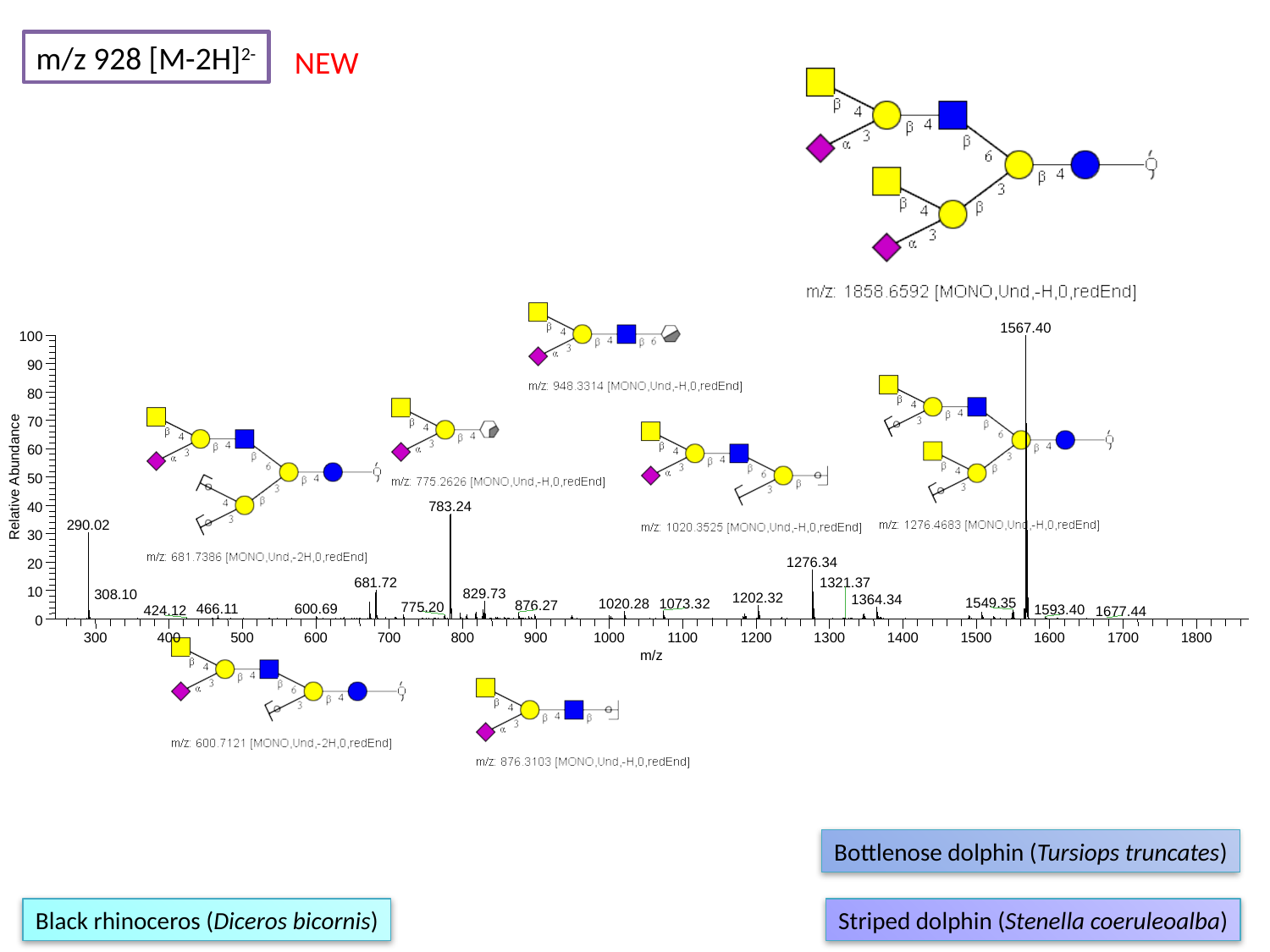

m/z 928 [M-2H]^2-^
NEW
1567.40
100
90
80
70
60
Relative Abundance
50
783.24
40
290.02
30
1276.34
20
681.72
10
829.73
1202.32
1364.34
1549.35
1020.28
1073.32
876.27
775.20
466.11
600.69
1593.40
0
300
400
500
600
700
800
900
1000
1100
1200
1300
1400
1500
1600
1700
1800
m/z
1321.37
308.10
424.12
1677.44
Bottlenose dolphin (Tursiops truncates)
Black rhinoceros (Diceros bicornis)
Striped dolphin (Stenella coeruleoalba)

### Slide 164
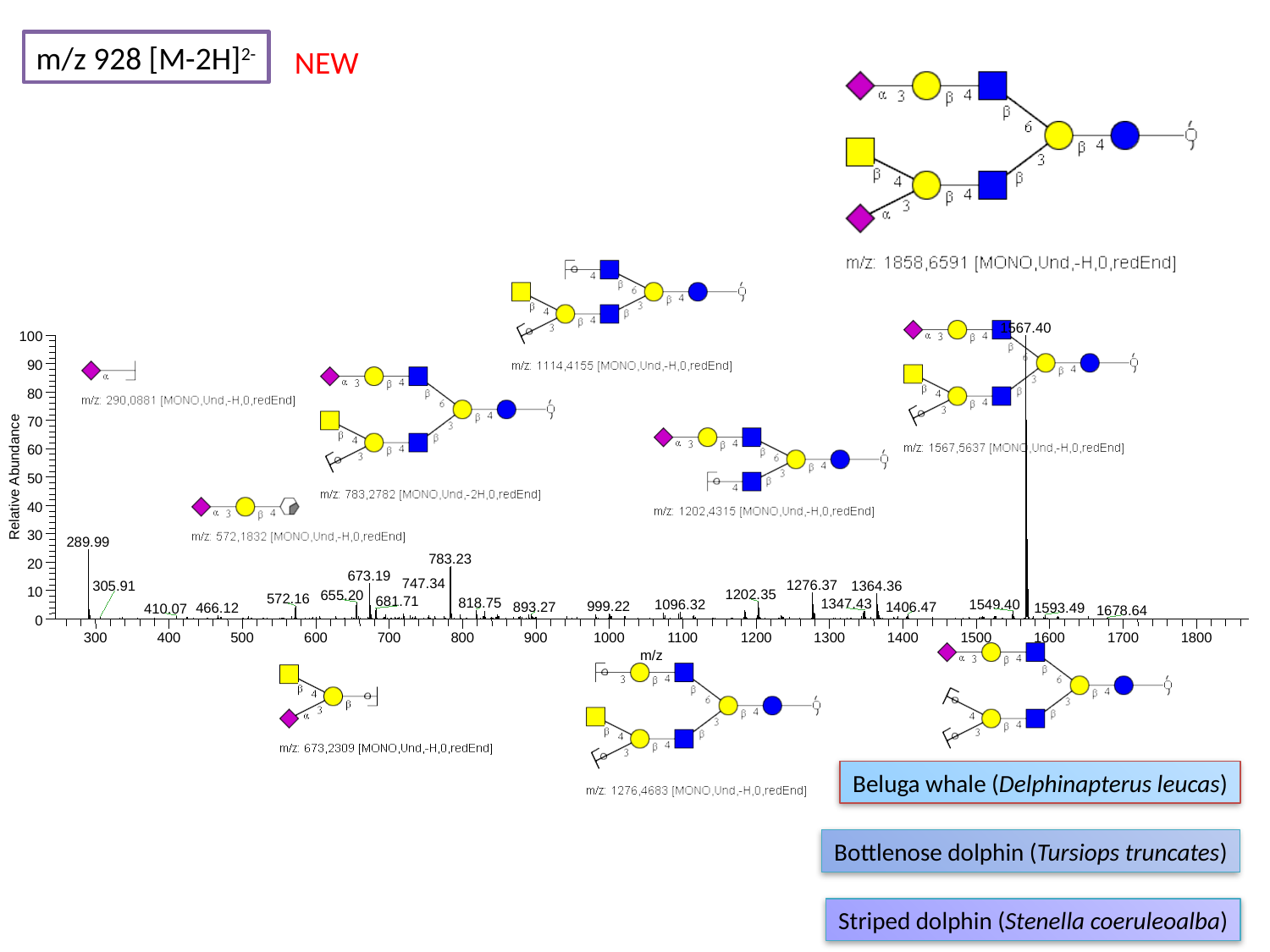

m/z 928 [M-2H]^2-^
NEW
1567.40
100
90
80
70
60
Relative Abundance
50
40
30
289.99
783.23
20
673.19
1276.37
1364.36
10
1202.35
655.20
572.16
681.71
818.75
1347.43
1549.40
0
300
400
500
600
700
800
900
1000
1100
1200
1300
1400
1500
1600
1700
1800
m/z
747.34
305.91
1096.32
999.22
893.27
1406.47
1593.49
466.12
410.07
1678.64
Beluga whale (Delphinapterus leucas)
Bottlenose dolphin (Tursiops truncates)
Striped dolphin (Stenella coeruleoalba)

### Slide 165
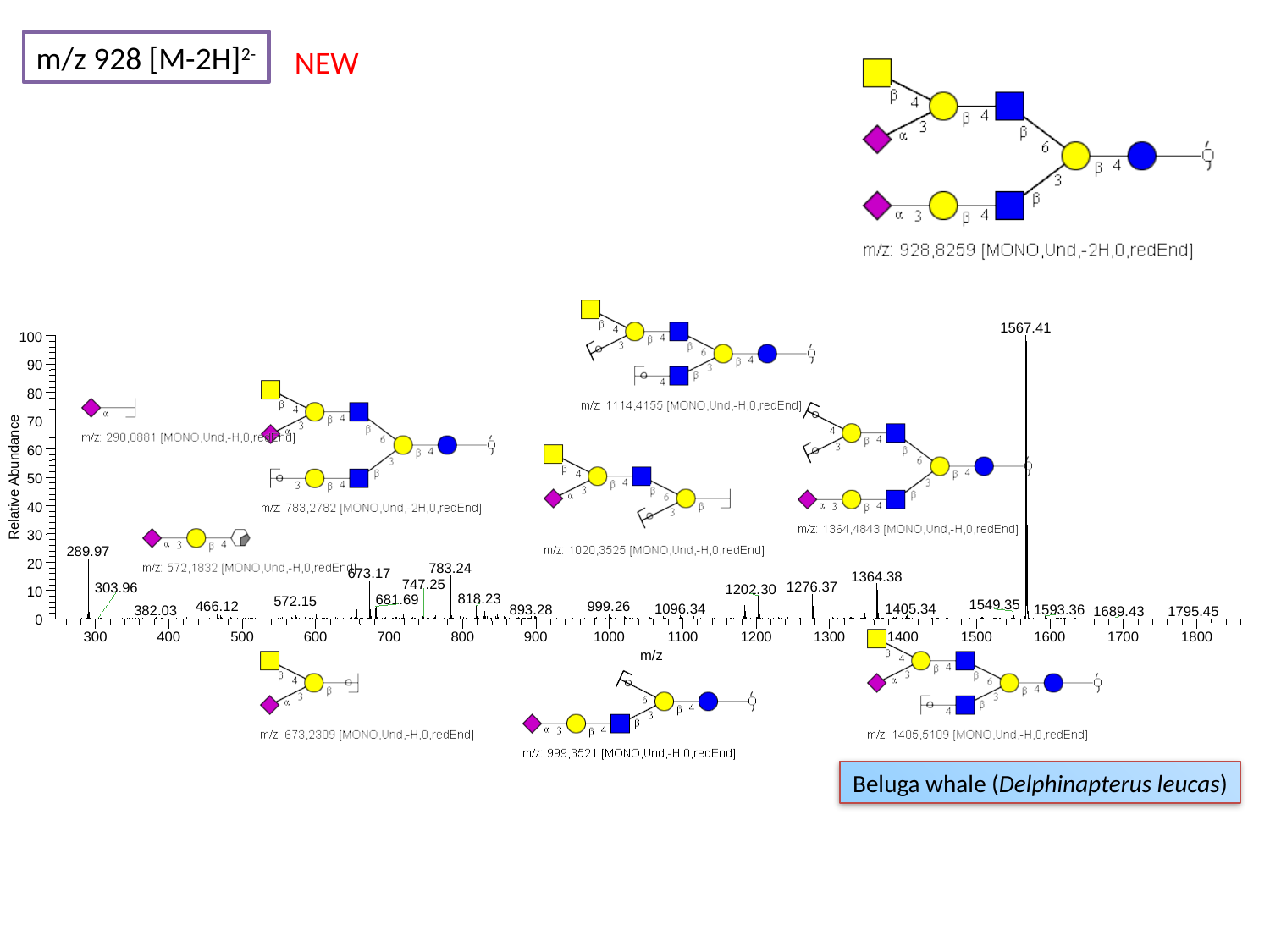

m/z 928 [M-2H]^2-^
NEW
1567.41
100
90
80
70
60
Relative Abundance
50
40
30
289.97
20
783.24
673.17
1364.38
1276.37
1202.30
10
818.23
681.69
572.15
1549.35
999.26
466.12
1096.34
1405.34
0
300
400
500
600
700
800
900
1000
1100
1200
1300
1400
1500
1600
1700
1800
m/z
747.25
303.96
893.28
1593.36
382.03
1689.43
1795.45
Beluga whale (Delphinapterus leucas)

### Slide 166
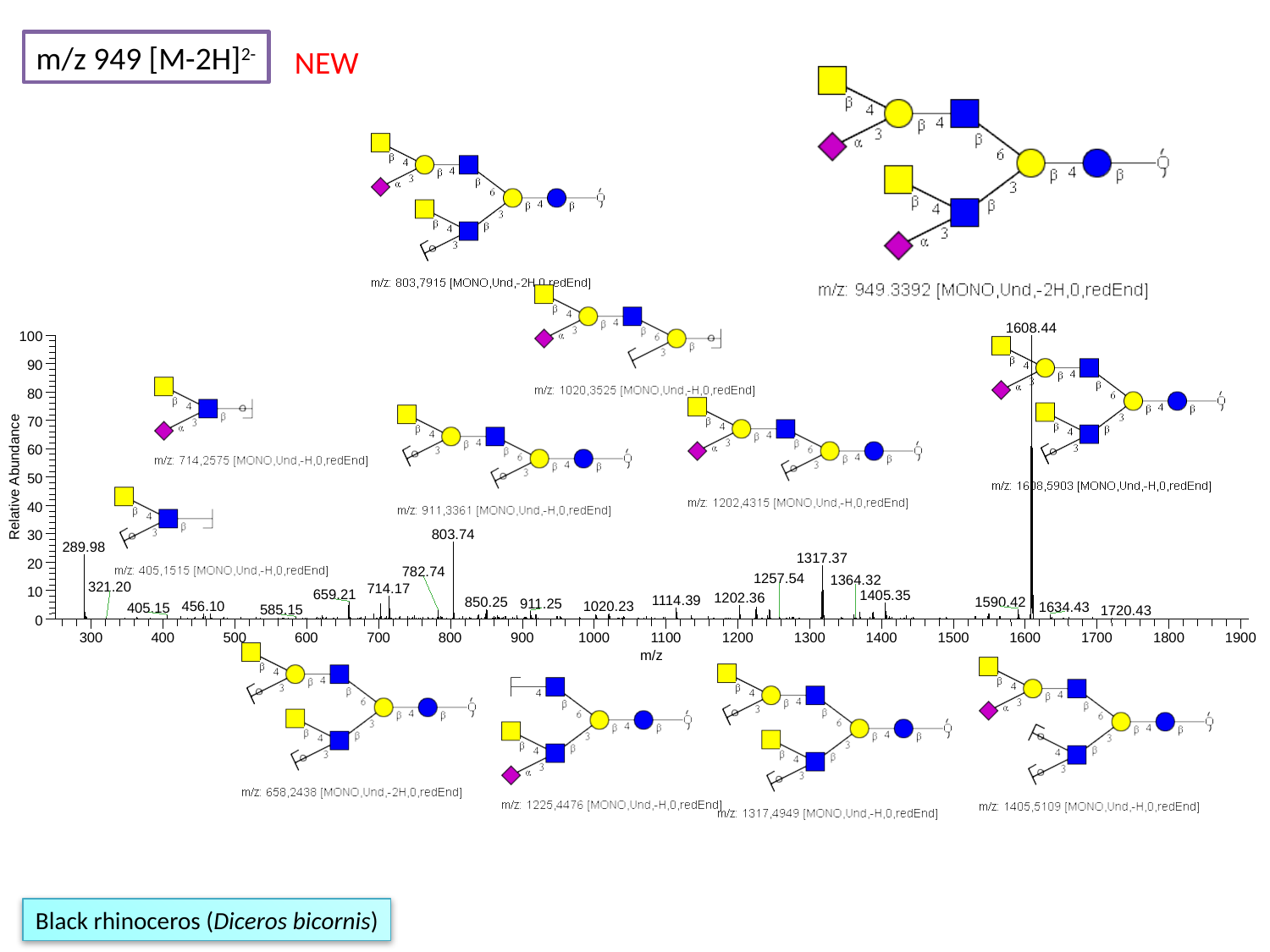

m/z 949 [M-2H]^2-^
NEW
1608.44
100
90
80
70
60
Relative Abundance
50
40
803.74
30
289.98
1317.37
20
782.74
714.17
10
659.21
1405.35
1202.36
1114.39
1590.42
850.25
911.25
0
300
400
500
600
700
800
900
1000
1100
1200
1300
1400
1500
1600
1700
1800
1900
m/z
1257.54
1364.32
321.20
1020.23
456.10
1634.43
405.15
585.15
1720.43
Black rhinoceros (Diceros bicornis)

### Slide 167
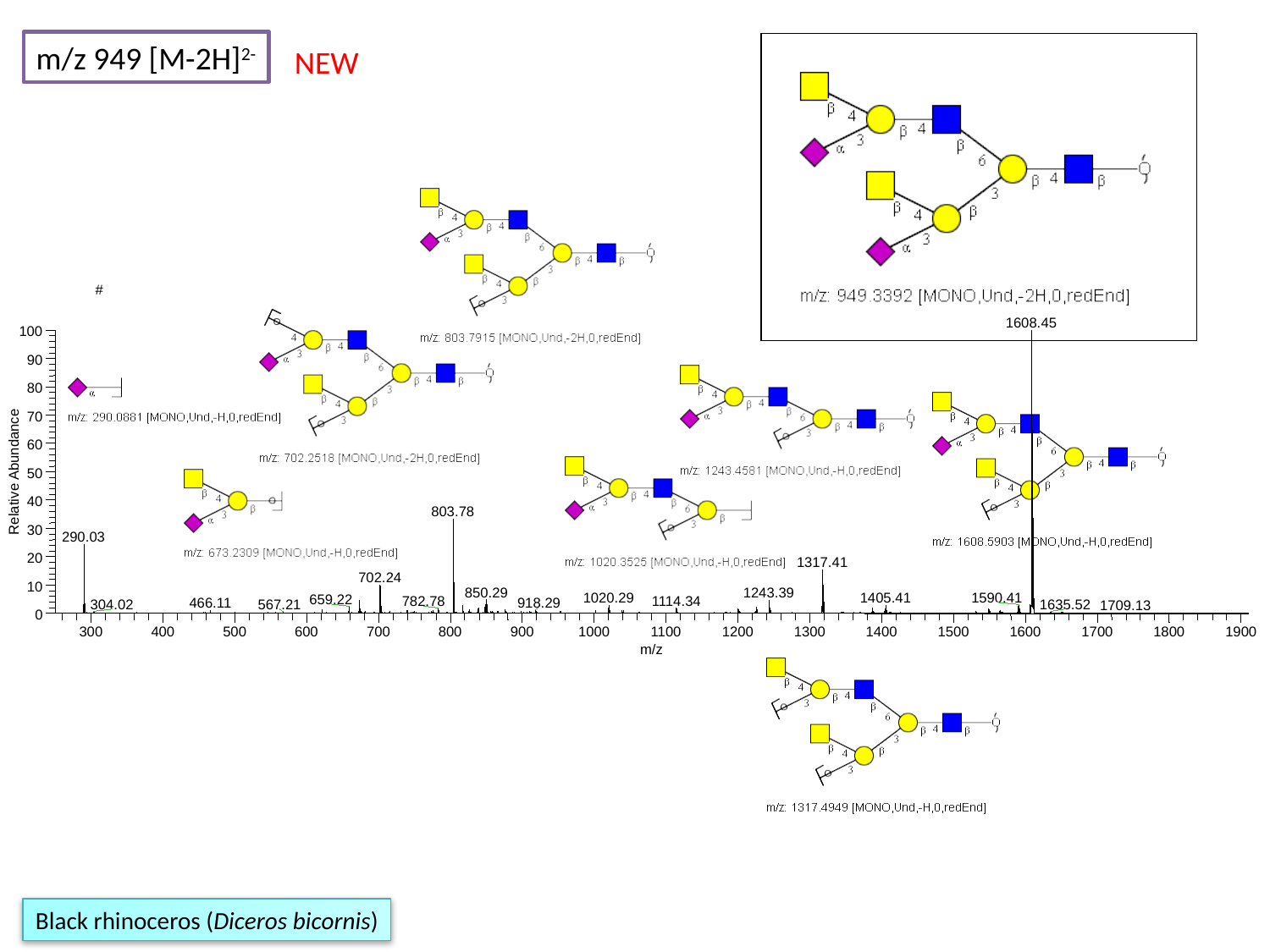

m/z 949 [M-2H]^2-^
NEW
#
1608.45
100
90
80
70
60
Relative Abundance
50
40
803.78
30
290.03
20
1317.41
702.24
10
850.29
1243.39
1590.41
1020.29
1405.41
659.22
1114.34
782.78
918.29
0
300
400
500
600
700
800
900
1000
1100
1200
1300
1400
1500
1600
1700
1800
1900
m/z
466.11
1635.52
304.02
567.21
1709.13
Black rhinoceros (Diceros bicornis)

### Slide 168
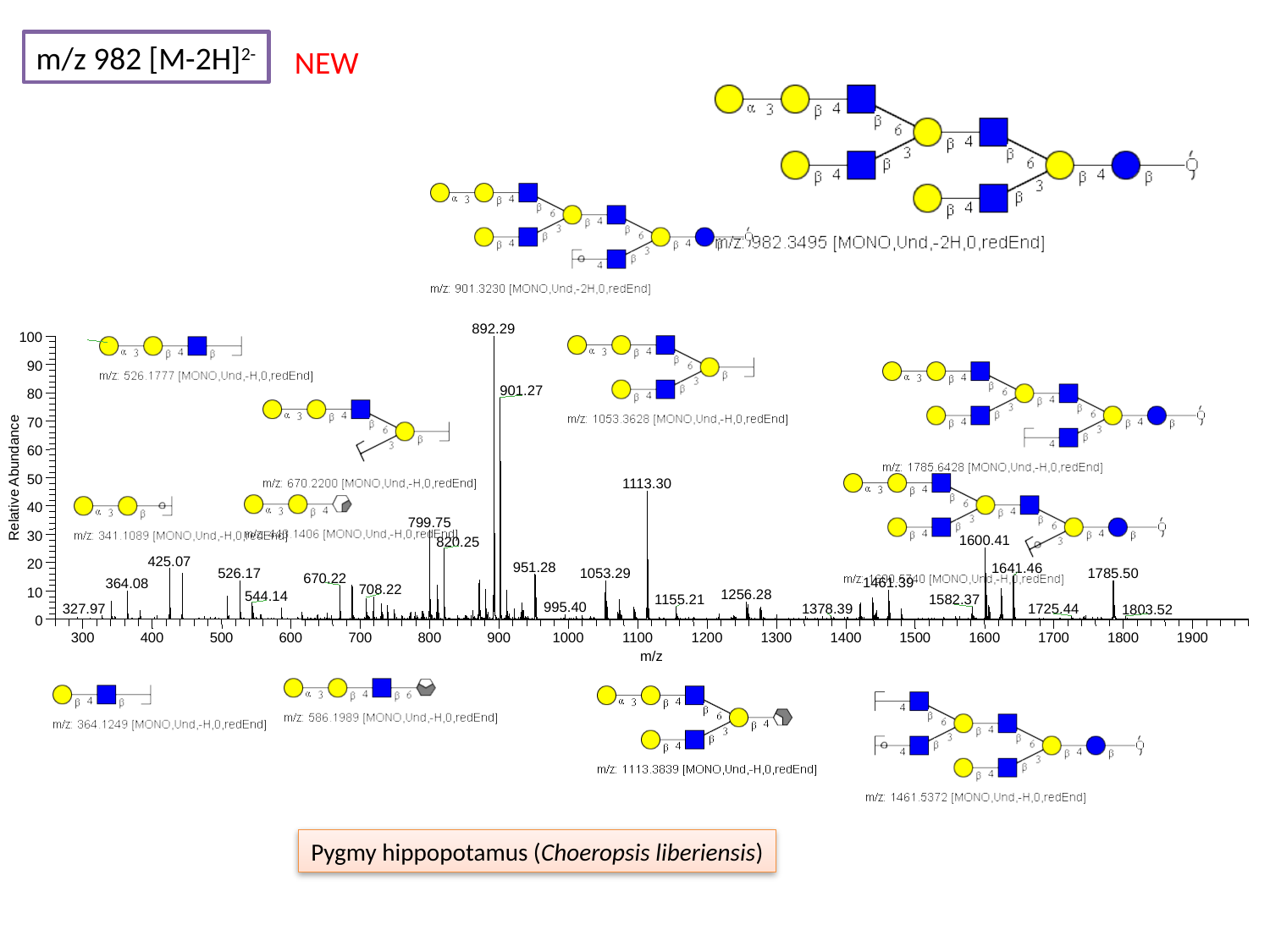

m/z 982 [M-2H]^2-^
NEW
892.29
100
90
901.27
80
70
60
Relative Abundance
50
1113.30
40
799.75
30
1600.41
820.25
425.07
20
951.28
1641.46
1785.50
10
0
300
400
500
600
700
800
900
1000
1100
1200
1300
1400
1500
1600
1700
1800
1900
m/z
1053.29
526.17
670.22
1461.39
364.08
708.22
1256.28
544.14
1155.21
1582.37
995.40
327.97
1378.39
1725.44
1803.52
Pygmy hippopotamus (Choeropsis liberiensis)

### Slide 169
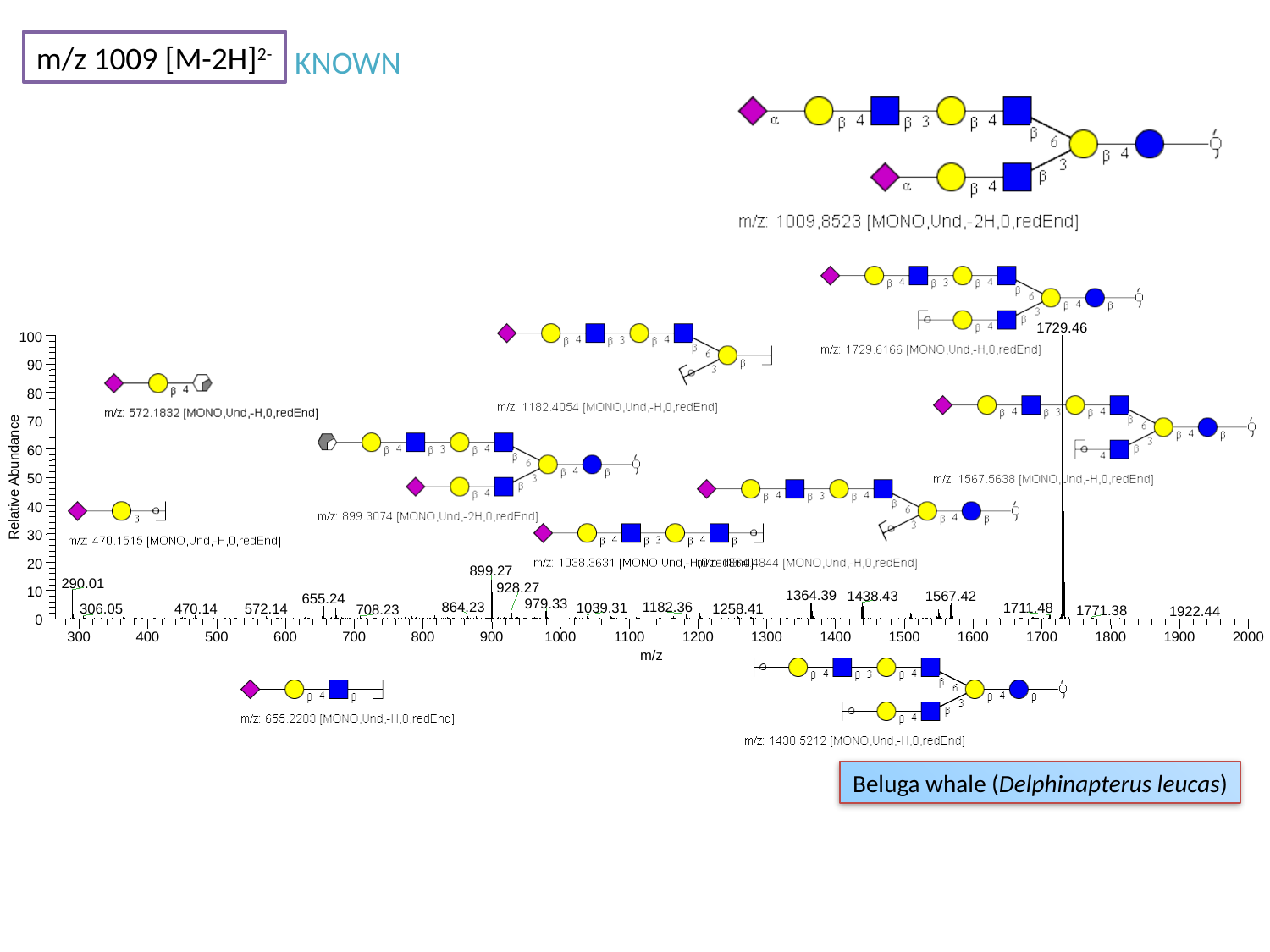

m/z 1009 [M-2H]^2-^
KNOWN
1729.46
100
90
80
70
60
Relative Abundance
50
40
30
20
899.27
290.01
10
1364.39
1438.43
1567.42
655.24
979.33
0
300
400
500
600
700
800
900
1000
1100
1200
1300
1400
1500
1600
1700
1800
1900
2000
m/z
928.27
864.23
1182.36
1039.31
1711.48
470.14
306.05
572.14
1258.41
708.23
1771.38
1922.44
Beluga whale (Delphinapterus leucas)

### Slide 170
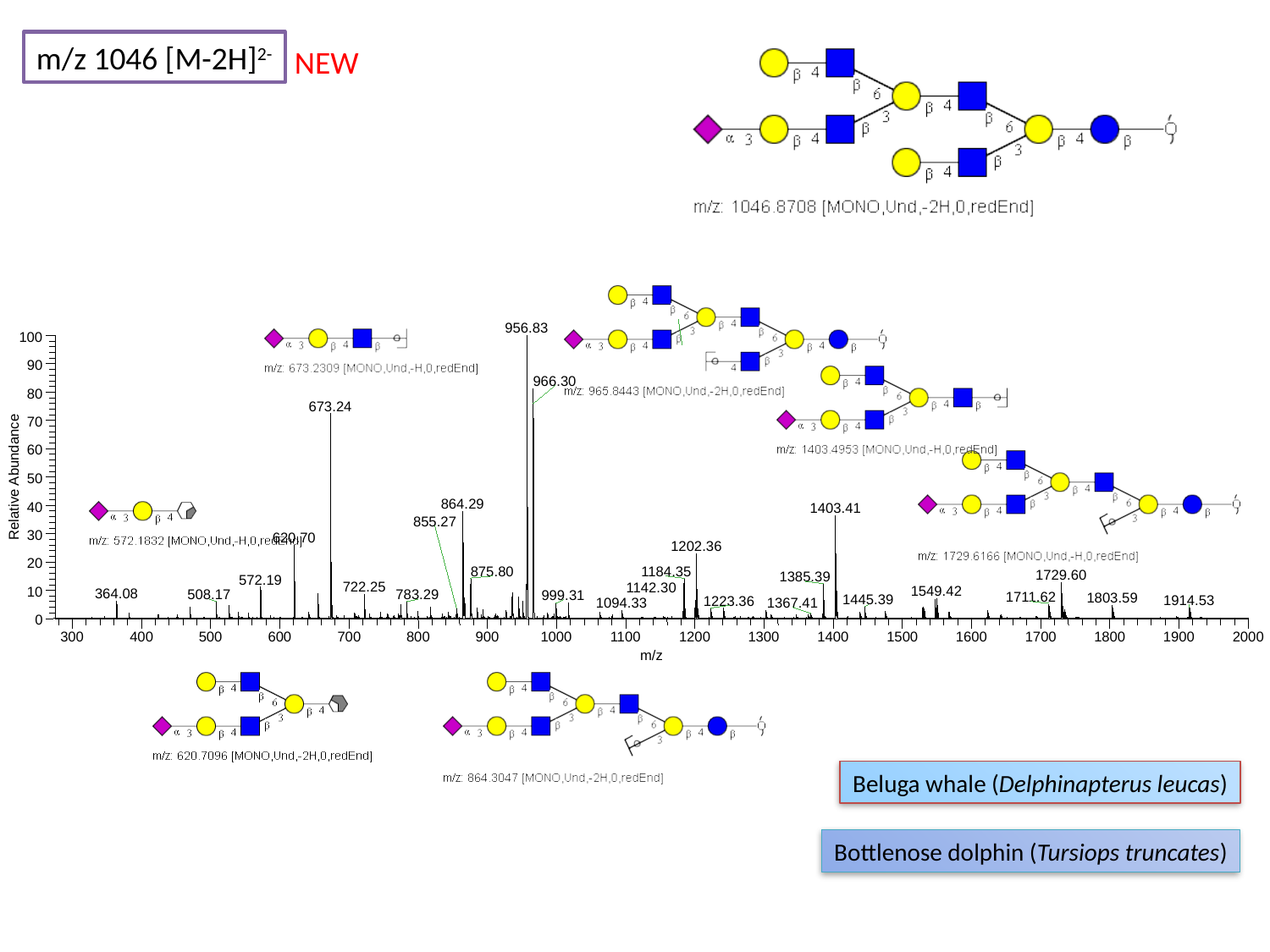

m/z 1046 [M-2H]^2-^
NEW
956.83
100
90
966.30
80
673.24
70
60
Relative Abundance
50
864.29
40
1403.41
30
620.70
1202.36
20
1184.35
1729.60
1385.39
10
0
300
400
500
600
700
800
900
1000
1100
1200
1300
1400
1500
1600
1700
1800
1900
2000
m/z
855.27
875.80
572.19
722.25
1142.30
1549.42
364.08
508.17
783.29
999.31
1711.62
1803.59
1445.39
1914.53
1223.36
1367.41
1094.33
Beluga whale (Delphinapterus leucas)
Bottlenose dolphin (Tursiops truncates)

### Slide 171
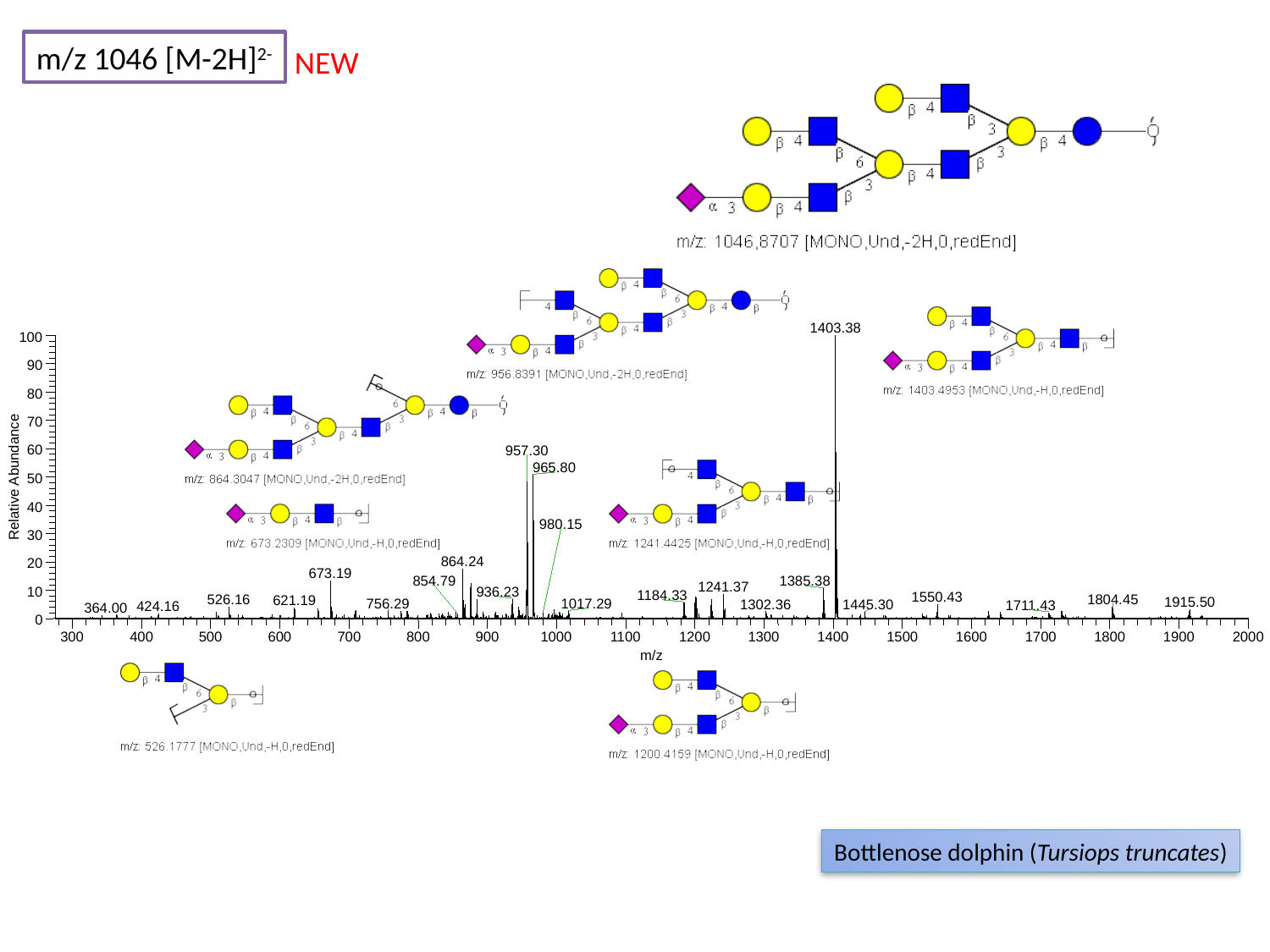

m/z 1046 [M-2H]^2-^
NEW
1403.38
100
90
80
70
60
957.30
965.80
Relative Abundance
50
40
30
864.24
20
673.19
1385.38
1241.37
10
1184.33
0
300
400
500
600
700
800
900
1000
1100
1200
1300
1400
1500
1600
1700
1800
1900
2000
m/z
980.15
854.79
936.23
1550.43
1804.45
526.16
621.19
1915.50
756.29
1017.29
1302.36
1445.30
1711.43
424.16
364.00
Bottlenose dolphin (Tursiops truncates)

### Slide 172
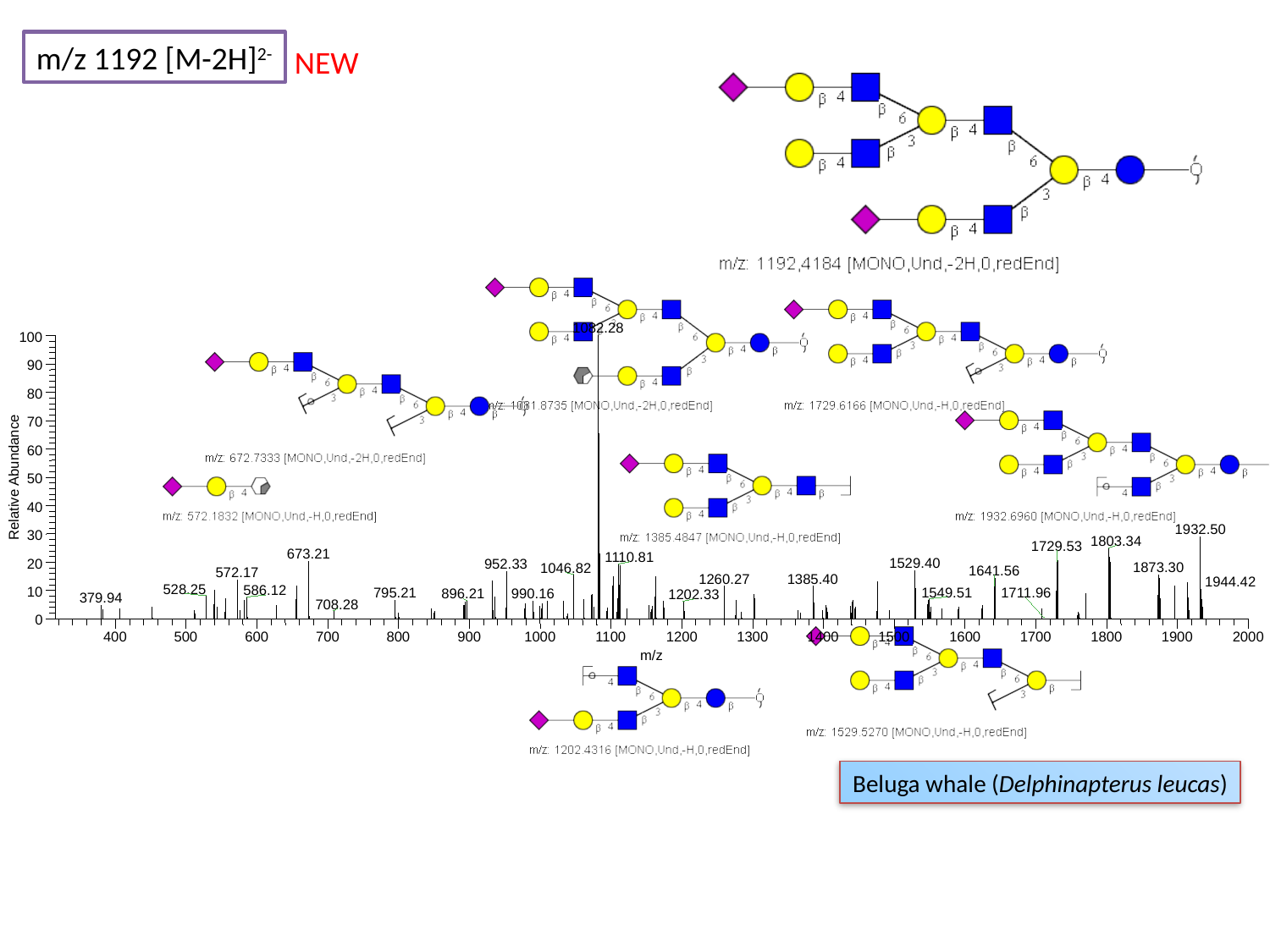

m/z 1192 [M-2H]^2-^
NEW
1082.28
100
90
80
70
60
Relative Abundance
50
40
1932.50
30
1803.34
1729.53
673.21
1110.81
1529.40
20
952.33
1873.30
1046.82
572.17
10
0
400
500
600
700
800
900
1000
1100
1200
1300
1400
1500
1600
1700
1800
1900
2000
m/z
1641.56
1260.27
1385.40
1944.42
528.25
586.12
1549.51
795.21
1711.96
990.16
896.21
1202.33
379.94
708.28
Beluga whale (Delphinapterus leucas)
